# High-resolution mapping of heteroduplex DNA formed during UV-induced and spontaneous mitotic recombination events in yeast

**DOI:** 10.1101/131995

**Authors:** Yi Yin, Margaret Dominska, Eunice Yim, Thomas D. Petes

**Affiliations:** Department of Molecular Genetics and Microbiology and University Program in Genetics and Genomics, Duke University Medical Center, Durham, NC 27710, USA

**Author notes:** Corresponding author: Thomas D. Petes Department of Molecular Genetics and Microbiology Box 3054, Duke University Medical Center Durham, NC 27710. Current address: Thomas D. Petes, Department of Molecular Genetics and Microbiology, Box 3054, Duke University Medical Center, Durham, NC 27710.

**Keywords:** mitotic recombination, mismatch repair, heteroduplex DNA, gene conversion, ultraviolet light

## Abstract

Double-stranded DNA breaks (DSBs) can be generated by both endogenous and exogenous agents. In diploid yeast strains, such breaks are usually repaired by homologous recombination (HR), and a number of different HR pathways have been described. An early step for all HR pathways is formation of a heteroduplex, in which a single-strand from the broken DNA molecule pairs with a strand derived from an intact DNA molecule. If the two strands of DNA are not identical, within the heteroduplex DNA (hetDNA), there will be mismatches. In a wild-type strain, these mismatches are removed by the mismatch repair (MMR) system. In strains lacking MMR, the mismatches persist and can be detected by a variety of genetic and physical techniques. Most previous studies involving hetDNA formed during mitotic recombination have been restricted to a single locus with DSBs induced at a defined position by a site-specific endonuclease. In addition, in most of these studies, recombination between repeated genes was examined; in such studies, the sequence homologies were usually less than 5 kb. In the present study, we present a global mapping of hetDNA formed in a UV-treated MMR-defective *mlh1* strain. Although about two-thirds of the recombination events were associated with hetDNA with a continuous array of unrepaired mismatches, in about one-third of the events, we found regions of unrepaired mismatches flanking regions without mismatches. We suggest that these discontinuous hetDNAs involve template switching during repair synthesis, repair of a double-stranded DNA gap, and/or Mlh1-independent MMR. Many of our observed events are not explicable by the simplest form of the double-strand break repair (DSBR) model of recombination. We also studied hetDNA associated with spontaneous recombination events selected on chromosomes IV and V in a wild-type strain. The interval on chromosome IV contained a hotspot for spontaneous crossovers generated by an inverted pair of transposable elements (HS4). We showed that HS4-induced recombination events are associated with the formation of very large (>30 kb) double-stranded DNA gaps.

## Introduction

Homologous recombination (HR) is important for repairing double-stranded DNA breaks (DSBs) in diploid yeast. Although three different HR pathways have been described (Symington *et al*., 2014), the earliest steps of all three pathways have a common intermediate in which a single DNA strand from one duplex invades the homologous template (green boxed region of Fig. 1). In Fig. 1, chromosomes are shown as double-stranded DNA structures with the two homologs drawn in different colors. Following the DSB, the broken ends are processed by 5’ to 3’ degradation in a two-step process with limited resection (about 100 bases) performed by Sae2 and the Mre11-Rad50-Xrs2 proteins, followed by more extensive resection (>1 kb) utilizing the redundant pathways of Exo1 or Dna2 with the Sgs1-Top3-Rmi1 complex (Symington, 2014). The 3′ single strand derived from the broken end then invades the unbroken template chromosome forming a heteroduplex, a region of duplex DNA derived from the different DNA molecules. In Fig. 1, heteroduplexes are shown as paired strands of different colors.

**Figure 1.**
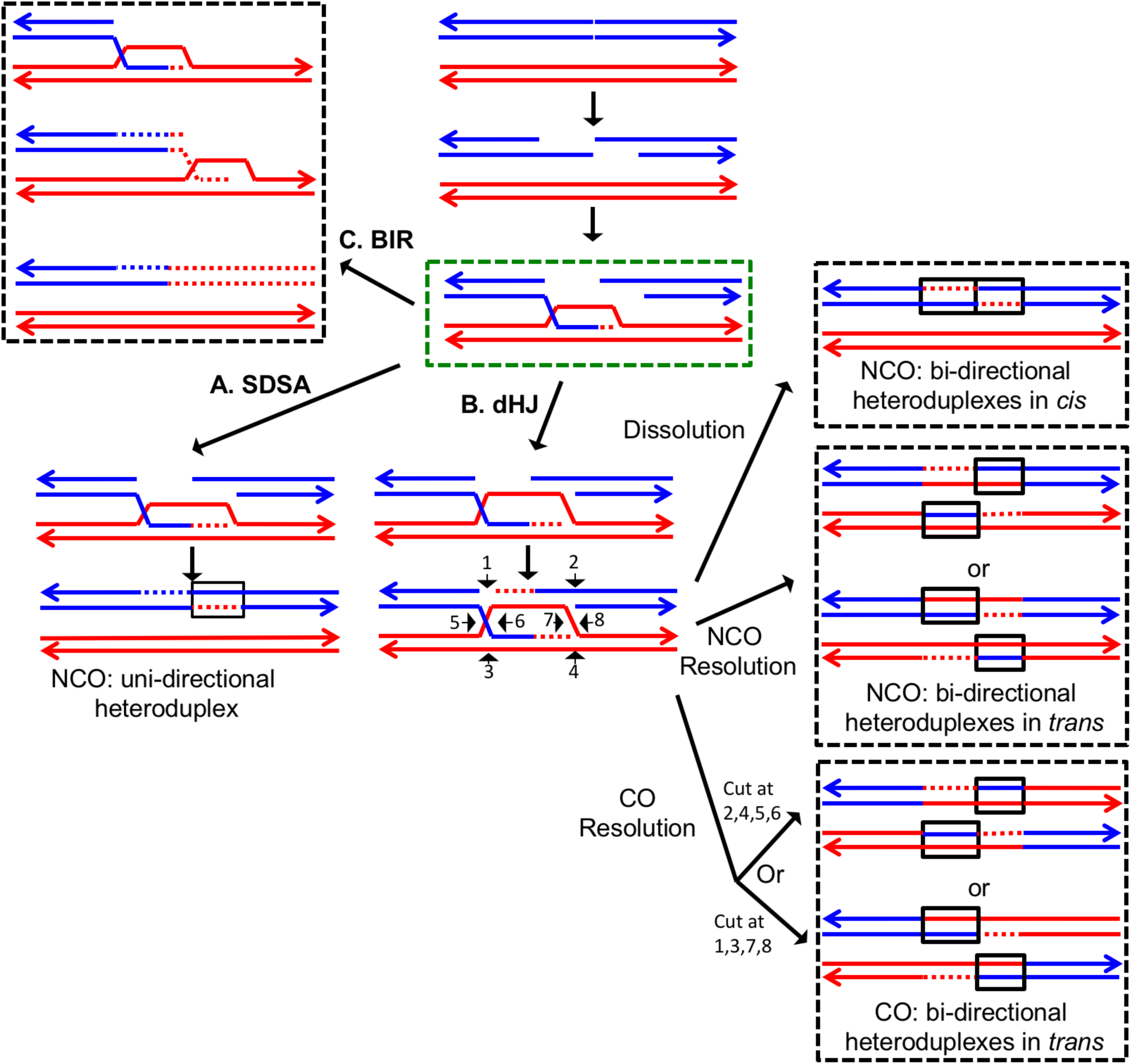
The DSBR repair pathways. The chromatids are shown as double-stranded molecules with arrows at the 3′ ends. The broken chromatid is colored blue and the intact template chromatid is red. Dotted lines indicate newly-synthesized DNA. Regions of heteroduplex DNA (hetDNA) are outlined by black boxes. All pathways are initiated by invasion of one processed broken end into the unbroken chromatid, forming a D-loop. **A.** Synthesis-dependent strand annealing (SDSA). Following DNA synthesis primed by the invading 3′ end, the invading end dissociates from the intact molecule, and re-anneals with the second broken end. The net result is an NCO event with hetDNA extending uni-directionally from the break. **B.** Double Holliday junction (dHJ) intermediate. Following DNA synthesis from one invading end, the second broken end pairs with the D-loop (second-end capture). formed in canonical double strand break repair (DSBR) pathway after second end capture. This intermediate can be dissolved by migrating the two junctions inwards, followed by decatenation; this pathway results in a NCO with heteroduplexes located in *cis* on one chromatid. Alternatively, it can also be resolved by cutting the junctions symmetrically to generate NCOs (cuts at arrows marked 1, 2, 3, and 4 or 5, 6, 7, and 8), or asymmetrically (cuts at arrows marked 2, 4, 5, and 6 or 1, 3, 7 and 8) to generate COs. For both types of resolution, the heteroduplexes are located in *trans* on the two chromatids, flanking the DSB site. Note that cuts at positions 1, 3, 7, and 8 reflect nick-directed resolution of the junctions. **C.** Break-induced replication (BIR). In this pathway, one broken end invades the intact template and copies DNA sequences by conservative replication to the end of the template. The other broken end is lost. Except for the initial strand invasion, heteroduplex intermediates are not relevant to this pathway.

For all three pathways, the invading 3’ end is used as a primer to catalyze DNA synthesis (shown as red dotted lines). The subsequent steps differ for each pathway. In the synthesis-dependent strand-annealing (SDSA) pathway, following DNA synthesis, the invading strand is displaced and reanneals to the other processed broken end. Repair of the single-stranded gap (dotted blue line) completes the event. In the SDSA pathway, a region of heteroduplex occurs on only one of the two homologs, and SDSA events are not associated with crossovers.

In the double-strand-break-repair (DSBR) pathway, the strand displaced by DNA synthesis from the invading strand pairs with the other broken end. Repair synthesis results in formation of a double Holliday junction (dHJ) intermediate. There are several mechanisms to resolve dHJs. First, the two junctions could be migrated toward each other, and decatenated by the Sgs1p-Top3p-Rmi1p complex (dissolution). By this mechanism, there are two regions of hetDNA flanking the DSB, both located on the molecule that was originally broken (*cis* hetDNA).

Alternatively, the dHJ could be resolved by HJ resolvases such as Mus81p and Yen1p to form either crossover (CO) or non-crossover (NCO) products. Resolution of both HJs in the same orientation (for example, cleavage of both junctions as shown by the horizontal arrows terminated by circles) results in NCO products, whereas cleavage in different orientations (one HJ cleaved as shown with vertical arrows terminated by diamonds, and the other cleaved as shown with horizontal arrows) results in CO products (Fig. 1). For both types of resolution, the hetDNA is on both sides of the DSB, and both the donor and recipient DNA molecule have hetDNA (*trans* hetDNA).

In the last pathway of HR, break-induced replication (BIR), the invading strand results in a migrating D-loop replication structure that duplicates chromosomal sequences conservatively from the point of strand invasion to the end of the chromosome (Donnianni and Symington, 2013; Saini *et al*., 2013). In this pathway, the second part of the broken DNA is lost, and no heteroduplex is associated with either the donor or recombinant DNA molecule.

If the two strands of DNA that compose the heteroduplex have non-identical sequences, the heteroduplex will contain mismatches (Fig. 2). Correction of these mismatches by the mismatch enzymes results in gene conversion, the non-reciprocal transfer of information between the two chromosomes (Symington *et al*., 2014). In Fig. 2A, the heteroduplex has four mismatches, if these mismatches are excised from the “blue” strand, and the resulting gap is filled in using the “red” strand as a template, a conversion event is observed. Alternatively, if the mismatches are removed from the red strand, no conversion would be observed; such events are termed “restorations”. A third possibility is that the mismatches would be excised from both strands resulting in “patchy” repair. In strains that lack MMR, mismatches are not repaired and two non-identical daughter cells are produced (Fig. 2B). In MMR-proficient strain, the patterns of heteroduplex formation may be obscured. For example, in the DSBR pathway, following MMR the *cis* and the *trans* pattern of heteroduplexes (Fig. 1) are indistinguishable because both yield the same type of conversion tracts (Fig. 2C and 2D).

**Figure 2.**
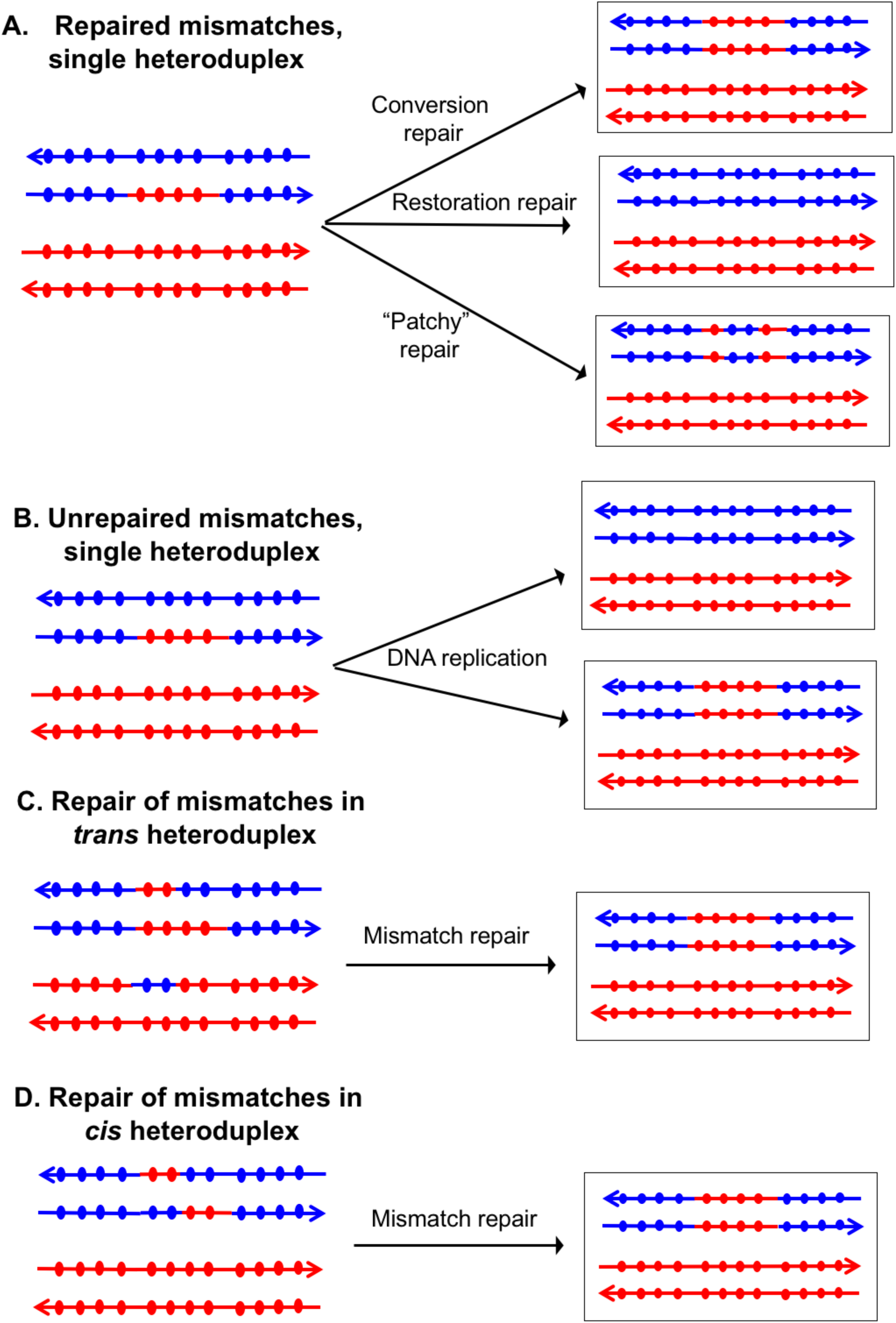
Patterns of mismatch repair in a MMR-proficient strain. Chromatids are shown as double-stranded DNA structures with circles indicating SNPs. For all parts of the figure, the blue chromatid is broken and, consequently, is the recipient of sequence information from the red chromatid. For these examples, all intermediates were resolved as NCO events. In Figs. 2A, 2C, and 2D, the products shown on the right side of each panel are a consequence of mismatch repair in the mother cell. In Fig. 2B, the products are shown in two daughter cells following replication of the chromosomes of the mother cell. A. Repair of mismatches within a single heteroduplex. In the top panel, the mismatches in the heteroduplex are repaired using the bottom strand as a template, resulting in a conversion event. In the middle panel, the upper strand is used as a template for MMR, resulting in a blue chromatid that is identical to an unbroken chromatid. In the bottom panel, some mismatches undergo conversion-type repair and others restoration-type repair. B. Loss of mismatches as a consequence of DNA replication. Replication of a chromatid with a heteroduplex results in one product that appears to have undergone conversion-type repair, and a second that is the same as a chromatid without a recombination event. C. Repair of mismatches in a *trans* heteroduplex. If mismatches within the two heteroduplexes are repaired using the red strand as a template, a conversion event that is identical to that shown in the upper panel of Fig. 2A would be generated. D. Repair of mismatches in a *cis* heteroduplex. If all mismatches are repaired using the red strand as a template, the resulting conversion product is identical to that shown in Figs. 2A and 2C.

Mitchel *et al*. (2010) examined patterns of heteroduplex formation in recombination events between a linearized plasmid and a chromosome in an MMR-deficient strain. There were multiple sequence differences spaced about 50 bp apart within an 800 bp region of homology. By sequencing the single-colony transformants, they could detect the unrepaired mismatches that defined the length of the heteroduplex. From their analysis, they concluded: 1) most of the heteroduplexes unassociated with crossovers (NCO) were restricted to one side of the DSB site, consistent with their formation by the SDSA pathway rather than the DSBR pathway, 2) most of the heteroduplexes associated with crossovers (CO) had the patterns consistent with the DSBR pathway, and 3) most of the heteroduplexes extended less than 500 bp from the DSB site on the plasmid. In general, the observations of Mitchel *et al*. were consistent with the predictions of the model described in Fig. 1.

In previous studies, we developed methods of detecting and mapping spontaneous and UV-induced crossovers and associated gene conversion events throughout the genome (St. Charles *et al*., 2012; St. Charles and Petes, 2013; Yin and Petes, 2013). Our analysis utilized diploids that were heterozygous for about 55,000 single-nucleotide polymorphisms (SNPs) and microarrays capable of distinguishing heterozygous SNPs and homozygous SNPs. One important conclusion from this analysis was that mitotic gene conversion tracts (median length of 11 kb, St. Charles and Petes, 2013) were generally much longer than meiotic conversion tracts (median length of about 2 kb; Mancera *et al*., 2008). This observation suggests that either heteroduplexes are much longer in mitosis than in meiosis or that long mitotic gene conversion tracts involve a different intermediate such as a large double-stranded DNA gap. Our previous experiments were done in MMR-proficient strains. In the current study, using *mlh1* (MMR-deficient) diploids, we show that some mitotic recombination events involve long heteroduplexes, although other events are consistent with gap repair. In addition, however, we find that recombination events initiated at a previously-described spontaneous mitotic recombination hotspot caused by an inverted pair of retrotransposons (St. Charles and Petes, 2013; Yim *et al*., 2014) often involve long (> 20 kb) double-stranded gaps rather than very long heteroduplexes. We previously showed that most spontaneous crossovers between homologs had patterns of gene conversion that indicate that recombinogenic DSBs occur in unreplicated DNA (Lee *et al*., 2009; Lee and Petes, 2010). Our current study is consistent with this conclusion.

In our previous mapping of gene conversion tracts in MMR-proficient strains, we found that about 20% of the UV-induced and spontaneous events (St. Charles and Petes, 2013; Yin and Petes, 2013) were “patchy” in which markers exhibiting gene conversion flanked markers that were not converted. One simple explanation of this observation is that mismatches within one heteroduplex are sometimes repaired in a non-concerted manner by conversion-type repair and restoration-type repair. If this explanation is correct then, in the absence of repair, mismatches within a heteroduplex should be continuous (Reyes *et al*., 2015). However, in the *mlh1* strain used in this study, we find that about one-third of heteroduplexes contain mismatches flanking regions without mismatches (discontinuous heteroduplexes). Such patterns can be explained by Mlh1-independent MMR, and/or template-switching during repair synthesis. We also find recombination events that are inconsistent with the canonical DSBR models, and require additional steps such as branch migration of the Holliday junctions and/or sequential invasion of broken DNA ends. Our results argue that mitotic recombination often occurs through pathways that are considerably more complex than those shown in Fig. 1. Similar conclusions have been reached through the analysis of meiotic (Mancera *et al*., 2008; Martini *et al*., 2011) and mitotic (Hum and Jinks-Robertson, accompanying paper) recombination events in yeast.

## Results

### Experimental system

The system that we previously used to analyze mitotic recombination in MMR-proficient strains is illustrated in Fig. 3. The experimental diploids were homozygous for the *ade2-1* ochre mutation located on chromosome XV. In the absence of a nonsense suppressor, such strains form red colonies. The chromosome arm to be assayed for recombination had a heterozygous insertion of the *SUP4-o* gene encoding an ochre suppressor located near the telomere. Diploids with zero, one, and two copies of *SUP4-o* form red, pink, and white colonies, respectively.

**Figure 3.**
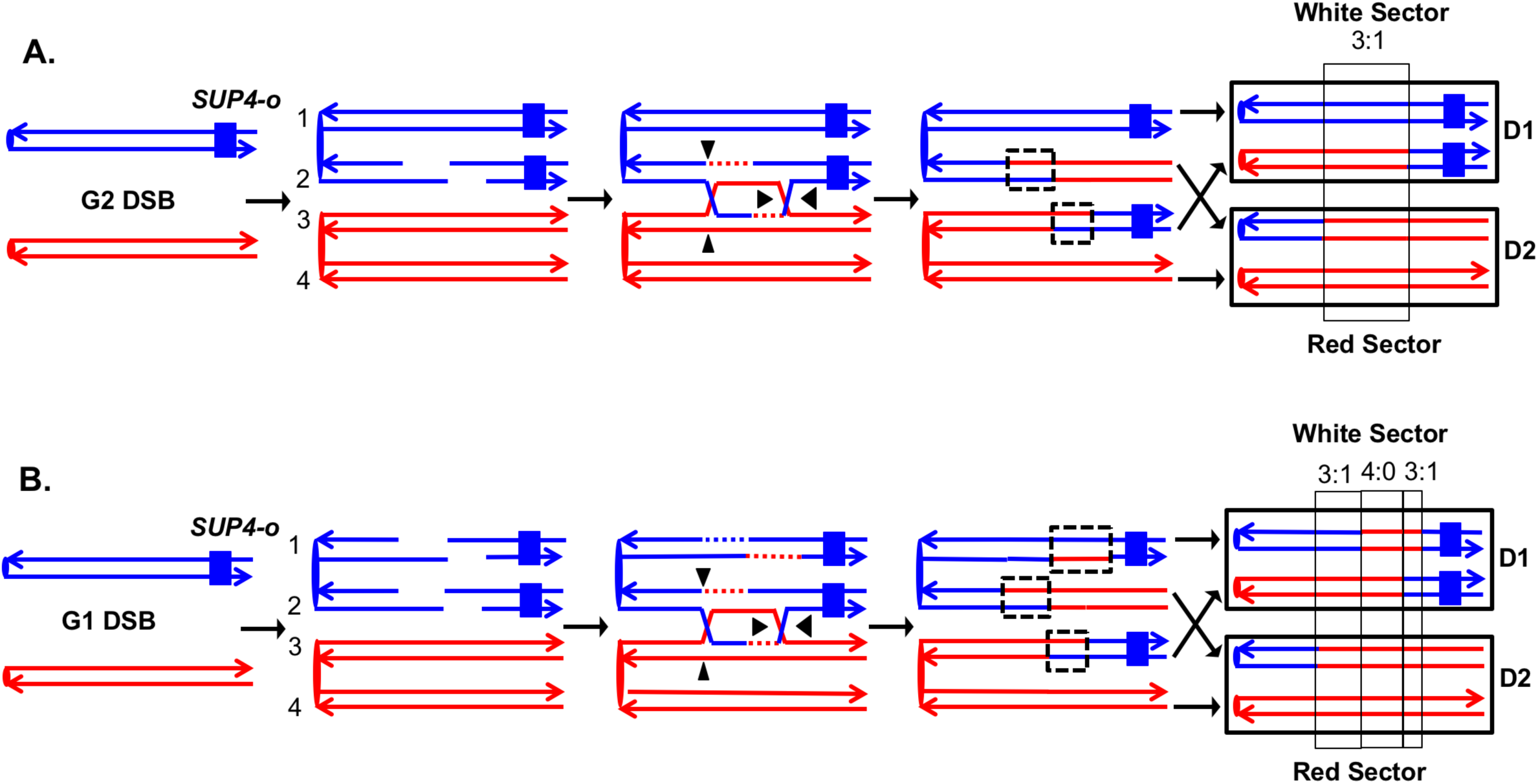
Detection and analysis of crossovers induced by DSBs in single chromatids or in unreplicated chromosomes in an MMR-proficient diploid. The lines show the two strands of each chromosome/chromatid, and ovals indicate centromeres. The strain is homozygous for *ade2-1*, an ochre allele that, in the absence of the *SUP4-o* ochre suppressor, forms a red colony. In the strains used in our study, one copy of *SUP4-o* is inserted near the telomere of one homolog. Strains with zero, one or two copies of the suppressors form red, pink and white colonies, respectively. Black triangles indicate the positions of cleavages of the dHJ. Heteroduplex intermediates are enclosed by dotted black lines, and conversion tracts are enclosed in thin black lines. Chromosomes in daughter cells are outlined by thick black rectangles, and D1 and D2 denote the two daughter cells resulting from the crossover. A. Crossover initiated by a single chromatid break (SCB). Following the crossover, segregation of one recombined and one parental chromatid into each cells will generate one cell homozygous for *SUP-o* and one cell lacking *SUP4-o*; subsequent divisions will lead to a red/white sectored colony. If mismatches in both heteroduplexes undergo conversion-type repair, a 3:1 conversion event would be observed; within the boxed region, three chromatids have information derived from the red chromatid and one has information derived from the blue chromatid. B. Crossover initiated by a DSB in an unreplicated blue chromosome, resulting in double sister-chromatid breaks (DSCB). DNA replication of the broken chromosome would result in,two broken sister chromatids. We show the middle pair of chromatids repaired as a crossover, and the top chromatid repair by an SDSA event. In this example, we show the heteroduplex associated with the SDSA event as longer than those of the CO-associated hetDNAs. If all of mismatches are repaired as conversion events, we would see a hybrid 3:1/4:0/3:1 conversion tract. The 4:0 pattern is diagnostic of a DSCB event.

Therefore, prior to the recombination event, cells have one copy of *SUP4-o* and form pink colonies. A cell that has a reciprocal crossover between the centromere and the *SUP4-o* marker will form a red/white sectored colony, assuming that both daughter cells have one recombinant and one non-recombinant chromosome (Fig. 3A). In addition to the heterozygous *SUP4-o* marker, the diploids were heterozygous for about 55,000 single-nucleotide polymorphisms (SNPs) because they were constructed by mating two sequence-diverged haploids, W303-1A and YJM789 (Lee *et al*., 2009; St. Charles *et al*., 2012). To determine the position of the crossover and to detect conversion events associated with the crossover, we used microarrays in which loss of heterozygosity (LOH) could be detected for about 13,000 SNPs. Each SNP was represented by four 25-base oligonucleotides, two containing the Watson and Crick strands of the W303-1A allele and two containing the Watson and Crick strands of the YJM789 allele. By hybridizing genomic DNA derived from the red and white sectors, we could map the position of the crossover and associated gene conversion tract (St. Charles *et al.*, 2012). Details of this analysis are provided in Materials and Methods.

In our previous analysis, we examined spontaneous recombination events in two types of MMR-proficient strains: those with the *SUP4-o* marker located about 100 kb from *CEN5* and those with the *SUP4-o* marker located about 1 Mb from *CEN4*. Most of the crossovers were associated with gene conversion events (solid-line boxes in Fig. 3). Although some of the sectored colonies had the expected pattern of conversion in which one daughter cell is homozygous for SNPs located near the crossover event and the daughter is heterozygous (Fig. 3A), more than half of the crossovers were associated with conversion events in which both daughter cells were homozygous for SNPs derived from one homolog (region marked 4:0 in Fig. 3B). This pattern indicates that two sister chromatids were broken at the same position, consistent with a model in which the recombinogenic DSB occurs in G_1_, and the broken chromosome is replicated to produce two broken sister chromatids. The 4:0 tracts are often adjacent to 3:1 tracts as in Fig. 3B. We interpret such hybrid tracts as reflecting the repair of two sister chromatids in which the extent of heteroduplex formation is different for the repair of each DSB (Lee *et al*., 2009; St. Charles *et al*., 2012).

To examine mismatch-containing heteroduplexes, we constructed two diploids in the hybrid genetic background that lacked the mismatch repair protein Mlh1p (details in Table S1). The heterozygous insertion of *SUP4-o* required for the sectoring assay was located near the right end of chromosome IV in diploid YYy311, and near the left end of chromosome V in diploid YYy310. In wild-type strains in which mismatches within the heteroduplex are corrected by the MMR enzymes, each sector will contain cells of only one genotype (Fig. 3). In contrast, in the MMR-deficient strains, there may be two different genotypes in one or both sectors. In Fig. 4A, we show the patterns of heteroduplex formation expected by the DSBR pathway in a MMR-deficient cell in which recombination initiates as a consequence of a single broken chromatid in G_2_ of the cell cycle. Both the D1 and D2 daughter cells retain heteroduplexes with mismatches. In Fig. 4A, the cells derived from D1 are in the white sector, and those derived from D2 are in the red sector. When the chromosomes in D1 are replicated, two different genotypes will be observed in the granddaughter cells, GD1-1 and GD 1-2. In addition, if the heteroduplexes are in the *trans* configuration as expected for the DSBR pathway, there will also be two different genotypes in cells derived from D2.

**Figure 4.**
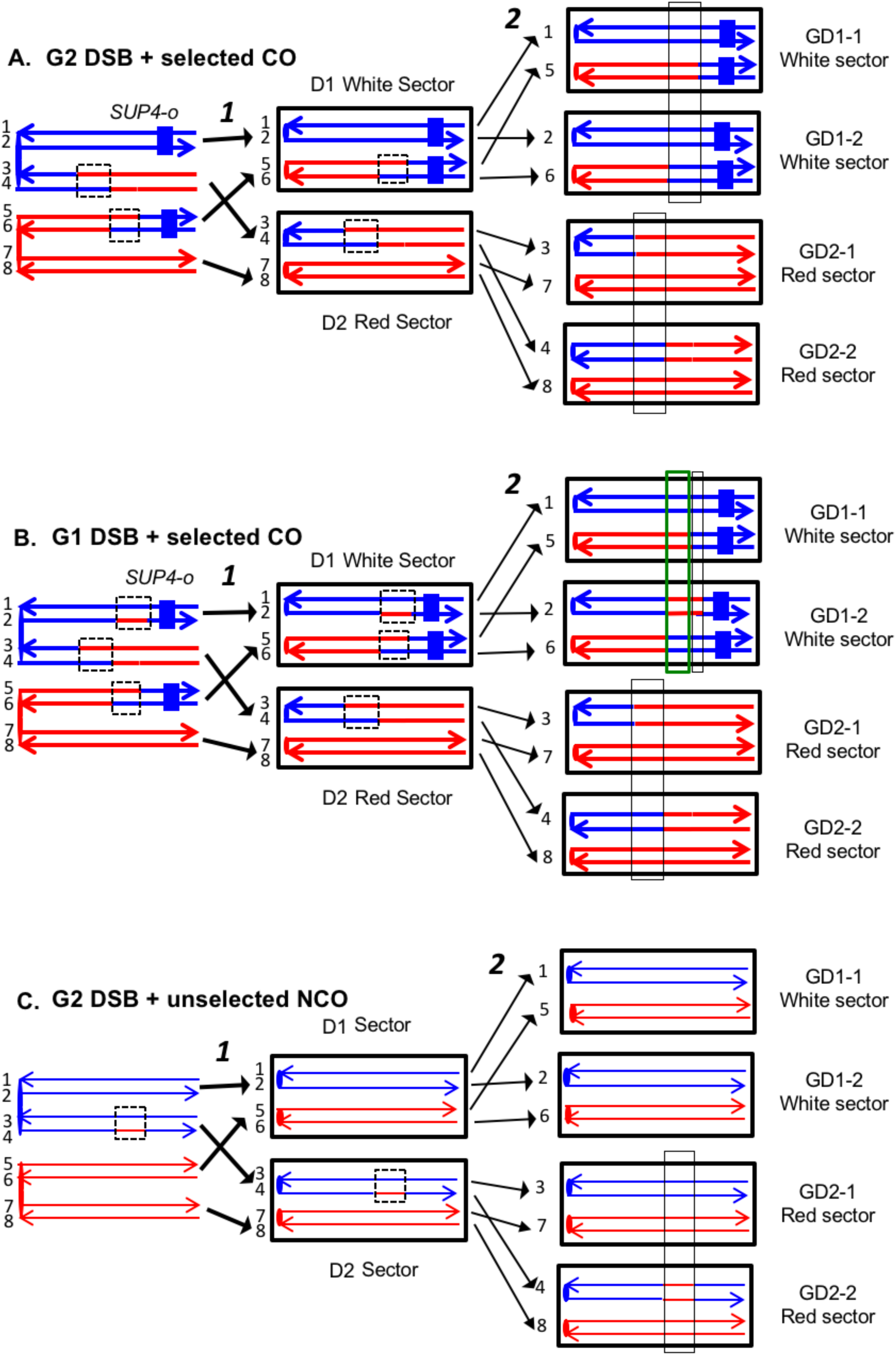
Expected recombination-associated segregation patterns of SNPs into daughter and granddaughter cells in MMR-deficient strains following a G_2_- or G_1_-induced DSB. We show heteroduplexes with unrepaired mismatches (outlined in dashed lines) in the daughter cells D1 and D2. Replication of unrepaired mismatches in heteroduplexes results in granddaughter cells with two different genotypes; these differences are outlined in the right side of the figure by thin black lines. The events shown in Figs. 4A and 4B were selected because the crossovers between the heterozygous *SUP4-o* marker generated daughter cells that had two copies (resulting in a white sector) or no copies (resulting in a red sector) of *SUP4-o*. The NCO event shown in Fig. 4C was on a homolog that did not contain the *SUP4-o* marker (shown in thin red and blue lines) and was unselected. A. CO-associated SCB (selected crossover). If the crossover occurred by the DSBR pathway, we would expect that both sectors would have granddaughter cells with different SNP patterns. B. CO-associated DSCB (selected crossover). As in Fig. 4A, both sectors would have granddaughter cells with different SNPs. One distinguishing feature of the DSCB is that both granddaughter cells of one sector (GD1-1 and GD1-2) have different coupling relationships for SNPs in the same region as outlined in the green rectangle. C. NCO-associated SCB (unselected). Strains treated with UV have many unselected events. Since these events are induced by UV at the same time as the selected event, we can use the information identifying GD1-1, GD1-2. GD2-1, and GD2-2 from the selected event to determine the patterns of heteroduplex formation for the unselected event by using whole-genome SNP arrays. In the depicted event, only one of the sectors had granddaughters with different genotypes.

On the left side of Fig. 4B, we show the pattern of heteroduplexes expected for an event initiated by a DSB in an unreplicated blue chromosome (producing two sister chromatids broken at the same position as the result of DNA replication) that was repaired in G_2_ of the cell cycle. Such cells can also produce two genotypes in each sector (Fig. 4B). In the white sector, two types of granddaughter cells with different genotypes are shown. In the region shown in the black rectangle, GD1-1 is homozygous for the blue-derived SNPs and GD1-2 is heterozygous. In addition, in the regions defined by the green rectangles for GD1-1 and GD1-2, although both genotypes are heterozygous, the coupling of the heterozygous regions is different. In GD1-1, the blue and red SNPs within the green rectangle are on the same chromosome as the centromere-proximal blue and red SNPs, whereas in GD1-2, the blue and red SNPs within the rectangle are coupled to the centromere-proximal red and blue SNPs, respectively. As described below, for our analysis, we looked for different genotypes within sectors by methods that allowed us to examine the locations of heterozygous and homozygous SNPs, and to determine the coupling of heterozygous regions. It is important to emphasize that there is a one-to-one correspondence between the chromosomes within the sectors (labeled 1-8 on the right side of the figure) and the DNA strands in the mother cell in which the recombination event occurred (strands labeled 1-8 on the left side of the figure).

Most of our data were obtained from G_1_-synchronized YYy310 cells treated with 15 J/m^2^ of UV. This treatment stimulates mitotic recombination on chromosome V >10^3^-fold and results in about ten unselected events on other chromosomes (Yin and Petes, 2013). To detect cells of different genotypes within each sector of red/white sectored colonies (reflecting a crossover on chromosome V), we first purified 10-20 white and red colonies from each sector. Initially, we performed a SNP-specific microarray array of genomic DNA isolated from one of the white colonies to locate the approximate position of the recombination event. For example, we analyzed one white colony (YYy310.9-5W1) derived from a sectored red/white colony number 5 derived from UV-treated YYy310.9 cells. In Fig. 5, the red and blue circles represent hybridization to W303-1A-and YJM789-derived-oligonucleotides, respectively. Hybridization values (Y-axis) are normalized such that a value of about 1 indicates that the experimental DNA sample is heterozygous for the W303-1A- and YJM789-derived-alleles (additional details in Materials and Methods); the X-axis shows Saccharomyces Genome Database (SGD) coordinates on chromosome V. LOH events that duplicate the sequences from one allele and remove sequences from the other are associated with hybridization values of about 1.5 and 0.3, respectively. Thus, by microarray analysis (Fig. 5A), the strain (a granddaughter derived from a white sector) shown in Fig. 5A was homozygous for YJM789-derived SNPs from the left end of chromosome V to SGD coordinate 51915. The strain was heterozygous for SNPs between 53612 to 54198, and then homozygous for W303-1A-derived SNPs from coordinates 56117 to 57170. Lastly, the strain was heterozygous for SNPs between coordinates 60701 and *CEN5*. As will be discussed below, this strain represents one of the two granddaughter genotypes observed in the white sector.

**Figure 5.**
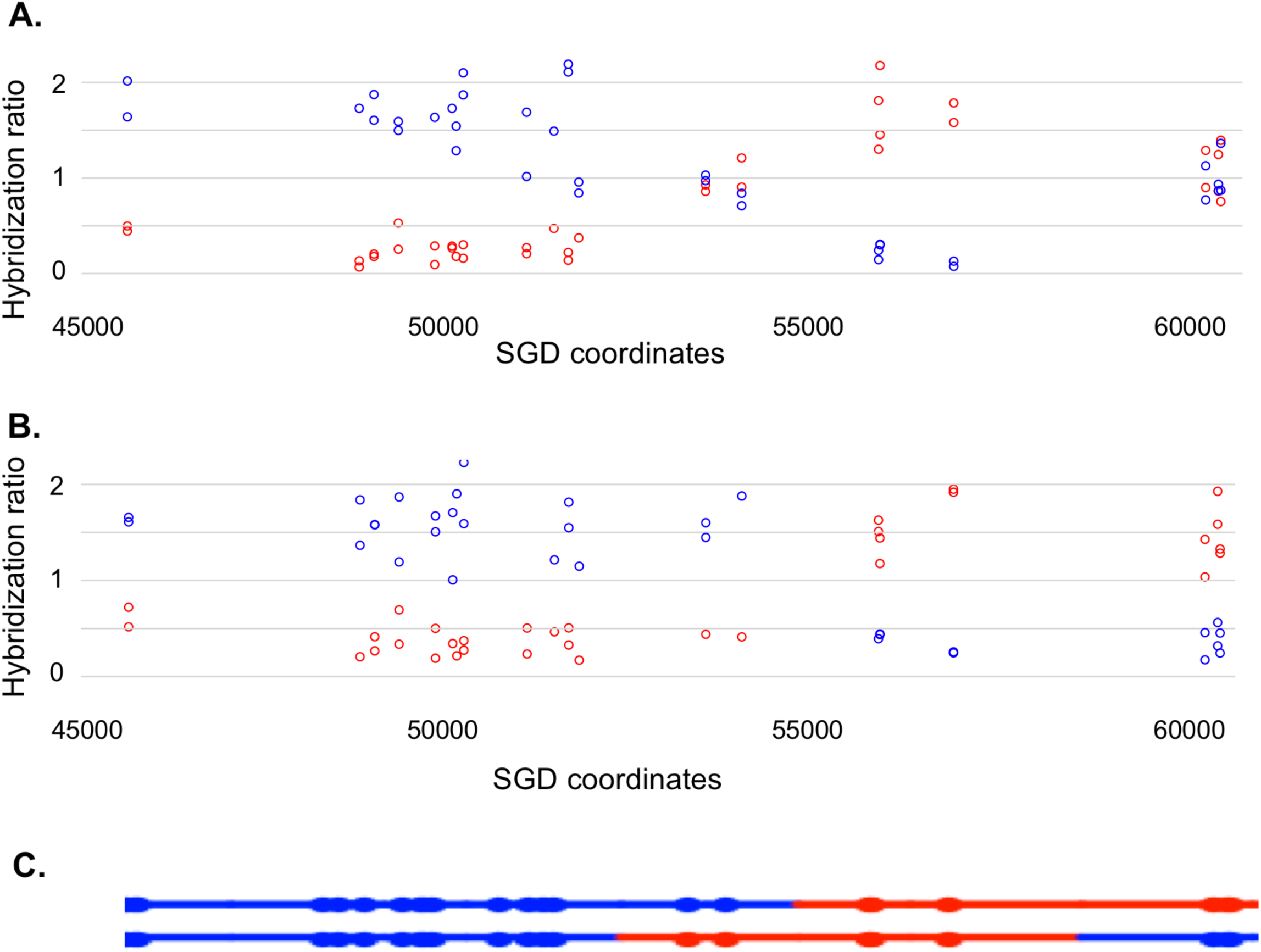
Microarray analysis of one granddaughter cell in a sectored colony resulting from a recombination event on chromosome V. By methods described in the text, we identified two granddaughter genotypes associated with the red and white sectors of the YYy310-9-5WR sectored colony. SNP-specific microarrays were done on genomic DNA isolated from all four granddaughter strains, and from spore derivatives of each of these granddaughters. A. Microarray analysis of the W1 granddaughter strain. Genomic DNA was isolated, and hybridized to SNP-specific microarrays in competition with differentially-labeled genomic DNA from a heterozygous control strain (details in Materials and Methods). The Y axis shows the hybridization ratio of the experimental samples to allele-specific SNPs normalized to the hybridization levels of the control heterozygous sample; blue and red circles indicate hybridization ratios to YJM789- and W303-1A-specific SNPs, respectively. Heterozygous samples have a ratio of about 1, samples in which the strain-specific allele is present in two copies have a ratio of about 1.7, and those in which the strain-specific allele is missing have a ratio of about 0.3. The × axis has SGD coordinates for chromosome V in the region of the recombination event. Between the left telomere (coordinate 1) and coordinate 51915, the W1 granddaughter is homozygous for the YJM789-derived SNPs, and heterozygous for SNPs between coordinates 60701 and the right telomere. In summary, chromosome V in this strain has a large terminal region that is homozygous for YJM789 SNPs, a short region of heterozygous SNPs, a short region that is homozygous for W303-1A SNPs, and a large heterozygous region that makes up the remainder of the chromosome. B. Microarray analysis of a spore derived from the W1 granddaughter strain. The W1 diploid was sporulated, and dissected. We examined the genomic DNA of one spore by SNP microarrays. From the pattern of SNPs in the spore, we conclude that one homolog in W1 has blue SNPs for the heterozygous region located near coordinates 55000 and red SNPs for the heterozygous region located between coordinates 60000, and the right telomere. The other homolog has the reciprocal pattern. C. Inferred arrangement of red and blue SNPs on the two homologs of W1 based on the microarray results shown in Figs. 5A and 5B. This pattern is reproduced in the top part of Fig. S8.

Based on this information, we examined genomic DNA from eight white colonies and eight red colonies by PCR analysis to look for different genotypes within one sector. We chose to examine SNPs that were near the transitions of heterozygous and LOH regions. For example, at position 51707, there is a polymorphism that distinguishes the YJM789- and W303-1A-derived SNPs (Lee *et al*., 2009). In the W303-1A genome, this polymorphism is part of a restriction enzyme recognition site for *Dra*l that is absent in YJM789. Consequently, we PCR-amplified the region containing this site from the individual white and red colonies, treated the resulting fragments with *Dra*l, and examined the resulting products by gel electrophoresis. Five of the white colonies were homozygous for the YJM789-form of the SNP, and three were heterozygous, demonstrating that this SNP was included as an unrepaired mismatch in one of the daughter cells. By a similar approach, we showed that SNPs heterozygous in the starting strain at coordinates 54915, 56166, and 57448 were homozygous for the W303-1A-form of the SNP in three red colonies, and heterozygous in the remaining five. Thus, this analysis allowed us to unambiguously define two different granddaughter genotypes in both the white (GD1-1, name shortened to W1 in Supplementary Figures; GD1-2, name shortened to W2) and red (GD2-1, name shortened to R1; GD2-2, name shortened to R2).

The complete analysis of the granddaughter genotypes required two additional steps. First, we mapped one representative of each granddaughter genotype using whole-genome SNP arrays. This analysis allows us to map heterozygous and homozygous SNPs on chromosome V and unselected events throughout the genome. The coordinates for transitions between heterozygous and homozygous SNPs for the UV-treated YYy310 samples are in Table S2. An example of the SNP microarray analysis for one granddaughter diploid strain derived from the white sector (YYy310.9-5W1) is shown in Fig. 5A.

Many of the granddaughter isolates, such as that shown in Fig. 5A, had two or more heterozygous regions. To examine the coupling relationships of the heterozygous regions, we sporulated the four granddaughter diploids derived from each sectored colony, and dissected tetrads. One spore derived from each strain was then examined by microarrays. The pattern shown in Fig. 5B represents the analysis of one haploid spore derived from granddaughter W1. This pattern shows the arrangement of SNPs on the top chromosome of Fig. 5C, and the arrangement of SNPs on the other homolog (bottom chromosome of Fig. 5C) can be directly inferred.

This analysis allowed a complete determination of the arrangement of SNPs in W1, W2, R1, and R2, and was performed for all sectored colonies. An example of how the patterns of LOH in the granddaughter cells can be used to figure out the pattern of heteroduplexes in the mother cell is shown in Fig. 6. Each chromosome in the granddaughter cells has the same pattern of SNPs as one of the DNA strands in the daughter cell. The source of the centromeres determines which of the chromosomes in the granddaughter cells was derived from which chromosome in the daughter cell. For example, in Fig. 6, the chromosomes labeled 1 and 2 have red centromeres, and were derived from the daughter chromosome with the red centromere. Chromosomes 5 and 6 were derived from the daughter chromosome with the blue centromere. Similarly, both the daughter chromosomes labeled 1 and 2 from the white sector and the daughter chromosomes labeled 3 and 4 from the red sector have red centromeres, and were connected as a pair of sister chromatids to the red centromere in the original mother cell.

**Figure 6.**
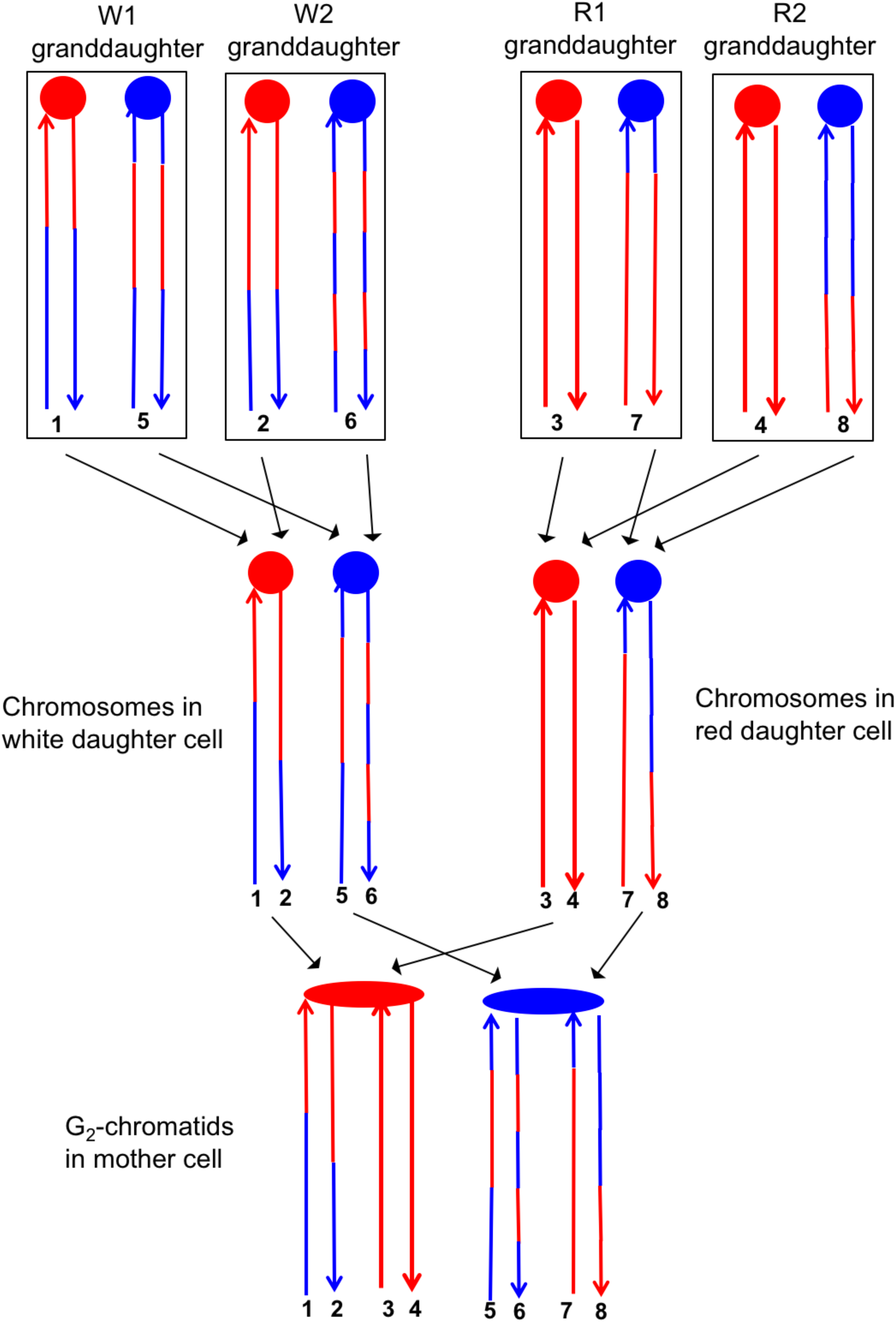
Correlation of SNP patterns in granddaughter cells with heteroduplex patterns in the mother cell. The example shown is the same crossover on chromosome V described in Fig. 5 (YYy310-9-5WR sectored colony). Each chromatid/chromosome is shown as a double-stranded DNA molecule with ovals and circles showing the centromeres. By the methods described in the text, we identified two genotypes within each sector. The microarray analysis illustrated in Fig. 5 allowed us to determine the patterns of LOH in each granddaughter cell (top of the figure). Each chromosome in the granddaughter cell was derived by replication by chromosomes in the daughter cells (middle of the figure). Chromosomes with the same color of centromere in the granddaughter cells must derive from a daughter chromosome with a centromere of the same color. For example, chromatids 1 and 2 in the W1 and W2 granddaughters must represent the strands labeled 1 and 2 in the white daughter cell. A similar procedure can be used to assign the strands to the chromosomes in all of the daughter cells. The chromosomes in the daughter cells represent segregants of the G_2_ chromatids in the mother cell. The daughter chromosomes with red centromeres must have been derived from the paired sister chromatids with the red oval centromere, and a similar conclusion can be drawn about the blue centromeres of the daughter chromosomes and the oval blue centromere in the mother cell. By the pattern of SNPs, we can also conclude that the chromatids in the mother cell with strands 1 and 2, and 7 and 8 were involved in a CO, whereas the other two chromatids were involved in the NCO mode of repair.

The transitions between regions of homozygous and heterozygous SNPs for all events are given in Table S2, and Table S3 summarizes the classes of these events as crossovers (CO) and non-crossovers (NCO). An important feature of our analysis is that once we have defined daughter and granddaughter cells based on the crossover on the selected chromosome, this lineage information also can be applied to unselected CO and NCO events on other chromosomes. For example, although the NCO event shown in Fig. 4C does not generate a sector, our whole-genome microarray analysis of granddaughter cells defined by the selected crossover allows us to detect and to fully describe the unselected event.

Our analysis of hetDNA included three types of experiments. First, we examined seven sectored colonies derived from the UV-treated YYy310 strain in which the *SUP4-o* marker was inserted near the left end of chromosome V. In addition to the selected crossover on chromosome V, these strains had about ten unselected events (both crossovers and conversions unassociated with crossovers) per strain because of the recombinogenic effects of UV (Yin and Petes, 2013). Second, we analyzed five spontaneous crossovers on chromosome V in YYy310. Lastly, we characterized six spontaneous crossovers on chromosome IV in strain YYy311 (heterozygous for the *SUP4-o* marker on the right end of chromosome IV) that were associated with the HS4 recombination hotspot.

### Analysis of selected crossovers on chromosome V and unselected recombination events induced by UV in YYy310

We examined seven sectored colonies derived from two isogenic independently-constructed YYy310 diploids (YYy310-9 and YYy310-10). By the methods described above, we identified two different genotypes within each sector in all seven sectored colonies; these genotypes are designated W1 and W2, and R1 and R2 for the white and red sectors, respectively. Our diagnosis of granddaughter genotypes was initially based on PCR analysis of SNPs associated with the selected crossover on chromosome V. For five of the seven sectored colonies, this analysis was sufficient to detect W1, W2, R1, and R2. For the two sectored colonies in which we could not identify two genotypes in the sectors using markers on V, we used unselected events on other chromosomes to identify the granddaughters.

In addition to the seven selected crossovers, there were 71 unselected crossover or gene conversion events among the sectored colonies. Interpretations of eight classes of relatively simple recombination events (described in detail below) are shown in Figs. S1 and S2. In Figs. S3-S80, all of the selected and unselected events are shown, with the upper part of the figure depicting the pattern of heterozygous and homozygous SNPs on the two chromosomes of the granddaughter of the white (W1 and W2) and red (R1 and R2) sectors. The bottom part of each figure shows the inferred patterns of heterozygous and homozygous SNPs in the mother cells, prior to the segregation of chromosomes into the daughters. Each chromatid is depicted as double-stranded DNA and circles of different colors at the same positions on the two strands indicate unrepaired mismatches in heteroduplexes. In the discussion below, SNPs from the W303-1A-derived homolog will be called “red SNPs” and those from the YJM789-derived homolog will be called “blue SNPs”, consistent with the figures. Below, we summarize some of our findings based on this complex dataset.

#### SCB and DSCB events are approximately equally frequent

First, we can divide the events into two major classes, those in which the event was initiated by a single broken chromatid in S or G_2_ (SCB, single chromatid break) and those initiated by two broken chromatids (DSCB, double-sister-chromatid break), likely reflecting replication of a chromosome broken in G_1_. The heteroduplex patterns inferred from our analysis of granddaughter cells for two events are shown in Figure 7. In the event shown in Fig. 7A, only chromatid 2 has a heteroduplex, suggesting that the initiating event was a SCB. In contrast, in Fig. 7B, chromatids 1 and 3 have the CO configuration of markers, whereas chromatid 4 has a NCO configuration. This observation suggests that this event was initiated by a DSCB. Of 77 events that could be unambiguously classified (Table S3), there were 39 SCB and 38 DSCB events. In previous studies of events in wild-type strains treated with the same dose of UV (Yin and Petes, 2013), we also found approximately equal frequencies of these two classes.

**Figure 7.**
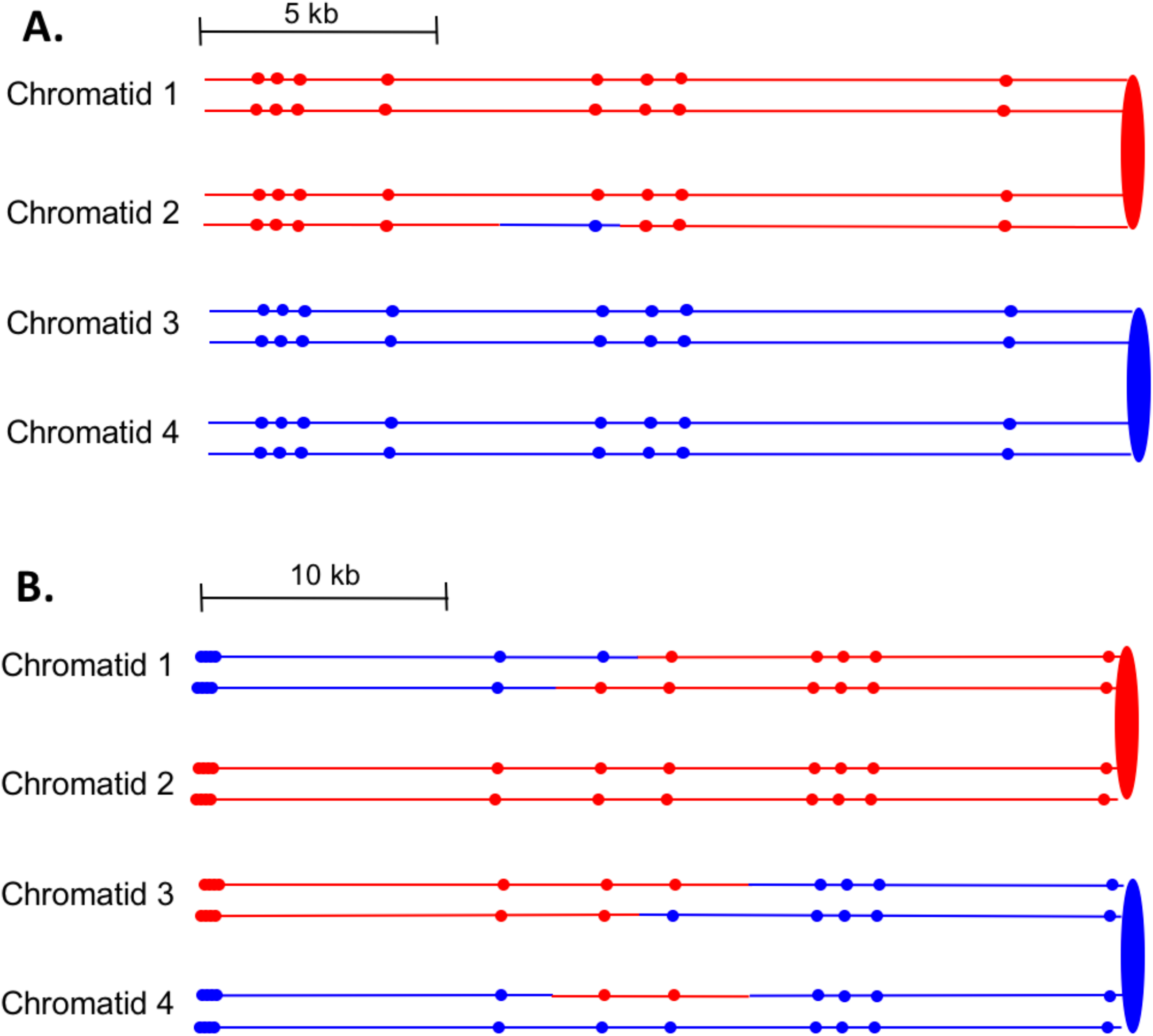
Examples of heteroduplex patterns associated with SCB- and DSCB-initiated events. Both the SCB event shown in 7A and the DSCB event shown in 7B were unselected events from the same sectored colony (YYy310-905WR; Table S3). The 7A event occurred on chromosome IV (labeled IV-2 in Table S3), and the 7B event was on chromosome XIV. The distances between SNPs (shown as red or blue circles) are to the scale shown above each set of chromatids. A. Inferred pattern of heteroduplexes in an SCB event. The patterns of markers in the granddaughter cells that define the patterns inferred in the G_2_ mother cell are in Fig. S6. A. Inferred pattern of heteroduplexes in an DSCB event. The patterns of markers in the granddaughter cells that define the patterns inferred in the G_2_ mother cell are in Fig. S16.

Excluding the selected CO events on chromosome V, of the 33 DSCB events, 22 had two NCOs, 9 had one NCO and one CO, and two had two COs. For our mechanistic interpretation of these events (to be described in detail below), we consider the two repair events resulting from the DSCB separately. The chromatids involved in a CO are almost always clear since the terminal SNPs at each end of the chromatid are altered in their coupling relative to the centromere as described above for chromatids 1 and 3 of Fig. 7B. For NCO events that have a single recombinant chromatid, we cannot determine which of the two non-recombinant chromatids was the donor of sequence information. For example, in Fig. 7A, the donor chromatid could be either chromatid 3 or 4 since these chromatids are identical. For such events, in Table S3, we arbitrarily assigned one chromatid as the donor.

It should be emphasized that in our experiments, as in prior studies (Lee *et al*., 2009; St. Charles and Petes, 2013; Yin and Petes, 2013), the DSCB events are far too frequent to represent two independent SCB events. The frequency of red/white sectored colonies (reflecting a crossover on chromosome V) in the UV-treated YYy310 strain was about 1.5%. About half of these events are a consequence of SCB. The likelihood of two independent SCB events resulting in an apparent DSCB is about (0.75%)^2^ or 5.6 × 10^−4^; we observed two orders of magnitude more DSCB events than this calculated frequency. In addition, the calculated frequency was based on double events occurring anywhere within the 120 kb between the *SUP4-o* insertion and *CEN5*. Most of the DSCB events involve DSBs that occur within a few kb of each other. Lastly, half of the time, two independent events would have donor chromatids derived from different homologs; only three of the DSCB events involved different homologs. Lastly, we point out that our measurement of the frequency of observed SCBs is an underestimate of the formation of SCBs, since many of the SCBs are likely to be repaired by sister-chromatid recombination (Kadyk and Hartwell, 1992).

Many previous studies of recombination in yeast show that the chromosome with the initiating DNA lesion acts a recipient of sequence information transferred from the intact donor chromosome (Symington *et al*., 2014). From our analysis, we can determine whether the homologs derived from the haploid parents YJM789 and W303-1A were equally susceptible to the initiating DNA lesion. As shown in Table S3, of the 38 DSCB events, 18 were initiated on the YJM789-derived homolog and 20 were initiated on the W303-1A-derived homolog. Of 40 SCBs (the event shown in Fig. S60 had two SCBs), the numbers initiated on the YJM789 and W303-1A homologs were 15 and 25, respectively; by chi-square analysis, these numbers are not significantly different from a 1:1 ratio (<u>p</u>=0.16). As expected, therefore, both homologs are equally susceptible to recombinogenic UV-induced DNA lesions.

#### Location of the DNA lesion that initiates recombination

For some events, the pattern of SNPs in recombinant chromatids allowed us to predict locations of the initiating DNA lesion, although most of these predictions had a degree of uncertainty. For single SCB events, the predicted position on the chromatid with the recombinogenic DNA lesion (likely a DSB) is shown as an arrow labeled DSB1 in Figs. S3-S80. For example, in Fig. S4, we show an arrow to the left of the conversion tract in chromatid 3. With equal validity, the arrow could have been placed to the right side or in the middle of the conversion tract. For DSCB events that are explicable by a single DSB on an unreplicated chromosome (Fig. S15), we show the DNA lesions by labeled arrows at the same positions on the replicated chromatids. In events that appear to require two independent DNA lesions (for example, Fig. S3), DSBs are labeled DSB1 and DSB2. For some DSCB events, we used information for all the recombinant chromatids to infer the position of the initiating DSB. For example, in Fig. S9, the initiating lesion for the NCO event on chromatid 3 could be placed at either end or in the middle of the conversion/heteroduplex tract. The pattern of the heteroduplex tract in chromatid 4, however, suggested the location of the initiating DSB shown by the arrow in Fig. S9.

#### Simple classes of CO and NCO chromatids

The 116 UV-induced CO and NCO events in YYy310 were classified into nine groups (Table S3), Classes 1-8 representing events with relatively straightforward interpretations (71 events) and a “Complex” class requiring more complicated mechanisms (45 events). In our discussion of mechanisms, we will assume that the recombinogenic lesion is a DSB, although there is evidence that DNA nicks are also recombinogenic (Fabre *et al*., 2002; Lettier *et al*., 2006; Davis and Maizels, 2014). As described above, the DSCB events are most easily explained as reflecting a DSB in G_1_; the SCB events could be initiated by either a DSB in S/G_2_ or a DNA molecule that is nicked in G_1_ and replicated to generate the DSB in S/G_2_.

In the discussion of classes below, we use the term “heteroduplex” to refer to duplex regions with mismatches, indicating DNA strands derived from different homologs. Conversion tracts are regions in which both of the interacting chromatids have homoduplexes derived from the donor chromatid (Fig. 2), and restoration tracts are regions in which the mismatches in heteroduplexes were inferred to be corrected to restore the original sequence of the recipient chromatid. The eight classes of events (Fig. 8, Classes 1-4 in Fig. S1 and Classes 5-8 in Fig. S2) are: Class 1 (NCO, single continuous heteroduplex tract on one side of putative DSB site; example: chromatid 2 of Fig. 7A), Class 2 (NCO, single continuous conversion tract on one side of putative DSB site; example: chromatid 3 of Fig. S4), Class 3 (NCO, hybrid heteroduplex/conversion tract located on one side of putative DSBs site; example: chromatid 4 of Fig. S5), Class 4 (CO, heteroduplex on only one side of putative DSB site; example: chromatids 1 and 3 of Fig. S28), Class 5 (NCO, heteroduplex on one side of putative DSB site, and conversion tract on the other; example: chromatid 3 of Fig. S14), Class 6 (CO, heteroduplexes in *trans* on recombinant chromatids; example: chromatids 1 and 3 of Fig. S16), Class 7 (CO, no observable heteroduplex or conversion tract on either recombinant chromatid; example: chromatids 1 and 3 of Fig. S24), Class 8 (NCO; two recombinant chromosomes, one containing a conversion tract, and the second with a heteroduplex involving SNPs at the same position; example: chromatids 2 and 3 of Fig. S67). The numbers of events in each of these classes (shown in parentheses) are: Class 1 (26), Class 2 (18), Class 3 (12), Class 4 (5), Class 5 (4), Class 6 (2), Class 7(2), and Class 8 (2).

**Figure 8.**
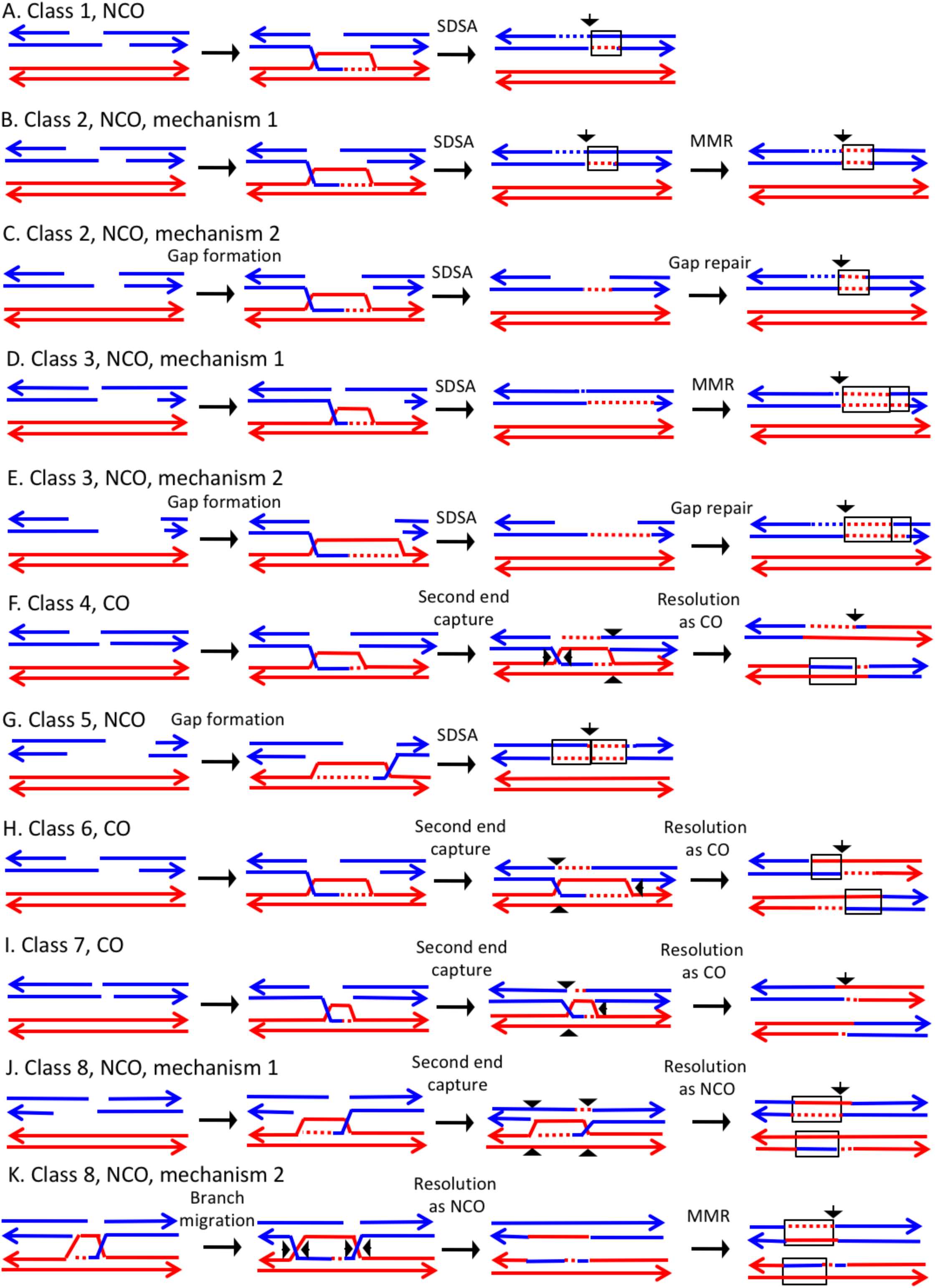
Mechanisms that generate Classes 1-8 recombination events. For all events, we show the initiating DSB on the blue chromatid. Dotted lines show sequences generated by replication or during mismatch repair. Black arrows indicate the position of the initiating DSB. The mechanisms are discussed in detail in the main text. A. Class 1. Class 1 events are NCOs formed by SDSA. B. Class 2. These NCO events could be generated by SDSA, followed by Mlh1p-independent MMR. C. Class 2. An alterative possibility is that Class 2 events reflect repair of a double-stranded DNA gap. D. Class 3. In this NCO class, the heteroduplex region is adjacent to a conversion tract. Such events could reflect a heteroduplex tract in which mismatches are repaired in one part of the tract and left unrepaired in the other. E. Class 3. An alternative model for this class is that the conversion tract is the result of repair of a double-stranded DNA gap with a heteroduplex region at one end. F. Class 4. For this CO class, a heteroduplex is observed on one chromatid but not the other. This class could be explained by the DSBR pathway in which heteroduplex region is short relative to the other; if the short heteroduplex does not contain a mismatch, it would be undetectable. G. Class 5. In this NCO class, the heteroduplex region is located on the opposite of the DSB site from the conversion region. This pattern is consistent with the repair of a double-stranded DNA gap that was restricted to one of the broken ends. H. Class 6. This CO class is identical to the pattern expected for the DSBR model. I. Class 7. In this CO class, no heteroduplexes or conversion tracts are observed adjacent to the crossover, consistent with the formation of a dHJ with short heteroduplex tracts that do not include mismatches. J. Class 8. In this NCO class, one chromatid has a conversion tract, and the other chromatid has a heteroduplex involving SNPs at the same position. This event could reflect resolution of a dHJ intermediate in the NCO model in which one region of heteroduplex is undetectable. K. Class 8. An alternative mechanism involves branch migration of a HJ, followed by resolution of the intermediate in a NCO mode. Mismatches in one of the two chromatids are repaired to generate a conversion tract.

Before discussing these classes in detail, we point out that heteroduplexes can be detected only if the heteroduplex includes a SNP. The diploid used in our study is heterozygous for about 55,000 SNPs, but only 13,000 of these SNPs are included on the microarrays (St. Charles *et al*., 2012). Since the average distance between SNPs, as assayed by the microarray, is about 1 kb, a fraction of short heteroduplex/conversion tracts will be undetectable (St. Charles *et al*., 2012; Zheng *et al*, 2016).

The largest class, Class 1, has the pattern of SNPs expected for events initiated by the SDSA pathway (Fig. 1A, Fig. 8A). Class 2 events could also reflect SDSA events, however, the observed conversion tracts are not consistent with the simplest form of the DSBR model. One possibility is that, following disengagement of the invading end, the mismatches in the heteroduplex are repaired by an Mlh1-independent pathway (Fig. 8B). Coïc *et al*. (2000) reported the existence of a short-patch (<12 bp) Msh2-independent repair pathway in yeast. Alternatively, short-batch mismatches close to the DSB could also depend on the proofreading activity of polymerase d (Anand *et al*., 2017). Although such pathways could explain conversion tracts limited to one or a few closely-spaced SNPs (Fig. S4, for example), it seems unlikely to explain conversion tracts extending over several kb (Fig. S23: for example).

An alternative model is that the Class 2 NCO events are a consequence of the repair of a double-stranded DNA gap (Fig. 8C). In experiments in which gapped plasmids are transformed into yeast, gap-repair occurs readily (Orr-Weaver *et al*.,1981), and such gap repair was an intrinsic part of the original DSBR model (Szostak *et al*., 1983). Since in most meiotic recombination events in yeast, DSBs are processed by degradation of only one strand of the duplex (De Massy *et al*., 1995), gap repair is usually not considered part of the standard DSBR model. Giannattasio *et al.* (2010), however, reported that the Exo1p could cause expansions of nucleotide excision repair tracts from 30 bases to several kb. Based on our previous observations that showed a two-fold reduction in UV-induced mitotic recombination in the *exo1* mutants, we previously suggested that Exo1-mediated tract expansion has a role in generating recombinogenic lesions in UV-treated cells (Yin and Petes 2014); the Exo1p could be involved in the long-tract Class 2 events. In summary, we suggest that events with long conversion tracts may reflect the repair of a double-stranded DNA gap, and short regions of conversion may represent Mlh1-independent MMR. Another possible mechanism for Class 2 events is that the broken end invades the homolog, and copies sequences by a BIR-like mechanism before disengaging from the homolog and re-engaging with the other broken end.

Class 3 NCO events are similar to Class 2, and have a similar mechanistic explanation. For Class 3 events, however, mismatches in one segment of the heteroduplex are repaired and those in the adjacent segment are not (Fig. 8D). Alternatively, the conversion tract could be generated by repair of a double-stranded DNA gap with a region of heteroduplex adjacent to the repair tract (Fig. 8E). The Class 4 CO events resemble the classic DSBR pathway except heteroduplexes are observed on only one site of the putative DSB site. One possibility is that the event occurs by the DSBR pathway but one heteroduplex does not include a heterozygous SNP and, therefore, is not observable (Fig. 8F). Studies of meiotic recombination suggest that heteroduplexes flanking the recombination-initiating DNA lesion are often of different lengths (Merker *et al*., 2003; Jessop *et al*., 2005).

Class 5 NCO events are explicable as reflecting the repair of one broken end with a double-stranded DNA gap and one broken end with the canonical processing of one strand (Fig. 8G) Following SDSA, a recombinant chromatid in which the putative DSB site is flanked by a heteroduplex on one side and a conversion tract on the other will be produced. The Class 6 CO events have the structure predicted for a CO in the DSBR pathway (Fig. 1, Fig. 8H). There were only two such events in our dataset. Class 7 events resemble Class 4 and Class 6 events except there are no observed heteroduplexes on either side of the putative DSB site. Class 7 COs might reflect intermediates with short heteroduplexes or heteroduplexes that are in regions that do not include SNPs (Fig. 8I).

In the Class 8 NCO events, both interacting chromatids have recombinant SNPs, and the conversion tract on one chromatid overlaps with the heteroduplex tract on the other (Fig. 8J and 8K). In Fig. 8J, this class is generated by formation of a dHJ with one region of short/undetectable heteroduplex, followed by resolution of the dHJ as a NCO. An alternative possibility is that branch migration occurs to generate regions of heteroduplex on both chromatids (Fig. 8K). Mlh1-independent repair of mismatches in one, but not both, of the heteroduplexes could produce the observed pattern.

If we assume that not all heteroduplexes are detectable, Classes 1, 4, 6, and 7 are consistent with the DSBR model of Fig. 1. For CO classes with *trans* heteroduplexes, all events had the pattern expected for nick-directed resolution of Holliday junctions. As shown in Fig. 1, nick-directed cleavage of the junctions (sites marked as 1, 2, 7 and 8) result in a different pattern of heteroduplexes than cleavage of unnicked junctions (sites marked as 2, 4, 5, and 6). Previously, a bias of the same type was detected for recombination events involving plasmid integration (Mitchel *et al.*, 2010). This observation is also consistent with previous studies that showed that *mus81* strains, but not *yen1* strains, had reduced frequencies of crossovers without an effect on gene conversions (Ho *et al.*, 2010; Yin and Petes, 2015); Mus81, but not Yen1, has a substrate preference for nicked junctions (Schwartz and Heyer, 2011).

Most of the other events require Mlh1-independent mismatch correction or the repair of a double-stranded DNA gap. These classes, however, represent only half of the total “simple” events. As discussed below, the complex classes require even more radical departures from the canonical recombination model.

#### Complex classes of CO and NCO chromatids

Those events that were classified as complex are shown in Table S3. A more detailed description of the complex classes is in Table S4, and possible interpretations of these events are shown schematically in Figs. S92-S124. Of the SCB events, 35 of 40 were “simple” (Classes 1-8) and only 5 were complex. Assuming the same distribution for the 38 DSCB events, the expected frequencies of DSCBs with two simple events, one simple and one complex event, and two complex events are 29, 8, and 1, respectively. We observed (Table S2) 12, 11, and 15 of these classes, respectively, a very significant (p<0.0001 by chi-square test) departure from this expectation. This observation suggests that the timing of the recombinogenic DSB (G_1_ versus G_2_) may influence the complexity of its repair. Below, we will discuss two of these complex events in detail. We note that many of these events can be explained by more than one mechanistic pathway, and we have tried to describe the simplest one.

The event depicted in Fig. S8 (re-drawn in Fig. 9A) is a DSCB event initiated on the red chromosome. The CO occurred between chromatids 1 and 4. One feature of the CO that is not consistent with the canonical DSBR model is that heteroduplexes are observed at the same position on both crossover chromatids. This pattern is consistent with the possibility of branch migration of one of the Holiday junctions in the rightward direction, resulting in a region of symmetric heteroduplexes (Fig. 9B). Since the heteroduplex tract is longer on chromatid 4 than on chromatid 1, there was likely a region of Mlh1-independent conversion-type repair of mismatches on chromatid 4. In the NCO event (involving chromatids 2 and 3), chromatid 2 had two regions of heteroduplex in which there was a strand switch at the junction of the putative DSB site. This pattern can be explained by formation of a dHJ that was resolved by dissolution (Fig. 9C). As for the CO event, we also need to postulate a region of Mlh1-independent MMR at one end of the heteroduplex tract.

**Figure 9.**
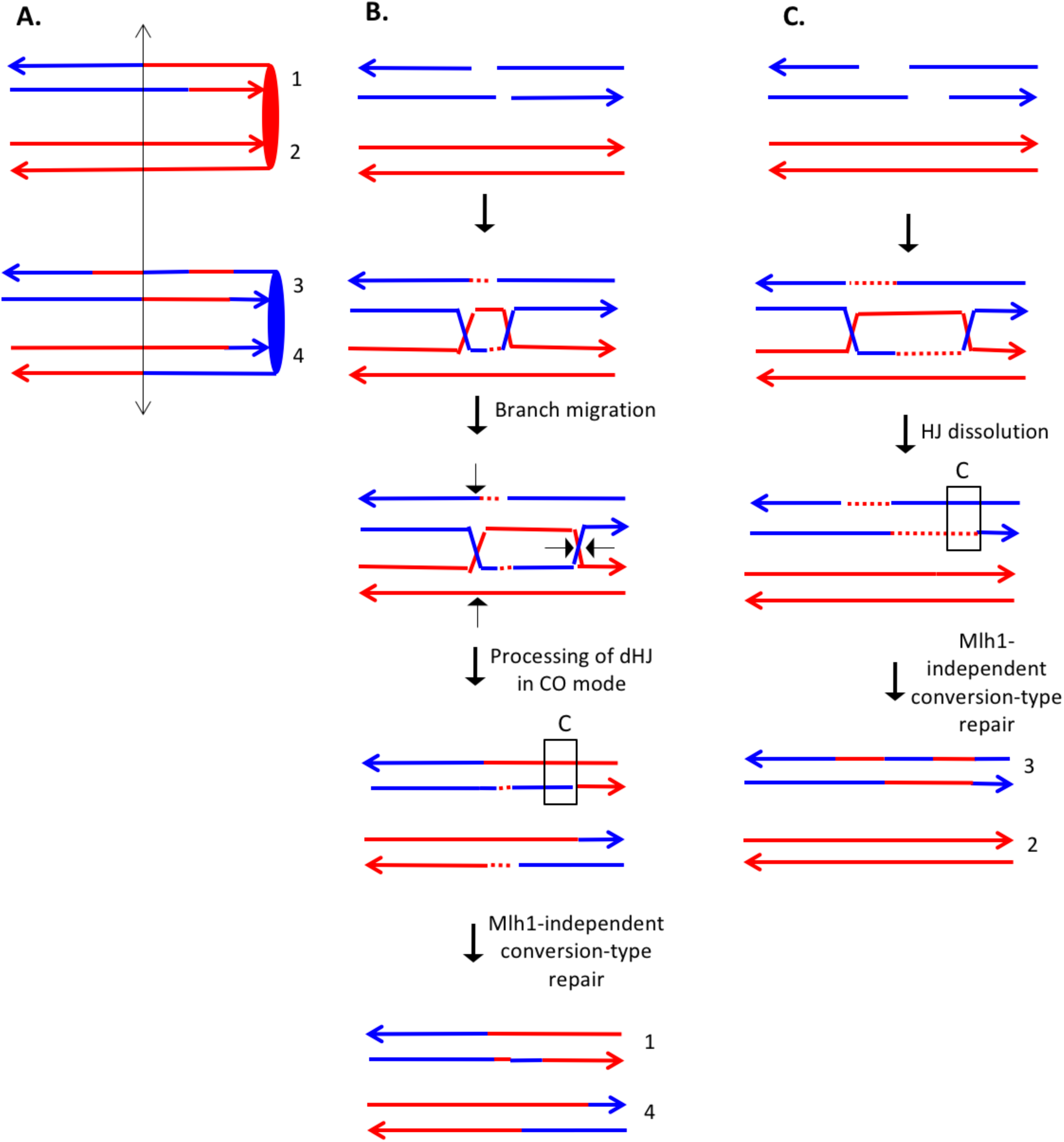
Complex DSCB event derived from analysis shown in Fig. S8. We infer that the recombination-initiating DSB was on a blue chromosome that was replicated to produce two broken blue chromatids. A. Depiction of heteroduplexes and conversion tracts present in the mother cell before chromosome segregation. The double-headed arrow indicates the position of the initiating DSB. B. Mechanism to produce the CO chromatids 1 and 4. Following invasion of the left end of the broken chromatid, one HJ underwent branch migration resulting in a region of symmetric heteroduplexes. The dHJ intermediate was processed to yield a CO as shown by the short arrows. Mismatches in a region of the heteroduplex on the upper chromatid were repaired to yield a conversion event. We suggest that the small homoduplex region near the DSB site did not include a SNP and was, therefore, undetectable. C. Mechanism to produce the NCO chromatids 2 and 3. Following dHJ formation, the intermediate was dissolved, resulting in bidirectional heteroduplexes in *cis* on chromatid 2. In addition, as in Fig. 8B, we hypothesize that there was a region of mismatch repair in one of the heteroduplexes.

The event shown in Fig. S16 (re-drawn in Fig. 10A) is also a DSCB event with the CO involving chromatids 1 and 3, and the NCO involving chromatid 4. The CO event is consistent with the classic DSBR model (Fig. 10B). If we assume that the NCO event was initiated by a DSB at the same position, the heteroduplex spanning the DSB site is inconsistent with the DSBR model. One mechanism to explain the observed NCO pattern is that the right broken end undergoes degradation of both strands, and the left end is extended by invasion of the sister chromatid (Fig. 10C). This invading end is then unpaired from the sister chromatid and pairs with the right end. After processing the dHJ in the NCO mode, the net result of these steps is a recombinant chromatid with a region of heteroduplex that spans the initiating DSB site.

**Figure 10.**
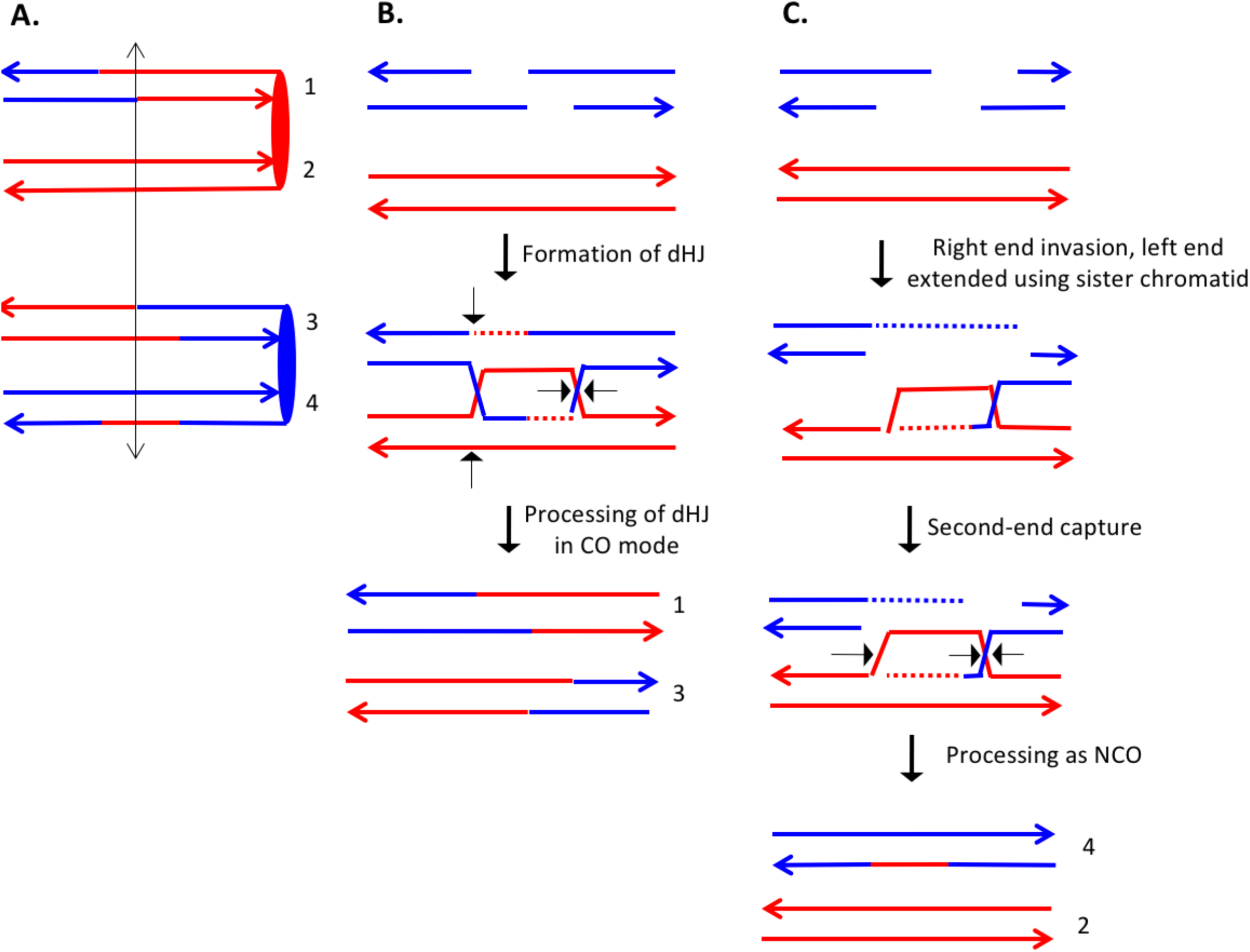
Complex DSCB event derived from the analysis shown in Fig. S16. As in Fig. 8, the initiating DSB was on the blue chromosome. A. Depiction of recombinant chromatids in mother cell before segregation of chromosomes into the daughters. B. CO between chromatids 1 and 3. The pattern of heteroduplexes on these chromatids is that expected from the DSBR model. C. NCO between chromatids 2 and 4. In this event, the heteroduplex spans the putative DSB site on chromatid 4. The right broken end underwent resection of both strands before strand invasion. The left end was extended by invasion of the sister chromatid. Following annealing of the extended end to the D-loop, the intermediate was processed as a NCO. We suggest that the small region of heteroduplex on the red chromatid did not contain a SNP and was not detectable.

Of the 46 complex events described in Tables S3 and S4, about 40% (18 of 46) can be explained by mechanisms involving template switching, branch migration, or independent invasion of two broken ends. In addition, this relatively large fraction of complex events is not solely a characteristic of UV-induced events in *mlh1* strains. About one-third of the spontaneous events in a wild-type strain (St. Charles and Petes, 2013), and 15% of the UV-induced events in a wild-type strain (Yin and Petes, 2013) appear associated with “patchy” repair of mismatches and/or branch migration.

#### Lengths of conversion/heteroduplex tracts in UV-treated YYy310

The median lengths of conversion/heteroduplex tracts in NCO and CO events in UV-treated YYy310 were 5.4 kb (95% confidence limits [CL] of 3.6-7.2 kb) and 10 kb (CL 5.7-17 kb), respectively; the median length of all tracts was 6.1 kb (CL 4.8-8.5 kb). For the CO events, we summed the lengths of the tracts on the two CO chromatids, since these events reflect the repair of a single DSB. These lengths are similar to those previously determined for the UV-treated isogenic wild-type strain (Yin and Petes, 2013): 5.7 kb (CL 4.5-6.6) for NCO events; 8.2 kb (CL 6.6-10.3) for CO events; 6.4 kb (CL 5.8-7.3) for all tracts. Thus, the Mlh1p does not have a significant role in determining the length of conversion tracts.

### Analysis of spontaneous crossovers on chromosome V in YYy310

Although most of the conclusions from our study are based on UV-induced events in YYy310, we also examined five spontaneous crossovers on chromosome V by similar procedures. No unselected crossover or BIR events were observed among these sectors. The coordinates for LOH transitions in the granddaughter cells of each colony, and the classification of these events as Classes 1-8 or complex events are shown in Tables S2 and S3, respectively. The events are depicted in Figs. S81-S85, and the mechanisms that explain these events are in Figs. S125-S129.

All five sectored colonies reflected DSCBs with one CO and one NCO event. Nine of the ten COs and NCOs were complex with similar features to the UV-induced complex events (Mlh1-independent repair, gap repair, branch migration, and independent invasion of two broken ends) (Table S4). Thus, the complexity of the patterns of recombination observed in the UV-induced events is not likely to be a consequence of the multiple DNA lesions introduced by UV.

### Analysis of spontaneous crossovers at the HS4 hotspot on chromosome IV in YYy311

Previously, we showed that an inverted pair of Ty elements resulted in a spontaneous mitotic recombination hotspot (termed “HS4”) on the right arm of chromosome IV (St. Charles and Petes, 2013). We subsequently showed that HS4-associated conversion events were associated with very long (>25 kb) conversion tracts (Yim *et al*., 2014); conversion tracts associated with a similar pair of inverted Ty elements on chromosome III also have very long conversion tracts (median length of 41 kb; Chumki *et al.*, 2016). To monitor the activity of this hotspot in an *mlh1* strain, we inserted the *hphMX4* and *URA3* markers centromere-proximal and centromere-distal to HS4 on one of the two homologs in the *mlh1* diploid YYy311. A crossover between the two markers results in a red/white sectored colony in which the white sector is resistant to hygromycin (Hyg^R^) and 5-fluoro-orotate (5-FOA^R^), and the red sector is Hyg^R^ 5-FOA^S^ (Fig. 11). In a wild-type strain with the *SUP4-o* marker located near the right end of chromosome IV, 17% of sectored red/white colonies result from a crossover in the *hphMX4-URA3* interval (St. Charles and Petes, 2013). 17% (101/593) of the spontaneous sectoring events in the isogenic *mlh1* strain YYy311 occur in this same interval, indicating that loss of Mlh1 does not reduce the activity of HS4. We examined six of the sectored colonies by SNP microarrays (Tables S2-S4), and five had patterns of recombination indicating an HS4-initiated event (shown in Figs. S87-S91). Since the recombination event in sectored colony shown in Fig. S86 is initiated somewhere in the interval 968-978 kb, and HS4 is located in the interval 981-993 kb, this event is not likely to be HS4-initiated (Fig. S130). Since the 56 kb *hphMX4-URA3* interval is considerably larger than the size of the HS4 hotspot (12 kb), it is not surprising that some crossovers in the *hphMX4-URA3* region are not HS4-mediated. Mechanisms consistent with the HS4-mediated events are shown in Figs. S131-S135.

**Figure 11.**
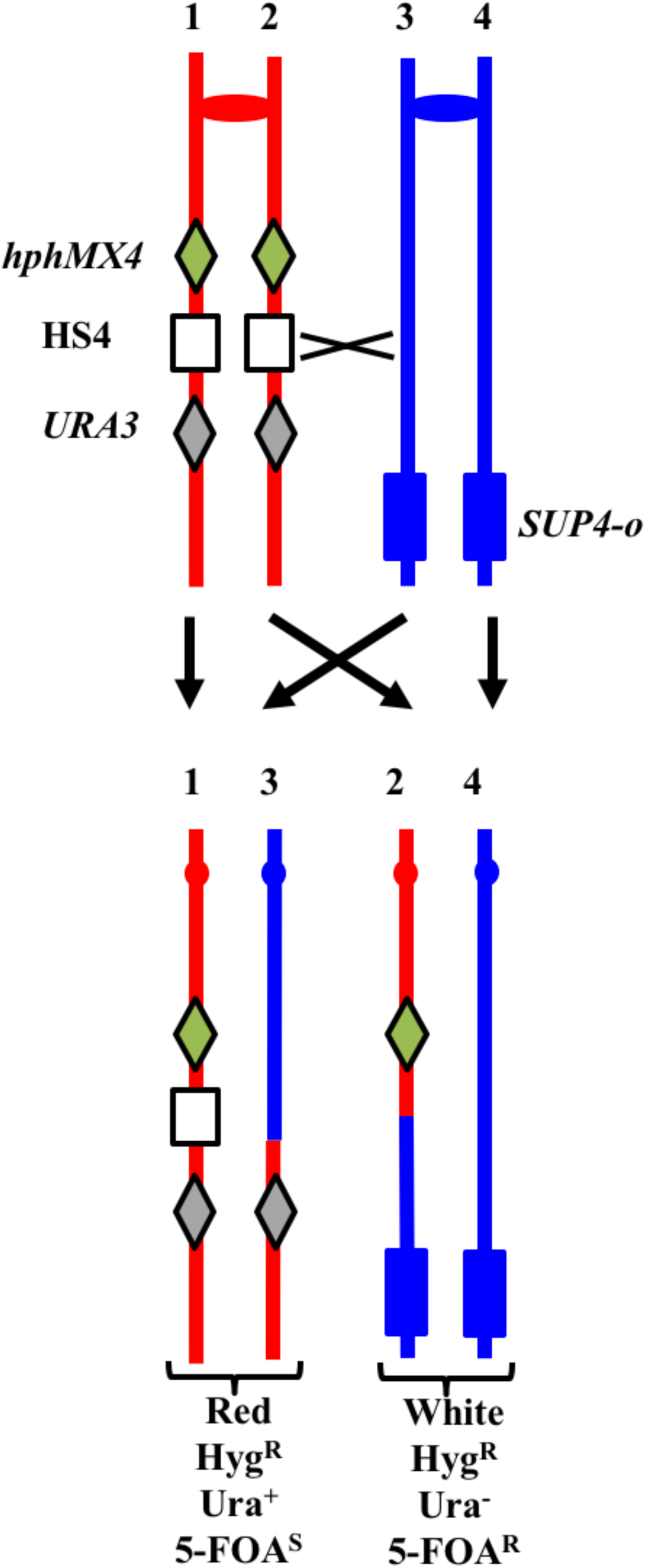
System for detecting crossovers at the HS4 hotspot on chromosome IV (strain YYy311). The W303-1A- and YJM789-derived chromatids are shown in red and blue, respectively. On the W303-1A-derived homolog, the HS4 hotspot is flanked by the *hphMX4* and *URA3* markers; *SUP4-o* is located near the telomere of the YJM789-derived homolog. Since the recombinogenic DSB is located at HS4, HS4-initiated events usually result in loss of HS4 as shown.

Based on the five HS4-initiated events, we can make several generalizations. In all events, at least three chromatids had transitions between W303-1A- and YJM789-derived sequences, indicating that all HS4-initiated events involved the repair of two sister chromatids broken at approximately the same position. Second, all events are initiated on the red (W303-1A-derived homolog), as expected since the YJM789-derived homolog lacks HS4 hotspot activity (St.Charles and Petes, 2013). Third, with the exception of the event in Fig. S88, gaps of at least 25 kb are observed in the HS4-initiated events; these gaps are much larger than necessary to remove the approximately 9 kb HS4-related heterology that is present in W303-1A but absent in YJM789-derived homolog (St. Charles and Petes, 2013). Fourth, many of these gaps are flanked by heteroduplexes on one or both sides. Gaps can be propagated symmetrically from HS4 (Fig. S89) or with a bias toward or away from the centromere. Fifth, some of the gaps (regions of 4:0 segregation without heteroduplexes) are interrupted by regions of heteroduplex (chromatid 4 of Fig. S91). Such a pattern can be explained by a template switch to a sister chromatid as shown in Fig. S135.

We also used SNP microarrays to examine twelve red/white sectored colonies with a crossover in the *hphMX4-URA3* interval in UV-treated YYy311. Since it was clear that none of these twelve colonies had an HS4-mediated event, we did not attempt to look for two genotypes within each sector, In addition, a smaller fraction of events (10%, 10/102) were in the *hphMX4-URA3* interval in the UV-treated samples of YYy311 than in the untreated samples. Thus, HS4 is a hotspot for spontaneous, but not UV-stimulated recombination events.

### Effect of the *mlh1* mutation on the frequency of mitotic crossovers

Two different assays can be used to monitor the effect of the *mlh1* mutation on crossovers: the frequencies of spontaneous and UV-induced red/white sectored colonies compared to wild-type, and the frequency of unselected crossovers in UV-treated cells. These two assays yield somewhat different results. We observed slightly more (<two-fold) spontaneous red/white sectored colonies in YYy311 than in the isogenic wild-type strain (53 sectored colonies/998395 total colonies [5.3 × 10^−5^/division]) in the *mlh1* strain versus 55 sectors/1761664 total [3.1 × 10^−5^/division; St. Charles and Petes, 2013]). A similar slight elevation was observed for UV-treated YYy311 cells: 378 sectors/3228 total (11.7%) in the *mlh1* strain versus 127 sectors/1420 total (8.9%) in the isogenic wild-type strain. In addition, the frequency of UV-stimulated sectored colonies in YYy310 (*SUP4-o* on chromosome V) is higher than the UV-stimulated events in the isogenic wild-type strain: 45 sectors/2980 colonies (1.5%) and 68 sectors/7194 colonies (0.9%), respectively.

An elevated frequency of crossovers could be explained by the heteroduplex-rejecting properties of Mlh1p (Harfe and Jinks-Robertson, 2000). If this mechanism was relevant to our system, we might expect a higher density of SNPs and in/dels in the heteroduplex tracts of *mlh1* strains than observed in tracts in the wild-type strain. In the UV-treated *mlh1* strain, we observed 2688 SNP+in/dels in tracts with a total length of 635041 bp, a density of 0.0042. In the UV-treated wild-type strain (Yin and Petes, 2013), we found 15387 SNP+in/dels in tracts totaling 4573874 bp, a density of 0.0034. By a Mann-Whitney comparison, the difference in density is of borderline significance (<u>p</u>=0.06).

In contrast to the first assay, the second assay indicates that Mlh1p positively influences the frequency of crossovers. In our previous study of unselected recombination events, in twenty UV-induced sectored colonies in the wild-type strain (Yin and Petes, 2013), we observed 141 interstitial LOH events, 50 COs, and 10 BIR events for a total of 201 LOH events (10.1/sectored colony). In seven sectored colonies of the *mlh1* YYy310 strain, we found 65 interstitial LOH events, 6 COs, and 0 BIR events for a total of 71 LOH events (10.1/sectored colony). The reduction in the number of COs is statistically significant (p=0.003 by Fisher exact test).

Although the discrepancy in the two assays prevents a strong conclusion about the effect of Mlh1p on mitotic crossovers, the direct assay of unselected crossovers by microarrays is less prone to artifacts than the frequency of red/white sectors since colony color can be affected by a variety of different types of genetic alterations including mitochondrial mutations (Kim *et al*., 2002).

## Discussion

The main conclusion from our analysis is that about one-third of mitotic recombination events are not explicable by the simplest forms of the double-strand-break repair model. More specifically, we need to invoke branch migration of Holliday junctions, template switching during repair synthesis, repair of double-stranded DNA gaps, and/or Mlh1-independent MMR. Our analysis confirms our previous conclusion that more than half of spontaneous crossovers and recombination events induced by high doses of UV reflect the repair of two sister chromatids broken at approximately the same position, likely resulting from replication of chromosome broken in G_1_. We also show that mitotic recombination events involve heteroduplexes that are longer than those observed for meiotic exchanges. In a previous study, we characterized a hotspot for spontaneous mitotic recombination events (HS4) associated with an inverted pair of Ty elements (St. Charles and Petes, 2013). In the current study, we show that the very long conversion tracts associated with HS4-initiated events (Yim *et al*., 2014) are associated with the repair of long double-stranded DNA gaps. In addition, HS4 hotspot activity is independent of Mlh1p and is not induced by UV.

### Complexity of recombination events

Before discussing exceptions to the recombination model shown in Fig. 1, it is important to point out that about two-thirds of the observed events <u>do</u> fit this model. Among the exceptional events are those in which the two interacting chromatids have heteroduplexes involving SNPs at the same positions (for example, the CO chromatids 1 and 4 in Fig. S8). Symmetric heteroduplexes of this type can be explained as a consequence of branch migration of a Holliday junction (Fig. S94). We previously suggested that branch migration may explain complex patterns of SNPs in MMR-proficient strains (St. Charles *et al*., 2012; Yin and Petes, 2013). Branch migration during meiotic recombination in yeast would be detectable as two spores with post-meiotic segregation within one tetrad (aberrant 4:4). Such events are rare in yeast, although a few percent of the total aberrant segregants had this pattern in an *msh2* strain (Martini *et al*., 2011).

In the current DSBR model, gene conversion events are a consequence of the repair of mismatches in heteroduplexes. Thus, in the absence of MMR, conversion events should be absent. In our study, we frequently observed conversion tracts, some adjacent to a heteroduplex tract (Fig. S5, chromatid 4) and some “solo” conversions (Fig. S4, chromatid 3) One possible source for such events is Mlh1-independent MMR. In cell-free extracts, MMR is only partially defective in the absence of Mlh1 if there is a 5’ nick near the mismatch (Genschel and Modrich, 2009). In addition, Coïc *et al*. (2000) identified a short-patch repair system in *S. cerevisiae* that was independent of the classic MMR system. The exonuclease activity of DNA polymerase d could also remove mismatches close to the DSB (Anand *et al.,* 2017). If these uncharacterized systems are responsible for the conversion events observed in our studies, they are most likely to be involved in those events with short conversion tracts. We suggest that long conversion tracts (chromatid 2, Fig. S33; Figs. S87-91) likely reflect the repair of a large double-stranded DNA gap. Although DSBs are usually thought to be processed by 5’-3’ degradation of the broken ends, resection of the 3’ “tails” has also been observed (Zierhut *et al*., 2008). It is clear that *S. cerevisiae* has the ability to repair double-stranded DNA gaps (Orr-Weaver *et al*., 1981) and, in the first version of the DSBR model, both gap repair and MMR were presented as mechanisms for generating a gene conversion event (Szostak *et al*., 1983).

We also observed both short and long tracts of restoration. For example, the long heteroduplex tract in chromatid 2 of Fig. S3 is interrupted by several one-SNP restoration tracts; such tracts may reflect Mlh1-independent MMR. In Fig. S44, a long heteroduplex tract on chromatid 1 is interrupted by a long tract of restoration. As shown in Fig. S105, this pattern can be explained by a template switch, followed by SDSA. In invoking template switching to explain certain patterns of SNP segregation, we point out that such switches were also proposed as intermediates in the production of complex meiotic recombination events (Martini *et al*., 2011). In addition, template switching is commonly observed during break-induced replication (Smith *et al*., 2007). Since the sister chromatid is a preferred substrate for the repair of DSBs in mitosis (Kadyk and Hartwell, 1992; Bzymek *et al*., 2010), a template switch to the sister chromatid would not be unexpected.

In previous mitotic (Miura *et al*., 2012) and meiotic recombination studies (Martini *et al*., 2011), some events appear to reflect independent interactions of the left and right broken ends with the intact template. For example, Martini *et al*. proposed that a double SDSA event could produce a NCO chromatid in which *trans* heteroduplexes were separated by a region of restoration. Similarly, in our analysis, some events are likely to reflect double-ended invasions. In Fig. S83, the CO chromatid 4 has a *trans* heteroduplex flanking the putative DSB site. This pattern, which is unexpected by the DSBR model, can be explained by invasion of the right broken end, followed by processing, and subsequent invasion of the left broken end (Fig. S126B).

The complex patterns of recombination observed in our study are also found in global analysis of meiotic recombination events in wild-type (Mancera *et al*., 2008) and *msh2* (Martini *et al*., 2011) strains. In the Mancera *et al*. study, more than 10% of the conversion tracts were described as “complex”, and, in the Martini *et al*. analysis, the majority of the CO-associated heteroduplex tracts required Msh2-independent MMR, branch migration, template switches, and/or the repair of double-stranded gaps. These observations contrast with other studies of mitotic recombination events induced with an endonuclease at a specific site; such events tend to have simpler patterns of marker segregation (Nickoloff *et al*., 1999; Mitchel *et al*., 2010; Hum and Jinks-Robertson, accompanying paper). Our studies differ from most previous studies in several ways: the nature of the recombinogenic lesions, the timing of the DSBs during the cell cycle, and the diverse genetic location of the recombinogenic lesions. With regard to the third point, if DNA lesions at different regions of the genome utilize different pathways of repair, as suggested by meiotic recombination studies (Medhi *et al*., 2016), then a global analysis of recombination would reveal more heterogeneity of recombinant products than studies based on a single locus. Lastly, we point out our analysis is performed in diploid cells whereas many of studies of HO-induced events are done in haploids with duplicated sequences. Any or all of these factors may result in the greater complexity observed in our recombination events.

### Comparison of recombination events in wild-type and *mlh1* strains

In general, our observations support the main conclusions of our previous studies of spontaneous (St. Charles and Petes, 2013) and UV-induced (Yin and Petes, 2013) mitotic recombination. In wild-type and *mlh1* strains, exchanges induced by high levels of UV in G_1_- synchronized cells represent DSCBs and SCBs with approximately the same frequency. All five of the spontaneous events selected on chromosome IV in the *mlh1* strain YYy311 were DSCB events as were the majority of spontaneous events in the wild-type strain (St. Charles and Petes, 2013). Events induced by I-SceI were almost exclusively the result of a DSCB (Hum and Jinks-Robertson, accompanying paper). ln contrast, recombination events induced by low levels of DNA polymerase alpha (Song *et al*., 2014) or delta (Zheng *et al*., 2016) usually reflect the break of a single chromatid.

Esposito (1978) previously proposed that mitotic recombination in yeast was initiated in G_1_-associated DSB that formed a Holliday junction connecting the two unreplicated homologs. Mismatch correction occurred in the heteroduplex regions, and the resulting intermediate was replicated to produce both CO and NCO recombinant products. By this model, granddaughter cells would not be expected to be genotypically different. Our analysis rules out this model.

In addition to its role in the repair of mismatches, the Mlh1p has two other functions related to recombination. First, in conjunction of Mlh3p, Mlh1p stimulates meiotic crossovers (Hunter and Borts, 1997; Wang *et al*., 1999); this function is likely related to its single-strand nicking activity (Rogacheva *et al*., 2014). If Mlh1p had a similar role in mitotic recombination, we would expect a hypo-Rec phenotype in *mlh1* strains. Conversely, for recombination substrates with diverged sequences, Mlh1 has anti-recombination activity (Nicholson *et al*., 2000). Strains lacking *mlh1* have seven-fold more crossovers than wild-type strains for substrates with 9% sequence divergence. Although this degree of divergence is much greater than the average 0.5% divergence existing between the homologs in YYy310 and YYy311, strains lacking the Msh2 and Msh3 repair proteins have an elevated frequency of crossovers relative to wild-type for 350 bp substrates with a single mismatched base (Datta *et al*., 1997).

Based on the data described above, it is difficult to predict the effect of the *mlh1* mutation on mitotic recombination in our system. As discussed in the Results section, we find an elevated frequency of sectored colonies in the *mlh1* strain compared to the wild-type strain, but a reduced frequency of crossovers among unselected UV-induced recombination events. Although the results indicating a several-fold reduced frequency of crossovers in the *mlh1* strain are likely to be less prone to artifacts than the sectored colony assay, further experiments are needed to substantiate them.

### Analysis of the HS4 mitotic recombination hotspot

Previously, we demonstrated that an inverted pair of Ty elements on the W303-1A-derived copy of chromosome IV stimulated mitotic recombination (St. Charles and Petes, 2013); this hotspot was called “HS4”. The HS4-associated recombination events have the following properties (St. Charles and Petes, 2013; Yin and Petes, 2014; Yim *et al*., 2014): 1) they are DSCB events, 2) both of the Ty elements are required for hotspot activity, 3) the events are initiated on the W303-1A homolog presumably because the YJM789 homolog lacks the inverted pair of elements, 4) HS4 activity requires the Exo1p, and 5) conversion events associated with HS4 are extraordinarily long with a median size >50 kb. Our analysis of five HS4-related events (Figs. S87-91; S131-135) shows that most have the properties expected for the repair of a large double-stranded gap (long conversion events) with flanking heteroduplex regions. Since the HS4 hotspot on the W303-1A-derived homolog is a pair of inverted repeats and the comparable region on the YJM789 homolog has only a partial Ty element in the same location (St. Charles and Petes, 2013), a DSB occurring within HS4 would require a region of end resection of at least 10 kb to expose sequence homologies between the two homologs. In general, the conversion/heteroduplex tracts for the HS4-induced events exceed 10 kb by a considerable amount. For example, in the NCO event on chromatid 2 of Fig. S91, the total length of the conversion/heteroduplex region is 39 kb, comparable to the lengths of conversion tracts observed in the wild-type strain (Yim *et al*., 2014).

Our data show that the very long conversion tracts observed in wild-type strains are not a consequence of extremely long heteroduplexes, but reflect an intermediate that does not have mismatched bases. Since we observe many other events that are explicable as gap repair, this is our preferred explanation for the HS4-associated events, which is consistent with the observation that the activity of HS4 requires Exo1p (Yin and Petes, 2014). The possible role of Exo1p is the expansion of a single-stranded nick into a large single-stranded gap before DNA replication, facilitating the formation of secondary structures involving the inverted Ty elements that could be subsequently processed by structure-specific nucleases. We cannot, however, rule out the generation of these long conversion tracts by a BIR-like mechanism (Yim *et al*., 2014). We found that HS4-associated hotspot was not stimulated by UV. This result argues that the activity of HS4 is not a consequence of random DNA breaks located near in the inverted repeat, but is likely a consequence of a DSB induced in a secondary structure in the DNA that is formed independently of nearby DSBs. Lastly, we suggest that the long conversion tracts associated with HS4 might be a general property of recombination events initiated in a large heterozygous insertion.

## Summary

In conclusion, our global analysis of UV-induced and spontaneous mitotic recombination events in an MMR-deficient yeast strain demonstrates that many of the events are more complex than predicted by the current DSBR model. We find that many gene conversion tracts are generated by a mechanism that does not involve the Mlh1-dependent repair of mismatches within a heteroduplex.

## Materials and Methods

### Strains

Constructions of the hybrid diploid strains YYy310 (isogenic isolates YYy310.9, YYy310.10) and YYy311 (isogenic isolates YYy311.1 and YYy311.3) are described in detail in Table S1. In brief, the *MLH1* gene was deleted from haploid derivatives of W303-1A and YJM789; these strains have about 0.5% sequence divergence (Wei *et al*., 2007). The primary difference between YYy310 and YYy311 is the location of the tRNA suppressor gene *SUP4-o*: located near the left telomere of the YJM789-derived chromosome V homolog in YYy310, and near the right telomere of the YJM789-derived chromosome IV homolog in YYy311. As explained in the text, this heterozygous insertion allows detection of crossovers as red/white sectored colonies. In addition, in the YYy311 strain on the W303-1A-derived homolog, the previously-described HS4 hotspot (St. Charles and Petes, 2013) was flanked by *hphMX4* and *URA3* markers. In both YYy310 and YYy311, the *MATα* locus was deleted to allow G_1_-synchronization with α-factor. This deletion also prevents the strains from undergoing meiosis in normal growth conditions. As noted below, however, such strains can be induced to undergo meiosis using special sporulation conditions.

### Isolation of granddaughter cells derived from red/white sectors

Most of the details of this analysis are described in the main text. In brief, we purified about 10-20 individual colonies from the red and white sectors. We analyzed one purified colony by SNP-specific microarrays to determine the approximate location of the selected recombination event. We then identified SNPs within these regions that could be assayed by a combination of PCR followed by treatment of the resulting PCR fragment with a diagnostic restriction enzyme. This procedure was previously used to detect LOH events on chromosome V (Lee *et al*., 2009). We employed this type of analysis with about 10-20 individual red and white colonies derived from each sector using the PCR primers and restriction enzymes listed in Table S5. If two PCR samples produced different fragment sizes after treatment with the restriction enzyme, we concluded that these strains represented the two granddaughters within the sector. We were able to detect most granddaughter cells using SNPs located on chromosomes V (YYy310) or IV (YYy311). However, for some sectored colonies, we used SNPs located on other chromosomes (Table S5). For some chromosomal regions, the SNPs did not result in an altered restriction enzyme recognition site. Consequently, for some events, we examined SNPs by sequencing PCR fragments from the relevant regions of the genome. The primers used in this analysis are also shown in Table S5.

### Meiotic analysis of marker coupling

For granddaughter cells with multiple heterozygous regions interspersed with homozygous regions (for example, Fig. 6A), we determined the coupling of these regions by examining microarrays of single spore cultures derived from the four granddaughters. Although the diploidsYYy310 and YYy311 lack *MATα* locus, these strains can be sporulated in medium containing 5 mM nicotinamide (St. Charles *et al*., 2012). By examining a single spore, we could generally determine the coupling relationships of the markers in each granddaughter. For example, the pattern of hybridization to SNPs shown for the spore in Fig. 68 demonstrates that the two regions of heteroduplex in the granddaughter cell are on different homologs. For most of the granddaughter cells, the SNP patterns of one daughter spore allowed unambiguous conclusions. However, if the pattern of SNPs in the spore was inconsistent with that observed in the diploid or if additional crossovers were observed within 50 kb of the mitotic recombination event, we considered the possibility of a meiotic crossover. Consequently, for such events, we examined the segregation pattern of SNPs flanking the region of mitotic exchange in several additional spores derived from the granddaughter cells. The primer pairs and restriction enzymes used in this analysis are described in Table S5. Complicating meiotic recombination events were rare, as expected, because the *mlh1* mutation substantially reduces the frequency of meiotic crossovers in yeast (Hunter and Borts, 1997).

Spontaneous red/white sectors generally had no unselected events. To determine coupling relationships within heterozygous regions, we used a different procedure from that described for the UV-induced events. We screened for monosomy on the chromosome with the selected event using a marker located centromere-proximal to the exchange. In YYy310, the heterozygous *URA3* gene was on the same chromosome arm (left arm of V) as the selected crossover, but was located centromere-proximal to the mitotic exchange. We selected for loss of the copy of V that had the *URA3* gene by using medium containing 5-fluoro-orotate. We confirmed the loss using PCR analysis, and restriction enzyme digestion of the resulting PCR product. Chromosome V-specific SNP microarrays were then used to determine the coupling of markers along the remaining homolog. A similar procedure was used to examine spontaneous events on yeast chromosome IV in YYy311. In YYy311, we selected monosomic strains that lacked the heterozygous *TRP1* marker located near *CEN4* using medium containing 5-fluoroanthranilic acid (Toyn *et al*., 2000).

### SNP microarrays

We used three types of SNP-Microarrays in our analysis: a whole-genome microarray (St. Charles *et al*., 2012) and microarrays to SNPs on chromosome IV (St. Charles and Petes, 2013) and chromosome V (Yin and Petes, 2013). The sequences of the oligonucleotides in the microarrays and their designs are on the Gene Expression Omnibus Website (https://www.ncbi.nlm.nih.gov/geo/) under the addresses GPL20144 (whole-genome), GPL21552 (chromosome IV), and GPL21274 (chromosome V). In brief, for each heterozygous SNP, we designed four 25-base oligonucleotides, two identical to the Watson and Crick strands of the W303-1A allele and two identical to the YJM789 allele. The heterozygous SNP was located near the middle of each oligonucleotide. Since the efficiency of hybridization is higher when genomic DNA is perfectly matched to the oligonucleotide than when there is a mismatch between the genomic DNA and the oligonucleotide, by measuring the ratio of hybridization of the experimental DNA to a control heterozygous strain for the four SNP-specific oligonucleotides, we can determine whether the genomic DNA is heterozygous for SNP or homozygous for either allele. In our experiments, the ratios of hybridization to the different oligonucleotides were normalized by comparing these ratios to a control heterozygous diploid. Genomic DNA from the experimental strains (derived from YYy310 or YYy311) was labeled with the Cy5-dUTP fluorescent nucleotide and DNA from an isogenic wild-type strain was labeled with Cy3-dUTP. The labeled samples were mixed and hybridized to the microarray. Details concerning the hybridization conditions and determining the levels of hybridization to the oligonucleotide probes are in St. Charles *et al*. (2012) and St. Charles *et al*. (2013). The data for all of these hybridization experiments are on the GEO Website (ref. number).

### Calculation of conversion/heteroduplex tract lengths

Two types of conversion/heteroduplex events were observed in our studies: events unassociated with crossovers and events associated with crossovers. For the first class, we averaged the distance between the closest heterozygous SNPs flanking the tract (the maximum tract length) and the homozygous SNPs located at the borders of the conversion/heteroduplex tract (the minimum tract length). For crossover-associated conversion/heteroduplex tracts, we averaged the distance between the homozygous SNPs flanking the tract (the maximum tract length), and the distance between the SNPs that were within the tract closest to the borders of the crossover.

### Bioinformatics and statistical analysis

The two haploid strains for W303 and YJM789 that were sequenced previously (St.Charles *et al*., 2012) were analyzed for the number of SNPs and insertions/deletions (in/dels) in gene conversion tracts. Paired-end reads were aligned to the S288c reference genome version 3 (downloaded from sad Cer3 in UCSC Genome Browser; https://genome.ucsc.edu) by BWA-MEM software (Li, 2013) and SNPs were determined using samtools (Li *et al*., 2009). Because our SNP-Microarray was based on S288c reference genome version 2 (downloaded from sacCer2 in UCSC Genome Browse), we translated the positions of the SNPs from the microarray to the version 3 reference genome before counting the number of SNPs and indels within each of the conversion tracts. We then used BEDTools (Quinlan and Hall, 2010) to count the number of SNPs within the conversion tracts. RStudio (http://www.rstudio.com/ used for various statistical tests used in this study.

## Acknowledgements

We thank members of the Petes and Jinks-Robertson labs for suggestions during the course of this work, and Sue Jinks-Robertson, Yee Fang Hum, Dao-Qiong Zheng and Ke Zhang for comments or help with the manuscript. The research was supported by NIH grants GM24110, GM52319, 1R35GM118020, and 5T32GM007754-37.

---

**Table S1.**
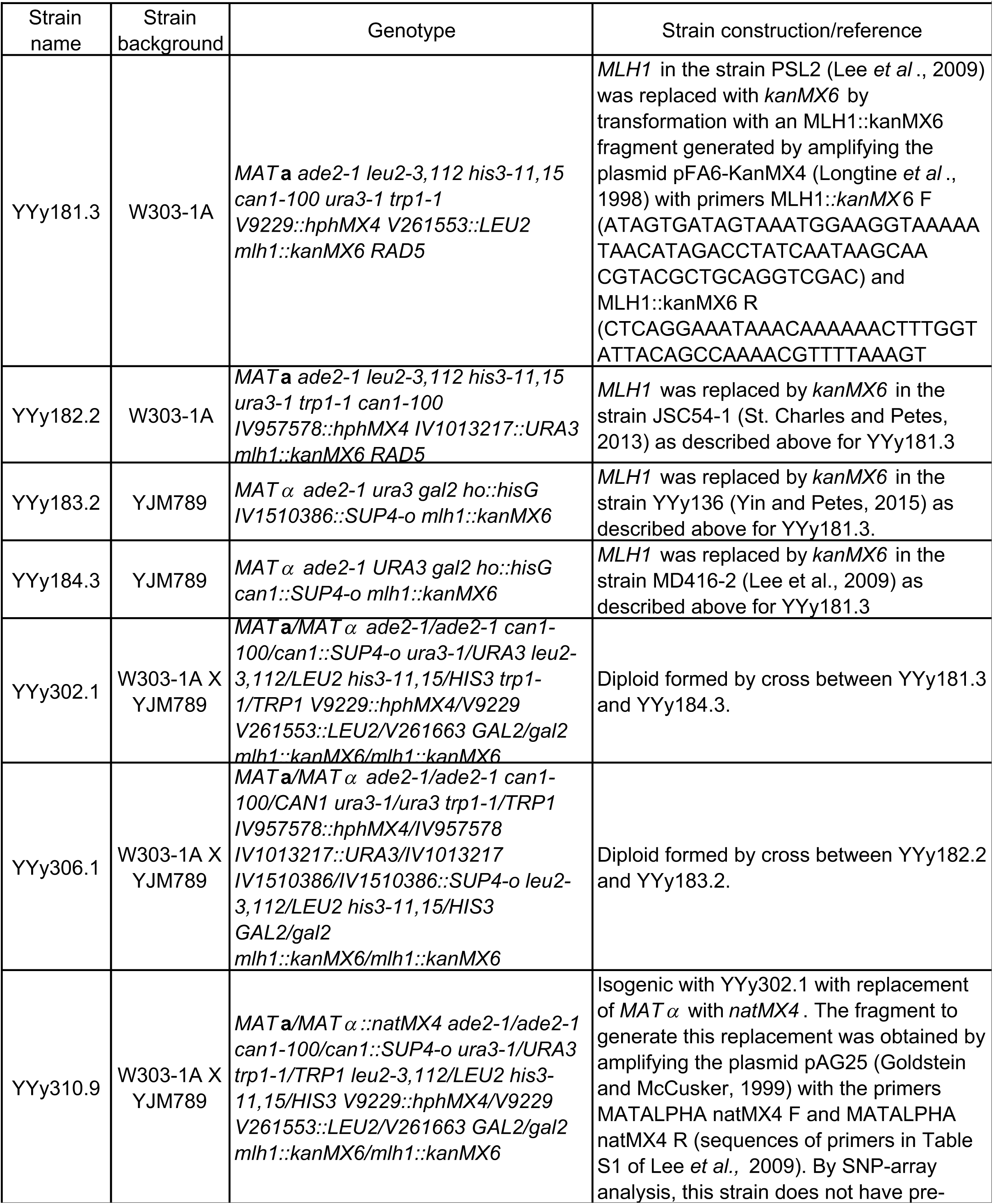
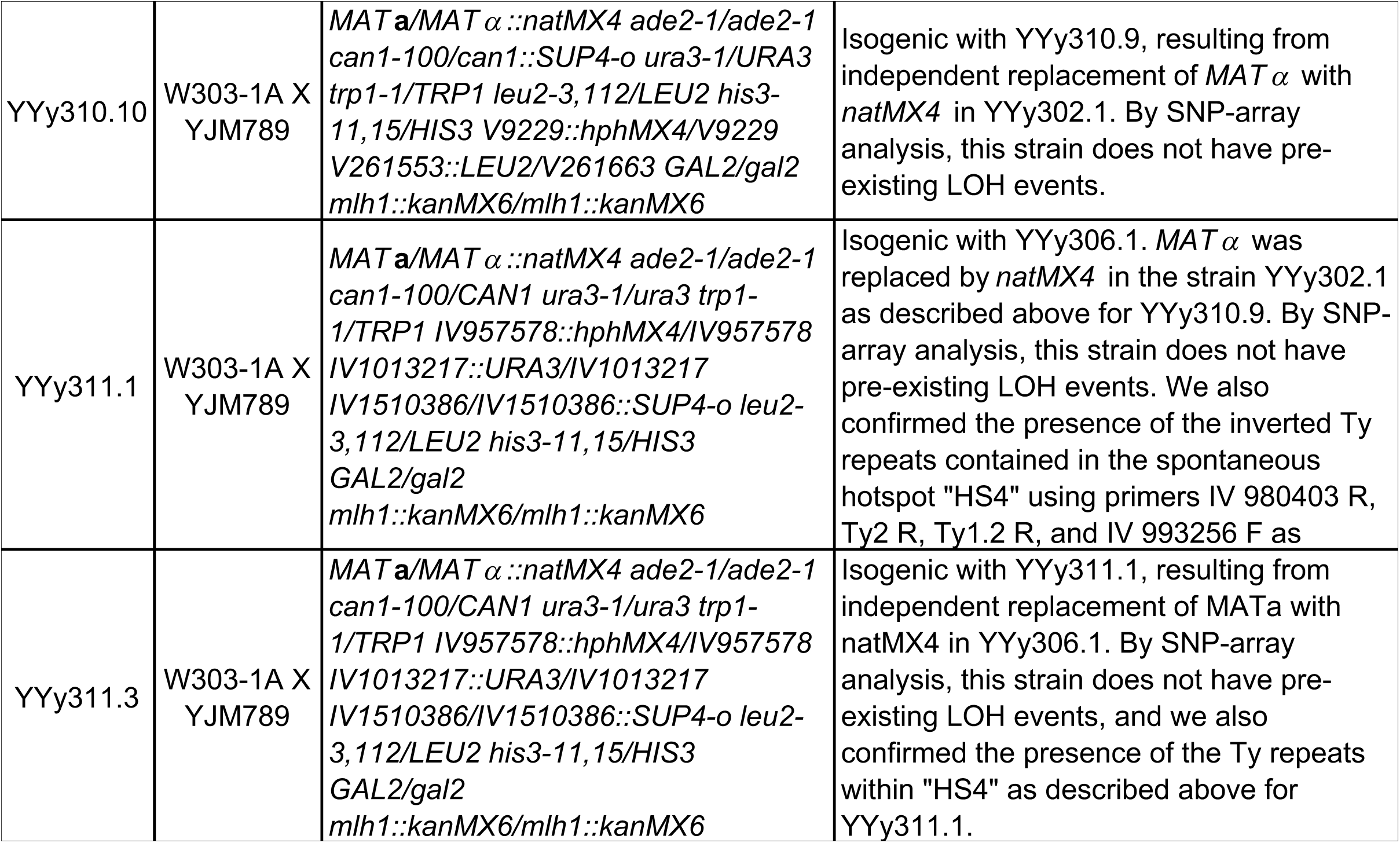
Strain genotypes and constructions.

**Table S2.** SGD coordinates for LOH transitions in the *mlh1* strain YYy310 induced by a UV dose of 15 J/m^2^ in G1-synchronized cells (Figs. S3-S80), spontaneous sectors in YYy310 selected on the left arm of chromosome V (Figs. S81-S85) and spontaneous sectors in YYy311 selected on the right arm of chromosome IV (Figs. S86-S91).

| Sector | Chromosome | Supplementary picture | Sub sector | Transition genotype |  | SNPs flanking transitions |  |
| --- | --- | --- | --- | --- | --- | --- | --- |
|  |  |  |  | Left | Right | Left | Right |
| YYy310-9-5WR | II | S3 | 5W1 | het | S | 235624 | 236376 |
| UV |  |  |  | S | het | 236383 | 236479 |
|  |  |  |  | het | Y | 248647 | 249558 |
|  |  |  |  | Y | het | 250304 | 251093 |
|  |  |  | 5W2 | het | het |  |  |
|  |  |  | 5R1 | het | het |  |  |
|  |  |  | 5R2 | het | Y | 238664 | 239853 |
|  |  |  |  | Y | het | 239940 | 240829 |
|  |  |  |  | het | Y | 240829 | 241471 |
|  |  |  |  | Y | het | 241471 | 241694 |
|  |  |  |  | het | Y | 241694 | 242027 |
|  |  |  |  | Y | het | 249558 | 250304 |
| YYy310-9-5WR | III | S4 | 5W1 | het | S | 60451 | 64292 |
| UV |  |  |  | S | het | 64292 | 67656 |
|  |  |  | 5W2 | het | S | 60451 | 64292 |
|  |  |  |  | S | het | 64292 | 67656 |
|  |  |  | 5R1 | het | het |  |  |
|  |  |  | 5R2 | het | het |  |  |
| YYy310-9-5WR | IV | S5 | 5W1 | het | het |  |  |
| UV |  |  | 5W2 | het | het |  |  |
|  |  |  | 5R1 | het | S | 325063 | 327230 |
|  |  |  |  | S | het | 330624 | 331783 |
|  |  |  | 5R2 | het | S | 325063 | 327230 |
|  |  |  |  | S | het | 329222 | 330624 |
| YYy310-9-5WR | IV | S6 | 5W1 | het | het |  |  |
| UV |  |  | 5W2 | het | het |  |  |
|  |  |  | 5R1 | het | het |  |  |
|  |  |  | 5R2 | het | Y | 356529 | 360199 |
|  |  |  |  | Y | het | 360199 | 361086 |
| YYy310-9-5WR | IV | S7 | 5W1 | het | het |  |  |
| UV |  |  | 5W2 | het | S | 505764 | 507165 |
|  |  |  |  | S | Y | 507165 | 507960 |
|  |  |  |  | Y | het | 512801 | 521609 |
|  |  |  | 5R1 | het | Y | 505764 | 507165 |
|  |  |  |  | Y | het | 509817 | 512801 |
|  |  |  | 5R2 | het | Y | 505764 | 507165 |
|  |  |  |  | Y | het | 509817 | 512801 |
| YYy310-9-5WR | V | S8 | 5W1 | Y | HET | 51915 | 53692 |
| UV |  |  |  | HET | S | 54198 | 56117 |
|  |  |  |  | S | het | 57170 | 60701 |
|  |  |  | 5W2 | Y | HET | 45587 | 48837 |
|  |  |  |  | HET | het | 51915 | 53692 |
|  |  |  |  | het | S | 54198 | 56117 |
|  |  |  |  | S | het | 57170 | 60701 |
|  |  |  | 5R1 | S | het | 57170 | 60701 |
|  |  |  | 5R2 | S | het | 51915 | 53692 |
| YYy310-9-5WR | VI | S9 | 5W1 | het | S | 185398 | 186565 |
| UV |  |  |  | S | het | 186912 | 189728 |
|  |  |  | 5W2 | het | S | 180526 | 185398 |
|  |  |  |  | S | het | 186912 | 189728 |
|  |  |  | 5R1 | het | het |  |  |
|  |  |  | 5R2 | het | S | 186643 | 186862 |
|  |  |  |  | S | het | 189839 | 191415 |
| YYy310-9-5WR | VII | S10 | 5W1 | het | S | 157428 | 159699 |
| UV |  |  |  | S | het | 159699 | 162937 |
|  |  |  | 5W2 | het | het |  |  |
|  |  |  | 5R1 | het | het |  |  |
|  |  |  | 5R2 | het | het |  |  |
| YYy310-9-5WR | VIII | S11 | 5W1 | het | het |  |  |
| UV |  |  | 5W2 | het | het |  |  |
|  |  |  | 5R1 | het | HET | 108370 | 110004 |
|  |  |  | 5R2 | het | Y | 108370 | 110004 |
|  |  |  |  | Y | HET | 110004 | 111606 |
| YYy310-9-5WR | X | S12 | 5W1 | het | het |  |  |
| UV |  |  | 5W2 | het | Y | 314934 | 315219 |
|  |  |  |  | Y | het | 320668 | 320792 |
|  |  |  | 5R1 | het | Y | 315219 | 317971 |
|  |  |  |  | Y | het | 323059 | 323805 |
|  |  |  | 5R2 | het | Y | 315219 | 317971 |
|  |  |  |  | Y | het | 320668 | 320792 |
| YYy310-9-5WR | XI | S13 | 5W1 | het | Y | 46433 | 46576 |
| UV |  |  |  | Y | het | 51547 | 52577 |
|  |  |  | 5W2 | het | het |  |  |
|  |  |  | 5R1 | het | Y | 50384 | 51547 |
|  |  |  |  | Y | het | 51547 | 52577 |
|  |  |  | 5R2 | het | Y | 46123 | 46433 |
|  |  |  |  | Y | het | 47424 | 50384 |
| YYy310-9-5WR | XII | S14 | 5W1 | het | S | 897133 | 897352 |
| UV |  |  |  | S | het | 901814 | 902318 |
|  |  |  | 5W2 | het | S | 890066 | 893936 |
|  |  |  |  | S | het | 901814 | 902318 |
|  |  |  | 5R1 | het | het |  |  |
|  |  |  | 5R2 | het | S | 897133 | 897352 |
|  |  |  |  | S | het | 898548 | 899569 |
| YYy310-9-5WR | XV | S15 | 5W1 | het | S | 1019448 | 1022697 |
| UV |  |  |  | S | het | 1023392 | 1024031 |
|  |  |  | 5W2 | het | het |  |  |
|  |  |  | 5R1 | het | S | 1014379 | 1018892 |
|  |  |  |  | S | het | 1023392 | 1024031 |
|  |  |  | 5R2 | het | het |  |  |
| YYy310-9-5WR | XVI | S16 | 5W1 | HET | S | 162425 | 165200 |
| UV |  |  |  | S | het | 165200 | 171020 |
|  |  |  | 5W2 | HET | S | 158225 | 162425 |
|  |  |  |  | S | het | 162425 | 165200 |
|  |  |  | 5R1 | het | S | 158225 | 162425 |
|  |  |  |  | S | het | 165200 | 171020 |
|  |  |  | 5R2 | het | het |  |  |
| YYy310-9-12WR | II | S17 | 12W1 | het | S | 407335 | 410831 |
| UV |  |  |  | S | het | 410985 | 411148 |
|  |  |  | 12W2 | het | S | 403952 | 406262 |
|  |  |  |  | S | het | 410985 | 411148 |
|  |  |  | 12R1 | het | het |  |  |
|  |  |  | 12R2 | het | S | 407335 | 410831 |
|  |  |  |  | S | het | 411214 | 420497 |
| YYy310-9-12WR | IV | S18 | 12W1 | het | Y | 29689 | 31400 |
| UV |  |  |  | Y | het | 33241 | 37025 |
|  |  |  | 12W2 | het | het |  |  |
|  |  |  | 12R1 | het | het |  |  |
|  |  |  | 12R2 | het | het |  |  |
| YYy310-9-12WR | IV | S19 | 12W1 | het | het |  |  |
| UV |  |  | 12W2 | het | het |  |  |
|  |  |  | 12R1 | het | Y | 426766 | 427154 |
|  |  |  |  | Y | het | 427315 | 427981 |
|  |  |  | 12R2 | het | Y | 426766 | 427154 |
|  |  |  |  | Y | het | 427315 | 427981 |
| YYy310-9-12WR | IV | S20 | 12W1 | het | het |  |  |
| UV |  |  | 12W2 | het | S | 454186 | 455000 |
|  |  |  |  | S | het | 458195 | 469765 |
|  |  |  | 12R1 | het | het |  |  |
|  |  |  | 12R2 | het | het |  |  |
| YYy310-9-12WR | V | S21 | 12W1 | Y | het | 50196 | 50295 |
| UV |  |  | 12W2 | Y | het | 51768 | 53692 |
|  |  |  | 12R1 | S | het | 50196 | 50295 |
|  |  |  | 12R2 | S | het | 50196 | 50295 |
| YYy310-9-12WR | VII | S22 | 12W1 | het | Y | 481769 | 482414 |
| UV |  |  |  | Y | het | 482414 | 484837 |
|  |  |  | 12W2 | het | Y | 481769 | 482414 |
|  |  |  |  | Y | het | 482414 | 484837 |
|  |  |  | 12R1 | het | het |  |  |
|  |  |  | 12R2 | het | het |  |  |
| YYy310-9-12WR | VIII | S23 | 12W1 | het | Y | 13263 | 13512 |
| UV |  |  |  | Y | het | 16312 | 17106 |
|  |  |  | 12W2 | het | Y | 13263 | 13512 |
|  |  |  |  | Y | het | 16312 | 17106 |
|  |  |  | 12R1 | het | het |  |  |
|  |  |  | 12R2 | het | het |  |  |
| YYy310-9-12WR | VIII | S24 | 12W1 | het | HET | 388782 | 393258 |
| UV |  |  | 12W2 | het | HET | 388782 | 393258 |
|  |  |  | 12R1 | het | HET | 388782 | 393258 |
|  |  |  | 12R2 | het | S | 378185 | 378225 |
|  |  |  |  | S | het | 382724 | 388782 |
|  |  |  |  | het | S | 388782 | 393258 |
|  |  |  |  | S | HET | 397033 | 399358 |
| YYy310-9-12WR | X | S25 | 12W1 | het | Y | 76156 | 77570 |
| UV |  |  |  | Y | het | 77717 | 78260 |
|  |  |  | 12W2 | het | Y | 76156 | 77570 |
|  |  |  |  | Y | het | 77717 | 78260 |
|  |  |  | 12R1 | het | het |  |  |
|  |  |  | 12R2 | het | het |  |  |
| YYy310-9-12WR | XI | S26 | 12W1 | het | HET | 74554 | 76907 |
| UV |  |  |  | HET | het | 78118 | 78701 |
|  |  |  |  | het | S | 78701 | 79174 |
|  |  |  |  | S | HET | 79174 | 79722 |
|  |  |  |  | HET | S | 79722 | 80717 |
|  |  |  |  | S | het | 81237 | 83093 |
|  |  |  |  | het | S | 83992 | 84050 |
|  |  |  |  | S | het | 84071 | 84394 |
|  |  |  | 12W2 | het | S | 78701 | 79174 |
|  |  |  |  | S | het | 79174 | 79722 |
|  |  |  |  | het | S | 79722 | 80717 |
|  |  |  |  | S | het | 84426 | 84604 |
|  |  |  |  | het | S | 84604 | 85461 |
|  |  |  |  | S | het | 85830 | 87504 |
|  |  |  | 12R1 | HET | S | 84604 | 85461 |
|  |  |  |  | S | het | 85830 | 87504 |
|  |  |  | 12R2 | HET | het | 85830 | 87504 |
| YYy310-9-12WR | XII | S27 | 12W1 | het | Y | 188741 | 190430 |
| UV |  |  |  | Y | het | 190736 | 191599 |
|  |  |  | 12W2 | het | Y | 188741 | 190430 |
|  |  |  |  | Y | het | 190736 | 191599 |
|  |  |  | 12R1 | het | het |  |  |
|  |  |  | 12R2 | het | het |  |  |
| YYy310-9-12WR | XII | S28 | 12W1 | het | S | 249041 | 249624 |
| UV |  |  |  | S | HET | 249888 | 250394 |
|  |  |  | 12W2 | het | HET | 249041 | 249624 |
|  |  |  | 12R1 | het | S | 249041 | 249624 |
|  |  |  |  | S | het | 250968 | 253263 |
|  |  |  | 12R2 | het | S | 249041 | 249624 |
|  |  |  |  | S | het | 250968 | 253263 |
| YYy310-9-12WR | XII | S29 | 12W1 | het | het |  |  |
| UV |  |  | 12W2 | het | S | 665333 | 665741 |
|  |  |  |  | S | het | 665742 | 677689 |
|  |  |  | 12R1 | het | Y | 657904 | 665226 |
|  |  |  |  | Y | het | 665742 | 677689 |
|  |  |  | 12R2 | het | Y | 657904 | 665226 |
|  |  |  |  | Y | het | 665742 | 677689 |
| YYy310-9-12WR | XII | S30 | 12W1 | het | het |  |  |
| UV |  |  | 12W2 | het | het |  |  |
|  |  |  | 12R1 | het | S | 693776 | 694201 |
|  |  |  |  | S | het | 706779 | 706956 |
|  |  |  | 12R2 | het | S | 693776 | 694201 |
|  |  |  |  | S | het | 696985 | 697188 |
| YYy310-9-12WR | XII | S31 | 12W1 | het | S | 834095 | 835000 |
| UV |  |  |  | S | het | 835000 | 835184 |
|  |  |  | 12W2 | het | S | 834095 | 835000 |
|  |  |  |  | S | het | 835000 | 835184 |
|  |  |  | 12R1 | het | S | 833747 | 833877 |
|  |  |  |  | S | het | 835000 | 835184 |
|  |  |  | 12R2 | het | S | 833877 | 833970 |
|  |  |  |  | S | het | 835000 | 835184 |
| YYy310-9-12WR | XIII | S32 | 12W1 | het | het |  |  |
| UV |  |  | 12W2 | het | Y | 296414 | 303742 |
|  |  |  |  | Y | het | 303910 | 307056 |
|  |  |  | 12R1 | het | het |  |  |
|  |  |  | 12R2 | het | het |  |  |
| YYy310-9-12WR | XIII | S33 | 12W1 | het | het |  |  |
| UV |  |  | 12W2 | het | het |  |  |
|  |  |  | 12R1 | het | Y | 370211 | 370688 |
|  |  |  |  | Y | het | 371928 | 378717 |
|  |  |  | 12R2 | het | Y | 370211 | 370688 |
|  |  |  |  | Y | het | 371928 | 378717 |
| YYy310-9-12WR | XVI | S34 | 12W1 | het | Y | 99390 | 101743 |
| UV |  |  |  | Y | het | 105255 | 105779 |
|  |  |  | 12W2 | het | Y | 99390 | 101743 |
|  |  |  |  | Y | het | 105941 | 108288 |
|  |  |  | 12R1 | het | Y | 99390 | 101743 |
|  |  |  |  | Y | het | 104035 | 104416 |
|  |  |  | 12R2 | het | Y | 105255 | 105779 |
|  |  |  |  | Y | het | 105941 | 108288 |
| YYy310-9-12WR | XVI | S35 | 12W1 | het | het |  |  |
| UV |  |  | 12W2 | het | het |  |  |
|  |  |  | 12R1 | het | S | 635100 | 637439 |
|  |  |  |  | S | het | 639408 | 644553 |
|  |  |  | 12R2 | het | S | 635100 | 637439 |
|  |  |  |  | S | het | 639408 | 644553 |
| YYy310-9-17WR | I | S36 | 17W1 | het | het |  |  |
| UV |  |  | 17W2 | het | Y | 124574 | 124635 |
|  |  |  |  | Y | het | 125371 | 125782 |
|  |  |  | 17R1 | het | het |  |  |
|  |  |  | 17R2 | het | het |  |  |
| YYy310-9-17WR | II | S37 | 17W1 | Y | het | 74904 | 79448 |
| UV |  |  | 17W2 | Y | het | 74904 | 79448 |
|  |  |  | 17R1 | S | het | 74904 | 79448 |
|  |  |  | 17R2 | S | het | 68229 | 69213 |
| YYy310-9-17WR | II | S38 | 17W1 | het | S | 173854 | 174748 |
| UV |  |  |  | S | het | 175099 | 175399 |
|  |  |  | 17W2 | het | S | 173854 | 174748 |
|  |  |  |  | S | het | 175099 | 175399 |
|  |  |  | 17R1 | het | S | 173854 | 174748 |
|  |  |  |  | S | het | 175099 | 175399 |
|  |  |  |  | het | Y | 179400 | 180245 |
|  |  |  |  | Y | het | 180245 | 182118 |
|  |  |  | 17R2 | het | het |  |  |
| YYy310-9-17WR | II | S39 | 17W1 | het | het |  |  |
| UV |  |  | 17W2 | het | S | 420548 | 427526 |
|  |  |  |  | S | het | 429164 | 431460 |
|  |  |  | 17R1 | het | het |  |  |
|  |  |  | 17R2 | het | het |  |  |
| YYy310-9-17WR | IV | S40 | 17W1 | het | het |  |  |
| UV |  |  | 17W2 | het | Y | 173405 | 184716 |
|  |  |  |  | Y | het | 184727 | 188296 |
|  |  |  | 17R1 | het | het |  |  |
|  |  |  | 17R2 | het | het |  |  |
| YYy310-9-17WR | IV | S41 | 17W1 | HET | Y | 409879 | 410333 |
| UV |  |  |  | Y | het | 417000 | 420543 |
|  |  |  | 17W2 | HET | Y | 409879 | 410333 |
|  |  |  |  | Y | het | 417000 | 420543 |
|  |  |  | 17R1 | het | HET | 409879 | 410333 |
|  |  |  |  | HET | S | 414508 | 415063 |
|  |  |  |  | S | het | 417000 | 420543 |
|  |  |  | 17R2 | het | HET | 409879 | 410333 |
|  |  |  |  | HET | het | 410349 | 413580 |
|  |  |  |  | het | S | 413580 | 414508 |
|  |  |  |  | S | het | 414508 | 415063 |
| YYy310-9-17WR | IV | S42 | 17W1 | het | Y | 1040924 | 1044556 |
| UV |  |  |  | Y | het | 1046711 | 1053213 |
|  |  |  | 17W2 | het | Y | 1040924 | 1044556 |
|  |  |  |  | Y | het | 1044648 | 1046711 |
|  |  |  | 17R1 | het | het |  |  |
|  |  |  | 17R2 | het | het |  |  |
| YYy310-9-17WR | V | S43 | 17W1 | Y | het | 78080 | 80291 |
| UV |  |  | 17W2 | Y | het | 78080 | 80291 |
|  |  |  | 17R1 | S | HET | 48837 | 49044 |
|  |  |  |  | HET | S | 60913 | 61808 |
|  |  |  |  | S | HET | 62013 | 62494 |
|  |  |  |  | HET | Y | 76261 | 77288 |
|  |  |  |  | Y | S | 78080 | 80291 |
|  |  |  |  | S | het | 80291 | 81264 |
|  |  |  | 17R2 | S | het | 72490 | 75719 |
|  |  |  |  | het | S | 78080 | 80291 |
|  |  |  |  | S | het | 80291 | 81264 |
| YYy310-9-17WR | V | S44 | 17W1 | het | het |  |  |
| UV |  |  | 17W2 | het | Y | 501972 | 502134 |
|  |  |  |  | Y | het | 502978 | 504162 |
|  |  |  |  | het | Y | 507136 | 507433 |
|  |  |  |  | Y | het | 507627 | 508392 |
|  |  |  | 17R1 | het | Y | 504309 | 504693 |
|  |  |  |  | Y | het | 504917 | 506853 |
|  |  |  |  | het | Y | 507433 | 507484 |
|  |  |  |  | Y | het | 507627 | 508392 |
|  |  |  | 17R2 | het | Y | 504309 | 504693 |
|  |  |  |  | Y | het | 504917 | 506853 |
|  |  |  |  | het | Y | 507136 | 507433 |
|  |  |  |  | Y | het | 507627 | 508392 |
| YYy310-9-17WR | IX | S45 | 17W1 | S | het | 132136 | 132892 |
| UV |  |  | 17W2 | S | HET | 119675 | 122608 |
|  |  |  |  | HET | het | 122608 | 129791 |
|  |  |  | 17R1 | Y | het | 119675 | 122608 |
|  |  |  |  | het | Y | 122608 | 129791 |
|  |  |  |  | Y | het | 132136 | 132892 |
|  |  |  | 17R2 | Y | het | 122608 | 129791 |
| YYy310-9-17WR | XV | S46 | 17W1 | het | Y | 182373 | 186468 |
| UV |  |  |  | Y | het | 196650 | 198759 |
|  |  |  | 17W2 | het | Y | 182373 | 186468 |
|  |  |  |  | Y | het | 196650 | 198759 |
|  |  |  | 17R1 | het | Y | 182373 | 186468 |
|  |  |  |  | Y | het | 196650 | 198759 |
|  |  |  | 17R2 | het | Y | 182373 | 186468 |
|  |  |  |  | Y | het | 196650 | 198759 |
| YYy310-10-14WR | II | S47 | 14W1 | het | S | 584379 | 584616 |
| UV |  |  |  | S | het | 586395 | 586943 |
|  |  |  | 14W2 | het | S | 584379 | 584616 |
|  |  |  |  | S | het | 586395 | 586943 |
|  |  |  | 14R1 | het | S | 584379 | 584616 |
|  |  |  |  | S | het | 586395 | 586943 |
|  |  |  | 14R2 | het | S | 584379 | 584616 |
|  |  |  |  | S | het | 586395 | 586943 |
| YYy310-10-14WR | V | S48 | 14W1 | Y | het | 94077 | 94339 |
| UV |  |  |  | het | Y | 95429 | 95684 |
|  |  |  |  | Y | het | 95906 | 96550 |
|  |  |  | 14W2 | Y | het | 91739 | 93200 |
|  |  |  | 14R1 | S | het | 80291 | 81264 |
|  |  |  | 14R2 | S | het | 80291 | 81264 |
|  |  |  |  | het | S | 85566 | 86732 |
|  |  |  |  | S | het | 90062 | 90200 |
|  |  |  |  | het | S | 90215 | 91703 |
|  |  |  |  | S | het | 91739 | 93200 |
| YYy310-10-14WR | IX | S49 | 14W1 | het | Y | 235680 | 236609 |
| UV |  |  |  | Y | het | 239260 | 240253 |
|  |  |  | 14W2 | het | Y | 235680 | 236609 |
|  |  |  |  | Y | het | 239260 | 240253 |
|  |  |  | 14R1 | het | het |  |  |
|  |  |  | 14R2 | het | het |  |  |
| YYy310-10-14WR | XI | S50 | 14W1 | het | het |  |  |
| UV |  |  | 14W2 | het | het |  |  |
|  |  |  | 14R1 | het | Y | 105100 | 112922 |
|  |  |  |  | Y | het | 113668 | 114642 |
|  |  |  | 14R2 | het | het |  |  |
| YYy310-10-14WR | XII | S51 | 14W1 | het | het |  |  |
| UV |  |  | 14W2 | het | het |  |  |
|  |  |  | 14R1 | het | S | 757465 | 757753 |
|  |  |  |  | S | het | 759448 | 760629 |
|  |  |  | 14R2 | het | het |  |  |
| YYy310-10-14WR | XIII | S52 | 14W1 | het | het |  |  |
| UV |  |  | 14W2 | het | het |  |  |
|  |  |  | 14R1 | het | S | 368237 | 368927 |
|  |  |  |  | S | het | 369406 | 370211 |
|  |  |  | 14R2 | het | S | 367185 | 367797 |
|  |  |  |  | S | het | 369406 | 370211 |
| YYy310-10-14WR | XIII | S53 | 14W1 | het | Y | 612380 | 613145 |
| UV |  |  |  | Y | het | 613247 | 613271 |
|  |  |  | 14W2 | het | Y | 612380 | 613145 |
|  |  |  |  | Y | het | 613247 | 613271 |
|  |  |  | 14R1 | het | Y | 613271 | 613724 |
|  |  |  |  | Y | het | 613724 | 614157 |
|  |  |  | 14R2 | het | Y | 613271 | 613724 |
|  |  |  |  | Y | het | 613724 | 614157 |
| YYy310-10-14WR | XV | S54 | 14W1 | het | Y | 1004852 | 1009367 |
| UV |  |  |  | Y | het | 1009367 | 1014379 |
|  |  |  | 14W2 | het | het |  |  |
|  |  |  | 14R1 | het | Y | 1004852 | 1009367 |
|  |  |  |  | Y | het | 1009367 | 1014379 |
|  |  |  | 14R2 | het | Y | 1004852 | 1009367 |
|  |  |  |  | Y | het | 1009367 | 1014379 |
| YYy310-10-14WR | XVI | S55 | 14W1 | het | S | 901548 | 907111 |
| UV |  |  |  | S | het | 912806 | 918183 |
|  |  |  | 14W2 | het | S | 901548 | 907111 |
|  |  |  |  | S | het | 911222 | 912157 |
|  |  |  | 14R1 | het | S | 901548 | 907111 |
|  |  |  |  | S | het | 909996 | 911222 |
|  |  |  | 14R2 | het | S | 901548 | 907111 |
|  |  |  |  | S | het | 912806 | 918183 |
| YYy310-10-17WR | IV | S56 | 17W1 | het | Y | 248076 | 249496 |
| UV |  |  |  | Y | het | 249496 | 251661 |
|  |  |  | 17W2 | het | Y | 248076 | 249496 |
|  |  |  |  | Y | het | 249496 | 251661 |
|  |  |  | 17R1 | het | Y | 232819 | 233059 |
|  |  |  |  | Y | het | 244267 | 246953 |
|  |  |  | 17R2 | het | Y | 242167 | 244267 |
|  |  |  |  | Y | het | 244267 | 246953 |
|  |  |  |  | het | Y | 248076 | 249496 |
|  |  |  |  | Y | het | 249496 | 251661 |
| YYy310-10-17WR | IV | S57 | 17W1 | het | het |  |  |
| UV |  |  | 17W2 | het | het |  |  |
|  |  |  | 17R1 | het | Y | 526934 | 540214 |
|  |  |  |  | Y | het | 540214 | 543869 |
|  |  |  | 17R2 | het | Y | 526934 | 540214 |
|  |  |  |  | Y | het | 540214 | 543869 |
| YYy310-10-17WR | IV | S58 | 17W1 | het | HET | 608319 | 611611 |
| UV |  |  |  | HET | Y | 611611 | 618459 |
|  |  |  | 17W2 | het | Y | 608319 | 611611 |
|  |  |  | 17R1 | het | S | 608319 | 611611 |
|  |  |  | 17R2 | het | S | 608319 | 611611 |
| YYy310-10-17WR | V | S59 | 17W1 | Y | HET | 121397 | 124190 |
| UV |  |  |  | HET | Y | 124505 | 124754 |
|  |  |  |  | Y | HET | 127038 | 128941 |
|  |  |  |  | HET | het | 131261 | 132281 |
|  |  |  | 17W2 | Y | het | 131261 | 132281 |
|  |  |  | 17R1 | S | het | 126304 | 127030 |
|  |  |  | 17R2 | S | het | 124754 | 125129 |
|  |  |  |  | het | Y | 126304 | 127030 |
|  |  |  |  | Y | het | 127038 | 128941 |
| YYy310-10-17WR | VIII | S60 | 17W1 | het | S | 139939 | 141697 |
| UV |  |  |  | S | het | 141697 | 144711 |
|  |  |  | 17W2 | het | Y | 117153 | 117396 |
|  |  |  |  | Y | het | 120428 | 139939 |
|  |  |  | 17R1 | het | het |  |  |
|  |  |  | 17R2 | het | het |  |  |
| YYy310-10-17WR | IX | S61 | 17W1 | HET | S | 108413 | 110547 |
| UV |  |  |  | S | het | 117444 | 119675 |
|  |  |  | 17W2 | HET | Y | 110547 | 111619 |
|  |  |  |  | Y | het | 115879 | 116682 |
|  |  |  | 17R1 | HET | het | 122608 | 129791 |
|  |  |  | 17R2 | HET | S | 117444 | 119675 |
|  |  |  |  | S | het | 122608 | 129791 |
| YYy310-10-17WR | X | S62 | 17W1 | S | HET | 237196 | 238853 |
| UV |  |  |  | HET | Y | 241092 | 242667 |
|  |  |  |  | Y | het | 242937 | 244592 |
|  |  |  | 17W2 | S | HET | 237196 | 238853 |
|  |  |  |  | HET | het | 242937 | 244592 |
|  |  |  | 17R1 | Y | het | 242937 | 244592 |
|  |  |  | 17R2 | Y | het | 237196 | 238853 |
|  |  |  |  | het | Y | 240830 | 242667 |
|  |  |  |  | Y | het | 242937 | 244592 |
| YYy310-10-17WR | XIII | S63 | 17W1 | het | het |  |  |
| UV |  |  | 17W2 | het | Y | 113881 | 114292 |
|  |  |  |  | Y | het | 114636 | 117330 |
|  |  |  | 17R1 | het | Y | 113881 | 114292 |
|  |  |  |  | Y | het | 114636 | 117330 |
|  |  |  | 17R2 | het | Y | 113881 | 114292 |
|  |  |  |  | Y | het | 114636 | 117330 |
| YYy310-10-18WR | II | S64 | 18W1 | het | het |  |  |
| UV |  |  | 18W2 | het | Y | 398295 | 399516 |
|  |  |  |  | Y | het | 400808 | 401593 |
|  |  |  | 18R1 | het | Y | 400808 | 401593 |
|  |  |  |  | Y | het | 401882 | 403262 |
|  |  |  | 18R2 | het | S | 400808 | 401593 |
|  |  |  |  | S | het | 401882 | 403262 |
| YYy310-10-18WR | V | S65 | 18W1 | Y | het | 44506 | 45284 |
| UV |  |  |  | het | Y | 45587 | 48837 |
|  |  |  |  | Y | het | 49385 | 49896 |
|  |  |  |  | het | Y | 50295 | 51183 |
|  |  |  |  | Y | het | 51183 | 51567 |
|  |  |  | 18W2 | Y | het | 37492 | 38068 |
|  |  |  | 18R1 | S | HET | 34235 | 34481 |
|  |  |  |  | HET | het | 34481 | 37492 |
|  |  |  |  | het | Y | 37492 | 38068 |
|  |  |  |  | Y | het | 41533 | 44506 |
|  |  |  | 18R2 | S | Y | 38068 | 41126 |
|  |  |  |  | Y | het | 41533 | 44506 |
| YYy310-10-18WR | VII | S66 | 18W1 | het | Y | 263911 | 264645 |
| UV |  |  |  | Y | het | 275506 | 275695 |
|  |  |  |  | het | HET | 277103 | 278627 |
|  |  |  |  | HET | het | 281296 | 284849 |
|  |  |  | 18W2 | het | Y | 262989 | 263911 |
|  |  |  |  | Y | het | 264645 | 267678 |
|  |  |  |  | het | Y | 268082 | 269269 |
|  |  |  |  | Y | het | 272660 | 273402 |
|  |  |  |  | het | Y | 277103 | 278627 |
|  |  |  |  | Y | het | 281296 | 284849 |
|  |  |  | 18R1 | het | S | 263911 | 264645 |
|  |  |  |  | S | het | 269529 | 272660 |
|  |  |  | 18R2 | het | het |  |  |
| YYy310-10-18WR | X | S67 | 18W1 | het | S | 690840 | 696274 |
| UV |  |  |  | S | het | 696274 | 697635 |
|  |  |  | 18W2 | het | S | 690840 | 696274 |
|  |  |  |  | S | het | 696274 | 697635 |
|  |  |  | 18R1 | het | Y | 690840 | 696274 |
|  |  |  |  | Y | het | 696274 | 697635 |
|  |  |  | 18R2 | het | het |  |  |
| YYy310-10-18WR | XII | S68 | 18W1 | het | S | 25673 | 27102 |
| UV |  |  |  | S | het | 27102 | 28812 |
|  |  |  | 18W2 | het | S | 25673 | 27102 |
|  |  |  |  | S | het | 29402 | 29600 |
|  |  |  | 18R1 | het | het |  |  |
|  |  |  | 18R2 | het | het |  |  |
| YYy310-10-18WR | XIV | S69 | 18W1 | het | S | 306383 | 310719 |
| UV |  |  |  | S | het | 310719 | 316099 |
|  |  |  | 18W2 | het | S | 301637 | 304112 |
|  |  |  |  | S | het | 310719 | 316099 |
|  |  |  | 18R1 | het | het |  |  |
|  |  |  | 18R2 | het | het |  |  |
| YYy310-10-18WR | XIV | S70 | 18W1 | het | Y | 540177 | 540710 |
| UV |  |  |  | Y | het | 541005 | 542559 |
|  |  |  | 18W2 | het | Y | 537970 | 538761 |
|  |  |  |  | Y | het | 539355 | 539640 |
|  |  |  | 18R1 | het | het |  |  |
|  |  |  | 18R2 | het | het |  |  |
| YYy310-10-18WR | XV | S71 | 18W1 | het | Y | 105667 | 108932 |
| UV |  |  |  | Y | het | 139346 | 140733 |
|  |  |  | 18W2 | Y | het | 141843 | 143584 |
|  |  |  | 18R1 | het | Y | 80436 | 86804 |
|  |  |  |  | Y | het | 108932 | 111280 |
|  |  |  | 18R2 | het | Y | 105667 | 108932 |
|  |  |  |  | Y | het | 108932 | 111280 |
| YYy310-10-18WR | XV | S72 | 18W1 | het | het |  |  |
| UV |  |  | 18W2 | het | Y | 252032 | 253952 |
|  |  |  |  | Y | het | 253952 | 254189 |
|  |  |  |  | het | Y | 254189 | 255004 |
|  |  |  |  | Y | het | 255254 | 259077 |
|  |  |  | 18R1 | het | het |  |  |
|  |  |  | 18R2 | het | het |  |  |
| YYy310-10-18WR | XV | S73 | 18W1 | het | S | 288101 | 288801 |
| UV |  |  |  | S | het | 288906 | 289639 |
|  |  |  | 18W2 | het | het |  |  |
|  |  |  | 18R1 | het | het |  |  |
|  |  |  | 18R2 | het | het |  |  |
| YYy310-10-23WR | I | S74 | 23W1 | S | het | 92538 | 93104 |
| UV |  |  |  | het | S | 95179 | 96036 |
|  |  |  |  | S | het | 99216 | 99918 |
|  |  |  | 23W2 | S | het | 92538 | 93104 |
|  |  |  |  | het | S | 95179 | 96036 |
|  |  |  |  | S | het | 98625 | 98874 |
|  |  |  | 23R1 | Y | het | 88734 | 90376 |
|  |  |  |  | het | S | 90970 | 92053 |
|  |  |  |  | S | het | 92521 | 92538 |
|  |  |  | 23R2 | Y | het | 92538 | 93104 |
| YYy310-10-23WR | IV | S75 | 23W1 | het | S | 1468302 | 1468543 |
| UV |  |  |  | S | het | 1469944 | 1471442 |
|  |  |  | 23W2 | het | S | 1468302 | 1468543 |
|  |  |  |  | S | het | 1468701 | 1469252 |
|  |  |  | 23R1 | het | S | 1468302 | 1468543 |
|  |  |  |  | S | het | 1468543 | 1468701 |
|  |  |  | 23R2 | het | S | 1468302 | 1468543 |
|  |  |  |  | S | het | 1469944 | 1471442 |
| YYy310-10-23WR | V | S76 | 23W1 | Y | het | 75719 | 76261 |
| UV |  |  |  | het | Y | 78080 | 80069 |
|  |  |  |  | Y | het | 80291 | 81264 |
|  |  |  | 23W2 | Y | het | 80291 | 81264 |
|  |  |  | 23R1 | S | het | 75719 | 76261 |
|  |  |  |  | het | Y | 78080 | 80069 |
|  |  |  |  | Y | het | 82770 | 84997 |
|  |  |  | 23R2 | S | het | 72490 | 75719 |
| YYy310-10-23WR | VIII | S77 | 23W1 | het | het |  |  |
| UV |  |  | 23W2 | het | het |  |  |
|  |  |  | 23R1 | het | Y | 95306 | 96110 |
|  |  |  |  | Y | het | 96638 | 97583 |
|  |  |  | 23R2 | het | S | 95306 | 96110 |
|  |  |  |  | S | HET | 96110 | 96638 |
|  |  |  |  | HET | het | 96638 | 97583 |
| YYy310-10-23WR | XII | S78 | 23W1 | het | Y | 234494 | 235321 |
| UV |  |  |  | Y | HET | 240616 | 241709 |
|  |  |  |  | HET | Y | 243415 | 243631 |
|  |  |  | 23W2 | het | Y | 235686 | 236112 |
|  |  |  |  | Y | HET | 240616 | 241709 |
|  |  |  |  | HET | Y | 255725 | 256514 |
|  |  |  | 23R1 | het | S | 227845 | 228409 |
|  |  |  | 23R2 | het | HET | 228409 | 234166 |
|  |  |  |  | HET | S | 234494 | 235321 |
| YYy310-10-23WR | XII | S79 | 23W1 | Y | het | 400760 | 401521 |
| UV |  |  |  | het | Y | 403077 | 407394 |
|  |  |  |  | Y | het | 407394 | 407628 |
|  |  |  |  | het | Y | 408418 | 412592 |
|  |  |  | 23W2 | Y | het | 400760 | 401521 |
|  |  |  |  | het | Y | 403077 | 407394 |
|  |  |  |  | Y | het | 407394 | 407628 |
|  |  |  |  | het | Y | 408418 | 412592 |
|  |  |  | 23R1 | S | S |  |  |
|  |  |  | 23R1 | S | S |  |  |
| YYy310-10-23WR | XVI | S80 | 23W1 | het | S | 676134 | 677039 |
| UV |  |  |  | S | het | 677039 | 677701 |
|  |  |  | 23W2 | het | S | 676134 | 677039 |
|  |  |  |  | S | het | 677039 | 677701 |
|  |  |  | 23R1 | het | het |  |  |
|  |  |  | 23R2 | het | het |  |  |
| YYy310-1WR | V | S81 | 1W1 | Y | het | 78080 | 80069 |
| spontaneous |  |  |  | het | S | 80069 | 80291 |
|  |  |  |  | S | het | 80291 | 81264 |
|  |  |  | 1W2 | Y | het | 78080 | 80069 |
|  |  |  |  | het | S | 80069 | 80291 |
|  |  |  |  | S | Y | 82770 | 84997 |
|  |  |  |  | Y | het | 84997 | 85566 |
|  |  |  | 1R1 | S | het | 78080 | 80069 |
|  |  |  | 1R2 | S | Y | 78080 | 80069 |
|  |  |  |  | Y | het | 80291 | 81264 |
| YYy310-4WR | V | S82 | 4W1 | Y | HET | 90215 | 91703 |
| spontaneous |  |  |  | HET | Y | 95906 | 97221 |
|  |  |  |  | Y | het | 97792 | 98369 |
|  |  |  |  | het | Y | 98369 | 98736 |
|  |  |  |  | Y | HET | 100439 | 101353 |
|  |  |  |  | HET | S | 104257 | 104636 |
|  |  |  |  | S | het | 105481 | 105830 |
|  |  |  | 4W2 | Y | HET | 90215 | 91703 |
|  |  |  |  | HET | Y | 91739 | 93200 |
|  |  |  |  | Y | het | 97792 | 98369 |
|  |  |  |  | het | HET | 100201 | 100439 |
|  |  |  |  | HET | S | 100439 | 101353 |
|  |  |  |  | S | het | 105481 | 105830 |
|  |  |  | 4R1 | S | het | 104257 | 104636 |
|  |  |  | 4R2 | S | HET | 90215 | 91703 |
|  |  |  |  | HET | S | 95906 | 97221 |
|  |  |  |  | S | het | 104636 | 105299 |
| YYy310-6WR | V | S83 | 6W1 | Y | het | 39153 | 41126 |
| spontaneous |  |  | 6W2 | Y | HET | 34799 | 37492 |
|  |  |  |  | HET | het | 39153 | 41126 |
|  |  |  | 6R1 | S | het | 34799 | 37492 |
|  |  |  | 6R2 | S | het | 33770 | 34235 |
|  |  |  |  | het | S | 34799 | 37492 |
|  |  |  |  | S | het | 39153 | 41126 |
| YYy310-9WR | V | S84 | 9W1 | Y | HET | 94077 | 94339 |
| spontaneous |  |  |  | HET | het | 100439 | 101353 |
|  |  |  |  | het | S | 107912 | 109029 |
|  |  |  |  | S | het | 109050 | 111741 |
|  |  |  | 9W2 | Y | HET | 90215 | 91703 |
|  |  |  |  | HET | S | 91739 | 93200 |
|  |  |  |  | S | HET | 93407 | 94077 |
|  |  |  |  | HET | S | 96550 | 97221 |
|  |  |  |  | S | HET | 98369 | 98736 |
|  |  |  |  | HET | S | 100439 | 101353 |
|  |  |  |  | S | het | 107912 | 109029 |
|  |  |  | 9R1 | S | het | 118782 | 119505 |
|  |  |  | 9R2 | S | het | 118605 | 118782 |
| YYy310-38WR | V | S85 | 38W1 | Y | het | 38068 | 41126 |
| spontaneous |  |  | 38W2 | Y | het | 34481 | 37492 |
|  |  |  |  | het | S | 41533 | 44506 |
|  |  |  |  | S | Y | 44506 | 45284 |
|  |  |  |  | Y | het | 45587 | 45951 |
|  |  |  | 38R1 | S | het | 45587 | 45951 |
|  |  |  | 38R2 | S | het | 46371 | 48837 |
| YYy311-10WR | IV | S86 | 10W1 | het | Y | 968316 | 968451 |
| spontaneous |  |  |  | Y | het | 971058 | 971688 |
|  |  |  |  | het | Y | 973167 | 973779 |
|  |  |  |  | Y | het | 976159 | 977291 |
|  |  |  |  | het | Y | 977470 | 977752 |
|  |  |  | 10W2 | het | Y | 969369 | 969944 |
|  |  |  |  | Y | het | 970646 | 971058 |
|  |  |  |  | het | Y | 971889 | 972525 |
|  |  |  | 10R1 | het | S | 973167 | 973779 |
|  |  |  | 10R2 | het | S | 968316 | 968451 |
|  |  |  |  | S | het | 971058 | 971688 |
|  |  |  |  | het | S | 976159 | 977291 |
| YYy311-21WR | IV | S87 | 21W1 | het | Y | 973959 | 974614 |
| spontaneous |  |  | 21W2 | het | Y | 973959 | 974614 |
|  |  |  |  | Y | het | 993347 | 993964 |
|  |  |  |  | het | Y | 996000 | 996527 |
|  |  |  | 21R1 | het | Y | 978621 | 979477 |
|  |  |  |  | Y | het | 996527 | 996677 |
|  |  |  |  | het | S | 1017595 | 1022402 |
|  |  |  | 21R2 | het | S | 977470 | 977752 |
|  |  |  |  | S | HET | 978621 | 979477 |
|  |  |  |  | HET | het | 980838 | 992782 |
|  |  |  |  | het | Y | 993347 | 993964 |
|  |  |  |  | Y | het | 995883 | 996527 |
|  |  |  |  | het | S | 1017595 | 1022402 |
| YYy311-25WR | IV | S88 | 25W1 | het | Y | 967772 | 968451 |
| spontaneous |  |  | 25W2 | het | Y | 978763 | 979777 |
|  |  |  | 25R1 | het | Y | 950685 | 950745 |
|  |  |  |  | Y | het | 978763 | 979777 |
|  |  |  |  | het | Y | 980838 | 992782 |
|  |  |  |  | Y | S | 993964 | 995822 |
|  |  |  |  | S | Y | 997558 | 997989 |
|  |  |  |  | Y | S | 999715 | 1002104 |
|  |  |  |  | S | Y | 1004465 | 1005641 |
|  |  |  |  | Y | S | 1011955 | 1013221 |
|  |  |  |  | S | Y | 1036164 | 1036844 |
|  |  |  |  | Y | S | 1037332 | 1039370 |
|  |  |  | 25R2 | het | Y | 950685 | 950745 |
|  |  |  |  | Y | het | 978763 | 979777 |
|  |  |  |  | het | Y | 980838 | 992782 |
|  |  |  |  | Y | S | 993964 | 995822 |
|  |  |  |  | S | Y | 997558 | 997989 |
|  |  |  |  | Y | S | 999715 | 1002104 |
|  |  |  |  | S | het | 1004465 | 1005641 |
|  |  |  |  | het | S | 1011955 | 1013221 |
|  |  |  |  | S | het | 1036164 | 1036844 |
|  |  |  |  | het | S | 1037332 | 1039370 |
| YYy311-27WR | IV | S89 | 27W1 | het | Y | 975687 | 976159 |
| spontaneous |  |  | 27W2 | het | Y | 974716 | 974975 |
|  |  |  | 27R1 | het | Y | 968451 | 969369 |
|  |  |  |  | Y | het | 995622 | 995822 |
|  |  |  |  | het | Y | 995883 | 996000 |
|  |  |  |  | Y | het | 1000365 | 1001602 |
|  |  |  |  | het | S | 1004465 | 1005641 |
|  |  |  | 27R2 | het | Y | 968451 | 969369 |
|  |  |  |  | Y | het | 993964 | 994215 |
|  |  |  |  | het | S | 1004465 | 1005641 |
| YYy311-30WR | IV | S90 | 30W1 | het | Y | 974674 | 974716 |
| spontaneous |  |  | 30W2 | het | Y | 974674 | 974716 |
|  |  |  |  | Y | HET | 980838 | 992782 |
|  |  |  |  | HET | Y | 1000365 | 1001602 |
|  |  |  | 30R1 | het | Y | 980838 | 992782 |
|  |  |  |  | Y | het | 992897 | 993113 |
|  |  |  |  | het | Y | 996000 | 996527 |
|  |  |  |  | Y | het | 997989 | 998469 |
|  |  |  |  | het | S | 1000365 | 1001602 |
|  |  |  | 30R2 | het | Y | 980838 | 992782 |
|  |  |  |  | Y | het | 992897 | 993113 |
|  |  |  |  | het | Y | 994215 | 994598 |
|  |  |  |  | Y | het | 994598 | 995622 |
|  |  |  |  | het | Y | 996000 | 996527 |
|  |  |  |  | Y | het | 997989 | 998469 |
|  |  |  |  | het | S | 1000365 | 1001602 |
| YYy311-32WR | IV | S91 | 32W1 | het | Y | 973959 | 974614 |
| spontaneous |  |  | 32W2 | het | Y | 964391 | 965936 |
|  |  |  | 32R1 | het | Y | 953272 | 955278 |
|  |  |  |  | Y | S | 993964 | 994598 |
|  |  |  | 32R2 | het | Y | 968451 | 969944 |
|  |  |  |  | Y | HET | 978358 | 978583 |
|  |  |  |  | HET | Y | 980838 | 992782 |
|  |  |  |  | Y | S | 993964 | 994598 |

**Table S3.** Classification of UV-induced mitotic recombination events in YYy310 (derivatives YYy310-9 and YYy310-10); Figs. S3-S80), spontaneous events in YYy310 (Figs. S81-S85), and spontaneous crossovers on chromosome IV in YYy311 (Figs. 86-91). Details of the positions of the transitions are given in Table S2. The table headings are described in the table legend.

| Sector | Chromosome | Type of break | Break 1 |  |  |  | Break 2 |  |  |  | Figure |
| --- | --- | --- | --- | --- | --- | --- | --- | --- | --- | --- | --- |
|  |  |  | NCO or | Chromatids | Homolog broken | Class | NCO or | Chromatids | Homolog broken | Class |  |
| YYy310-9-5WR | II | DSCB | NCO | 1 & 4 | S | 1 | NCO | 2 & 3 | S | Complex | S3 |
|  | III | SCB | NCO | 1 & 3 | Y | 2 |  |  |  |  | S4 |
|  | IV-1 | SCB | NCO | 1 & 4 | Y | 3 |  |  |  |  | S5 |
|  | IV-2 | SCB | NCO | 2 & 3 | S | 1 |  |  |  |  | S6 |
|  | IV-3 | DSCB | NCO | 1 & 4 | S | 1 | NCO | 2 & 3 | S | Complex | S7 |
|  | V | DSCB | CO | 1 & 4 | Y | Complex | NCO | 2 & 3 | Y | Complex | S8 |
|  | VI | DSCB | NCO | 1 & 3 | Y | 3 | NCO | 2 & 4 | Y | 1 | S9 |
|  | VII | SCB | NCO | 2 & 3 | Y | 1 |  |  |  |  | S10 |
|  | VIII | SCB | CO | 2 & 4 | S | 4 |  |  |  |  | S11 |
|  | X | DSCB | NCO | 1 & 3 | S | 1 | NCO | 2 & 4 | S | 5 | S12 |
|  | XI | DSCB | NCO | 1 & 3 | S | Complex | NCO | 2 & 4 | S | Complex | S13 |
|  | XII | DSCB | NCO | 1 & 3 | Y | 5 | NCO | 2 & 4 | Y | 1 | S14 |
|  | XV | DSCB | NCO | 1 & 3 | Y | 1 | NCO | 2 & 4 | Y | 1 | S15 |
|  | XVI | DSCB | CO | 1 & 3 | Y | 6 | NCO | 2 & 4 | Y | Complex | S16 |
| YYy310-9-12WR | II | DSCB | NCO | 1 & 3 | Y | 5 | NCO | 2 & 4 | Y | 1 | S17 |
|  | IV-1 | SCB | NCO | 1 & 3 | S | 1 |  |  |  |  | S18 |
|  | IV-2 | SCB | NCO | 2 & 3 | S | 2 |  |  |  |  | S19 |
|  | IV-3 | SCB | NCO | 2 & 3 | Y | 1 |  |  |  |  | S20 |
|  | V | SCB | CO | 1 & 4 | S | 4 |  |  |  |  | S21 |
|  | VII | SCB | NCO | 1 & 3 | S | 2 |  |  |  |  | S22 |
|  | VIII-1 | SCB | NCO | 1 & 3 | S | 2 |  |  |  |  | S23 |
|  | VIII-2 | DSCB | CO | 1 & 3 | Y | 7 | CO | 2 & 4 | Y | Complex | S24 |
|  | X | SCB | NCO | 1 & 3 | S | 2 |  |  |  |  | S25 |
|  | XI-1 | DSCB | NCO | 1 & 3 | Y | Complex | CO | 2 & 4 | Y | 4 | S26 |
|  | XI-2 | SCB | NCO | 1 & 3 | S | 2 |  |  |  |  | S27 |
|  | XII-1 | DSCB | CO | 1 & 3 | Y | 4 | NCO | 2 & 4 | Y | Complex | S28 |
|  | XII-2 | SCB | NCO | 2 & 3 | S | 8 |  |  |  |  | S29 |
|  | XII-3 | SCB | NCO | 1 & 4 | Y | 3 |  |  |  |  | S30 |
|  | XII-4 | DSCB | NCO | 1 & 3 | Y | 2 | NCO | 2 & 4 | Y | 3 | S31 |
|  | XIII-1 | SCB | NCO | 1 & 3 | S | 1 |  |  |  |  | S32 |
|  | XIII-2 | SCB | NCO | 2 & 3 | S | 2 |  |  |  |  | S33 |
|  | XVI-1 | DSCB | NCO | 1 & 3 | S | 5 | NCO | 2 & 4 | S | Complex | S34 |
|  | XVI-2 | SCB | NCO | 2 & 4 | Y | 2 |  |  |  |  | S35 |
| YYy310-9-17WR | I | SCB | NCO | 1 & 3 | S | 1 |  |  |  |  | S36 |
|  | II-1 | SCB | CO | 1 & 4 | S | 4 |  |  |  |  | S37 |
|  | II-2 | DSCB | NCO | 2 & 3 | S | Complex | NCO | 3 & 4 | Y | Complex | S38 |
|  | II-3 | SCB | NCO | 1 & 3 | Y | 1 |  |  |  |  | S39 |
|  | IV-1 | SCB | NCO | 1 & 3 | S | 1 |  |  |  |  | S40 |
|  | IV-2 | DSCB | CO | 1 & 3 | S | Complex | NCO | 2 & 4 | S | Complex | S41 |
|  | IV-3 | SCB | NCO | 1 & 3 | S | 3 |  |  |  |  | S42 |
|  | V-1 | DSCB | CO | 1 & 4 | S | Complex | NCO | 2 & 3 | S | Complex | S43 |
|  | V-2 | DSCB | NCO | 1 & 3 | S | Complex | NCO | 2 & 4 | S | Complex | S44 |
|  | IX | DSCB | NCO | 1 & 4 | S | 1 | CO | 2 & 3 | S | Complex | S45 |
|  | XV | DSCB | NCO | 1 & 3 | S | 2 | NCO | 2 & 4 | S | 2 | S46 |
| YYy310-10-14WR | II | DSCB | NCO | 1 & 3 | Y | 2 | NCO | 2 & 4 | Y | 2 | S47 |
|  | V | SCB | CO | 1 & 4 | S | Complex |  |  |  |  | S48 |
|  | IX | SCB | NCO | 1 & 3 | S | 2 |  |  |  |  | S49 |
|  | XI | SCB | NCO | 2 & 3 | S | 1 |  |  |  |  | S50 |
|  | XII | SCB | NCO | 1 & 4 | Y | 1 |  |  |  |  | S51 |
|  | XIII-1 | SCB | NCO | 1 & 4 | Y | 3 |  |  |  |  | S52 |
|  | XIII-2 | DSCB | NCO | 1 & 3 | S | Complex | NCO | 2 & 4 | S | Complex | S53 |
|  | XV | DSCB | NCO | 1 & 3 | S | 1 | NCO | 2 & 4 | S | 2 | S54 |
|  | XVI | DSCB | NCO | 1 & 3 | Y | 3 | NCO | 2 & 4 | Y | 3 | S55 |
| YYy310-10-17WR | IV-1 | DSCB | NCO | 1 & 3 | S | Complex | NCO | 2 & 4 | S | Complex | S56 |
|  | IV-2 | SCB | NCO | 2 & 3 | S | 2 |  |  |  |  | S57 |
|  | IV-3 | DSCB | CO | 1 & 4 | Y | 7 | NCO | 2 & 3 | Y | 1 | S58 |
|  | V | DSCB | CO | 1 & 4 | S | Complex | NCO | 2 & 3 | S | Complex | S59 |
|  | VIII | 2SCBs | NCO | 1 & 4 | S | 1 | NCO | 2 & 3 | Y | 1 | S60 |
|  | IX | DSCB | CO | 1 & 4 | Y | Complex | CO | 2 & 3 | S | 6 | S61 |
|  | X | DSCB | NCO | 1 & 4 | S | Complex | CO | 2 & 3 | S | Complex | S62 |
|  | XIII | DSCB | NCO | 1 & 3 | S | 1 | NCO | 2 & 4 | S | 2 | S63 |
| YYy310-10-18WR | II | DSCB | NCO | 1 & 3 | S | 1 | NCO | 2 & 4 | S | Complex | S64 |
|  | V | DSCB | CO | 1 & 4 | S | Complex | NCO | 2 & 3 | S | Complex | S65 |
|  | VII | DSCB | NCO | 2 & 3 | Y | Complex | NCO | 1 & 4 | Y | Complex | S66 |
|  | X | SCB | NCO | 2 & 3 | Y | 8 |  |  |  |  | S67 |
|  | XII | SCB | NCO | 2 & 3 | Y | 3 |  |  |  |  | S68 |
|  | XIV-1 | SCB | NCO | 2 & 3 | Y | 3 |  |  |  |  | S69 |
|  | XIV-2 | SCB | NCO | 1 & 3 | S | Complex |  |  |  |  | S70 |
|  | XV-1 | See table legend. |  |  |  |  |  |  |  |  | S71 |
|  | XV-2 | SCB | NCO | 1 & 3 | S | Complex |  |  |  |  | S72 |
|  | XV-3 | SCB | NCO | 1 & 3 | Y | 1 |  |  |  |  | S73 |
| YYy310-10-23WR | I | DSCB | NCO | 1 & 4 | Y | Complex | CO | 2 & 3 | Y | Complex | S74 |
|  | IV | DSCB | NCO | 1 & 3 | Y | 3 | NCO | 2 & 4 | Y | 3 | S75 |
|  | V | DSCB | CO | 1 & 4 | S | Complex | NCO | 2 & 3 | S | Complex | S76 |
|  | VIII | SCB | NCO | 2 & 4 | S | Complex |  |  |  |  | S77 |
|  | XII-1 | DSCB | CO | 1 & 4 | Y | Complex | NCO | 2 & 3 | Y | Complex | S78 |
|  | XII-2 | SCB | NCO | 1 & 3 | Y | Complex |  |  |  |  | S79 |
|  | XVI | SCB | NCO | 2 & 3 | Y | 2 |  |  |  |  | S80 |
| YYy310-1WR | V | DSCB | CO | 1 & 4 | Y | Complex | NCO | 2 & 3 | Y | Complex | S81 |
| YYy310-4WR | V | DSCB | CO | 1 & 4 | Y | Complex | NCO | 2 & 3 | Y | Complex | S82 |
| YYy310-6WR | V | DSCB | CO | 1 & 4 | S | Complex | NCO | 2 & 3 | S | Complex | S83 |
| YYy310-9WR | V | DSCB | CO | 1 & 4 | Y | Complex | NCO | 2 & 3 | Y | Complex | S84 |
| YYy310-38WR | V | DSCB | CO | 1 & 4 | Y | Complex | NCO | 2 & 3 | Y | 1 | S85 |
| YYy311-10WR | IV | SCB | CO | 1 & 4 | S | Complex |  |  |  |  | S86 |
| YYy311-21WR | IV | DSCB | CO | 1 & 4 | S | Complex | NCO | 2 & 3 | S | Complex | S87 |
| YYy311-25WR | IV | DSCB | CO | 1 & 4 | S | Complex | NCO | 2 & 3 | S | Complex | S88 |
| YYy311-27WR | IV | DSCB | CO | 1 & 4 | S | Complex | NCO | 2 & 3 | S | Complex | S89 |
| YYy311-30WR | IV | DSCB | CO | 1 & 4 | S | Complex | NCO | 2 & 3 | S | Complex | S90 |
| YYy311-32WR | IV | DSCB | CO | 1 & 4 | S | Complex | NCO | 2 & 3 | S | Complex | S91 |

**Table S4.** Summary of complex patterns of recombination.

| Supp. Fig. # | CO or NCO | Observations | Interpretations |
| --- | --- | --- | --- |
| S3 | NCO | Simple heteroduplex. DSB on blue chromatid. | SDSA |
|  | CO | Long heteroduplex with regions of homoduplex. DSB on red chromatid. | Standard DSBR event except Mlh1-independent MMR. Two independent DSBs. |
| S7 | NCO1 | Large homoduplex region separating DSB site from heteroduplex tract (chromatids 2 and 3). | Repair of double-stranded DNA gap |
|  | NCO2 | Simple heteroduplex on chromatid 1 | SDSA |
| S8 | CO | Symmetric heteroduplexes | Branch migration following formation of double Holliday junction (dHJ) |
|  | NCO | Strand switches of heteroduplexes on NCO chromatid, region of conversion at junction of heteroduplex | dHJ dissolution and region of Mlh1-independent MMR |
| S13 | NCO1 | Heteroduplex spanning DSB site in chromatid 1 | Extension of broken end by interaction with sister chromatid |
|  | NCO2 | Heteroduplexes in trans on chromatid 2 separated by homoduplex | Dissolution of dHJ followed by Mlh1-independent MMR |
| S16 | CO | Heteroduplexes in *trans* flanking DSB site | Standard CO according to DSBR model |
|  | NCO | Heteroduplex spanning DSB site on NCO chromatid 4 | Extension of broken end by interaction with sister chromatid |
| S17 | NCO1 | Heteroduplex with interspersed conversion and restoration tracts (chromatid 3) | SDSA with Mlh1-independent MMR |
|  | NCO2 | Simple heteroduplex (chromatid 4) | SDSA |
| S24 | CO1 | CO between chromatids 1 and 3 with no heteroduplex | Standard DSBR model with limited strand invasion and limited DNA synthesis |
|  | CO2 | CO between chromatids 2 and 4 with restoration tract between *trans* heteroduplexes | Standard DSBR model with tract of Mlh1-independent restoration repair |
| S26 | NCO | Symmetric heteroduplexes; regions of homoduplex within heteroduplex; strand switch within heteroduplex | Branch migration; Mlh1-independent MMR; DSB between regions of strand switch, followed by dissolution |
|  | CO | CO between chromatids 2 and 4 with heteroduplex on only one chromatid | Standard CO according to DSBR with one heteroduplex occurring in region without SNP |
| S28 | NCO | Long conversion tract spanning putative DSB site | Repair of double-stranded DNA gap |
|  | CO | Heteroduplex tract on only one side of DSB site | Standard CO according to DSBR with one heteroduplex occurring in region without SNP |
| S34 | NCO1 | Long conversion-tract on chromatid 1 | Repair of double-stranded DNA gap |
|  | NCO2 | Heteroduplexes in trans on chromatid 2 separated by homoduplex | Dissolution of dHJ associated with restoration repair |
| S38 | NCO1 | Conversion tract adjacent to DSB1 site on chromatid 3 | SDSA, Mlh1-independent MMR |
|  | NCO2 | Heteroduplex adjacent to DSB1 site on chromatid 4 | SDSA |
|  | NCO3 | Heteroduplex adjacent to DSB2 site on chromatid 2 | SDSA |
| S41 | NCO | Symmetric heteroduplexes on chromatids 2 and 4; homoduplex regions within heteroduplex | Branch migration; Mlh1-independent MMR |
|  | CO | Crossover between chromatids 1 and 3 associated with long conversion tract | Repair of double-stranded DNA gap |
| S43 | CO | Restoration repair on one side of CO chromatid 4 and conversion repair on the other | Branch migration; Mlh1-independent MMR |
|  | NCO | Long heteroduplex region with short restoration tract | SDSA with Mlh1-independent MMR |
| S44 | NCO1 | Chromatid 1 has heteroduplex region with long restoration tract in the middle | Template switch during replication to sister strand; SDSA |
|  | NCO2 | Chromatid 2 has mixture of homoduplex conversion and restoration tracts | Mlh1-independent patchy MMR |
| S45 | NCO | Simple heteroduplex (chromatid 1) | SDSA |
|  | CO | Regions of heteroduplex with switched strands on chromatid 3; symmetric heteroduplexes | Independent invasion of two broken ends; branch migration |
| S48 | CO | Regions of heteroduplex interspersed with homoduplex regions | Mlh1-independent patchy MMR |
| S53 | NCO1 | Conversion tract with no heteroduplex in chromatid 1 | Mlh1-independent MMR |
|  | NCO2 | Conversion and restoration tracts in chromatid 2 | Mlh1-independent patchy MMR |
| S56 | NCO1 | Conversion tract with no heteroduplex in chromatid 1 | Mlh1-independent MMR |
|  | NCO2 | Conversion and restoration tracts in chromatid 2 | Mlh1-independent patchy MMR |
| S59 | CO | Long conversion tract between heteroduplex and putative DSB site on chromatid 4 | Repair of double-stranded DNA gap |
|  | NCO | Regions of conversion/restoration and heteroduplex on same side of putative DSB site in chromatids 3 and 4 | Branch migration; NCO resolution of dHJ; Mlh1-independent patchy repair |
| S61 | CO1 | *Trans* heteroduplexes on chromatids 2 and 3 following DSB on red chromatid | Crossover by standard DSBR model |
|  | CO2 | DSB on blue chromatid, large conversion tract | Repair of double-stranded DNA gap; followed by cleavage of dHJ to yield a CO |
| S62 | NCO | Conversion tract spanning putative DSB site in chromatid 1 | Repair of double-stranded DNA gap; dissolution |
|  | CO | *Trans* heteroduplexes on chromatids 2 and 3; interspersed conversion and restoration tracts | Mlh1-independent MMR |
| S64 | NCO1 | Simple heteroduplex (chromatid 1) | SDSA |
|  | NCO2 | Heteroduplexes on same side of DSB in chromatids 2 and 4 | Branch migration; NCO mode of dHJ resolution |
| S65 | CO | Heteroduplex on same side of DSB site in chromatids 1 and 4; heteroduplex spans DSB site in chromatid 4; tracts of restoration within heteroduplex in chromatid 2 | Extension of broken end by interaction with sister chromatid; Mlh1-independent patchy repair; CO resolution of dHJ |
|  | NCO | Heteroduplex spans DSB site in chromatid 2; tracts of conversion and restoration interspersed with heteroduplex regions | Extension of broken end by interaction with sister chromatid; Mlh1-independent patchy repair; SDSA |
| S66 | NCO1 | Heteroduplex on chromatid 3 separated from putative DSB site by long region of homoduplex | Extension of broken end by interaction with sister chromatid, followed by SDSA |
|  | NCO2 | Heteroduplexes on both chromatids 1 and 4; chromatid 1 has trans heteroduplexes flanking the DSB site in addition to regions of homoduplex within the heteroduplex | Invasion of right end, followed by cleavage of junction and invasion of left end; branch migration; Mlh1-independent patchy repair |
| S70 | NCO | *Trans* heteroduplexes separated by restoration tract on chromatid 1 | Formation of dHJ event; dissolution followed by Mlh1-independent repair |
| S72 | NCO | Heteroduplex tract with an internal restoration tract | Mlh1-independent MMR, SDSA |
| S74 | NCO | Region of restoration repair separating DSB site from heteroduplex on chromatid 4 | Mlh1-independent MMR, SDSA |
|  | CO | Long regions of restoration and conversion repair separating DSB site from heteroduplex tract | Mlh1-independent MMR or template switching |
| S76 | CO | Conversion event at the end of heteroduplex tract | Mlh1-independent MMR, resolution of dHJ in CO mode |
|  | NCO | Displacement of heteroduplex tract from DSB site | Interaction of broken end with sister chromatid or long restoration tract by Mlh1-independent MMR |
| S77 | NCO | Symmetric heteroduplex; conversion tract at end of heteroduplex tract | Branch migration; Mlh1-independent MMR; NCO processing of dHJ |
| S78 | NCO | Regions of homoduplex separating heteroduplex region from DSB site | Mlh1-independent MMR or template switching; resolution of dHJ in NCO mode |
|  | CO | Region of conversion separating heteroduplex region from DSB site on chromatid 4 | Mlh1-independent MMR or gap repair; resolution of dHJ in CO mode |
| S79 |  | In regions centromere-proximal to event, both homologs in white sector derived from one parental homolog and both in red sector derived from the other; regions of conversion and restoration homoduplexes on chromatid 1 | Crossover on same chromosomes located centromere-proximal to the event in S79 (described in S78) produced centromere-proximal regions. Homoduplex regions produced by template switching or by Mlh1-independent MMR |
| S81 | CO | Long tracts of homoduplex on chromatids 1 and 4. | Extension of broken end by interaction with sister chromatid; Mlh1-independent restoration repair; resolution of dHJ in CO mode |
|  | NCO | Heteroduplex DNA on same side of DSB site in NCO chromatids | Branch migration followed by processing of dHJ in CO mode |
| S82 | CO | Heteroduplex with interspersed conversion and restoration tracts (chromatid 1); initiated | Mlh1-independent MMR; processing of dHJ in CO mode |
|  | NCO | Symmetric heteroduplex tracts; heteroduplex tract interspersed with homoduplex regions; strand switch on chromatid 3; initiated at DSB2 | Branch migration; Mlh1-independent MMR; dissolution of dHJ |
| S83 | CO | Strand switch of heteroduplex in CO chromatid 4 | Invasion of the right broken end, followed by processing of junction; extension of right end by sister-chromatid interaction; invasion of the left end and processing dHJ in crossover mode |
|  | NCO | DSB on red chromatid, but red chromatid acts as donor | Following strand invasion, junction cleaved before initiating DNA synthesis |
| S84 | CO | Long conversion tract spanning DSBs in both CO chromatids 1 and 4; heteroduplex in chromatid 1 interrupted by restoration tracts | Gap repair; Mlh1-independent MMR; resolution of dHJ in CO mode |
|  | NCO | NCO chromatid 3 has strand switch of heteroduplexes; conversion event within | Resolution of dHJ by dissolution; Mlh1-independent MMR |
| S85 | CO | Strand switch in CO chromatid 1; heteroduplexes in trans on chromatids 1 and 4; long conversion tract adjacent to heteroduplex region in chromatid 1 | Invasion of right end, followed by branch migration and cutting of junction. Extension of right end by interaction with sister chromatid; invasion of left end and resolution of dHJ in CO |
|  | NCO | Simple heteroduplex | SDSA |
| S86 | CO | Strand switches in heteroduplexes in both CO chromatids 1 and 4; both chromatids have homoduplex regions interspersed with heteroduplex | Invasion of right broken end, followed by branch migration; resolution of junctions in NCO mode, followed by left end invasion and branch migration; cleavage in CO mode and |
| S87 | CO | Large region of conversion spanning putative DSB site; heteroduplex regions on chromatids 1 and 4 in *trans*; conversion tract distal to heteroduplex tract on chromatid 4 | Repair of large gap associated with HS4 hotspot; processing dHJ in CO mode; Mlh1-independent MMR |
|  | NCO | Region of conversion spanning putative DSB site; conversion tract adjacent to heteroduplex | Repair of large gap; SDSA; Mlh1-independent MMR |
| S88 | CO | Region of conversion spanning putative DSB site; heteroduplex tracts in *trans* on chromatids 1 and 4; multiple homoduplex tracts within heteroduplex tract on chromatid | Repair of large gap; patchy Mlh1-independent MMR; resolution of dHJ in CO mode |
|  | NCO | Large conversion tract spanning putative DSB site; chromatid 3 has a pattern of heteroduplex and homoduplex tracts that is very similar to that of chromatid 4 | Repair of large gap; chromatid 4 CO chromatid used as template for part of repair; also, switch of repair templates to sister chromatid |
| S89 | CO | Very large conversion tract spanning putative DSB site; heteroduplex on chromatid 1 but not | Repair of large gap; resolution of dHJ in CO mode |
|  | NCO | Very large conversion tract spanning putative DSB site; restoration tract in middle of | Repair of large gap; resolution of dHJ in NCO mode; Mlh1-independent MMR |
| S90 | CO | Very large conversion tract spanning DSB site; no heteroduplexes observed | Repair of large double-stranded DNA gap, resolution of dHJ in CO mode |
|  | NCO | Heteroduplexes in both NCO chromatids 2 and 3 on same side of DSB site; homoduplex tracts in heteroduplex | Invasion of broken end, followed by branch migration; resolution of dHJ structure in NCO mode with regions of Mlh1-independent MMR |
| S91 | CO | Heteroduplexes on both chromatids 1 and 4 on same side of putative DSB site; large conversion tract flanking DSB site | Invasion of right broken end, followed by synthesis off of homolog; template switch to sister chromatid, and then re-invasion of sister chromatid; resolution of dHJ in CO mode |
|  | NCO | Large conversion tract spanning putative DSB | Repair of large gap; SDSA |

**Table S5.**
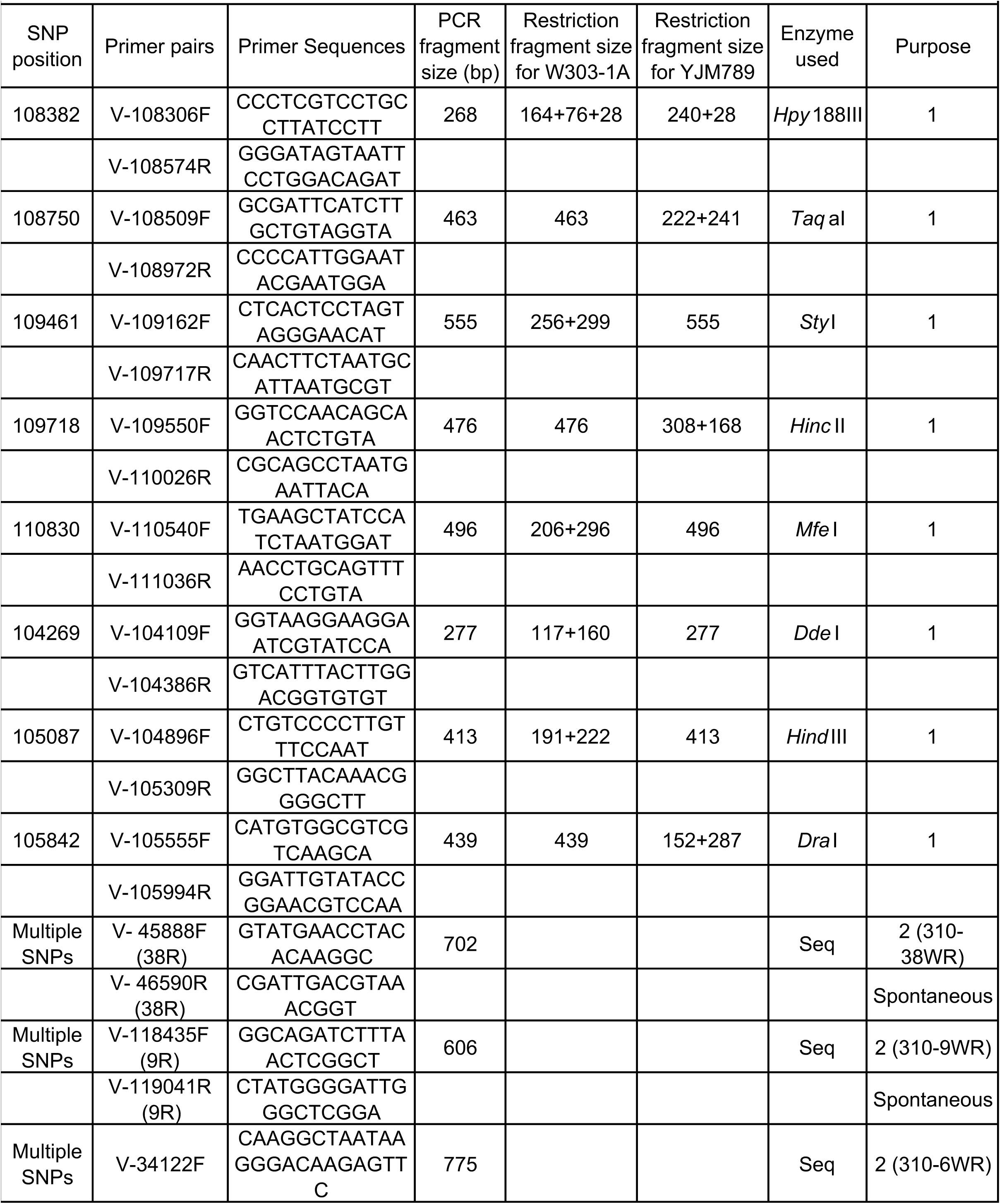
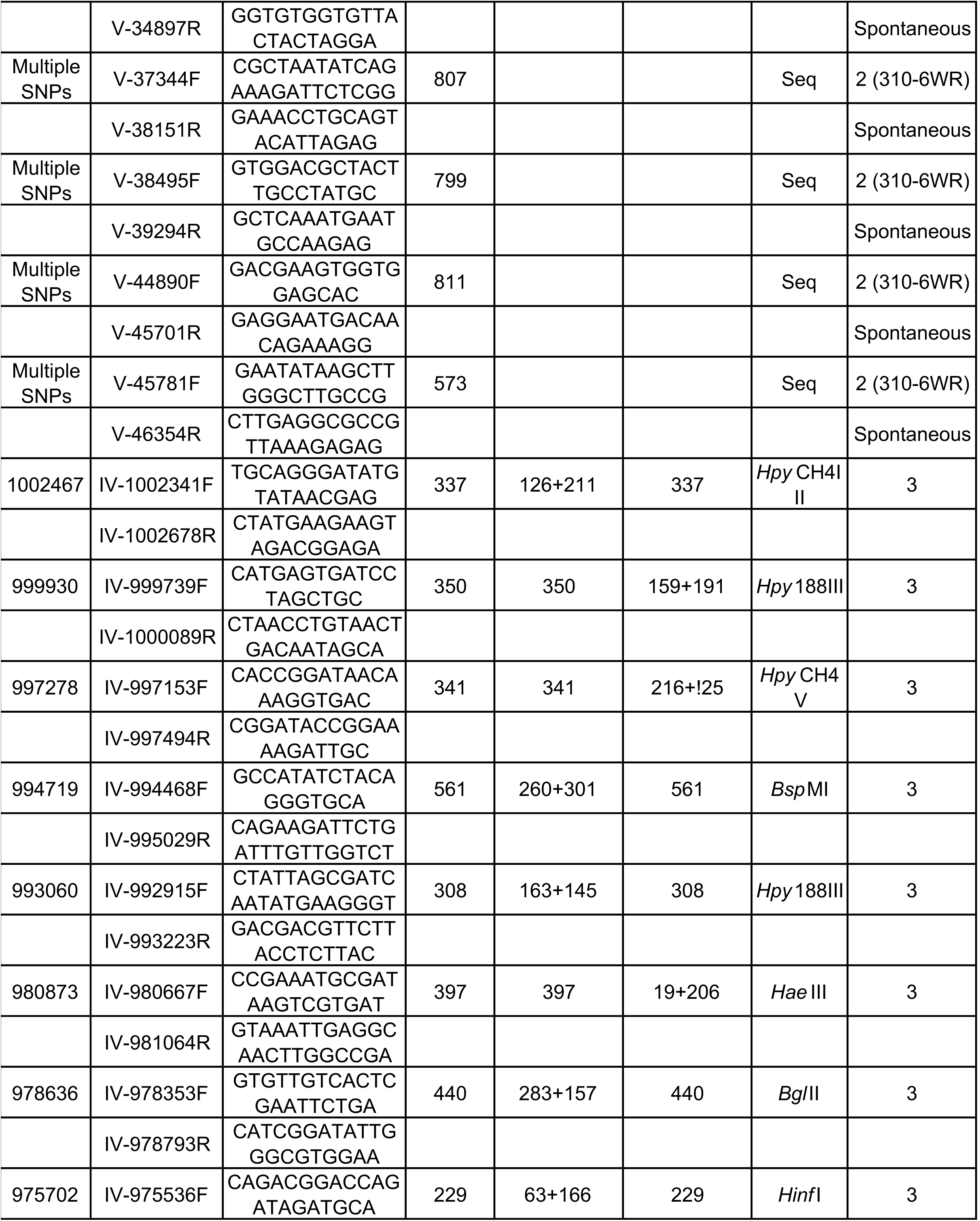
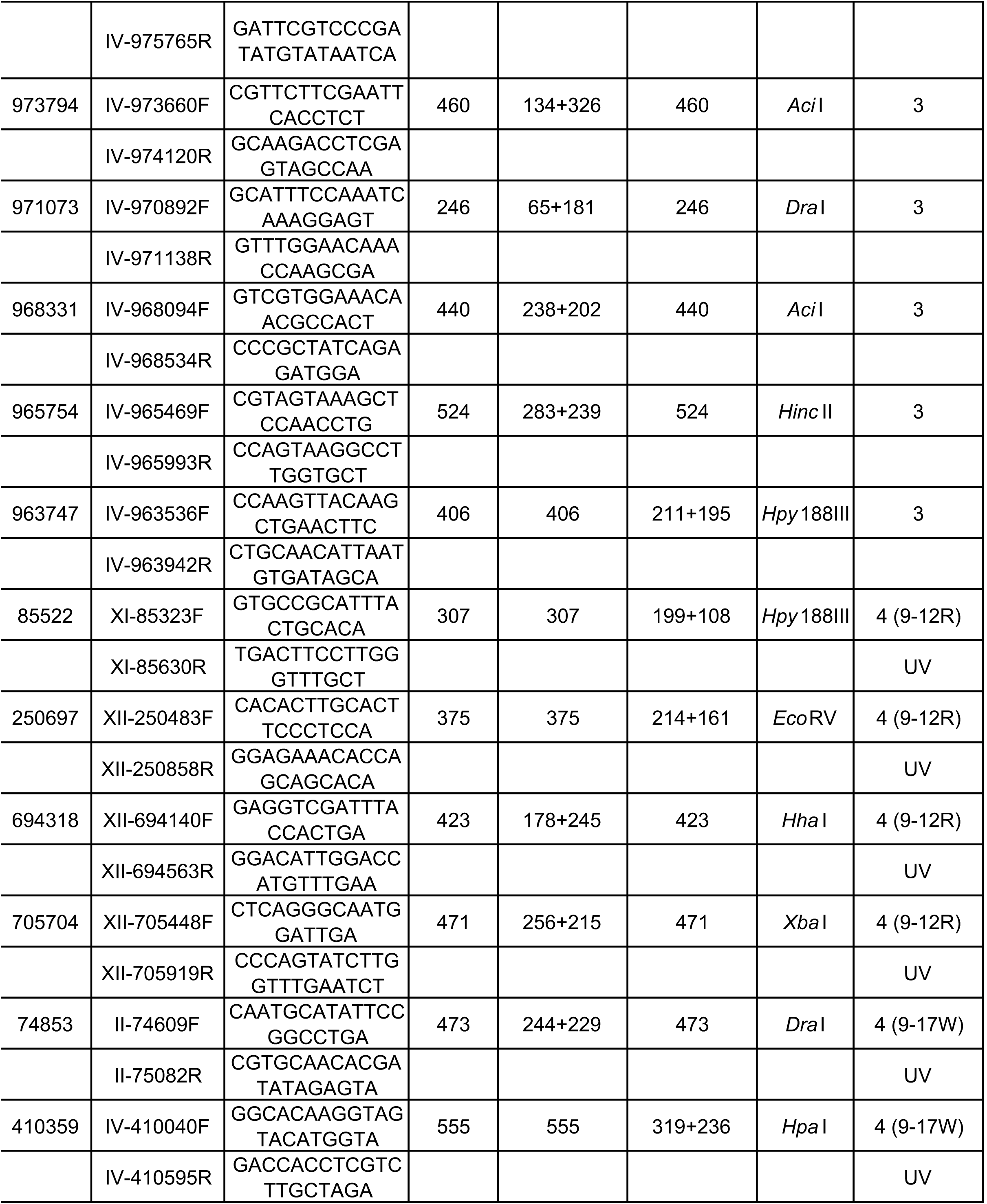
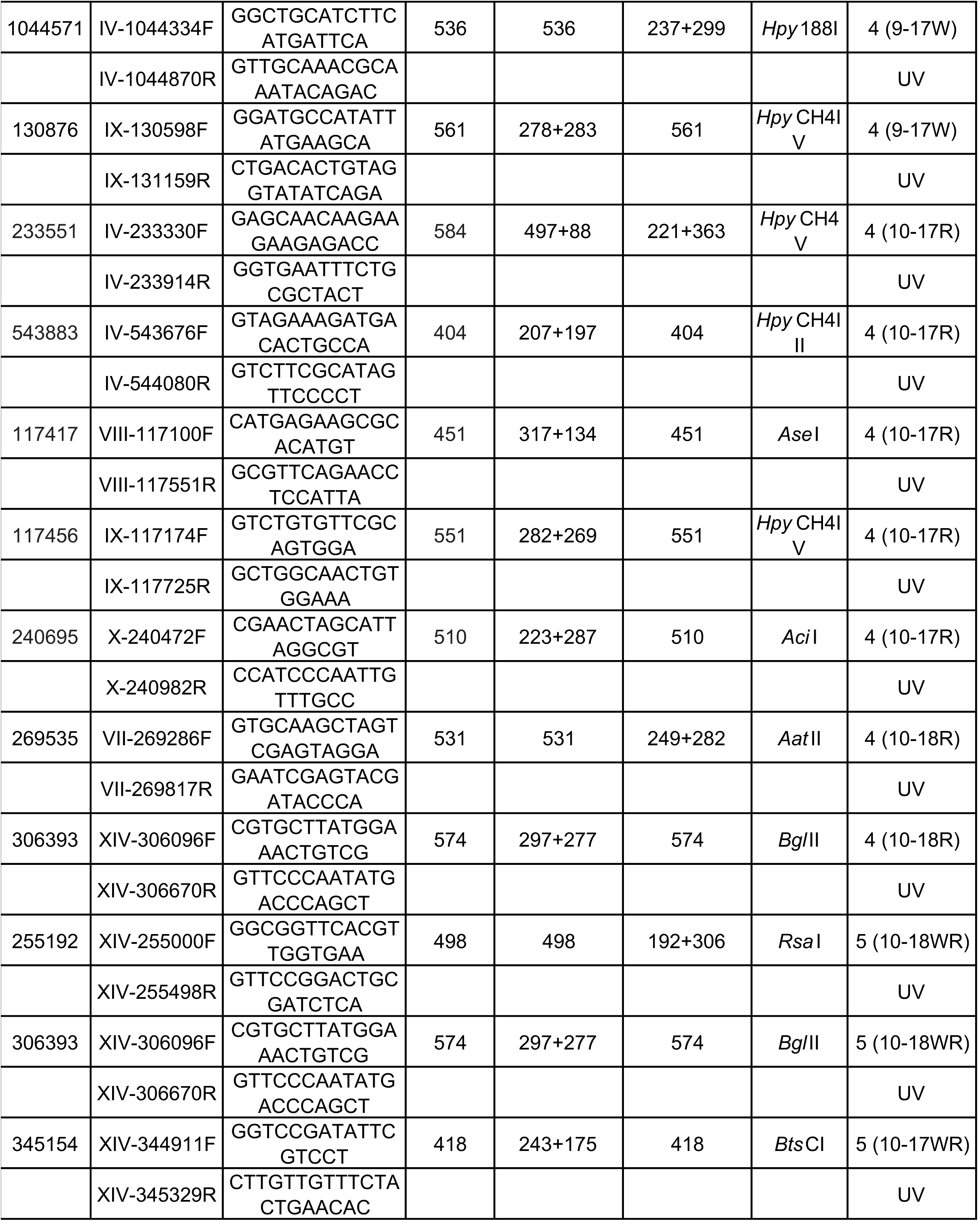
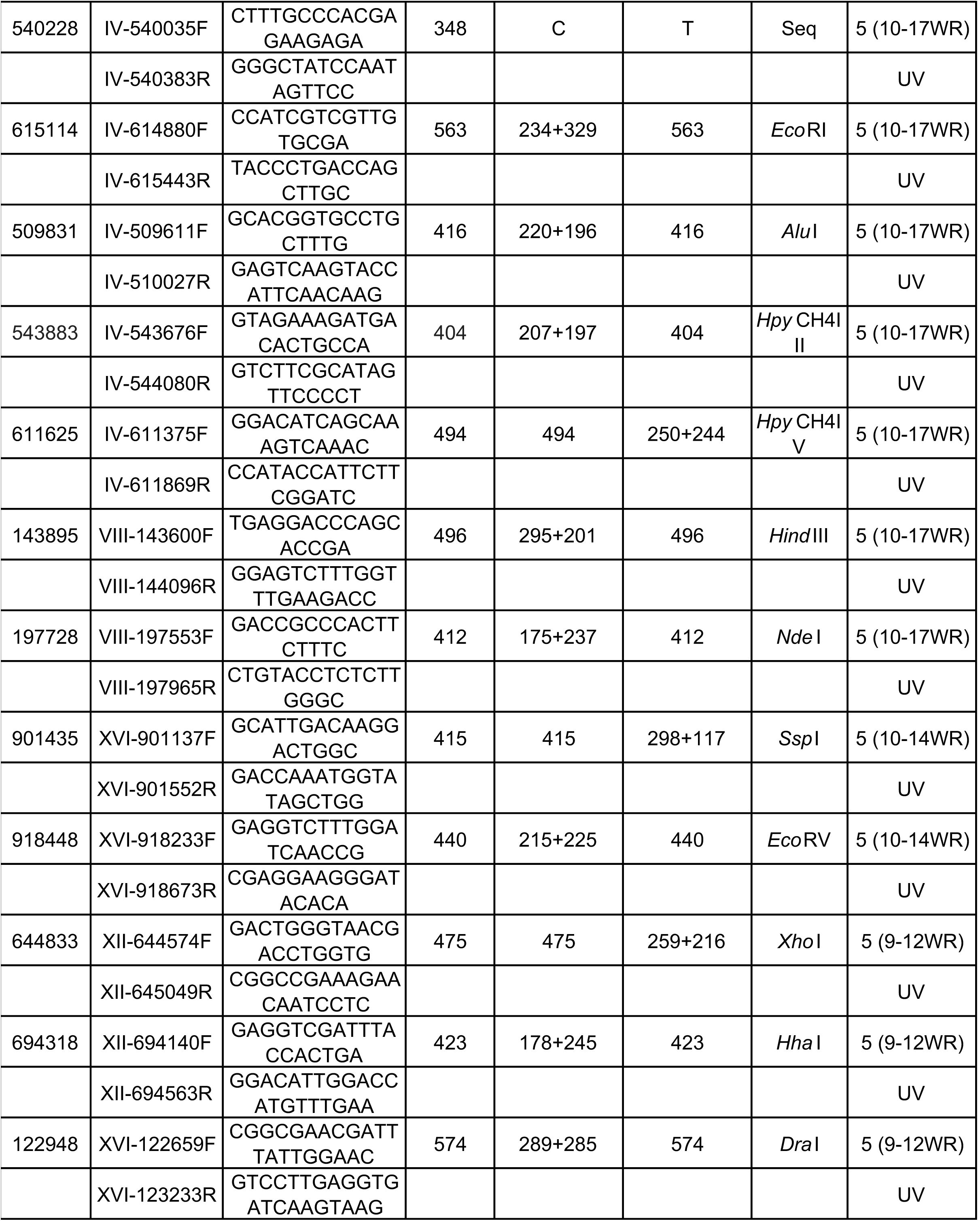
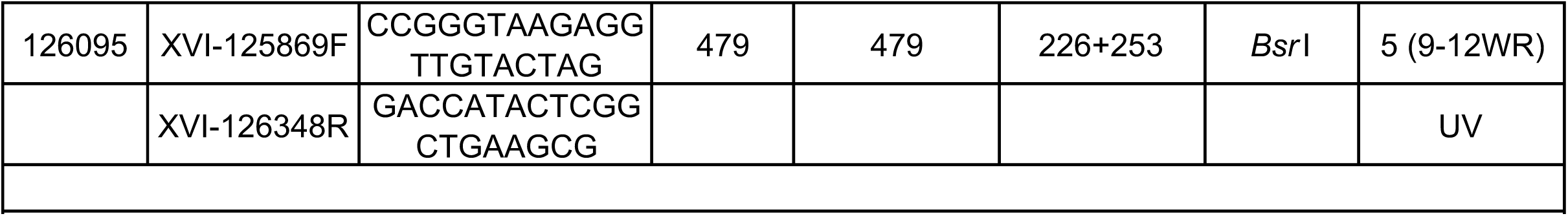
Primers and enzymes used in PCR and restriction enzyme analysis.

## Supplemental figure legends

**Fig. S1.**
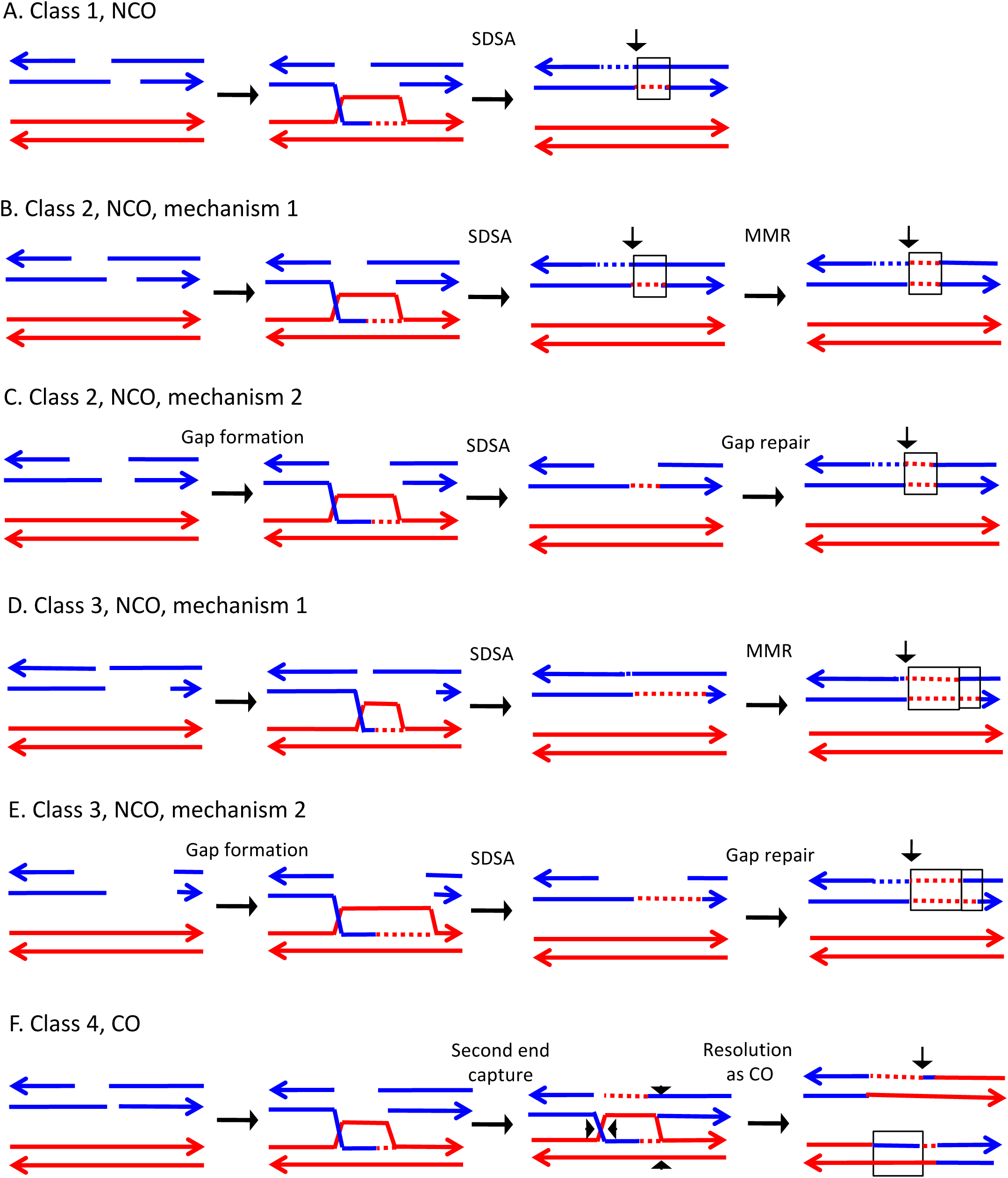
Mechanisms that generate Classes 1-5 recombination events. For all events, we show the initiating DSB on the blue chromatid. Dotted lines show sequences generated by replication or during mismatch repair. Black arrows indicate the position of the initiating DSB. The mechanisms are discussed in detail in the main text. A. Class 1. Class 1 events are NCOs formed by SDSA. B. Class 2. These NCO events could be generated by SDSA, followed by Mlh1p-independent MMR. C. Class 2. An alterative possibility is that Class 2 events reflect repair of a double-stranded DNA gap. D. Class 3. In this NCO class, the heteroduplex region is adjacent to a conversion tract. Such events could reflect a heteroduplex tract in which mismatches are repaired in one part of the tract and left unrepaired in the other. E. Class 3. An alternative model for this class is that the conversion tract is the result of repair of a double-stranded DNA gap with a heteroduplex region at one end. F. Class 4. For this CO class, a heteroduplex is observed on one chromatid but not the other. This class could be explained by the DSBR pathway in which heteroduplex region is short relative to the other; if the short heteroduplex does not contain a mismatch, it would be undetectable.

**Fig. S2.**
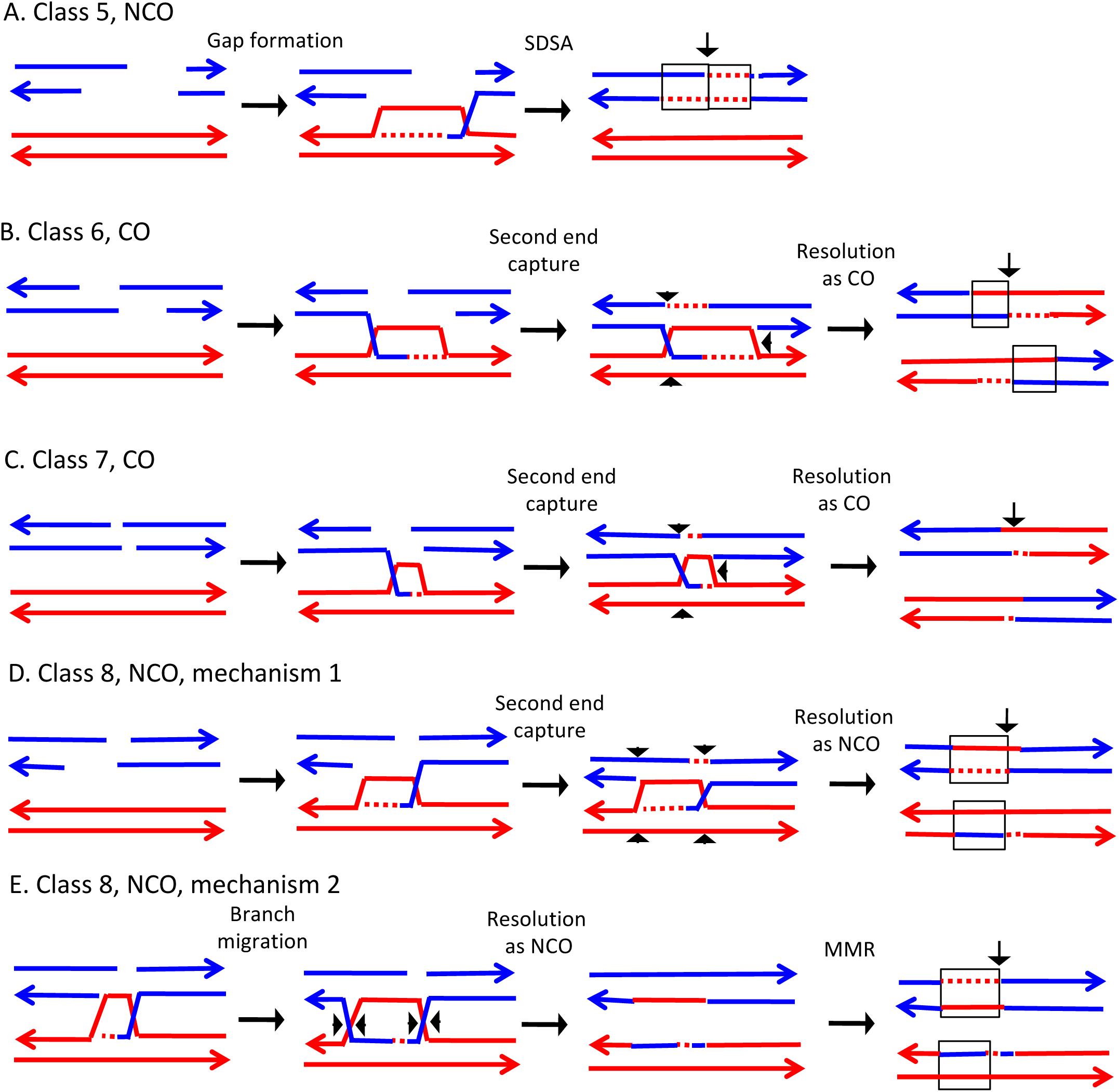
Mechanisms that generate Classes 6-8 recombination events. Events are depicted as in Fig. S2. A. Class 5. In this NCO class, the heteroduplex region is located on the opposite of the DSB site from the conversion region. This pattern is consistent with the repair of a double-stranded DNA gap that was restricted to one of the broken ends. B. Class 6. This CO class is identical to the pattern expected for the DSBR model. C. Class 7. In this CO class, no heteroduplexes or conversion tracts are observed adjacent to the crossover, consistent with the formation of a dHJ with short heteroduplex tracts that do not include mismatches. D. Class 8, mechanism 1. In this NCO class, one chromatid has a conversion tract, and the other chromatid has a heteroduplex involving SNPs at the same position. This event could reflect resolution of a dHJ intermediate in the NCO model in which one region of heteroduplex is undetectable. E. Class 8, mechanism 2. An alternative mechanism involves branch migration of a HJ, followed by resolution of the intermediate in a NCO mode. Mismatches in one of the two chromatids are repaired to generate a conversion tract.

**Fig. S3-S91.**
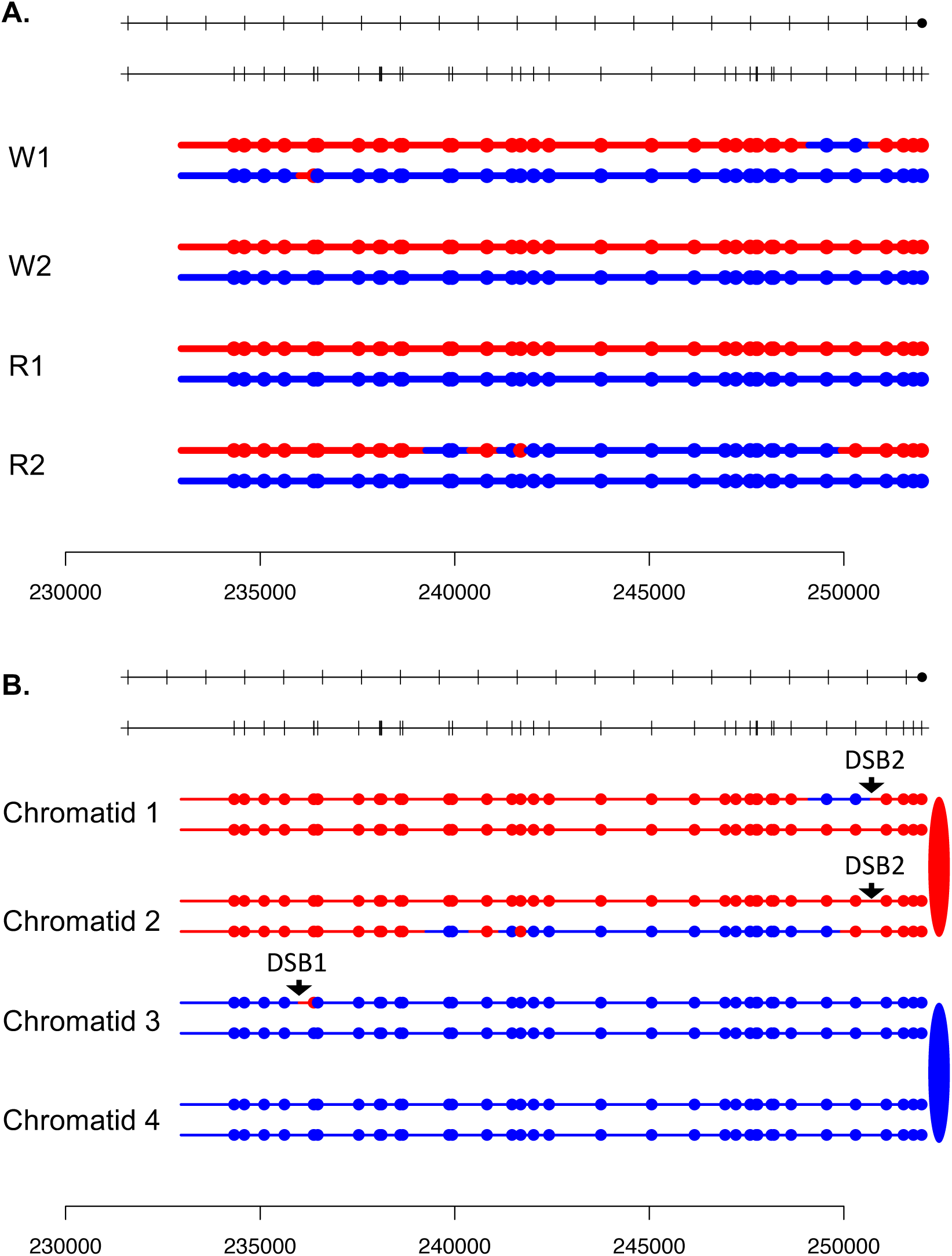
In the upper part of the figure (labeled A), we show the patterns of SNPs derived from the W303-1A homolog (red) and YJM789 homolog (blue). Each pair of lines is derived from one of the granddaughter cells in the white (W1 or W2) or red (R1 or R2) sectors. The locations of the SNPs, represented by colored circles, are drawn proportionally with a scale showing SGD coordinates at the bottom of A. In the B part of the figures, we show the arrangement of SNPs in double-stranded chromatids in the mother cell before chromosome segregation. The inferred positions of the initiating DSBs are indicated with arrows. Chromatids 1 and 2 are sisters, and chromatids 3 and 4 are sisters. Figure S3-S80 show selected and unselected UV-induced events in strain YYy310. Figures S81-S85 show selected spontaneous events on chromosome V in strain YYy310, and Figures 86-91 represent spontaneous HS4-related crossovers on chromosome V in strain YYy311. The coordinates and classifications of the events represented by these supplemental figures are in Tables S2 and S3, respectively. For those events described as “Complex”, we show possible mechanistic pathways for their formation in Figs. S92-S135. Lastly, we note that the event depicted in Fig. S71 has the pattern expected for a BIR event in one of the daughter cells. Because of this complication, we did not attempt to infer the locations of the initiating DSBs.

**Fig. S4.**
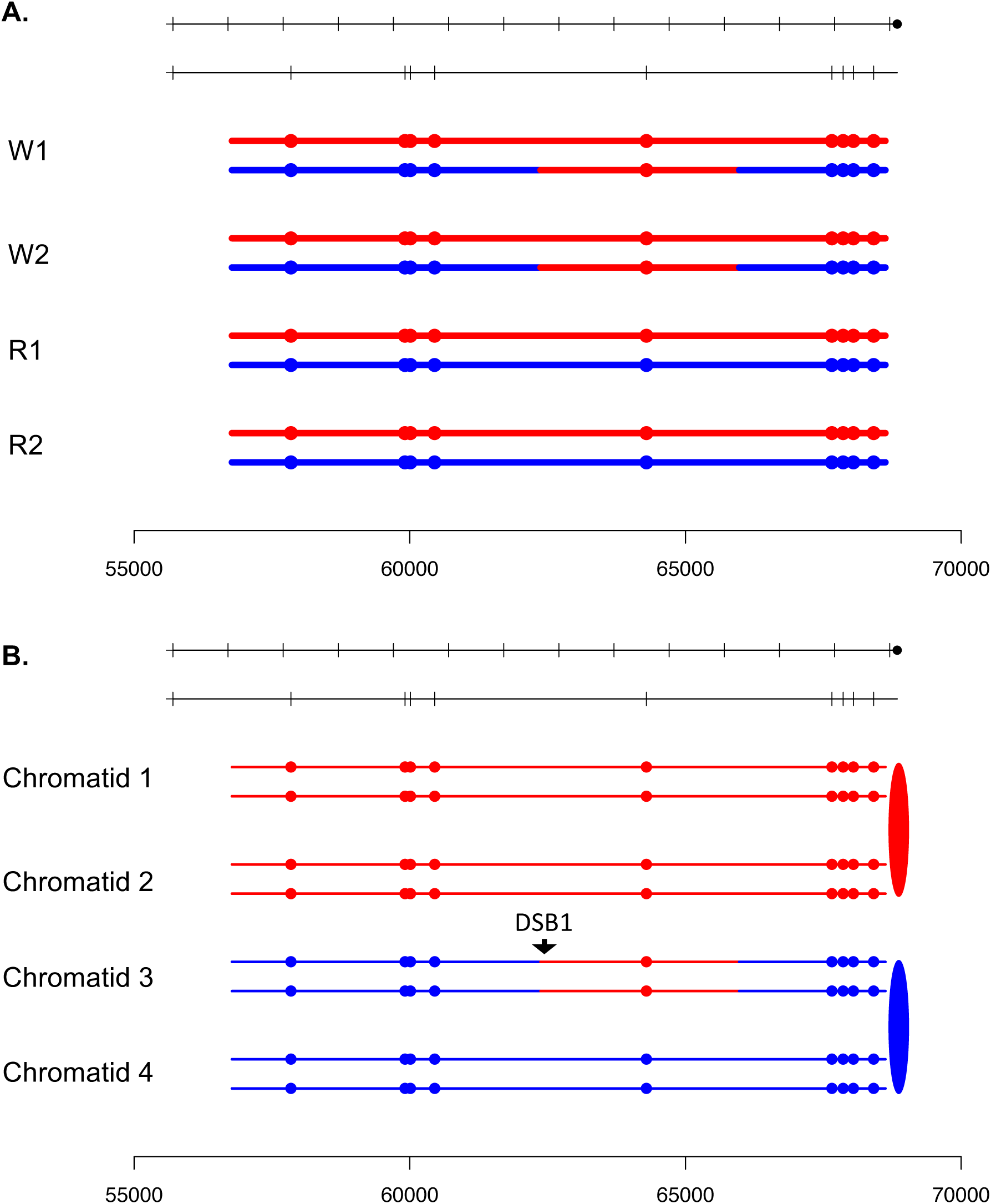

**Fig. S5.**
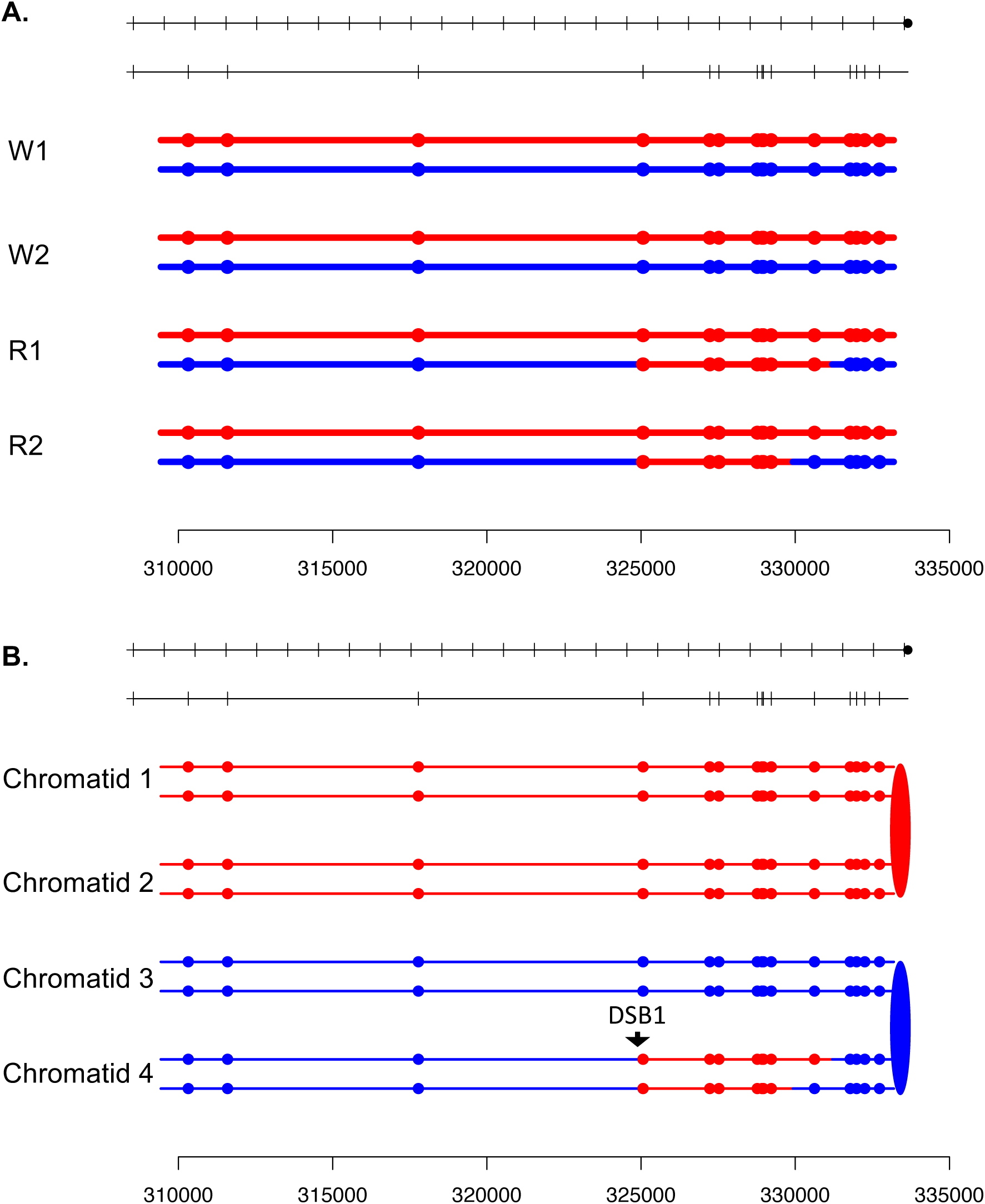

**Fig. S6.**
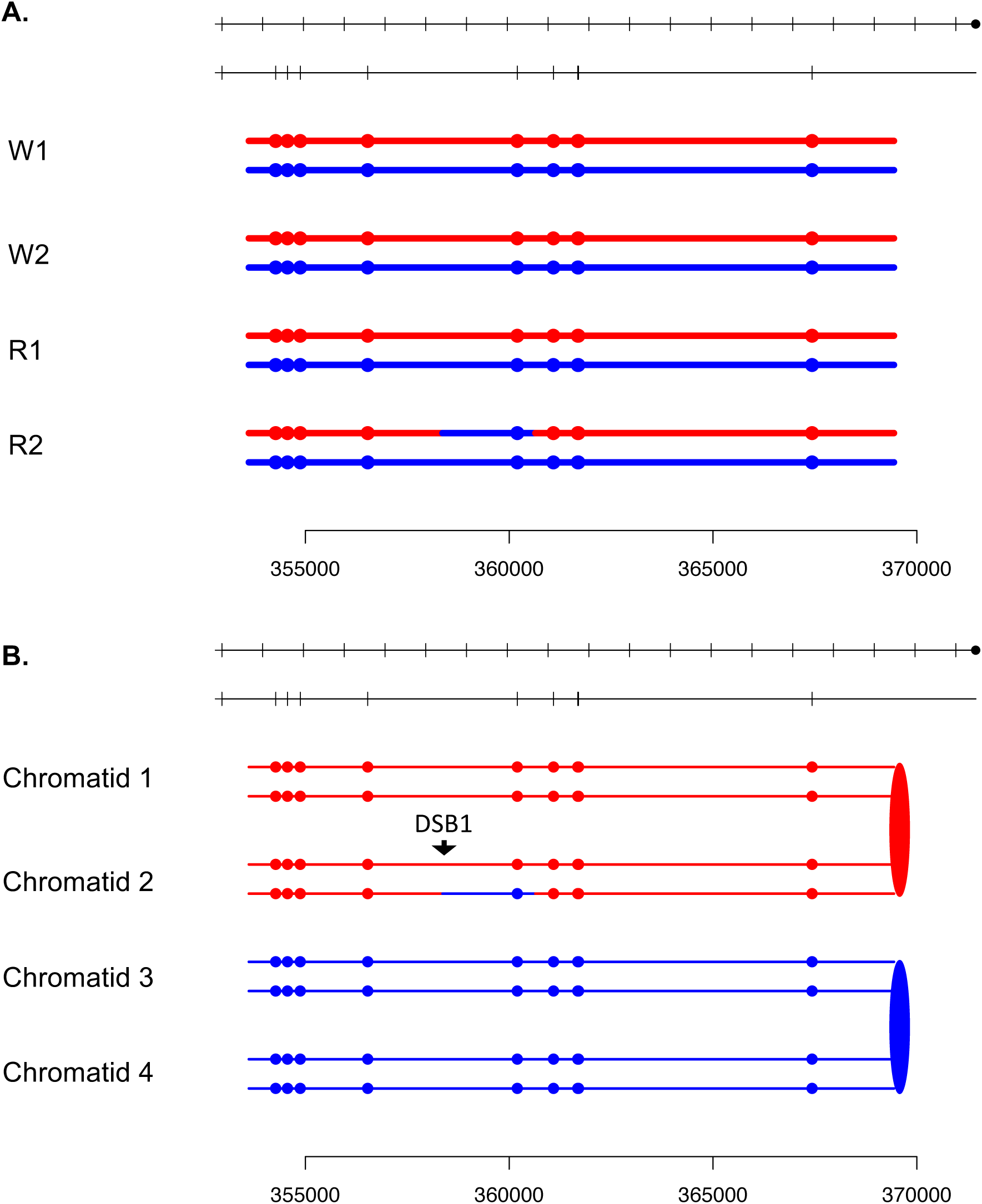

**Fig. S7.**
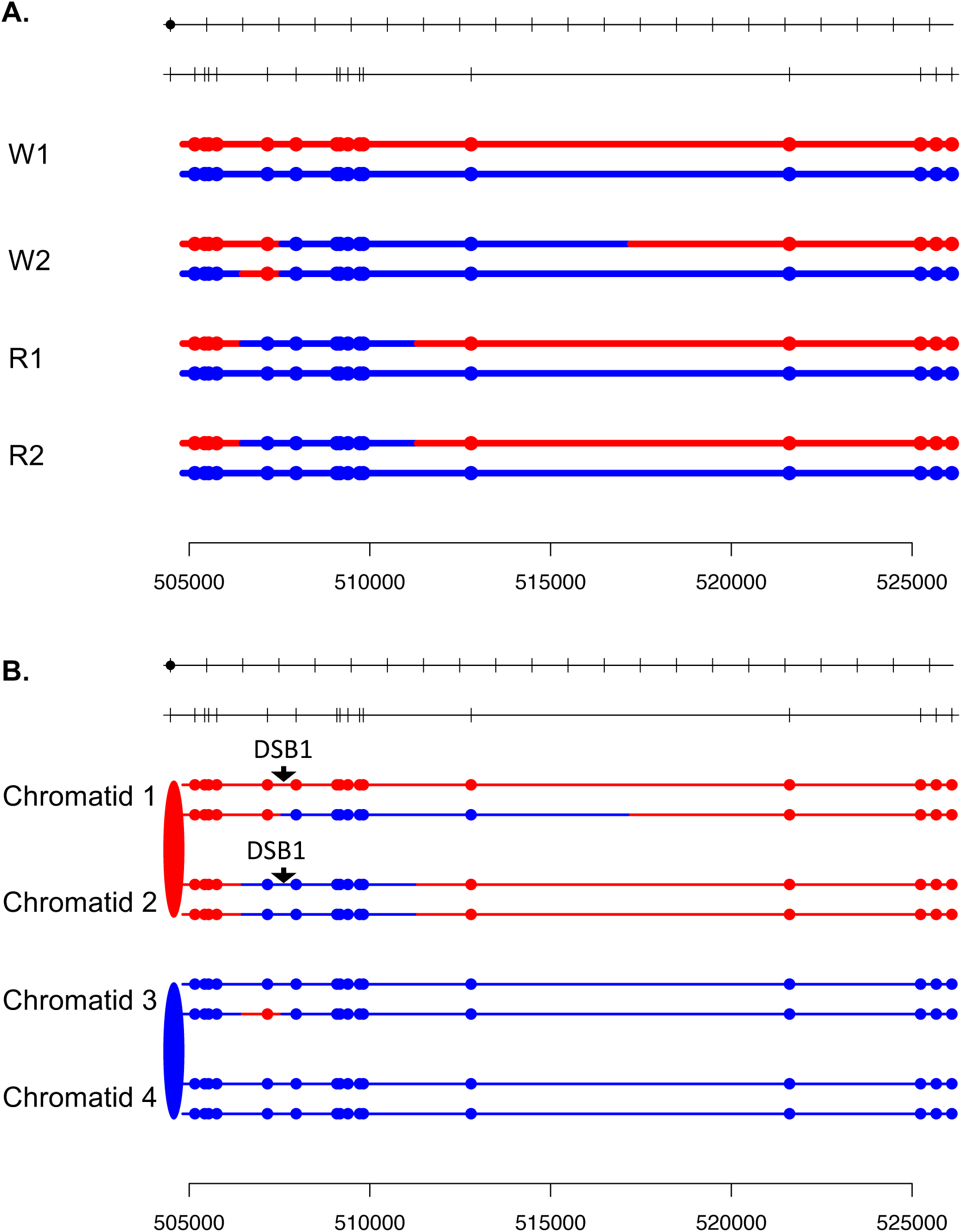

**Fig. S8.**
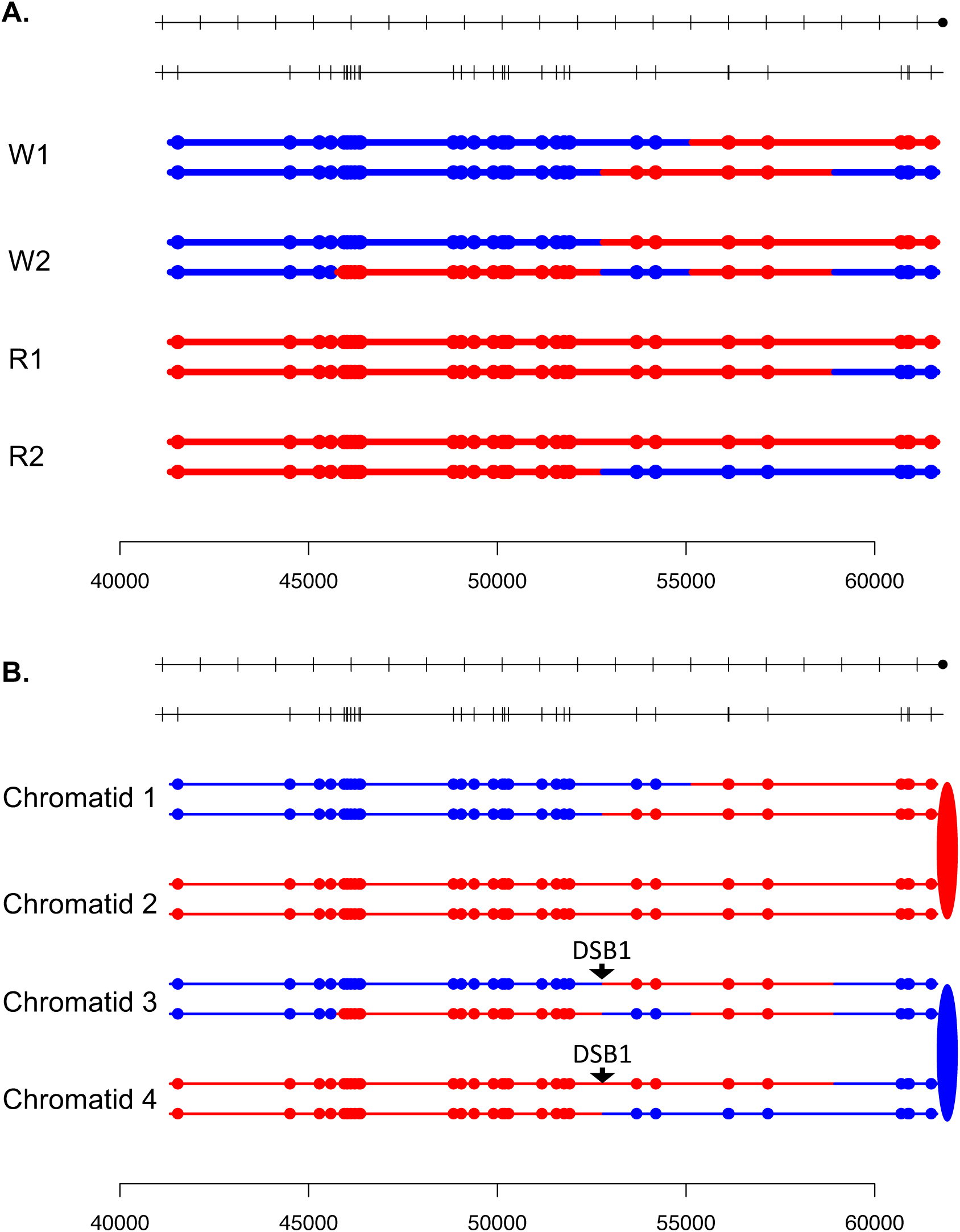

**Fig. S9.**
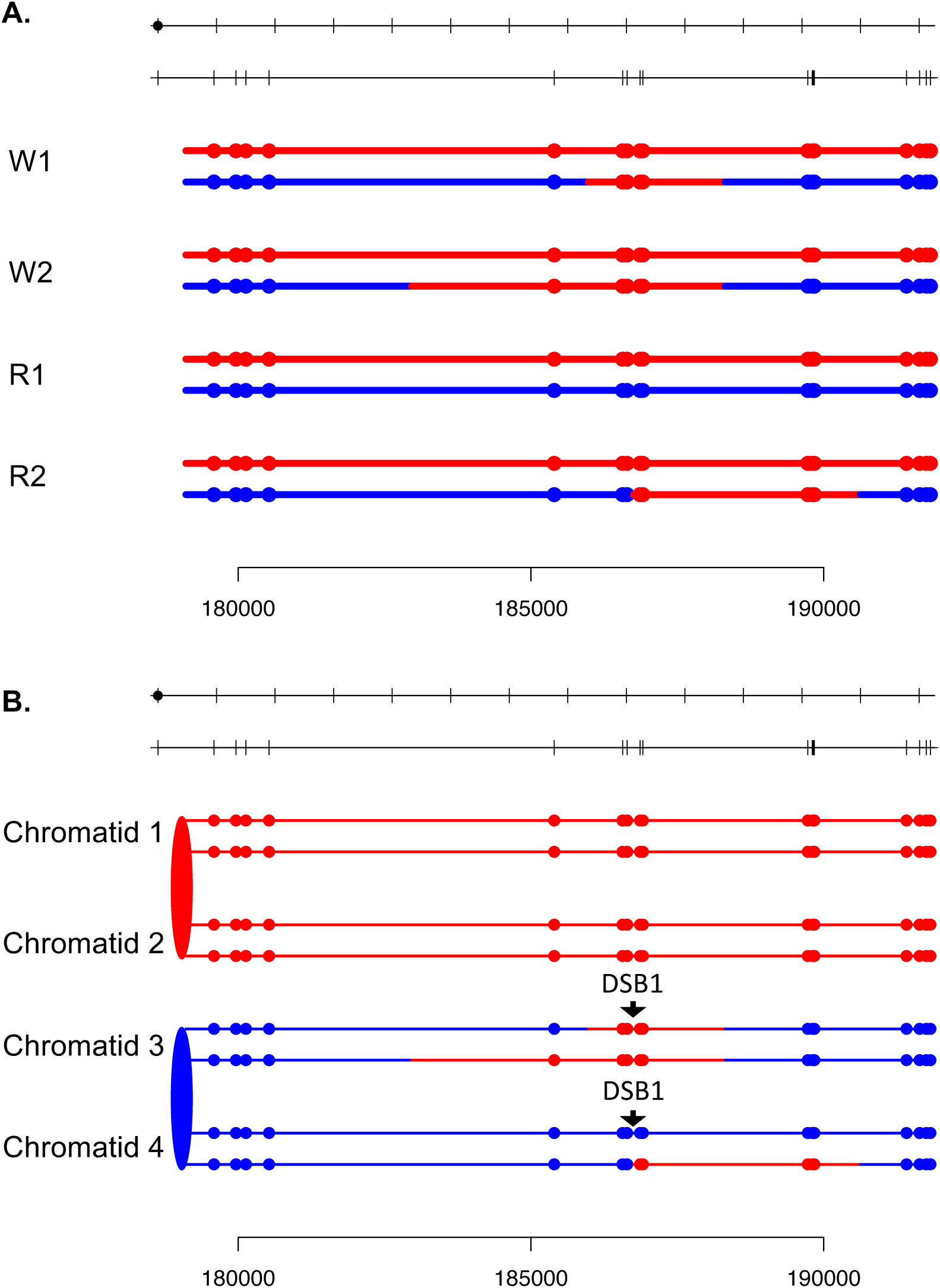

**Fig. S10.**
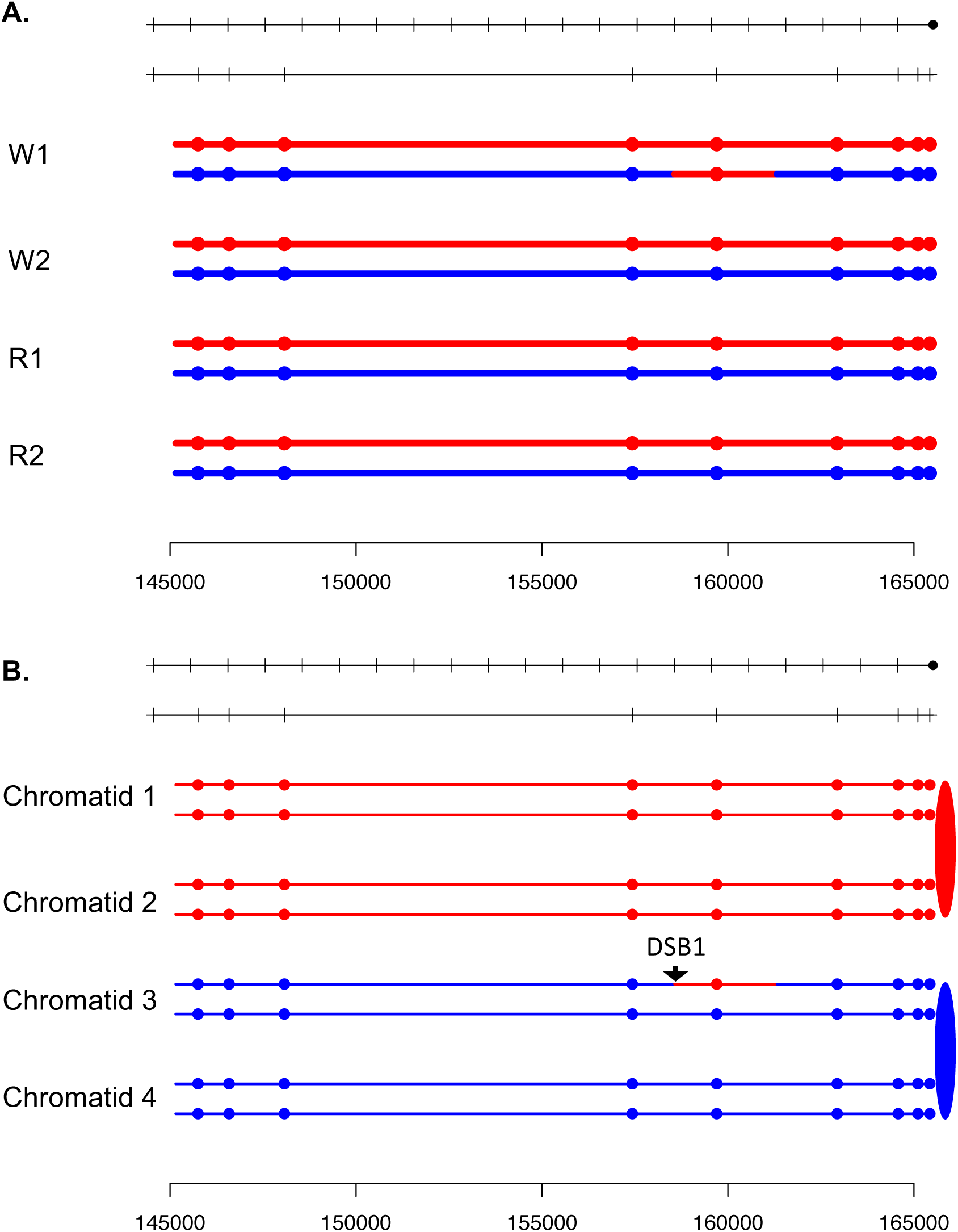

**Fig. S11.**
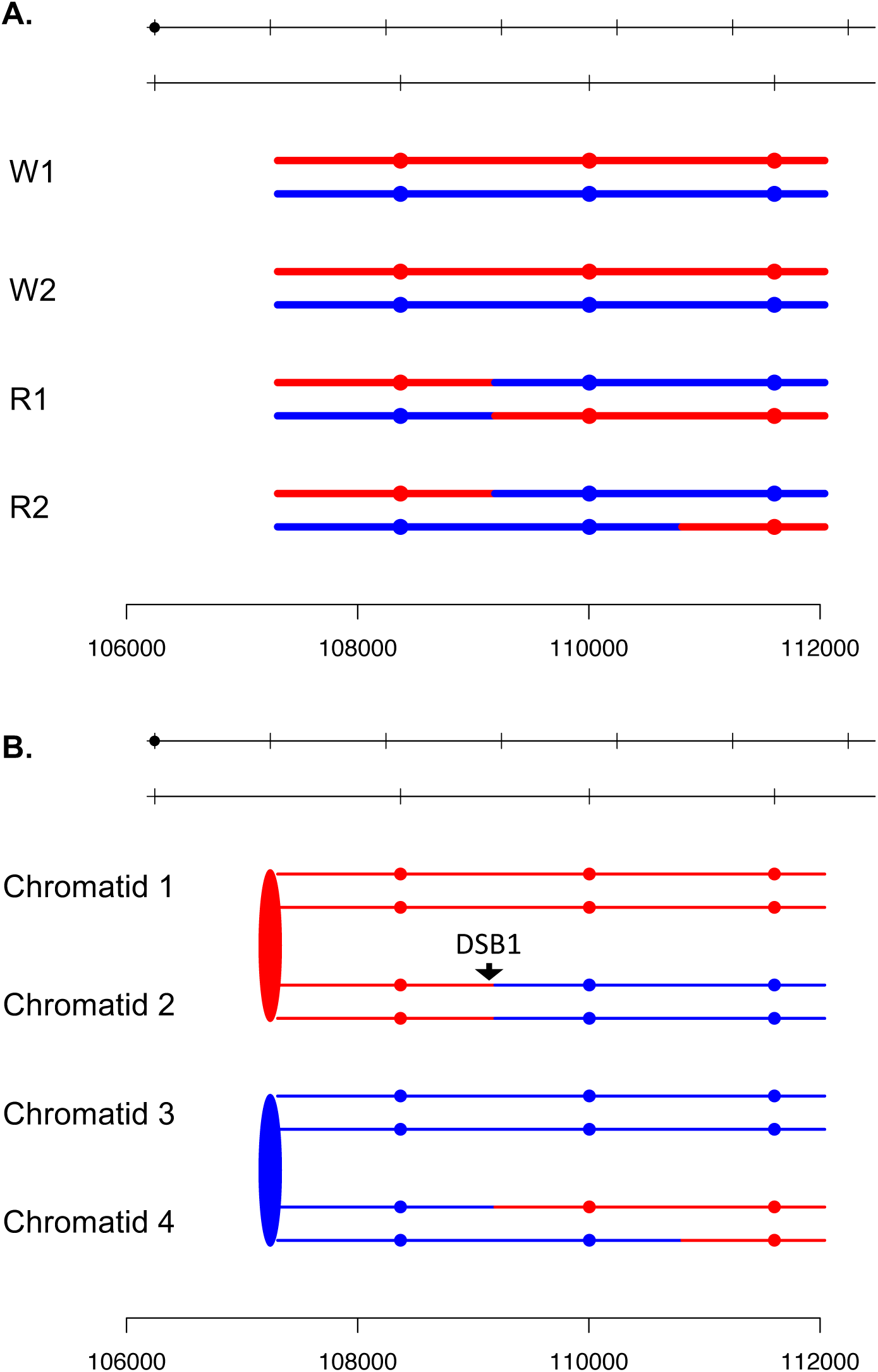

**Fig. S12.**
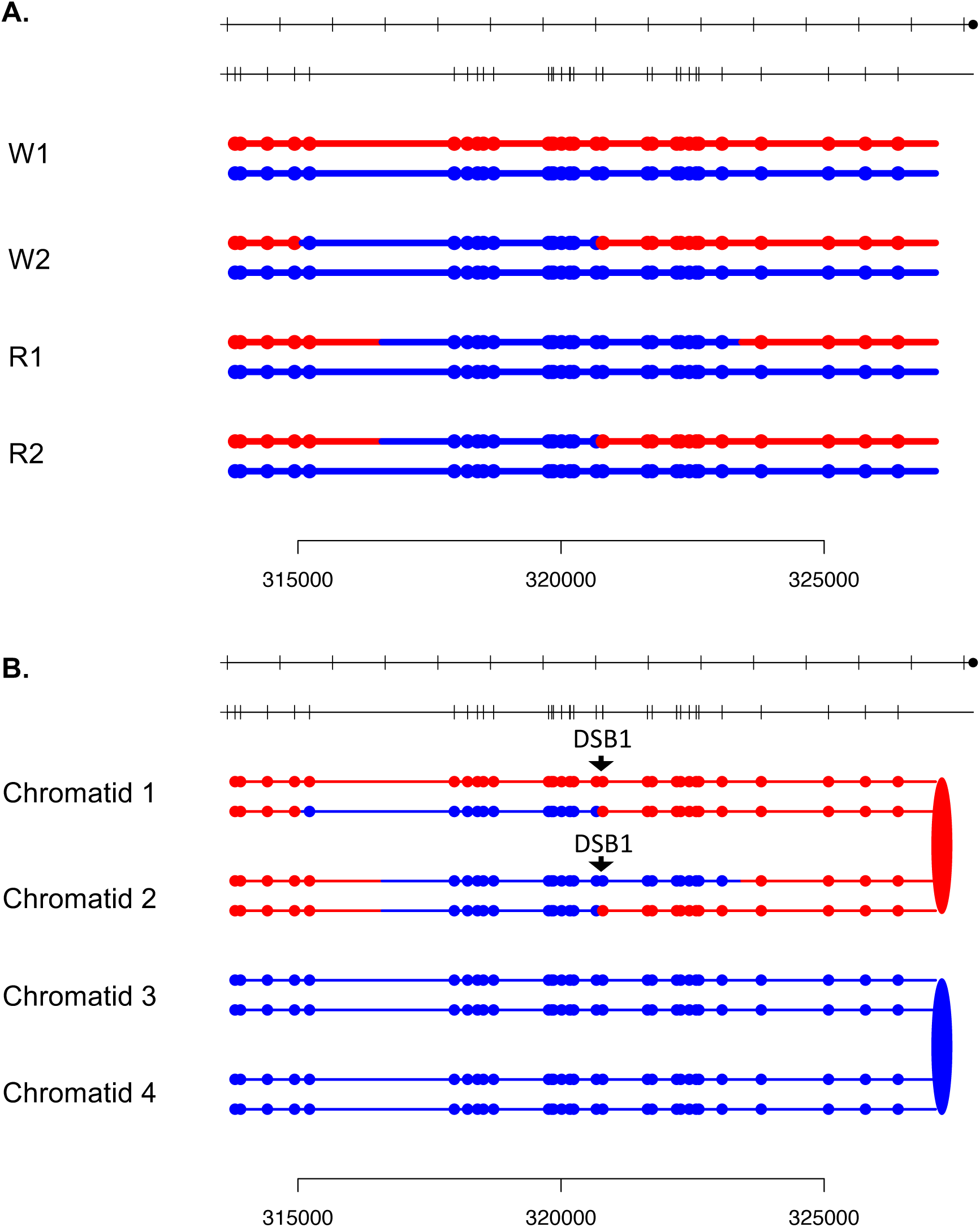

**Fig. S13.**
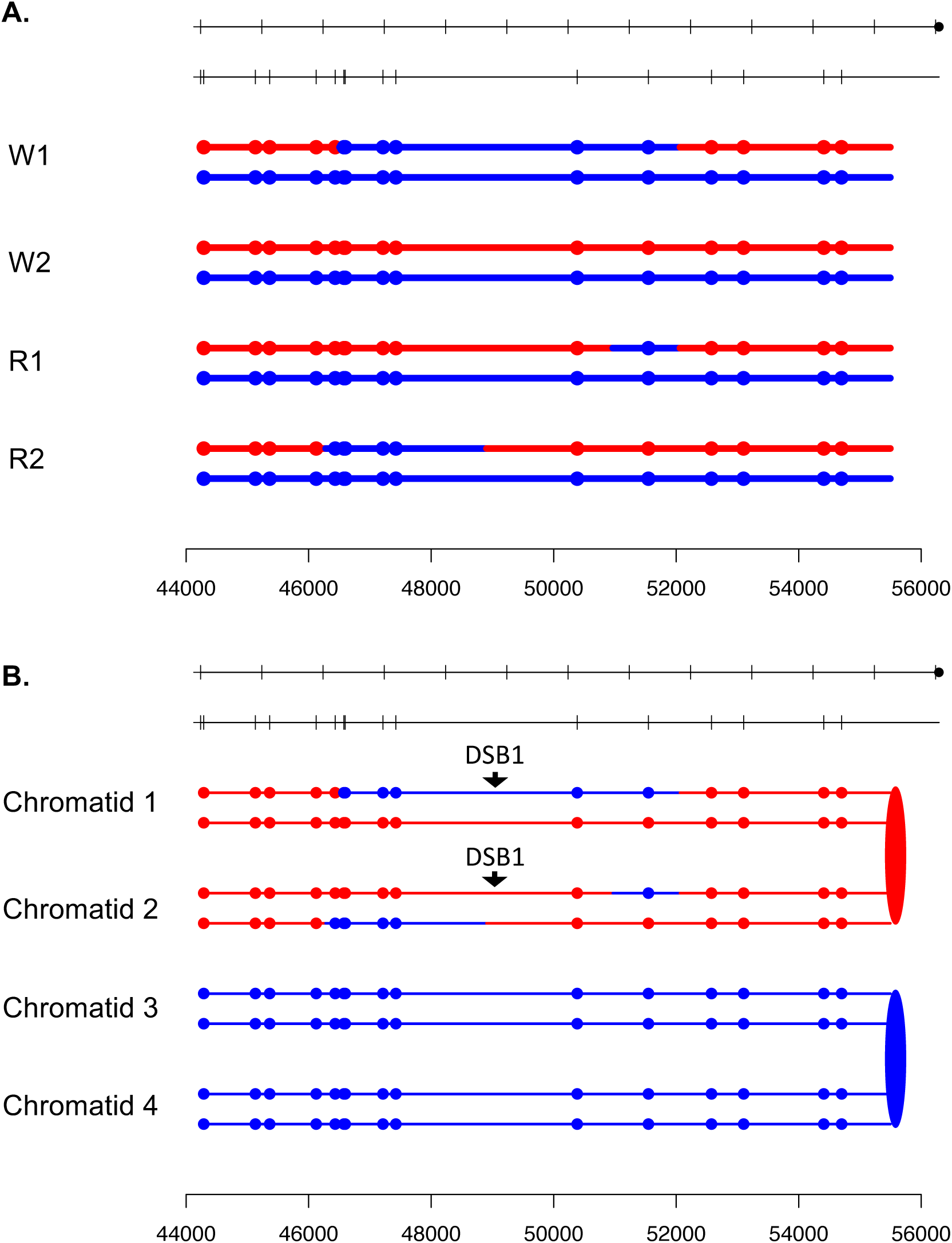

**Fig. S14.**
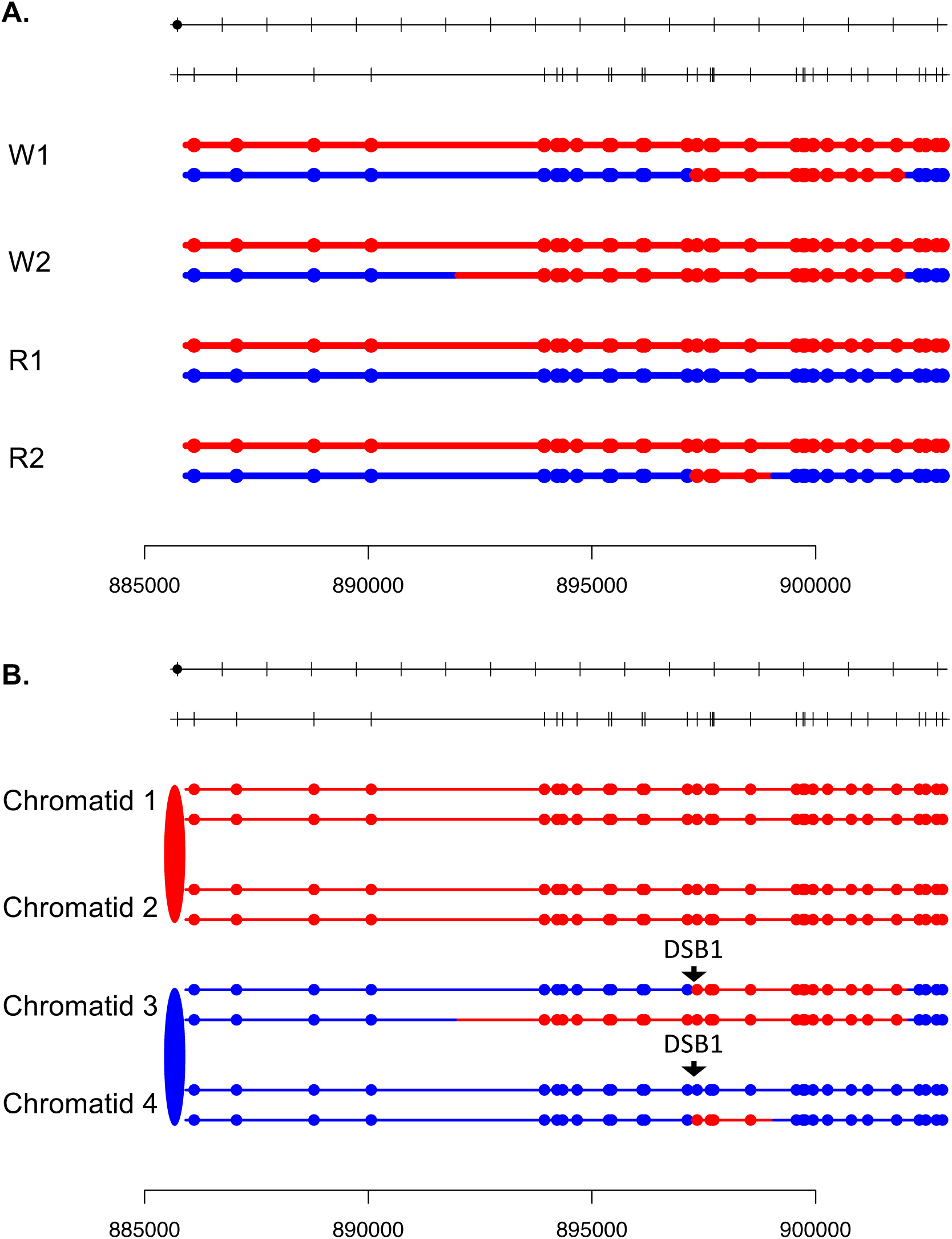

**Fig. S15.**
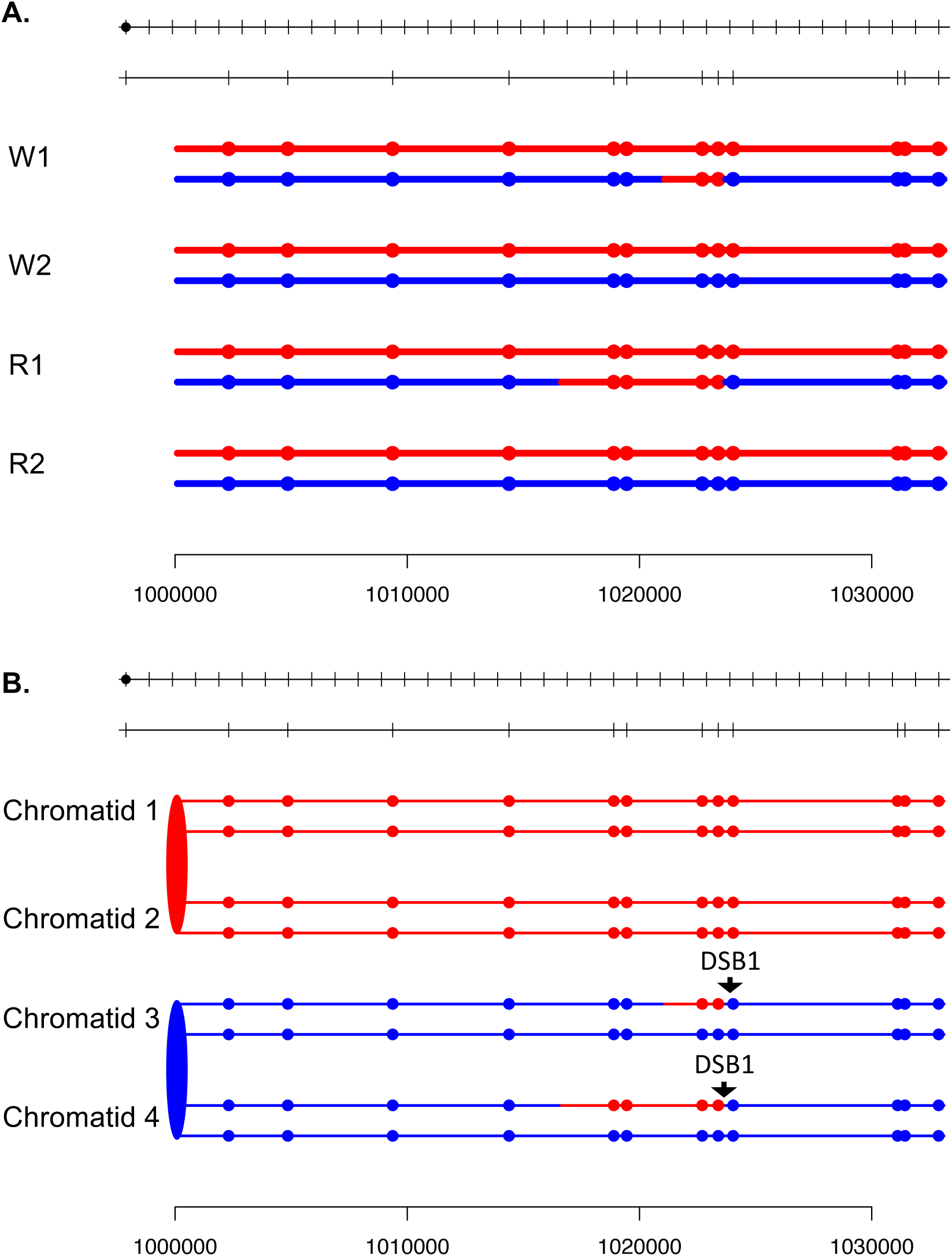

**Fig. S16.**
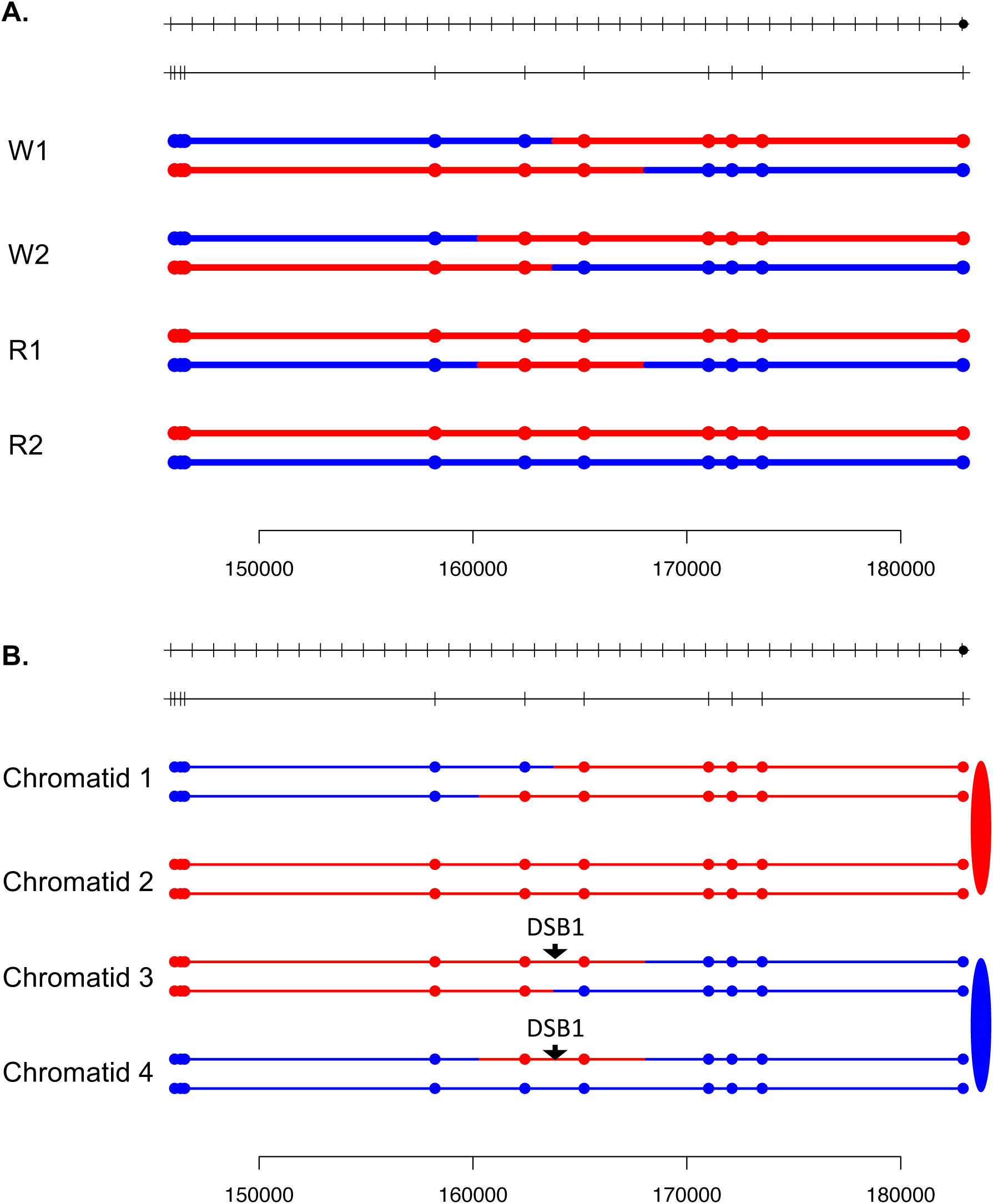

**Fig. S17.**
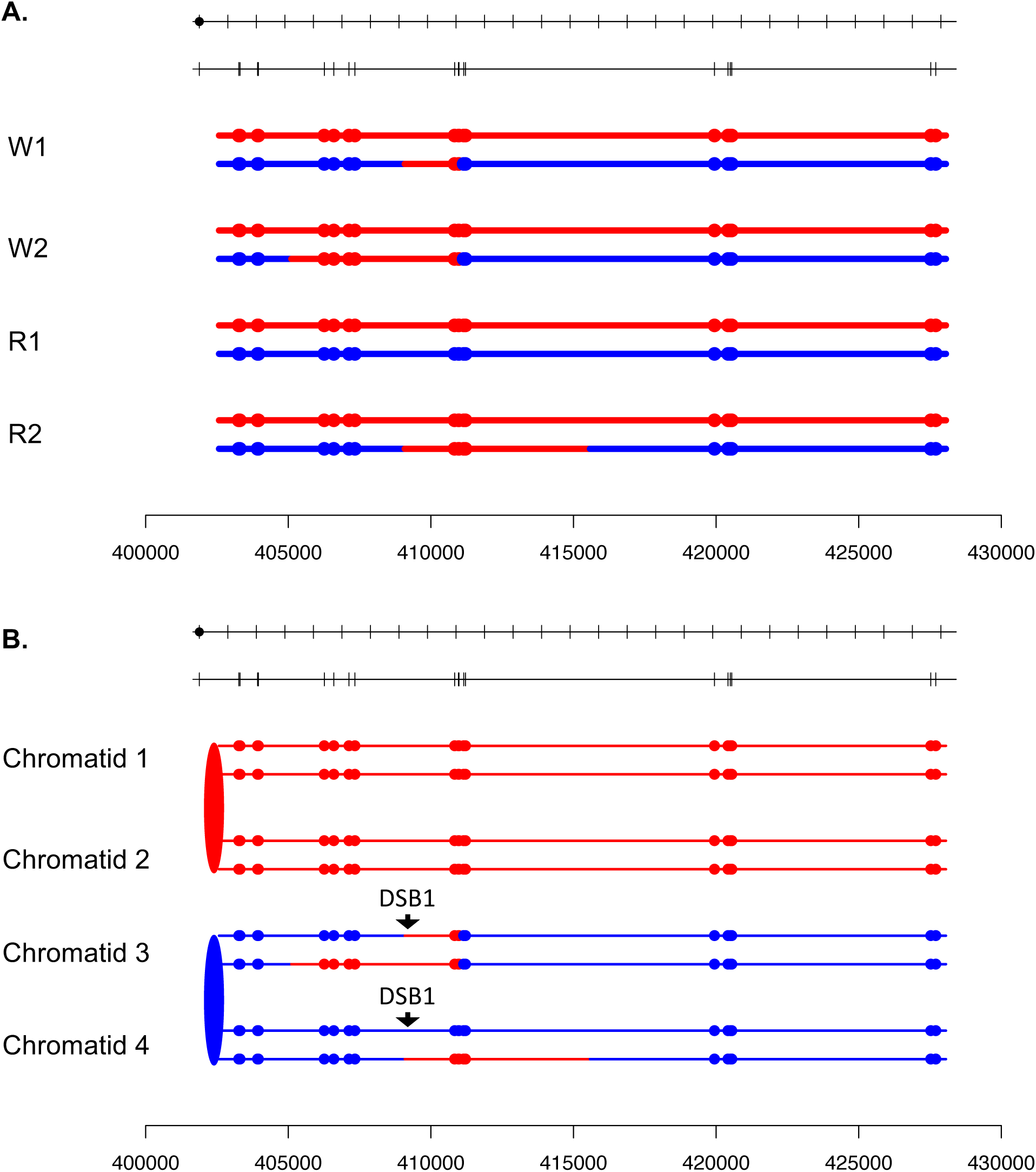

**Fig. S18.**
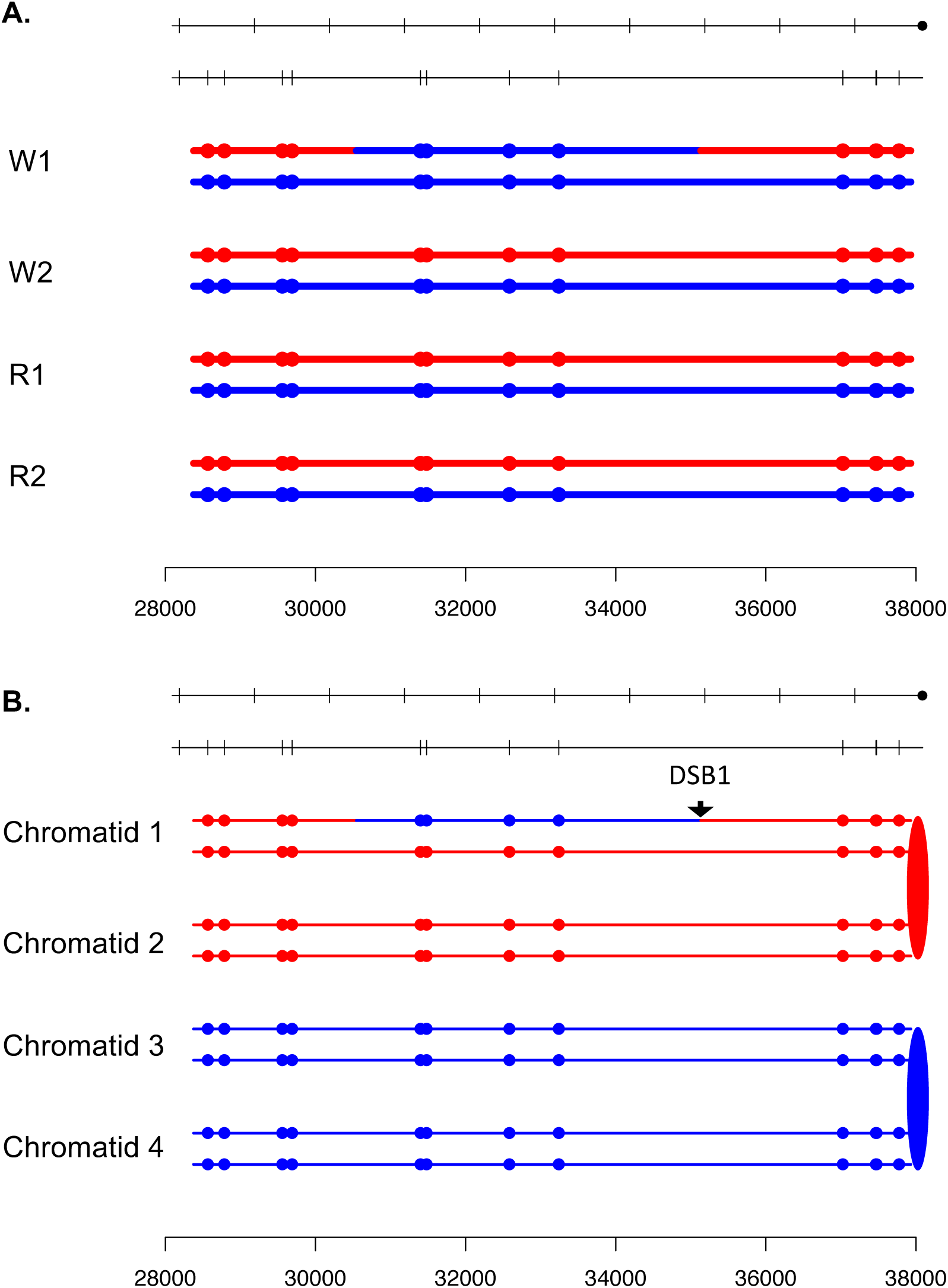

**Fig. S19.**
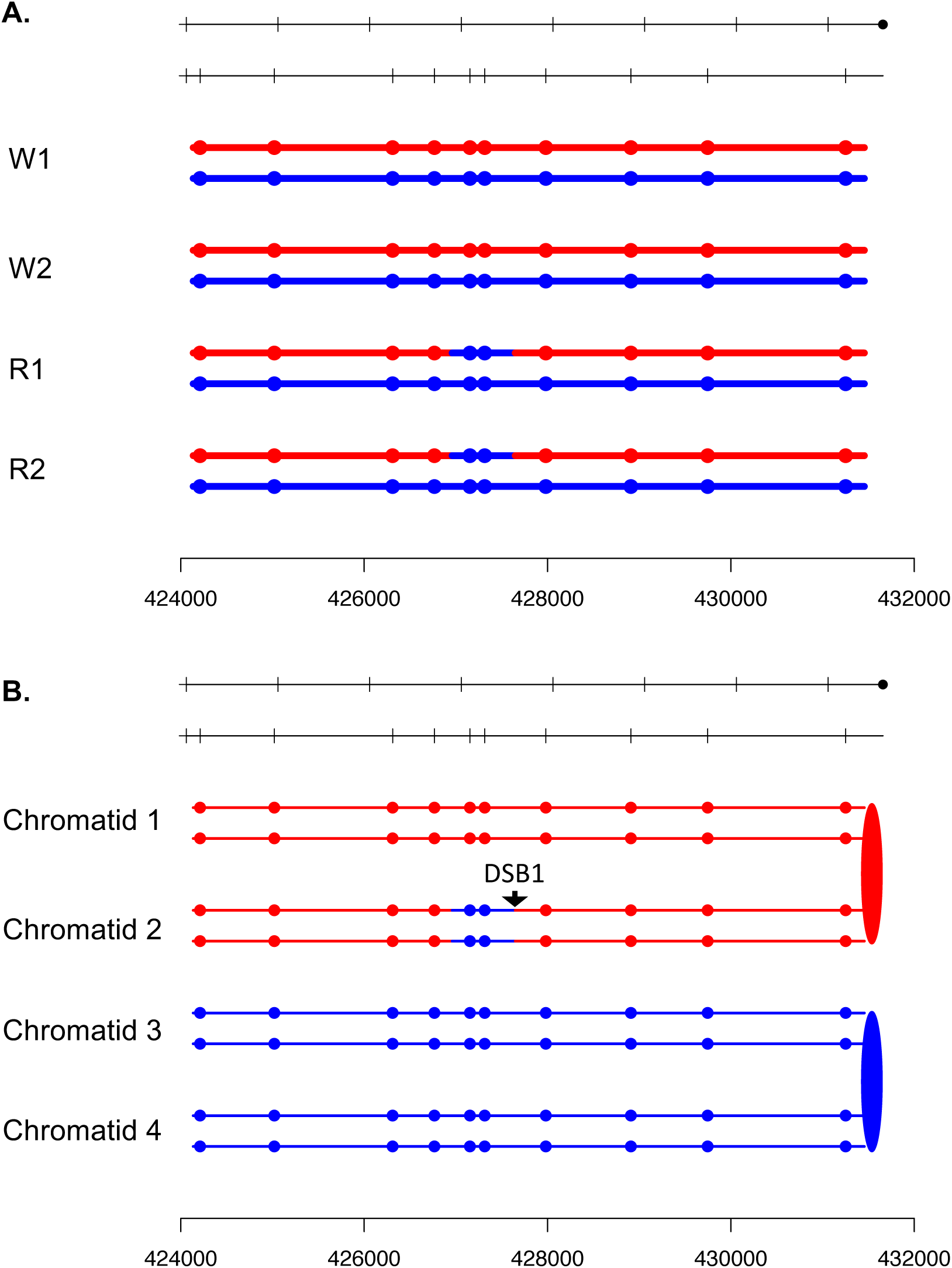

**Fig. S20.**
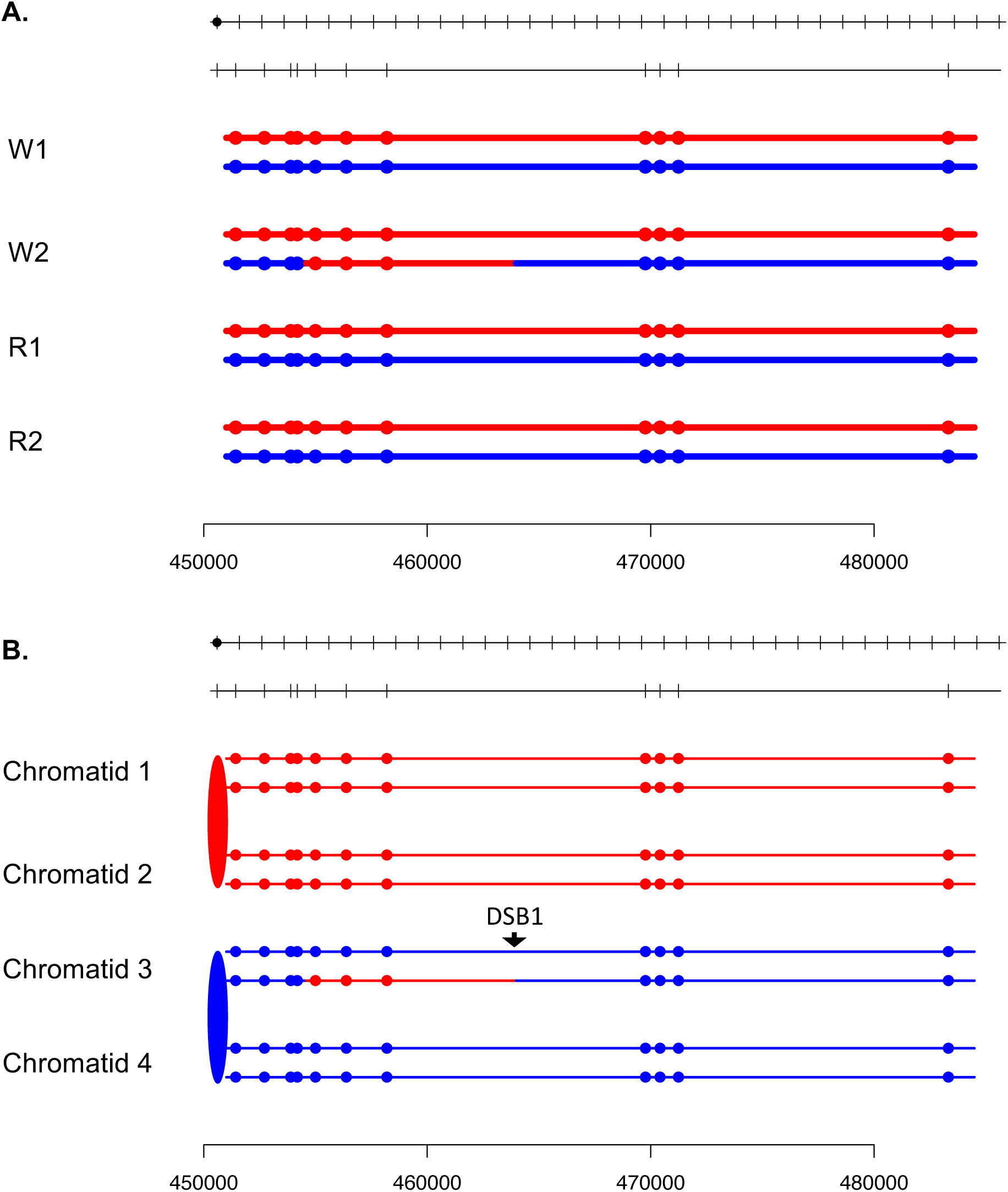

**Fig. S21.**
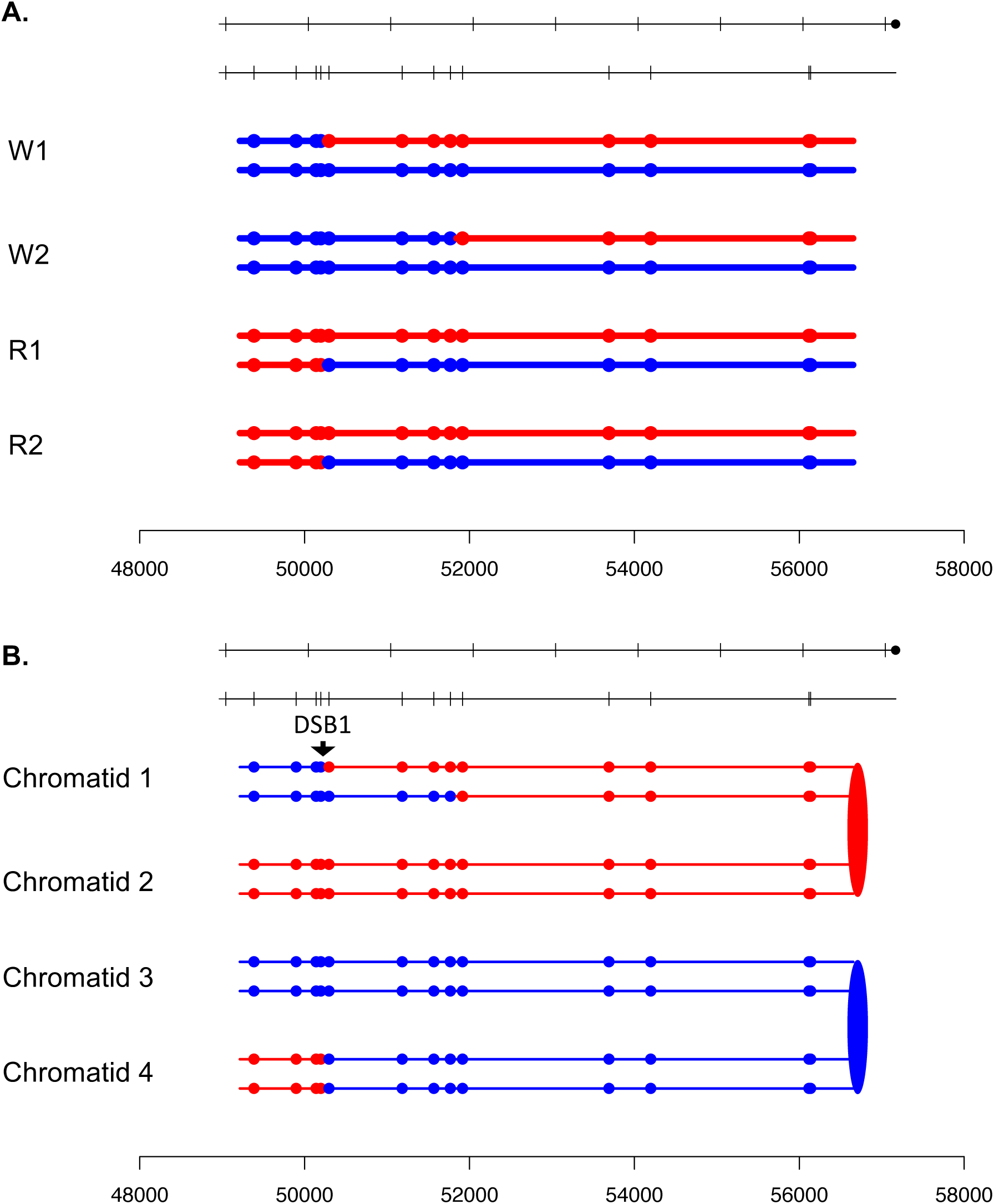

**Fig. S22.**
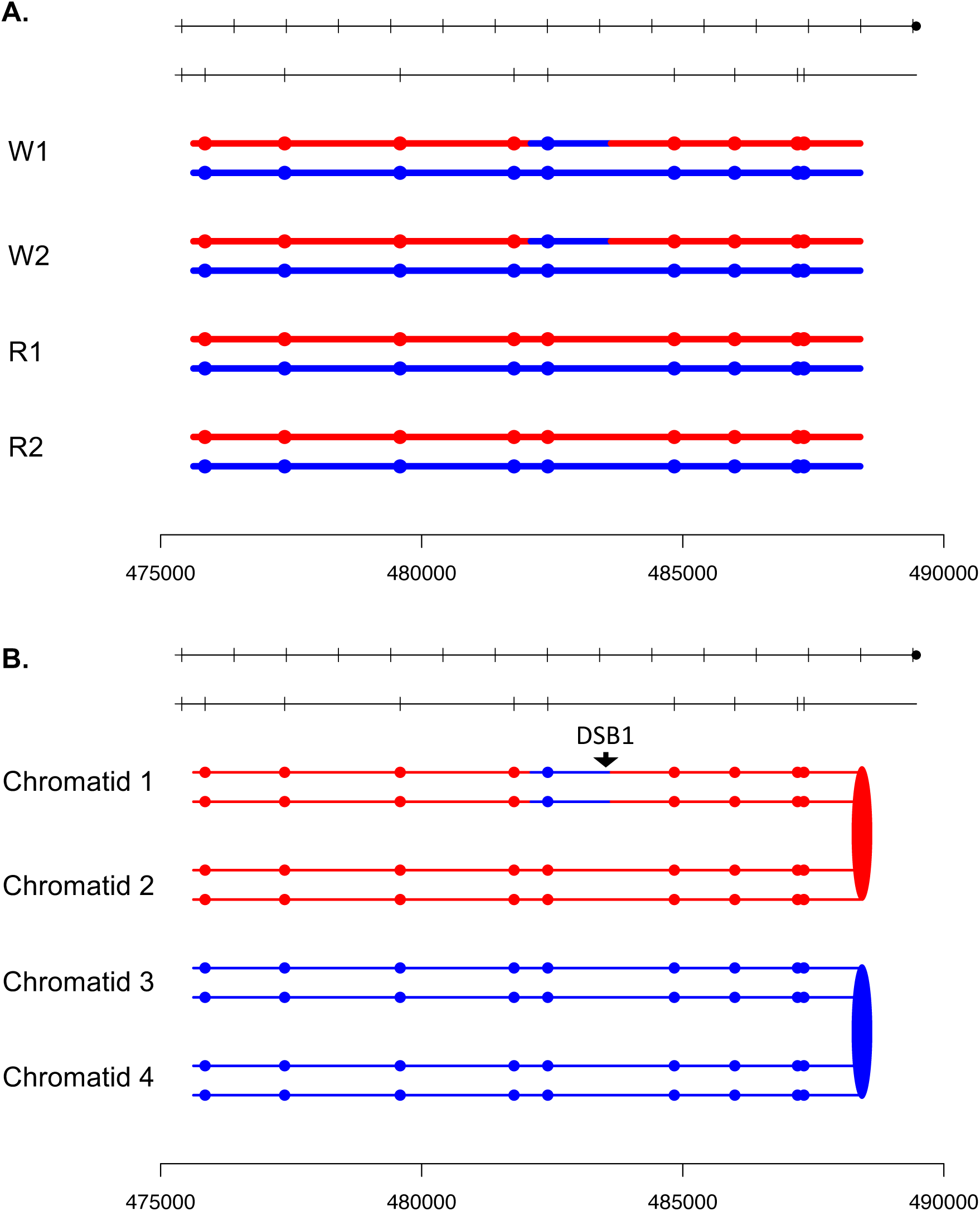

**Fig. S23.**
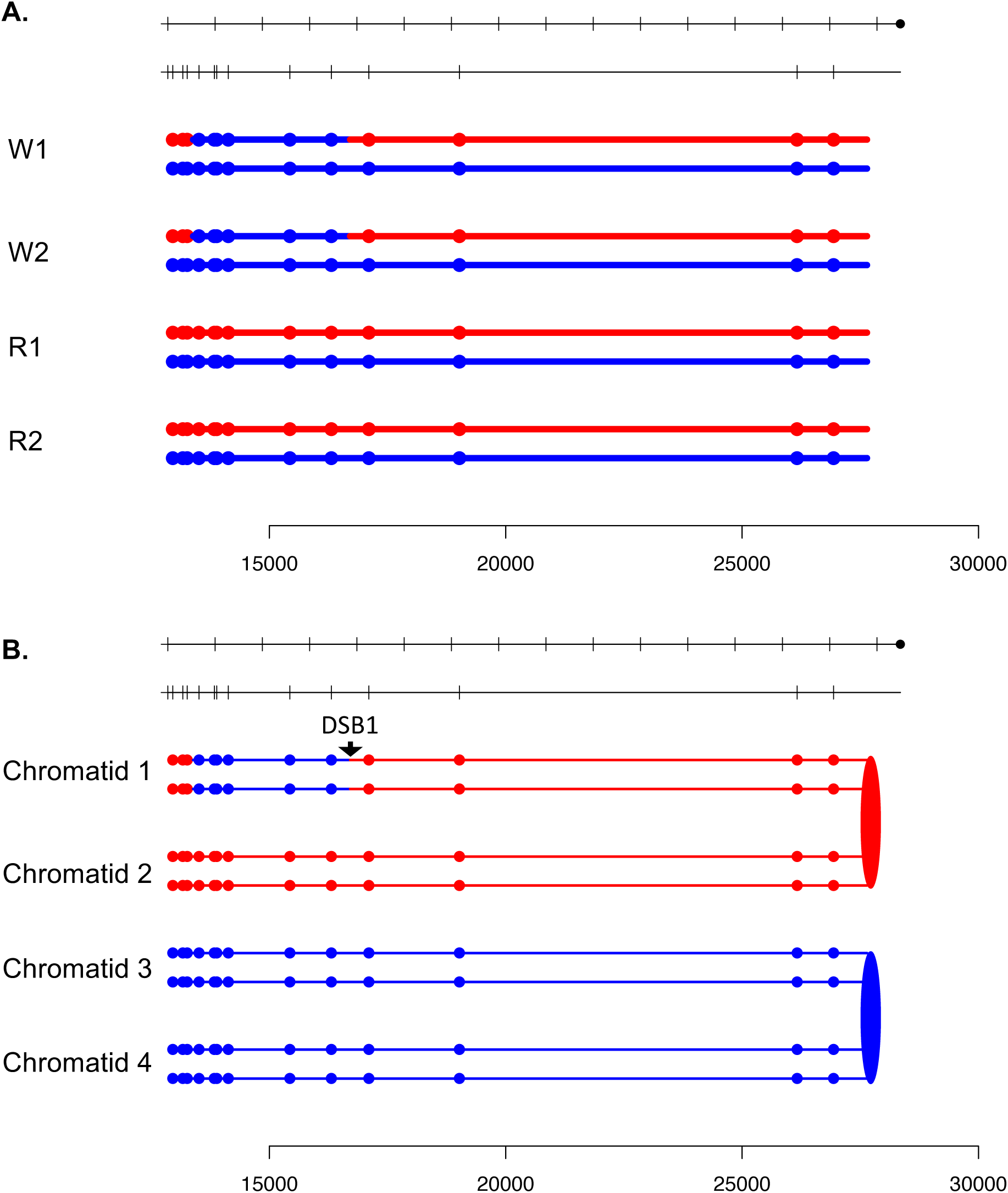

**Fig. S24.**
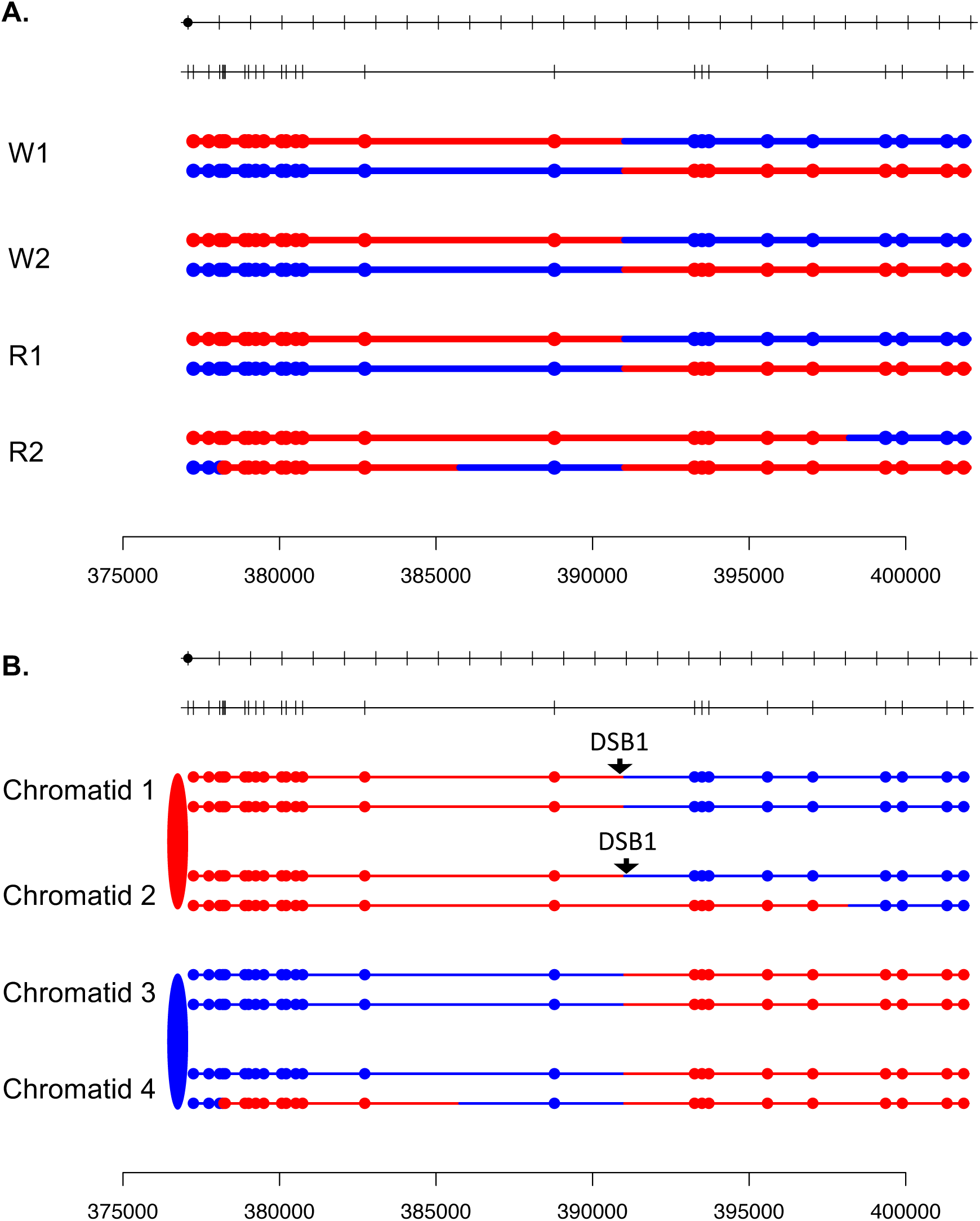

**Fig. S25.**
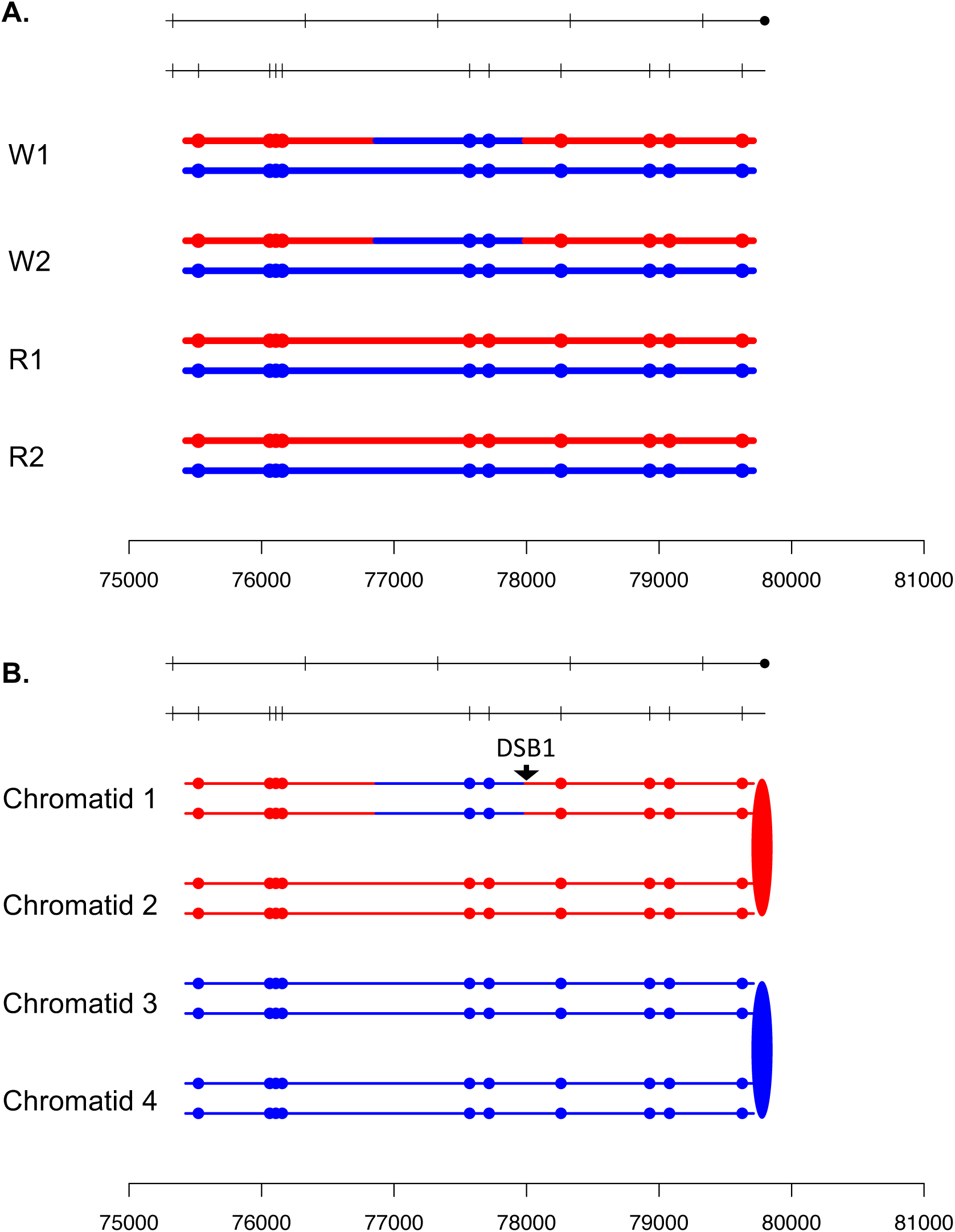

**Fig. S26.**
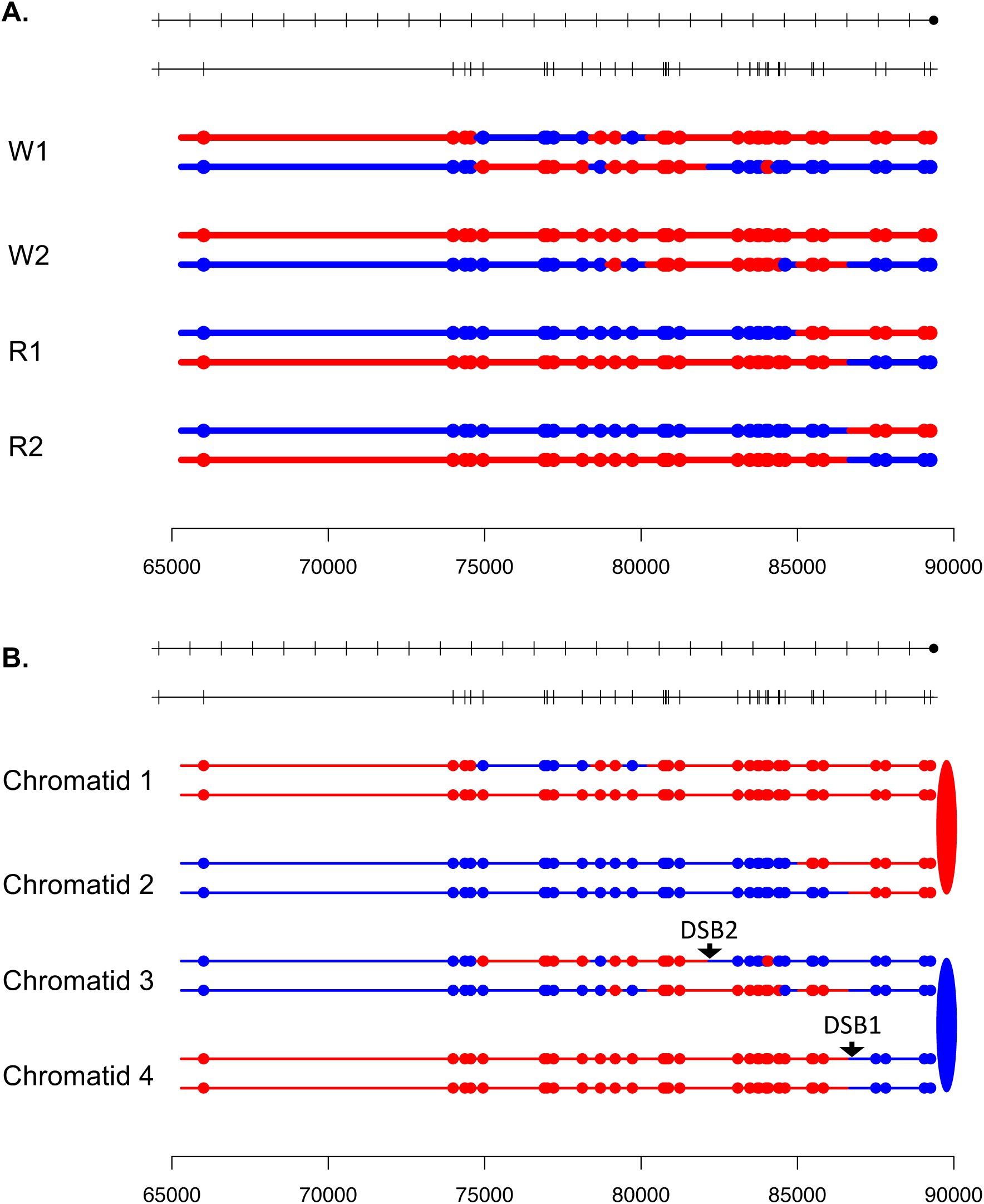

**Fig. S27.**
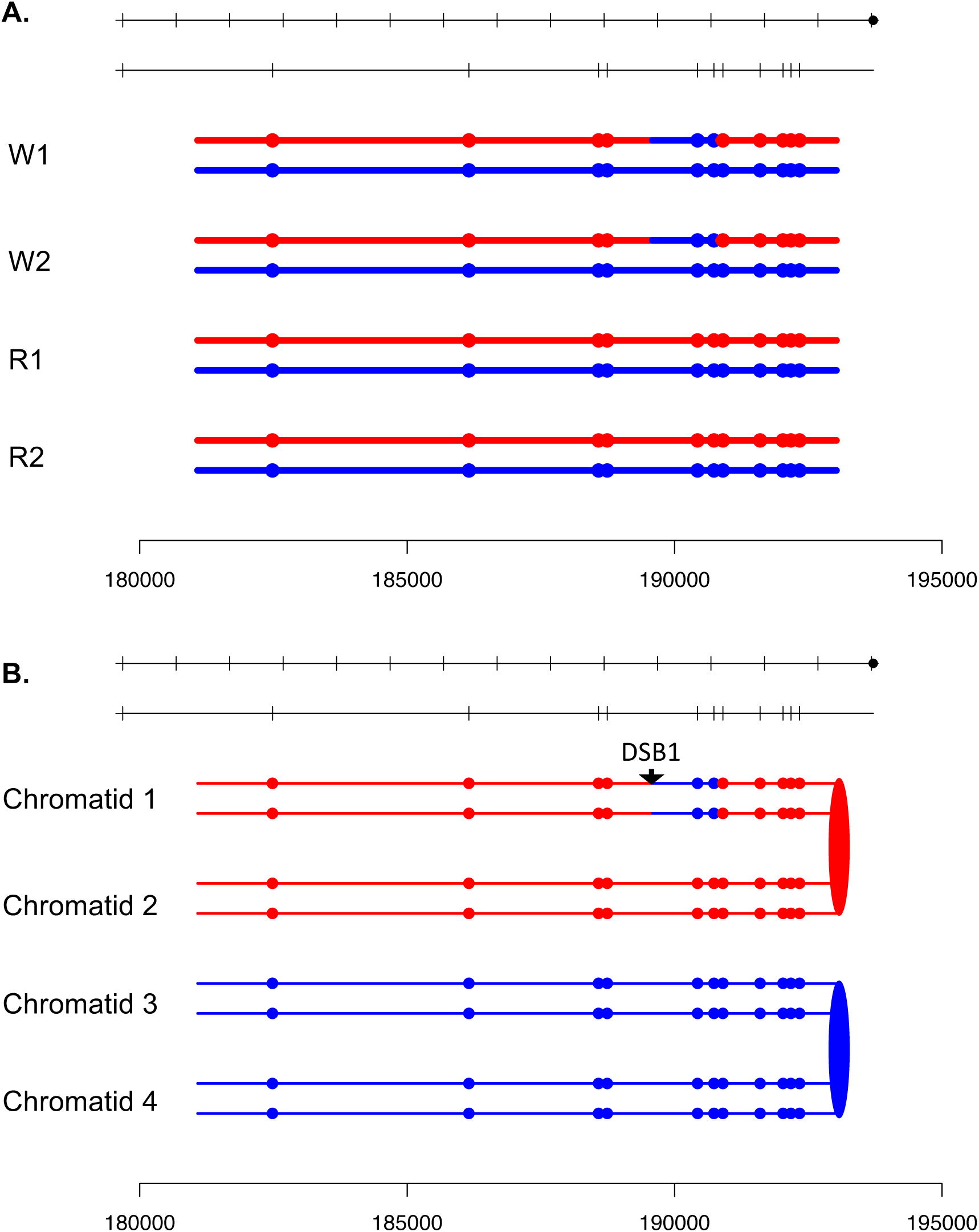

**Fig. S28.**
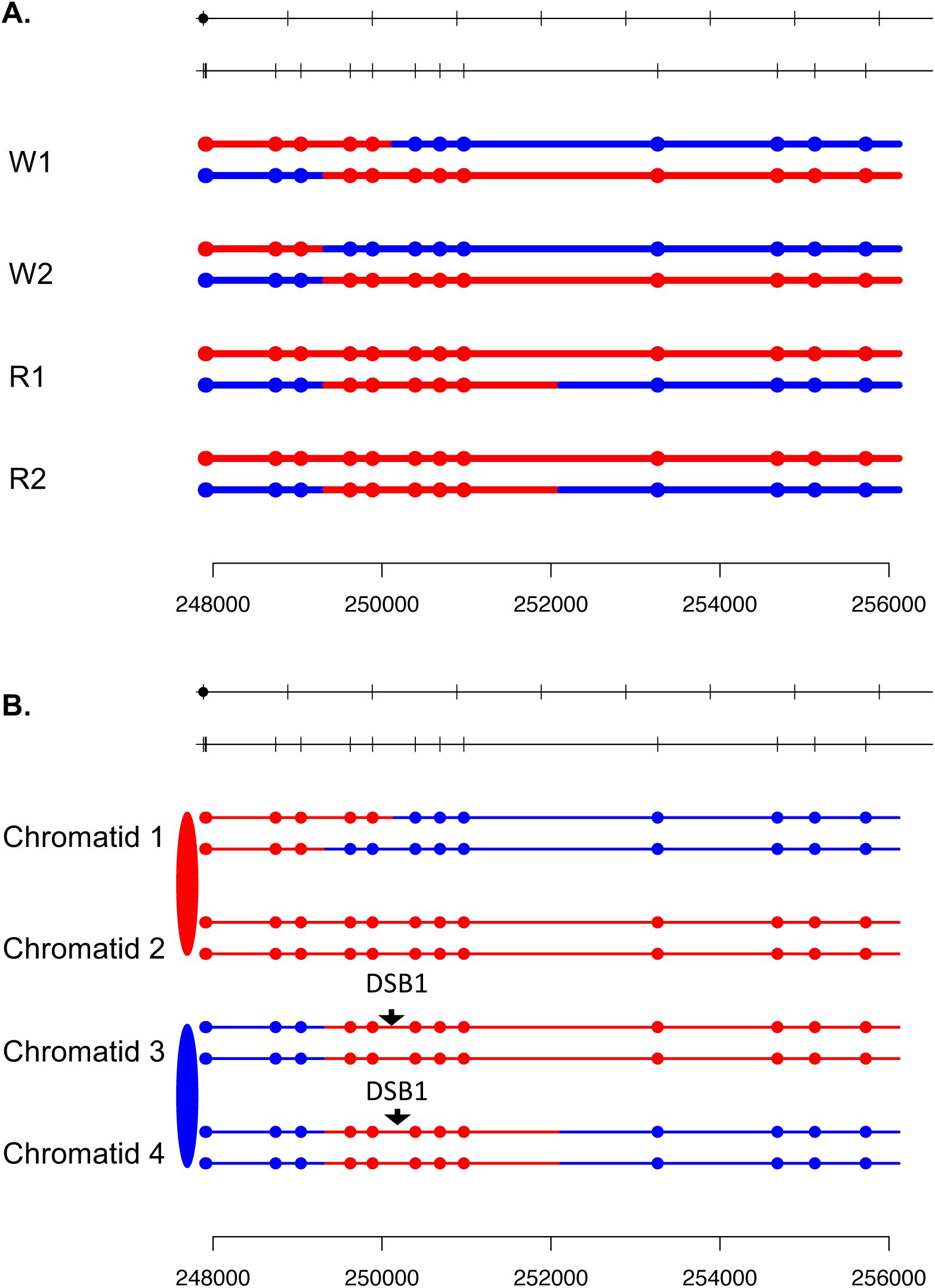

**Fig. S29.**
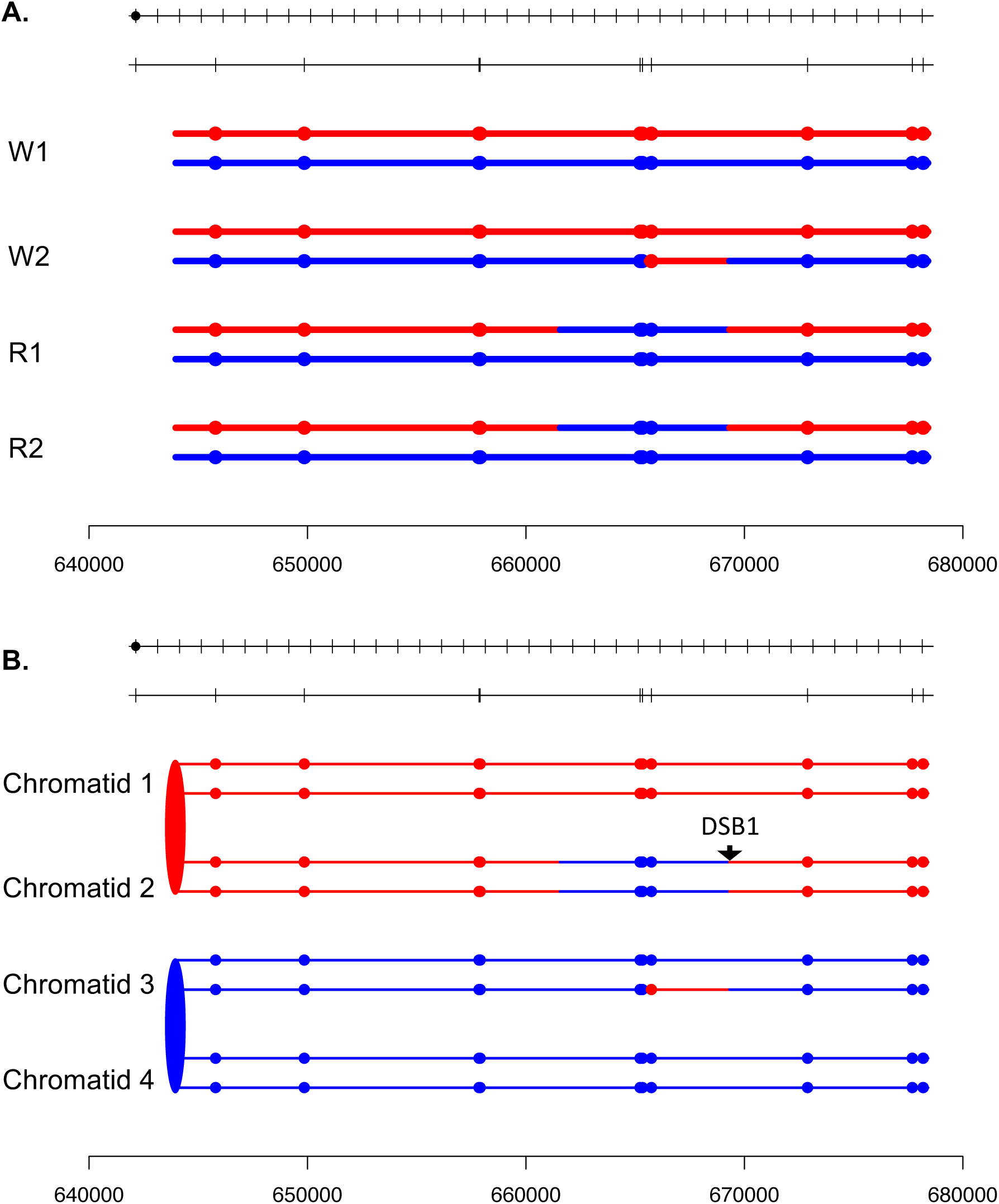

**Fig. S30.**
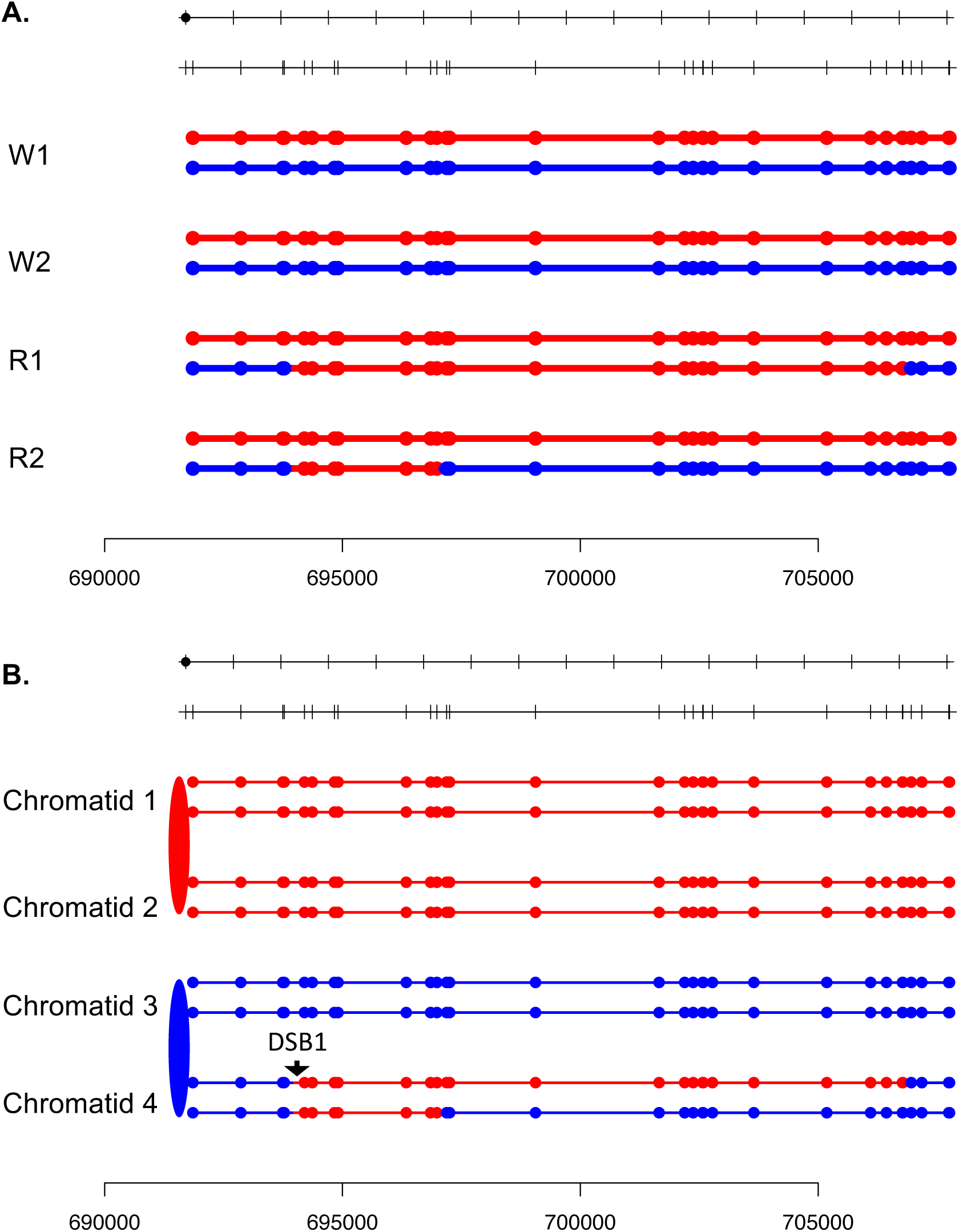

**Fig. S31.**
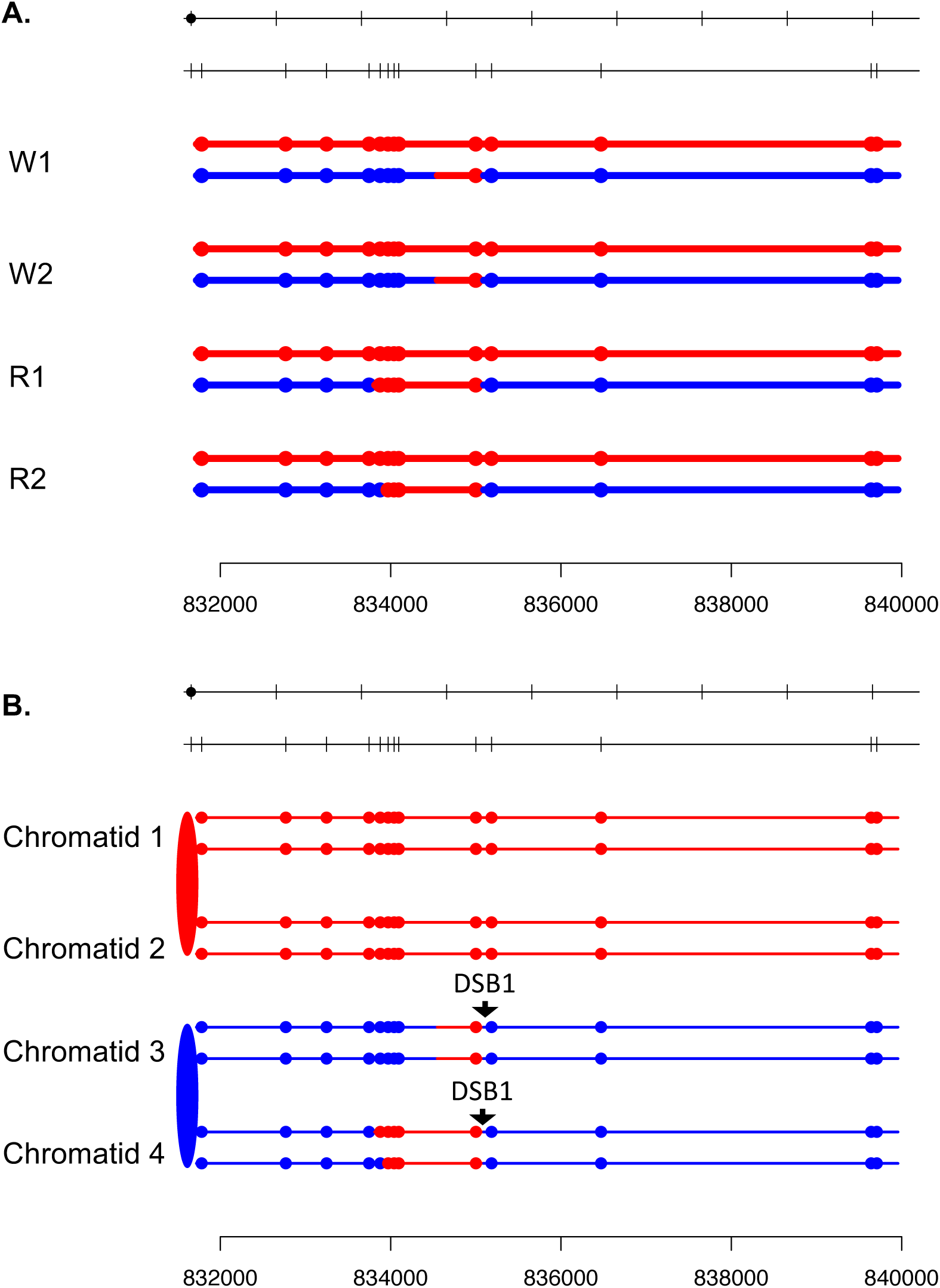

**Fig. S32.**
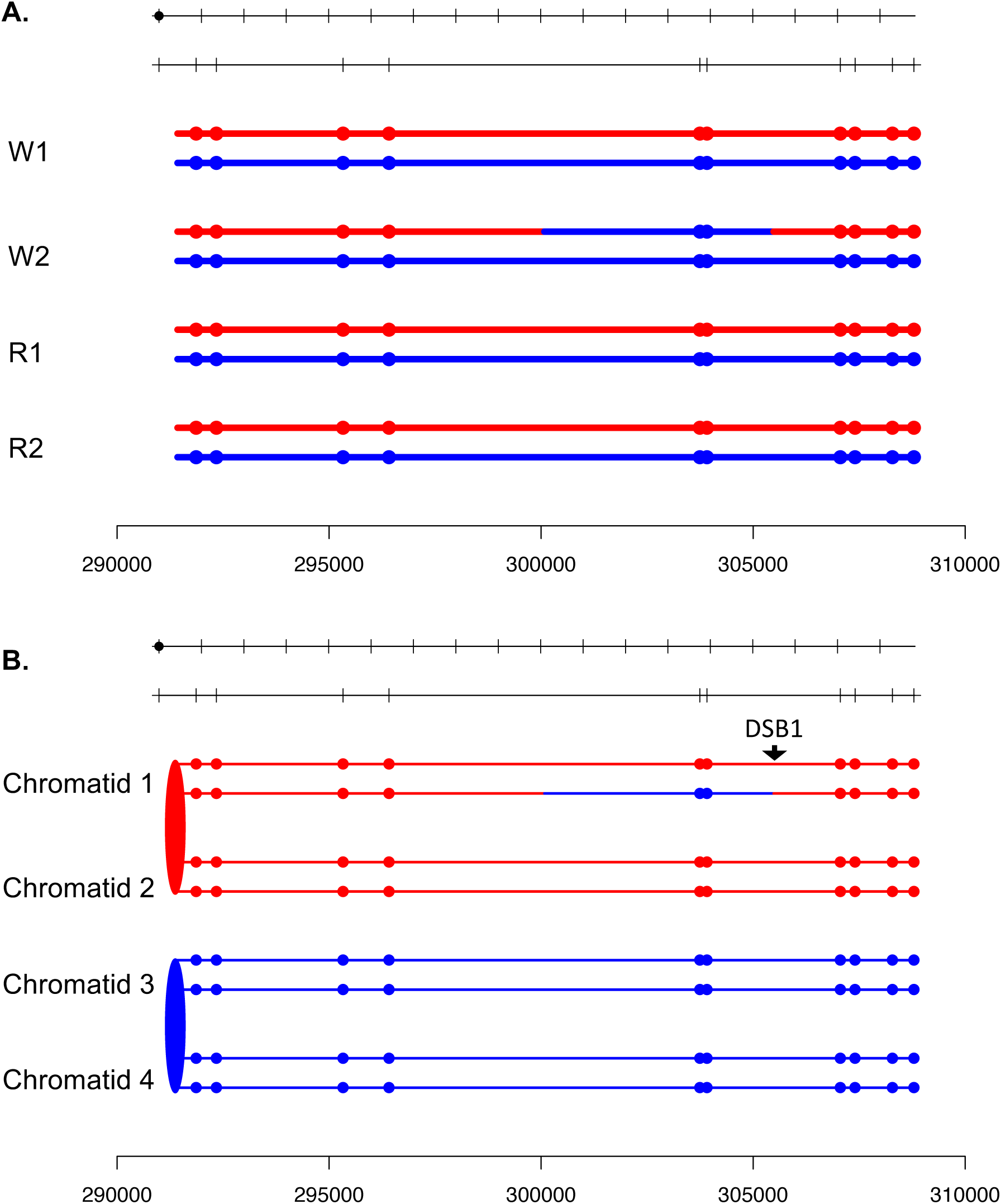

**Fig. S33.**
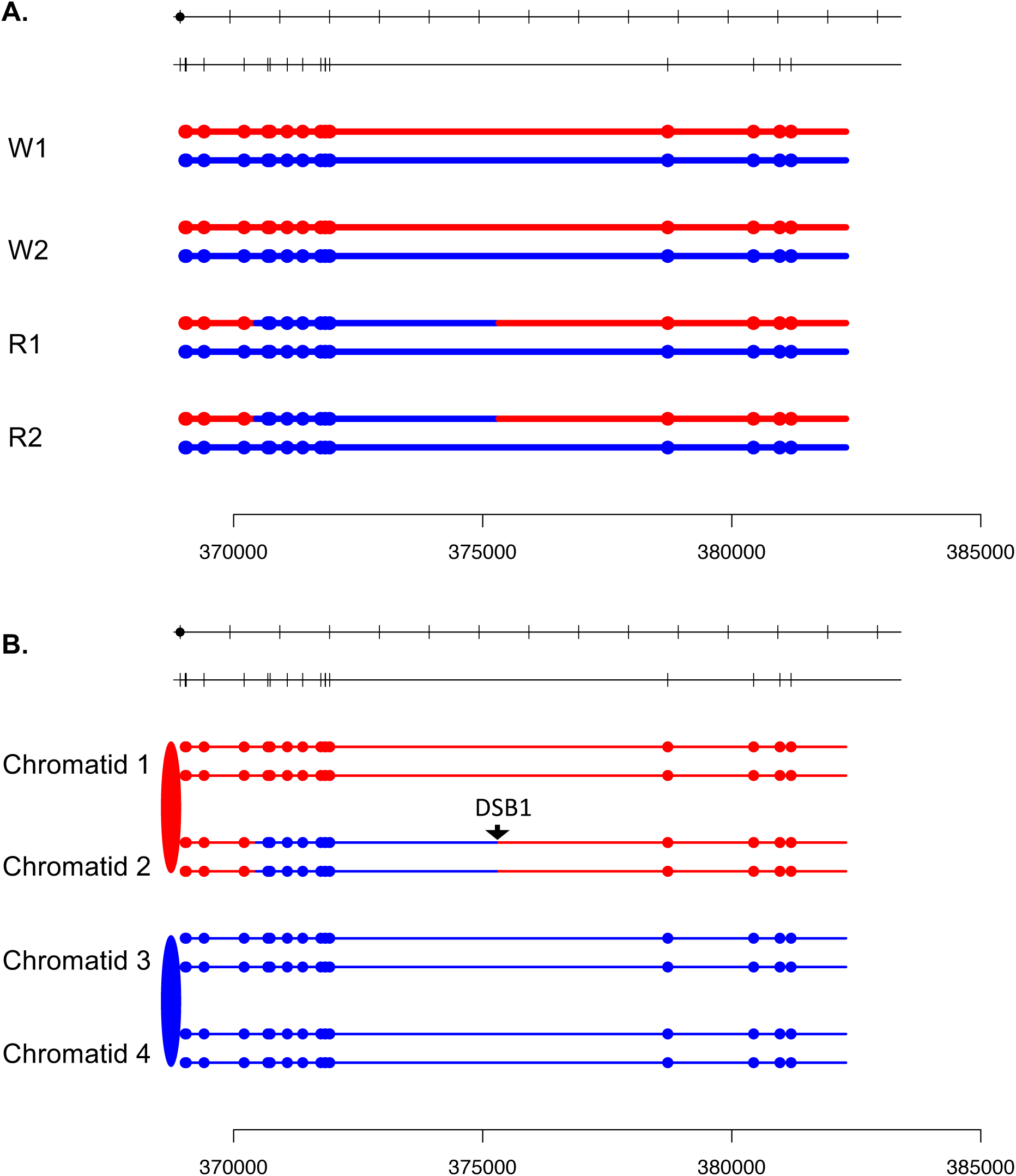

**Fig. S34.**
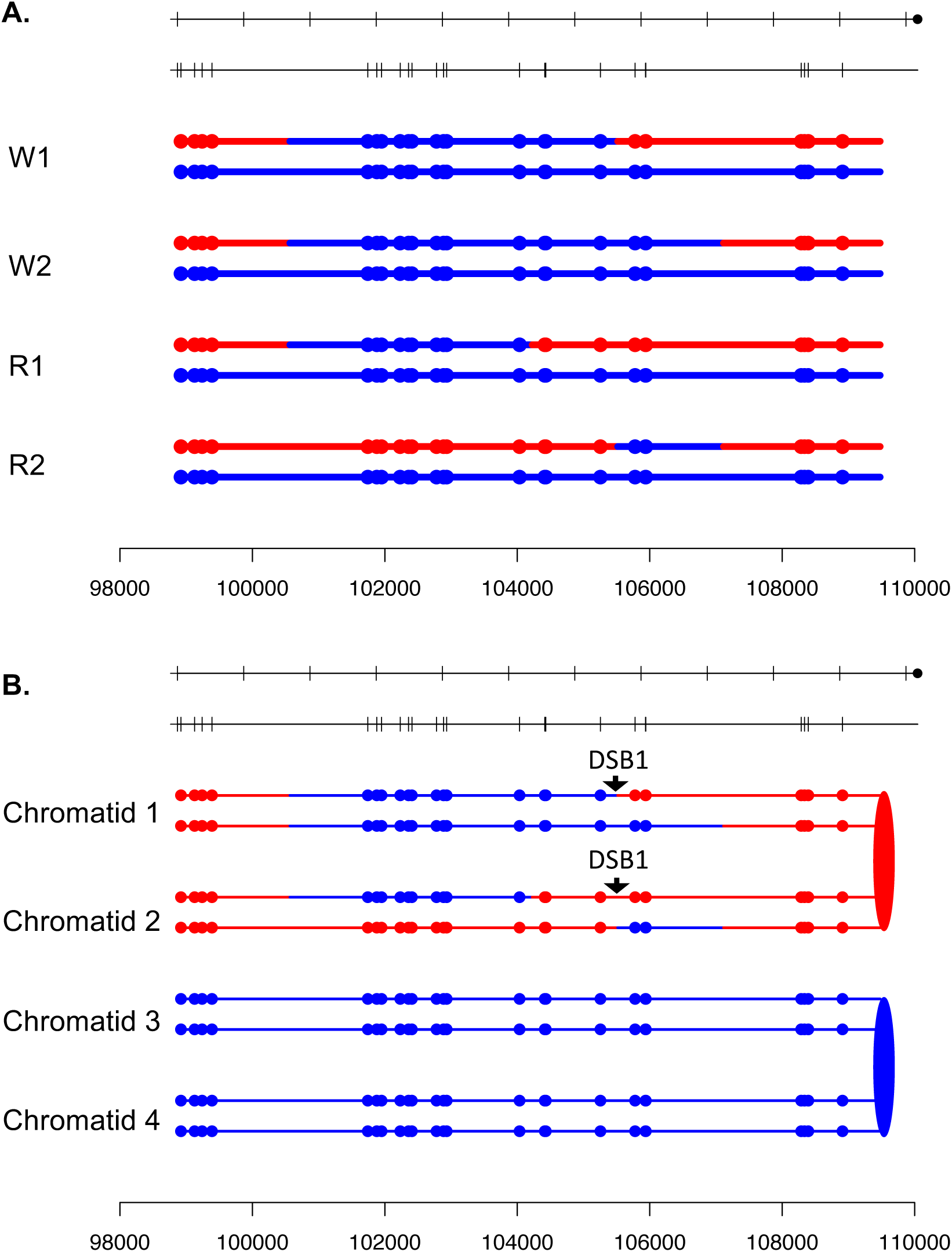

**Fig. S35.**
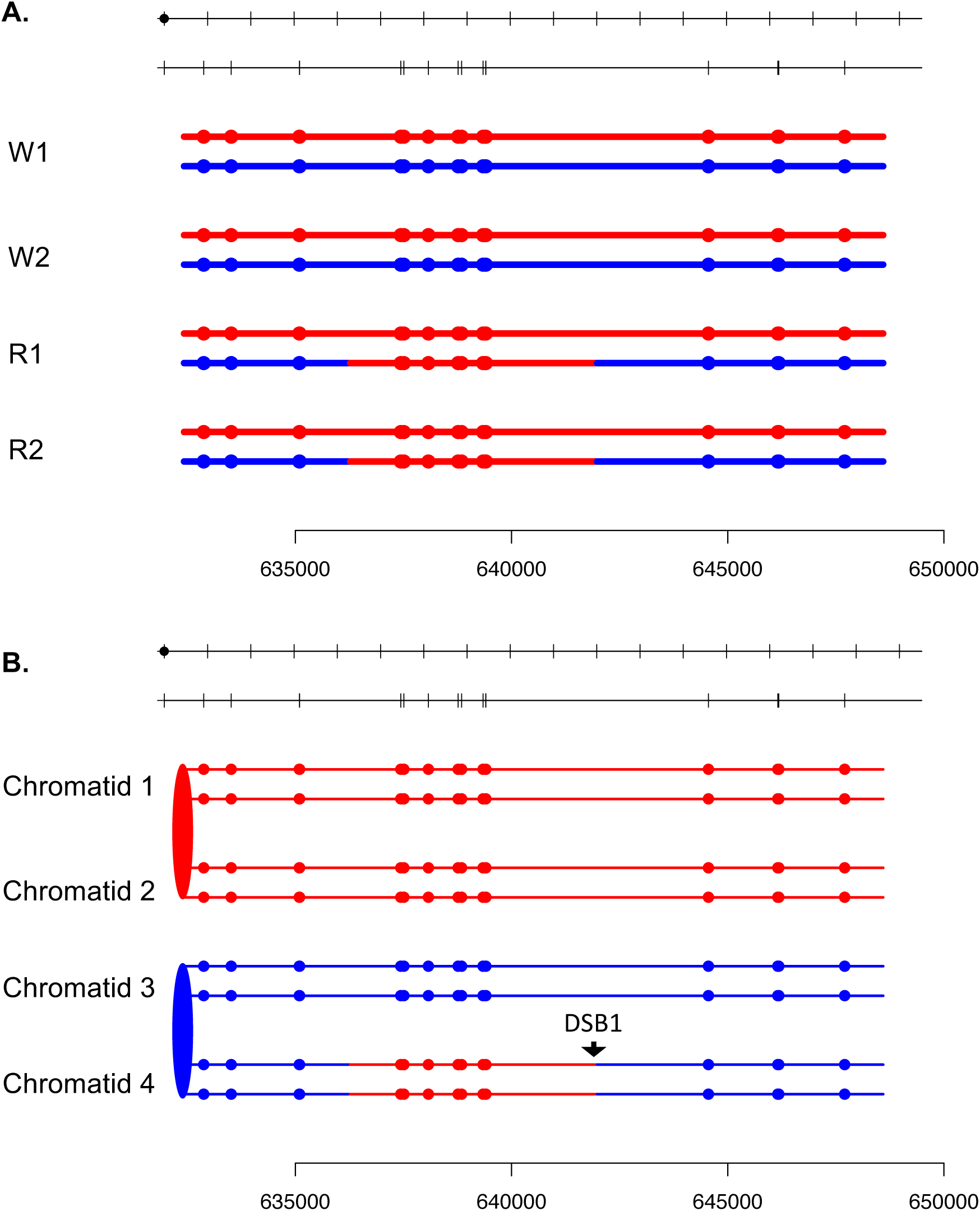

**Fig. S36.**
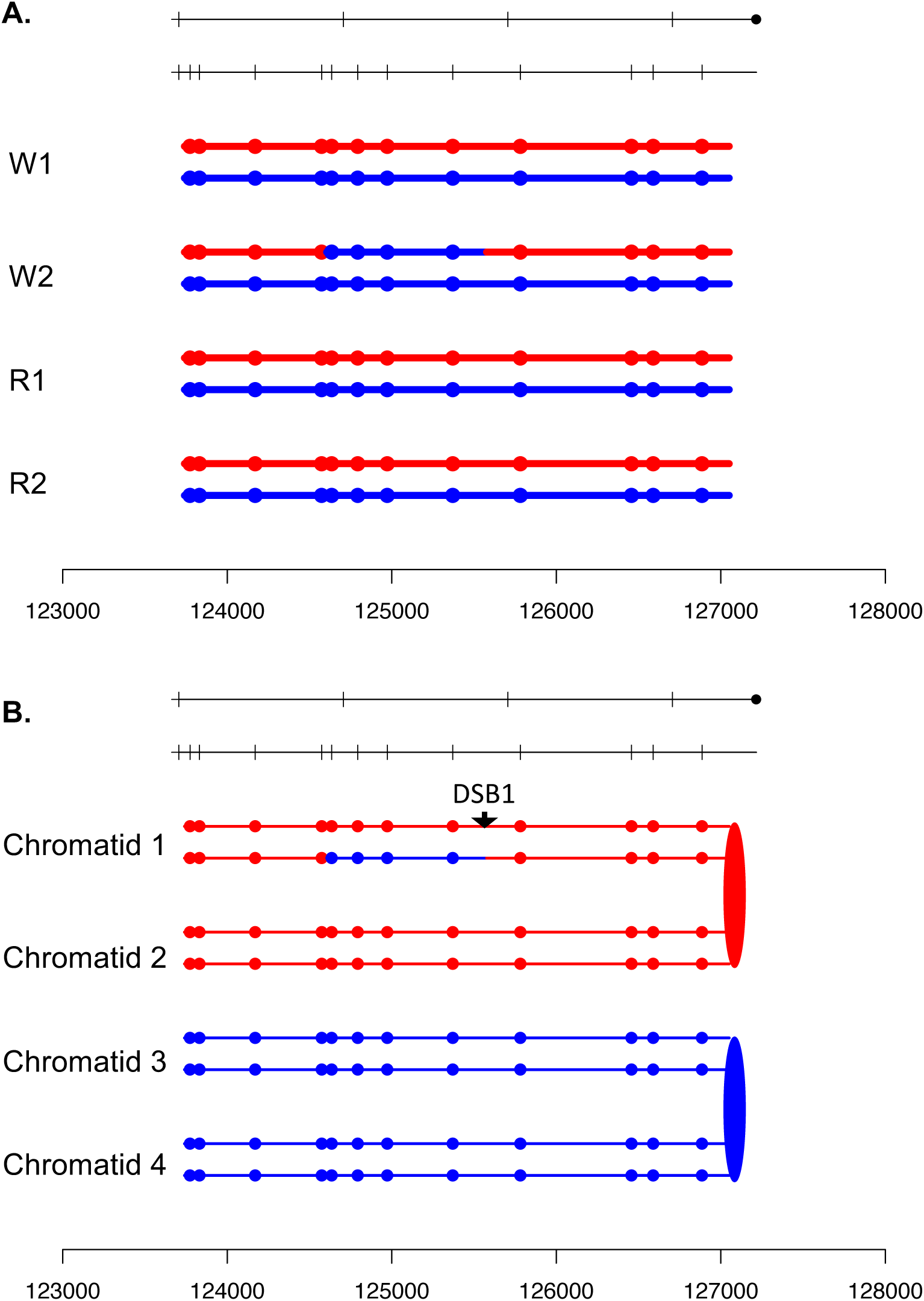

**Fig. S37.**
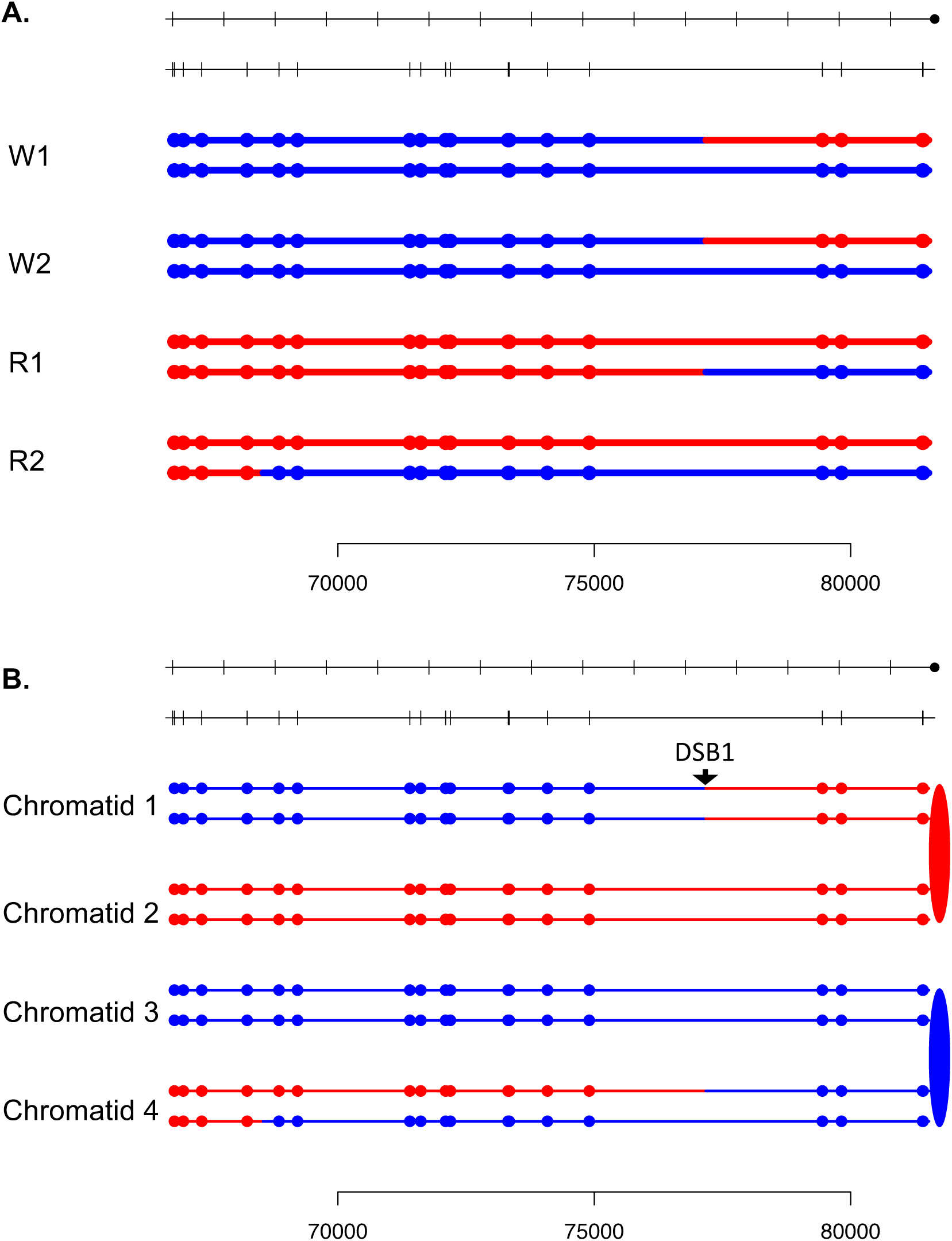

**Fig. S38.**
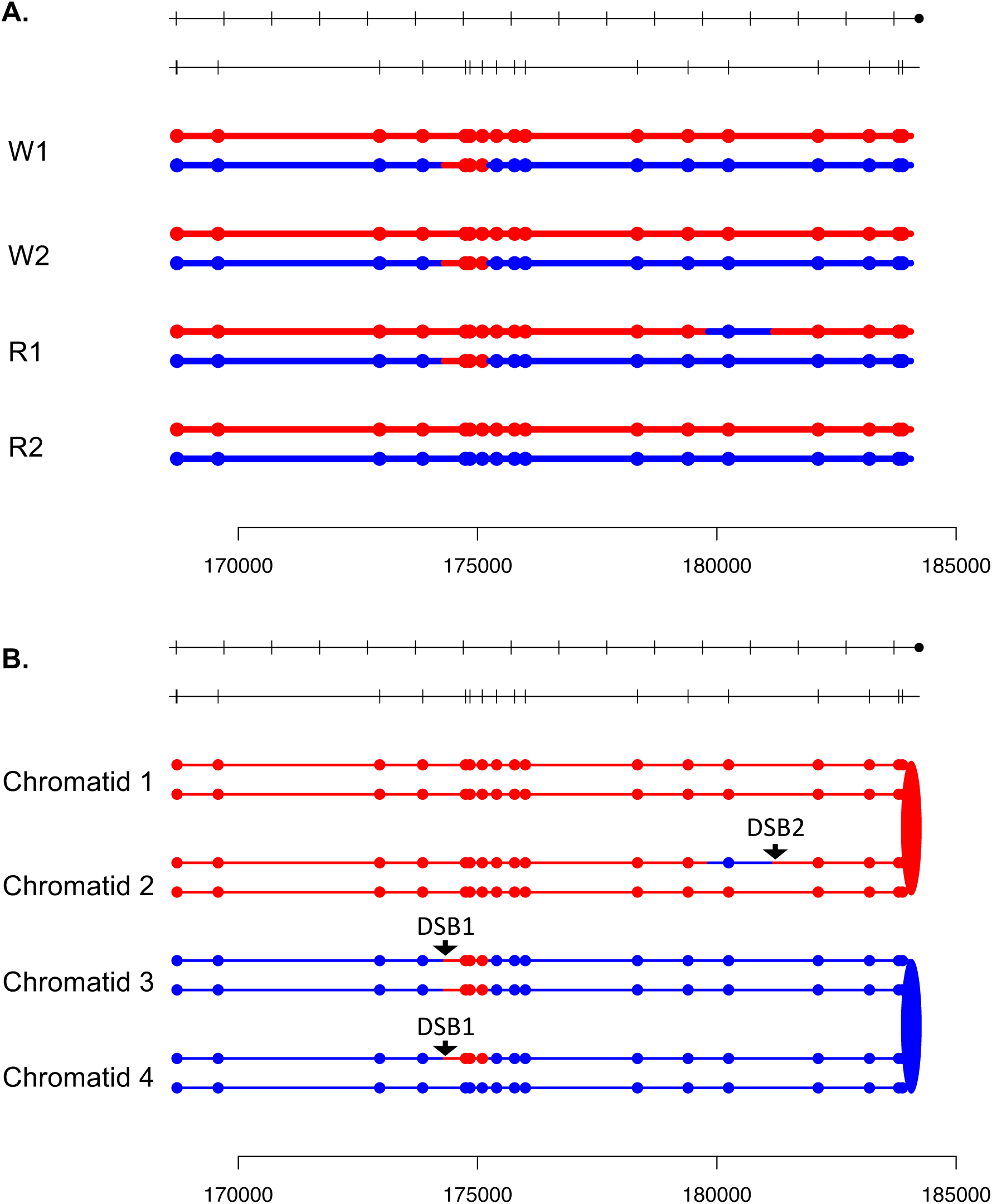

**Fig. S39.**
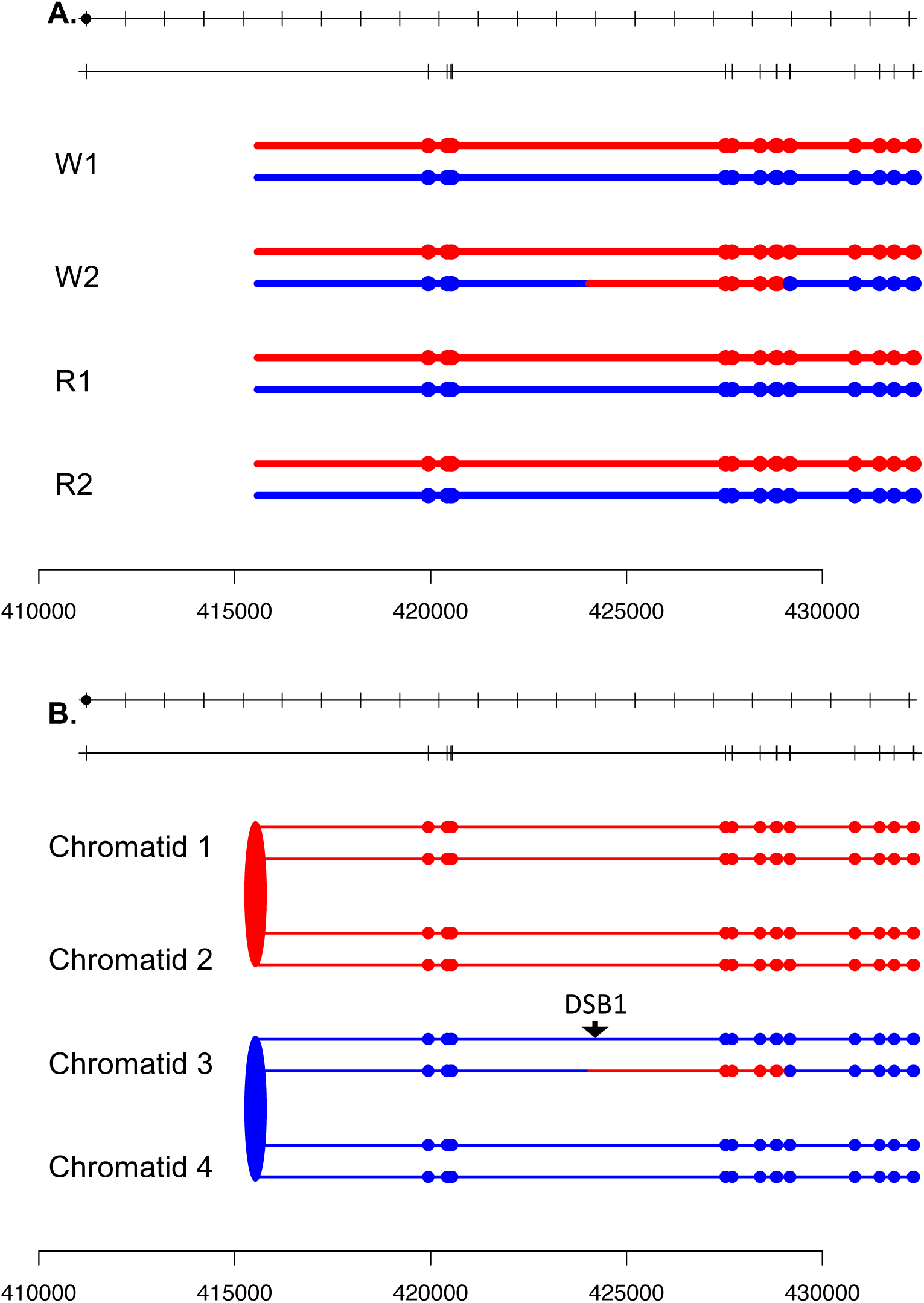

**Fig. S40.**
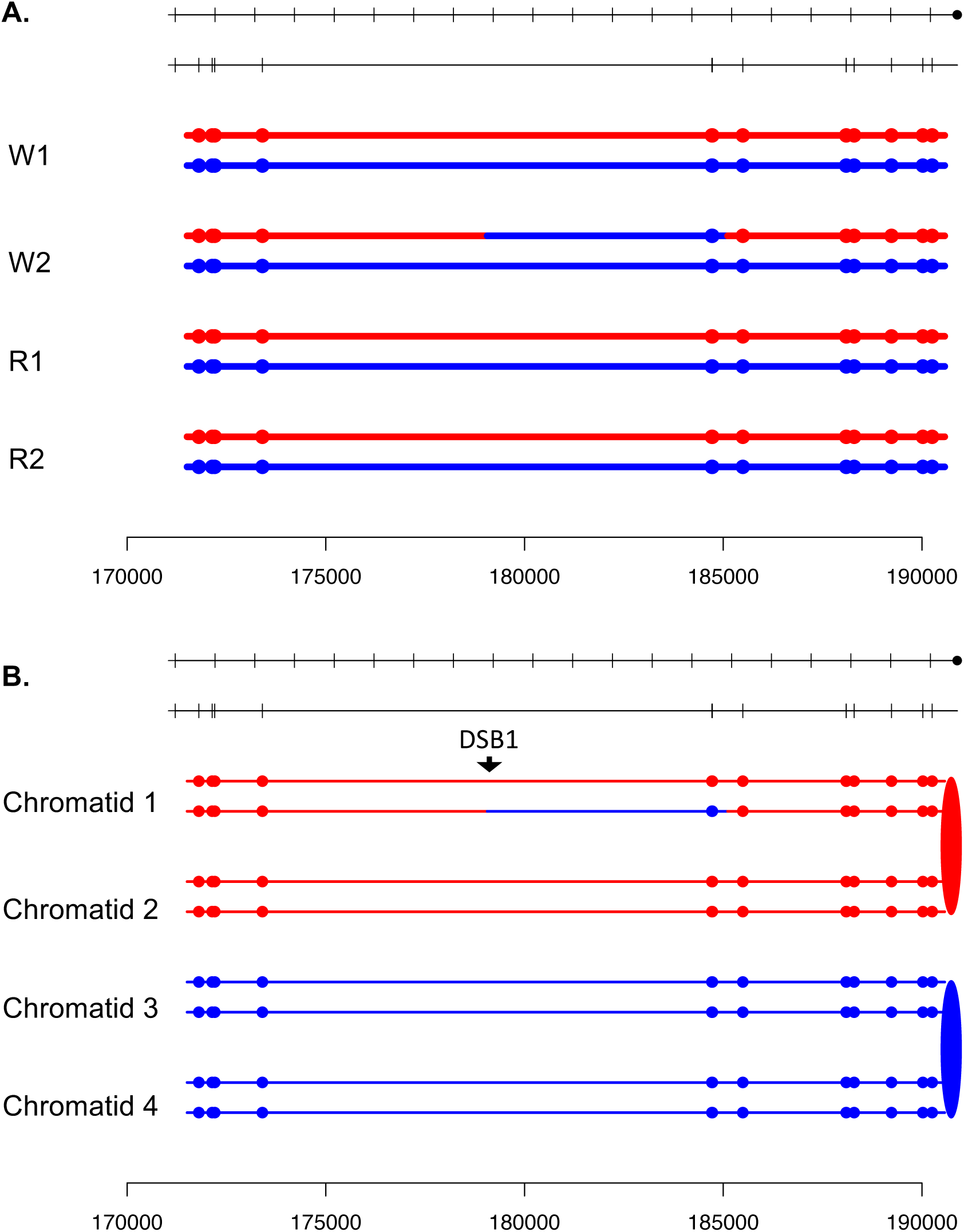

**Fig. S41.**
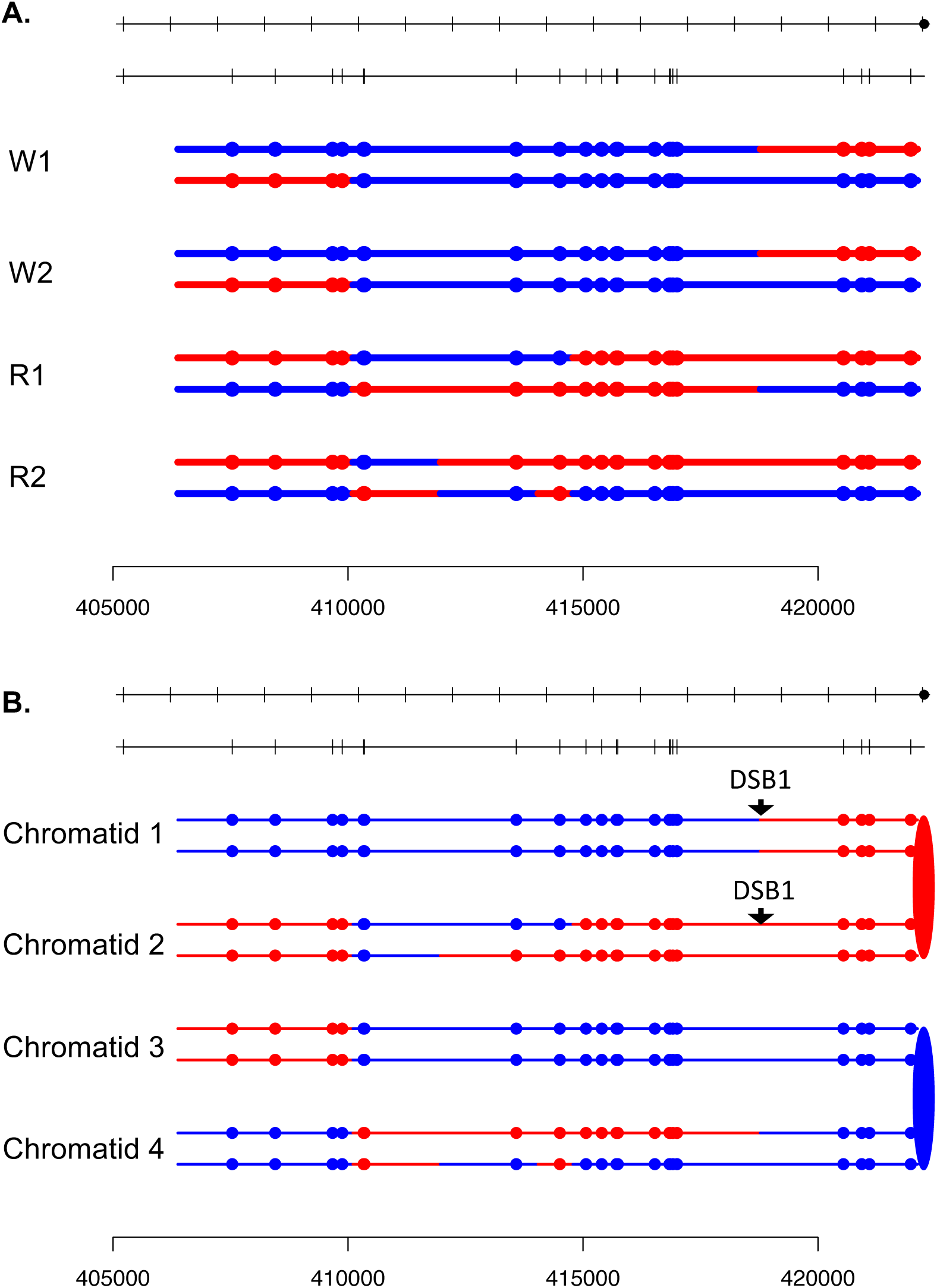

**Fig. S42.**
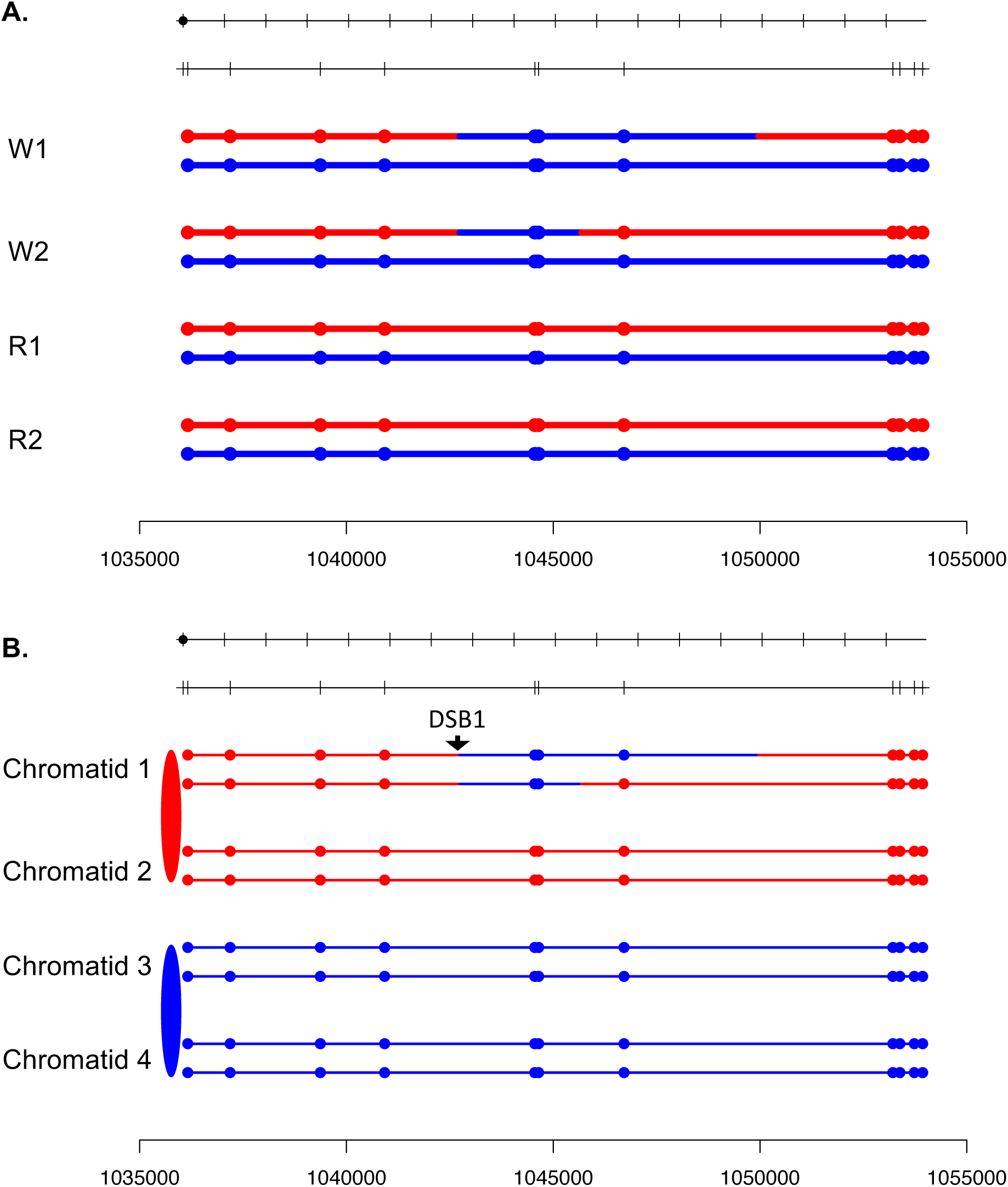

**Fig. S43.**
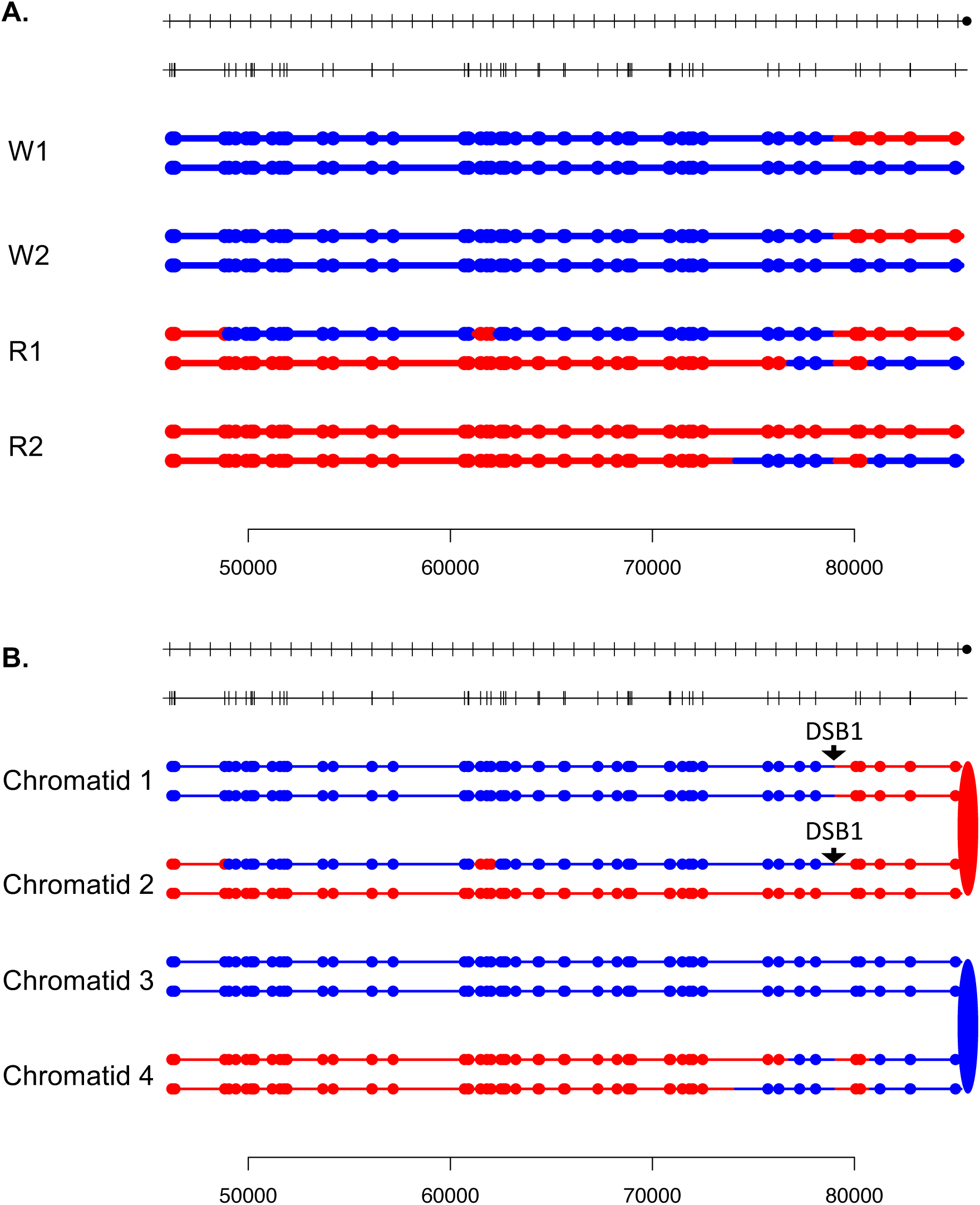

**Fig. S44.**
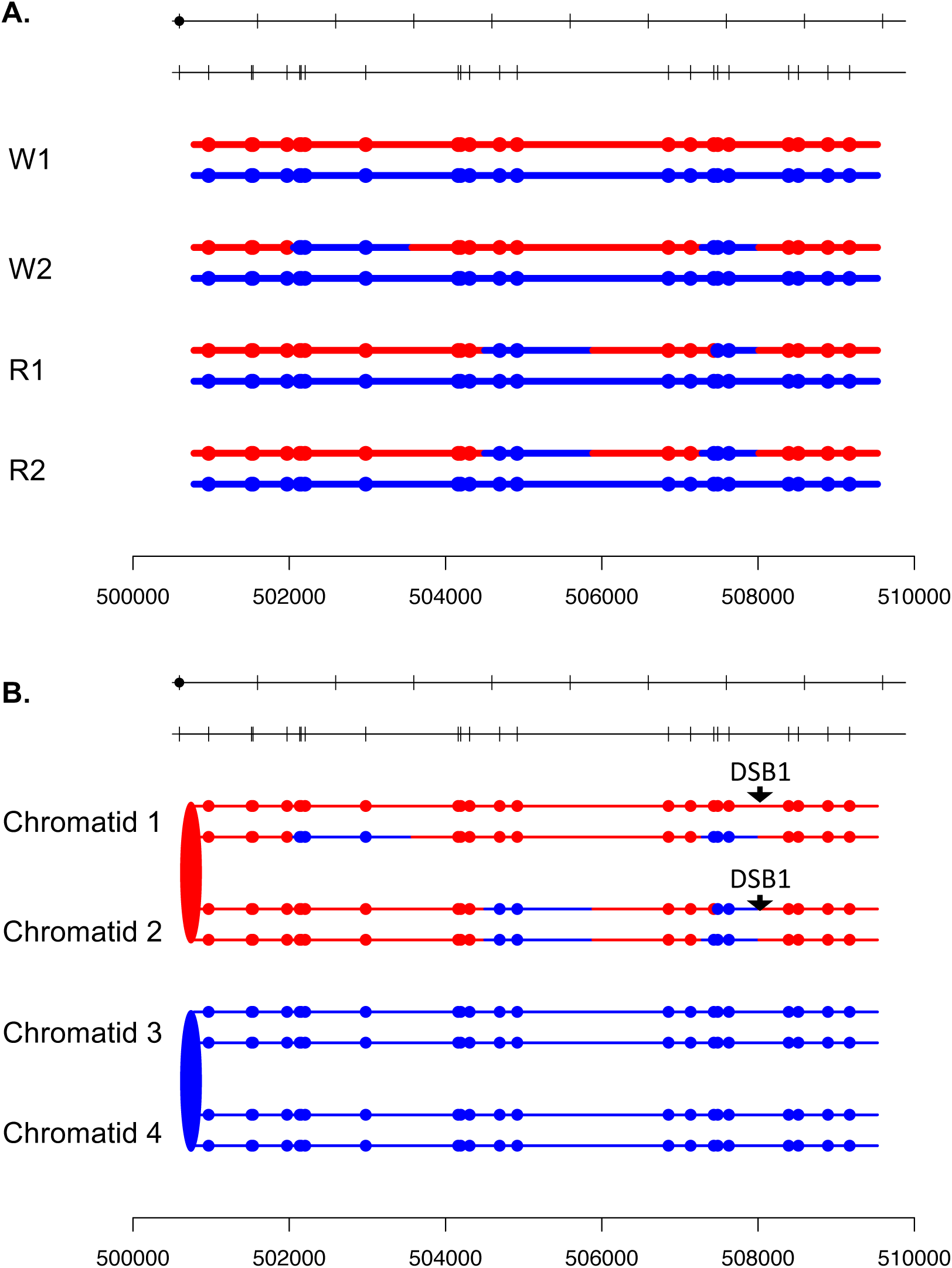

**Fig. S45.**
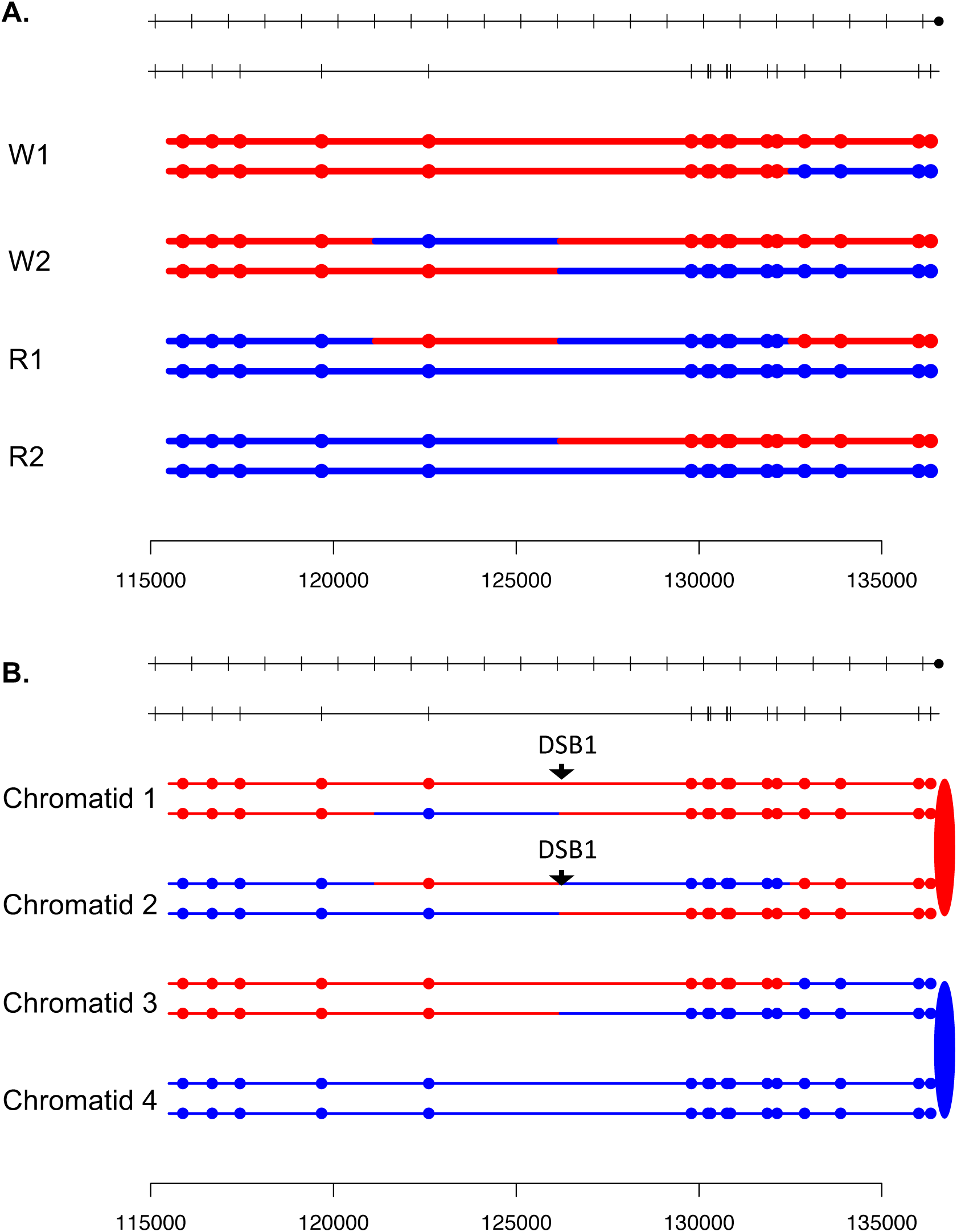

**Fig. S46.**
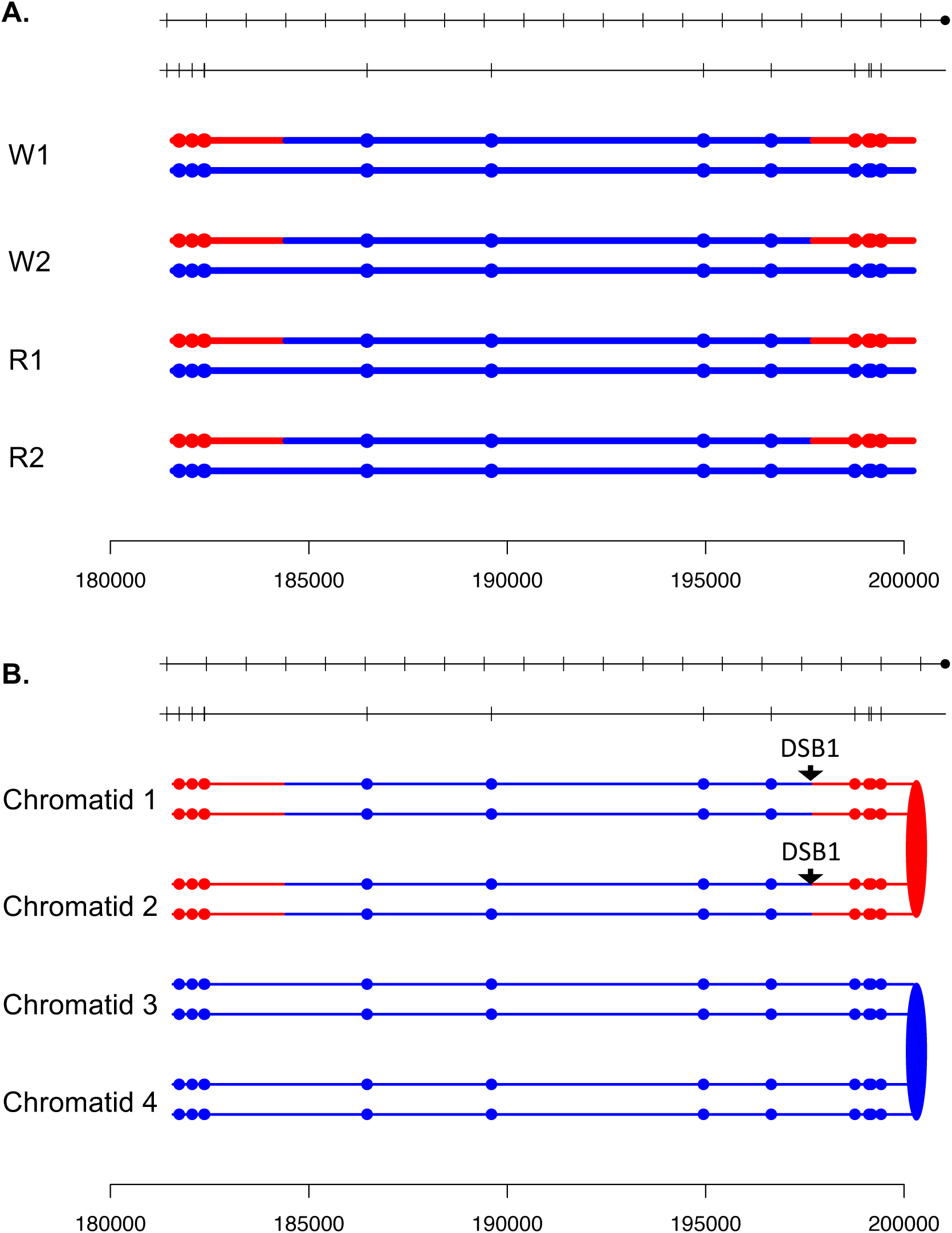

**Fig. S47.**
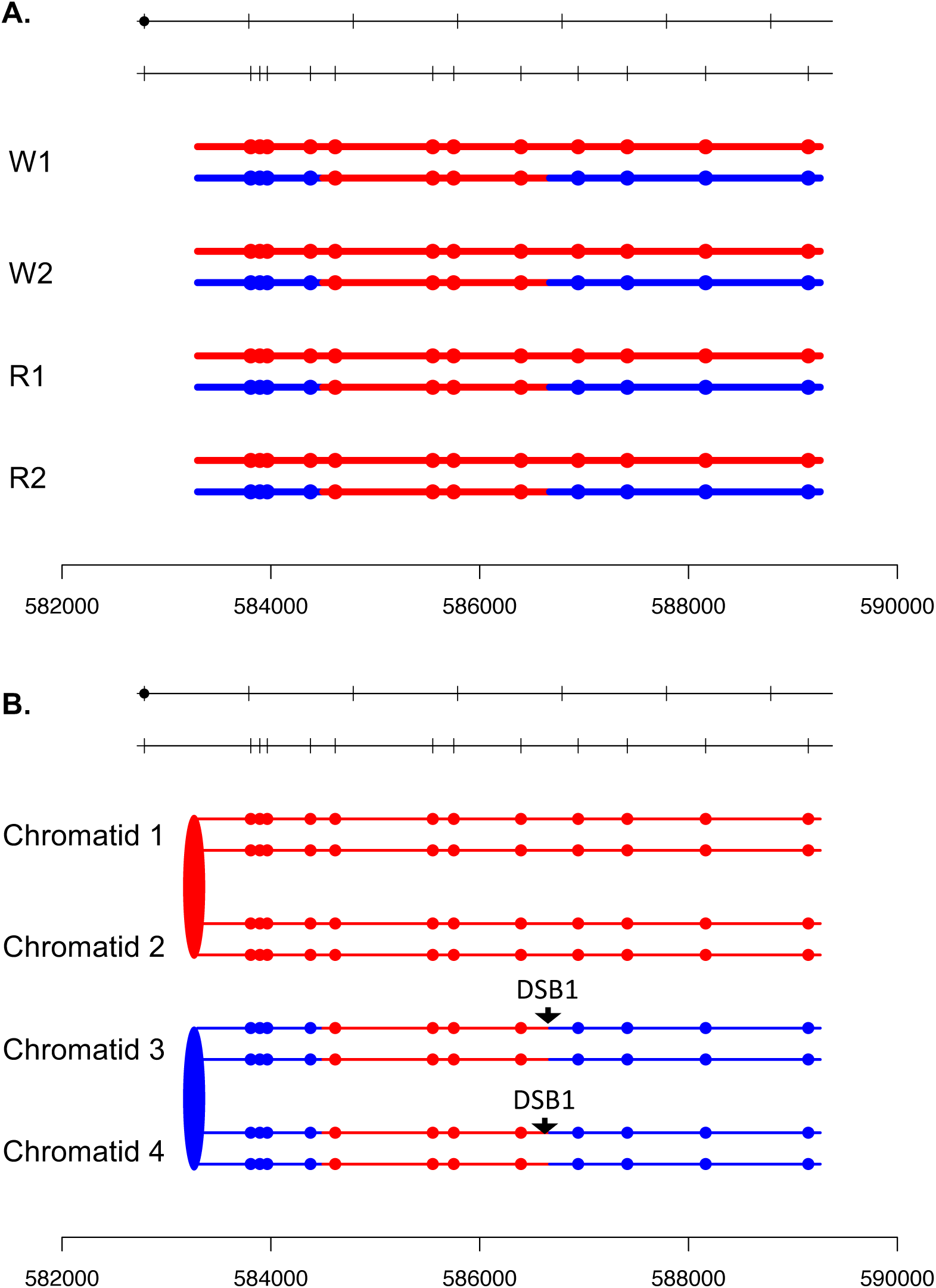

**Fig. S48.**
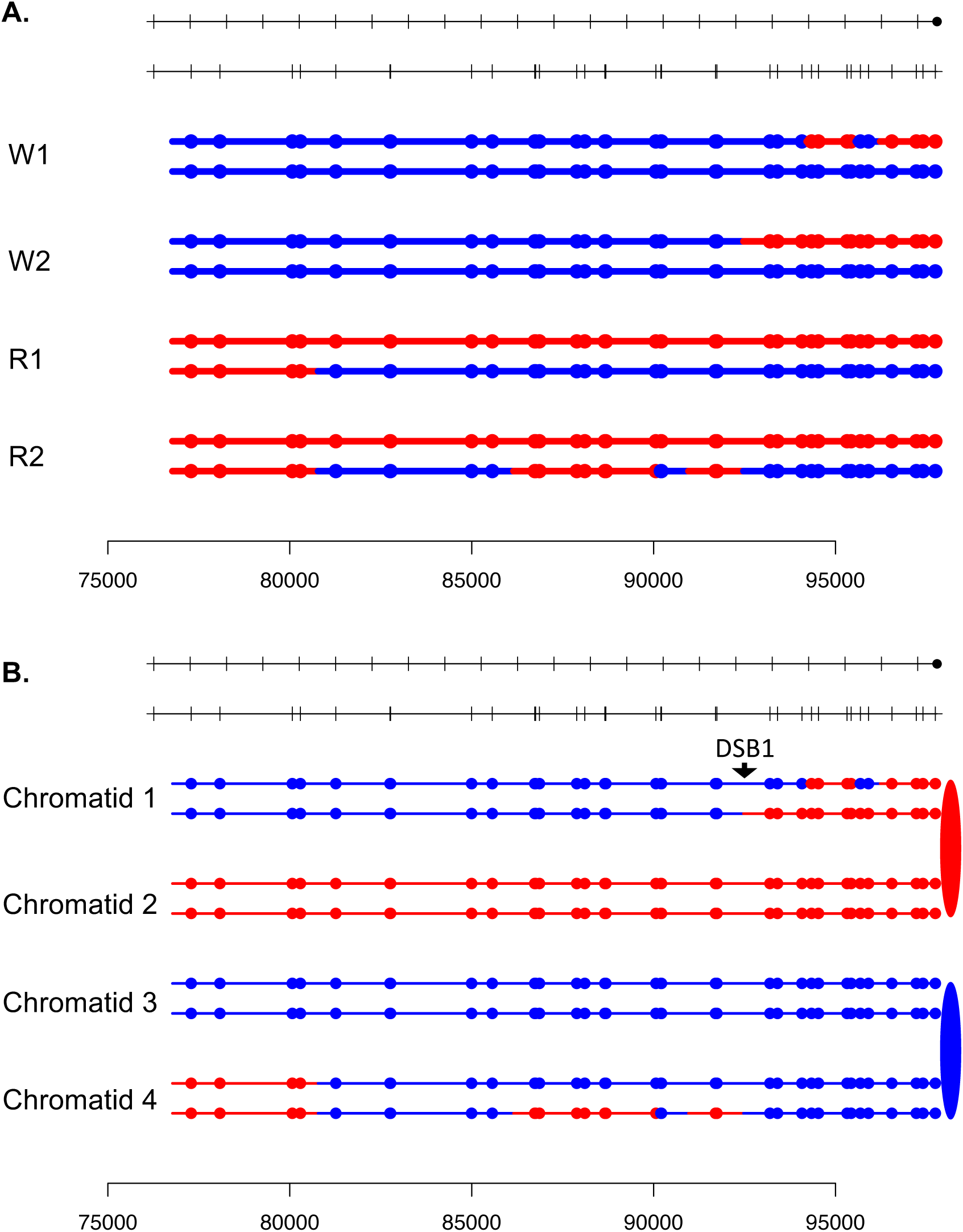

**Fig. S49.**
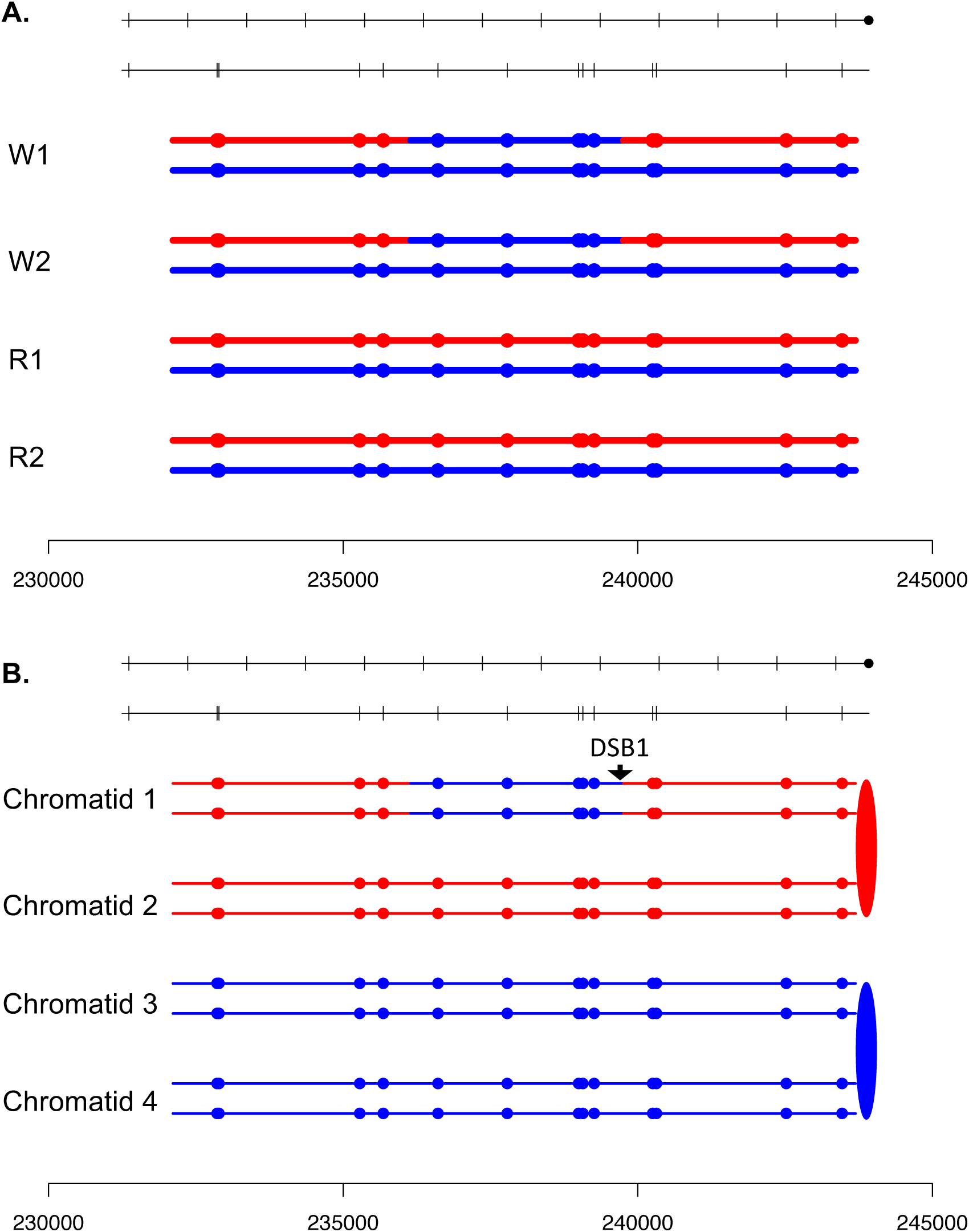

**Fig. S50.**
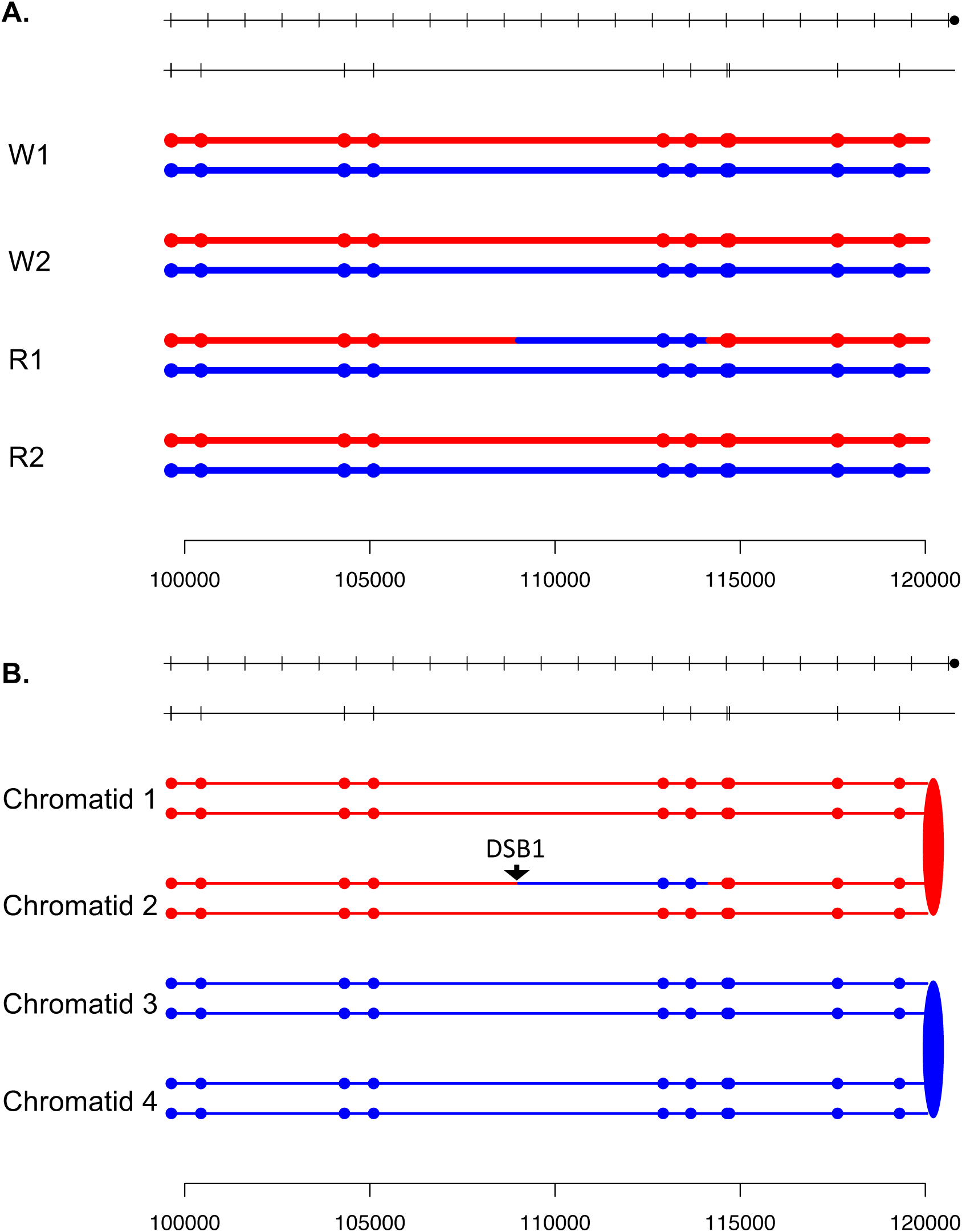

**Fig. S51.**
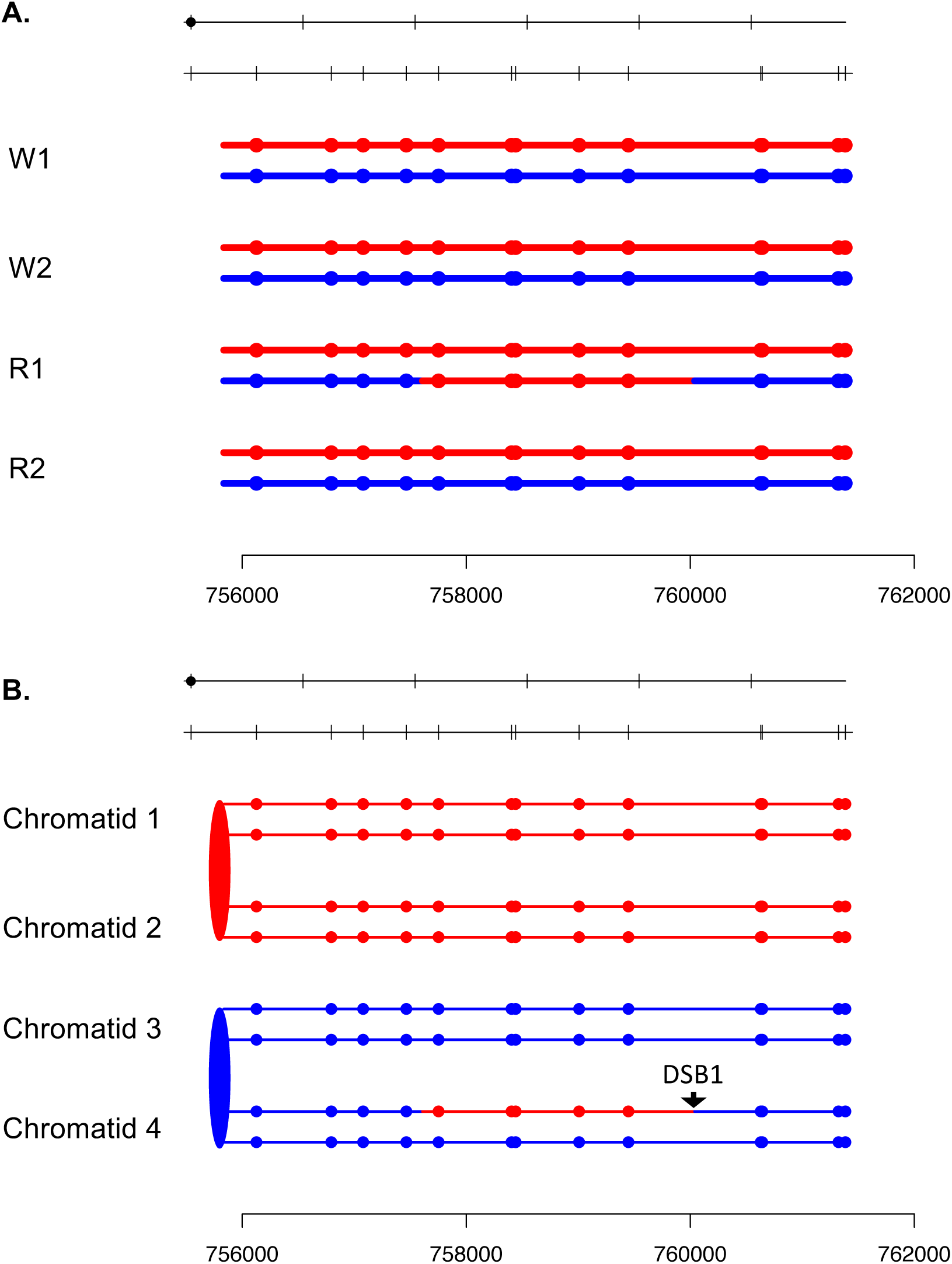

**Fig. S52.**
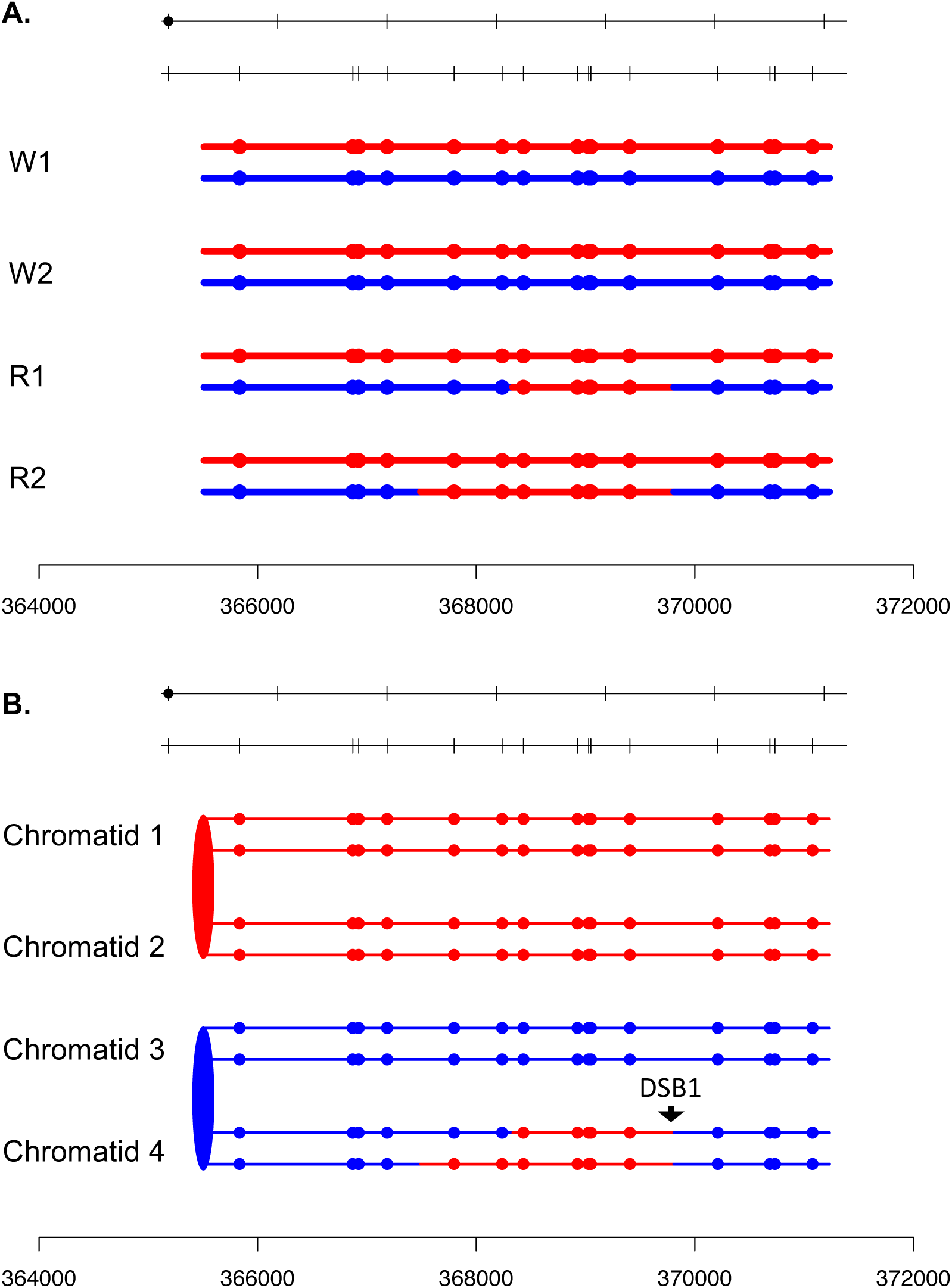

**Fig. S53.**
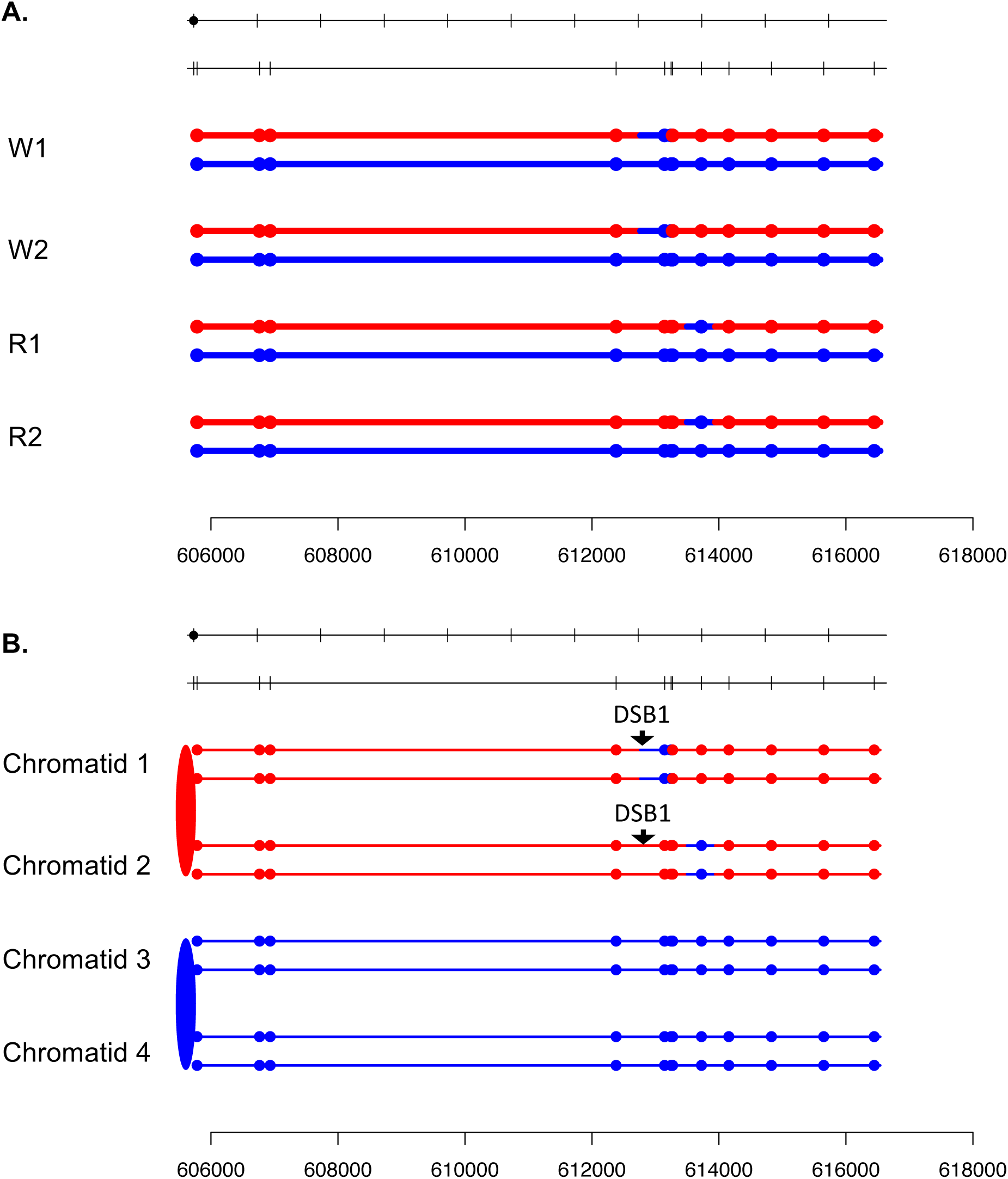

**Fig. S54.**
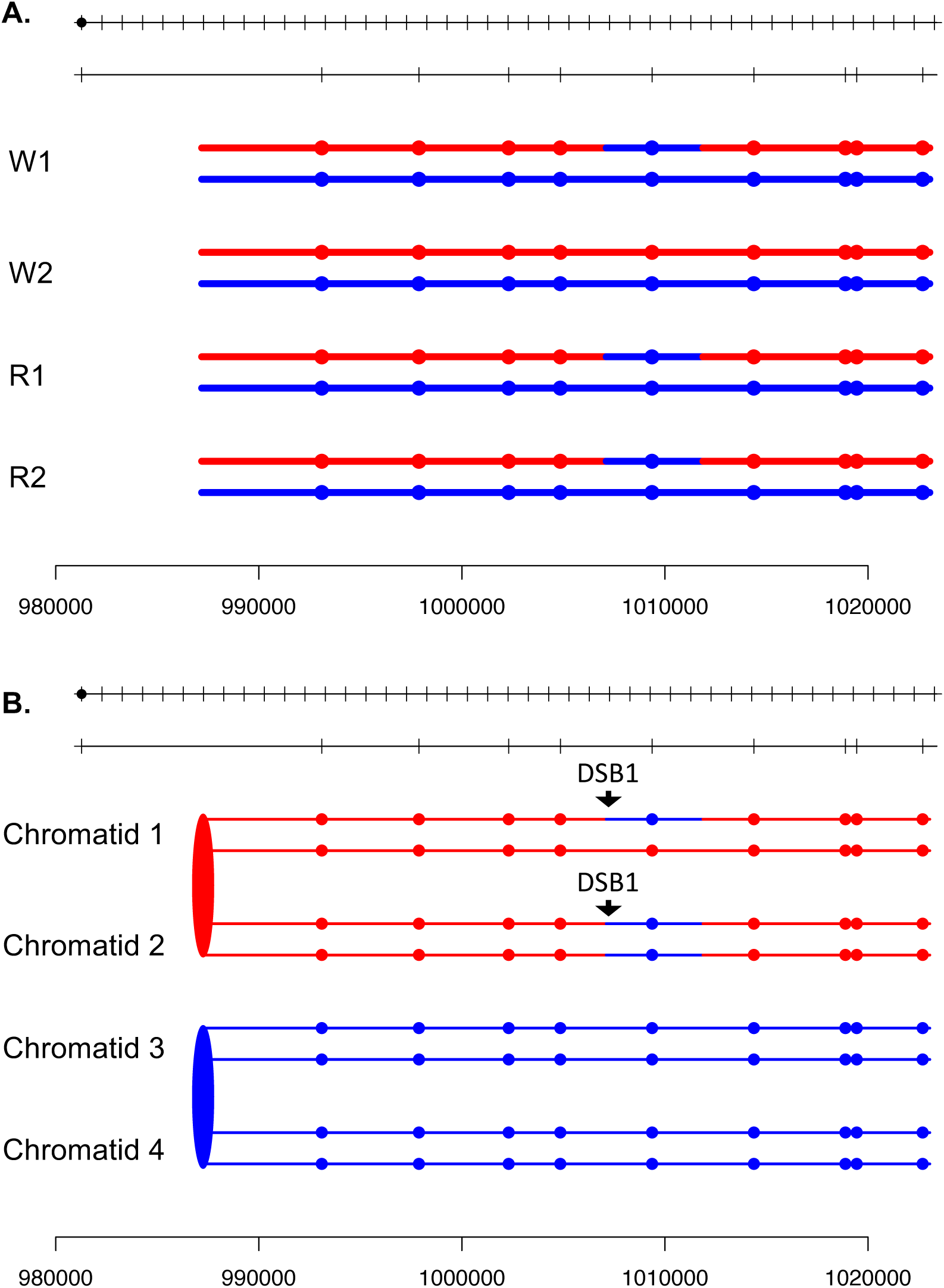

**Fig. S55.**
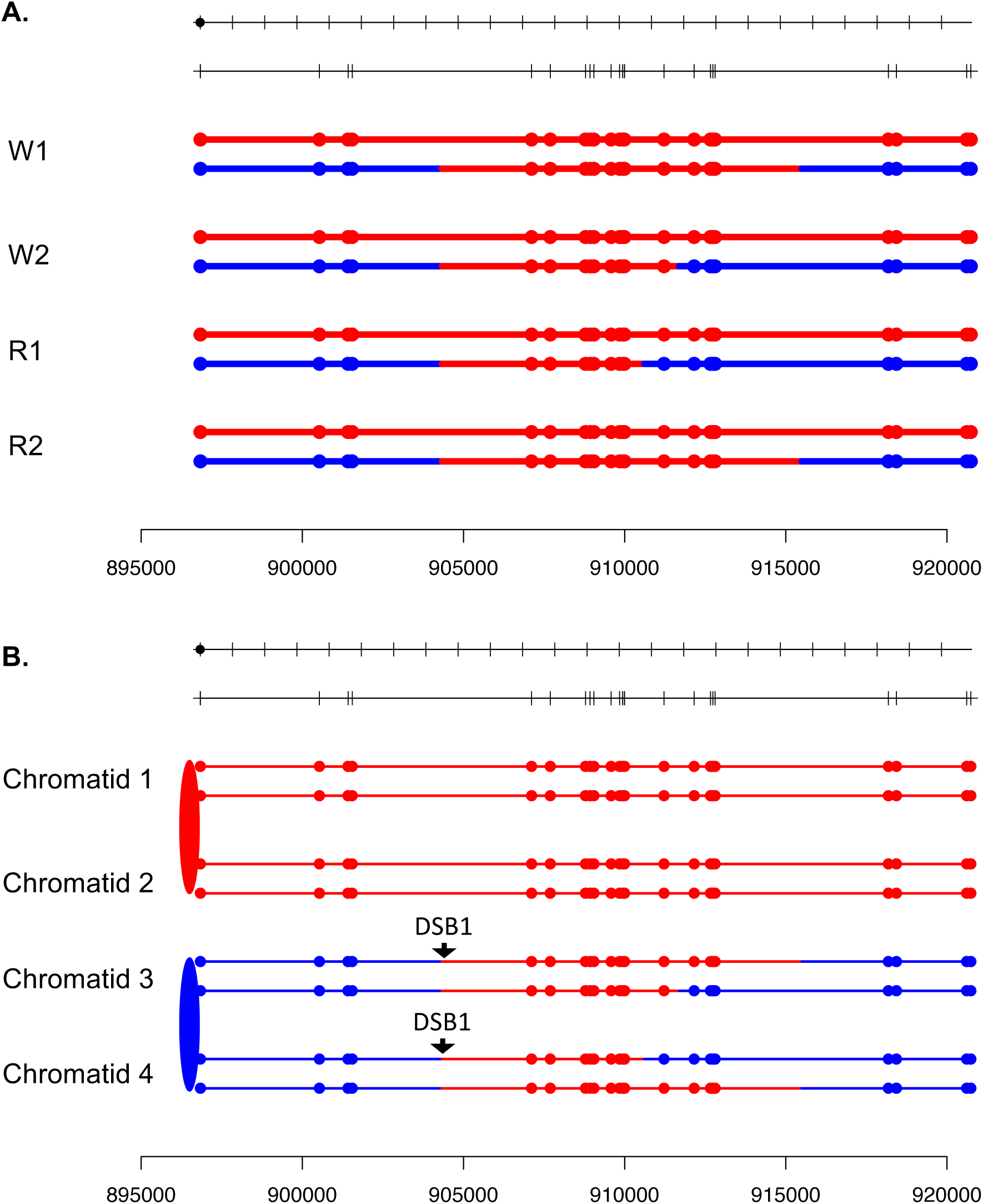

**Fig. S56.**
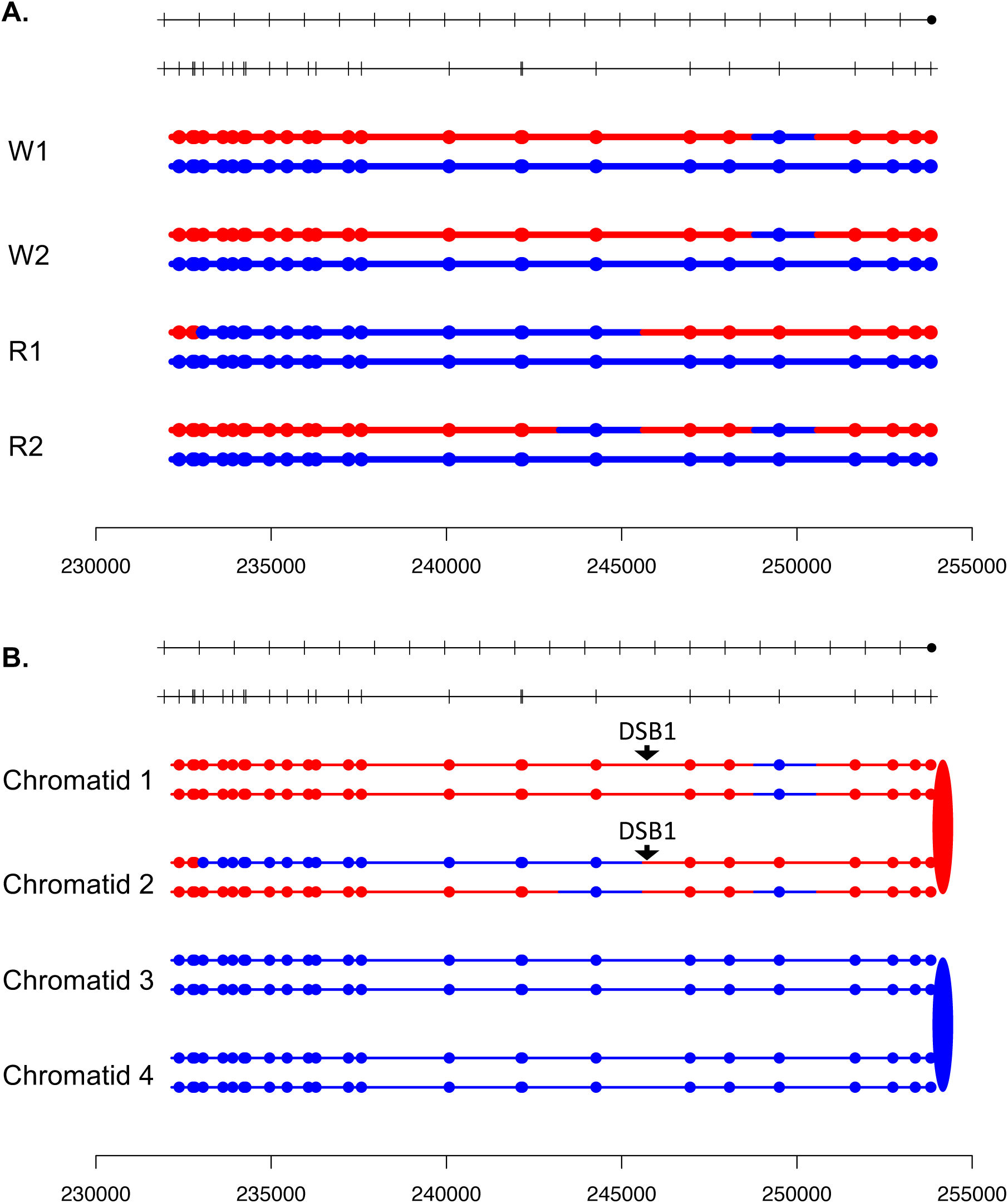

**Fig. S57.**
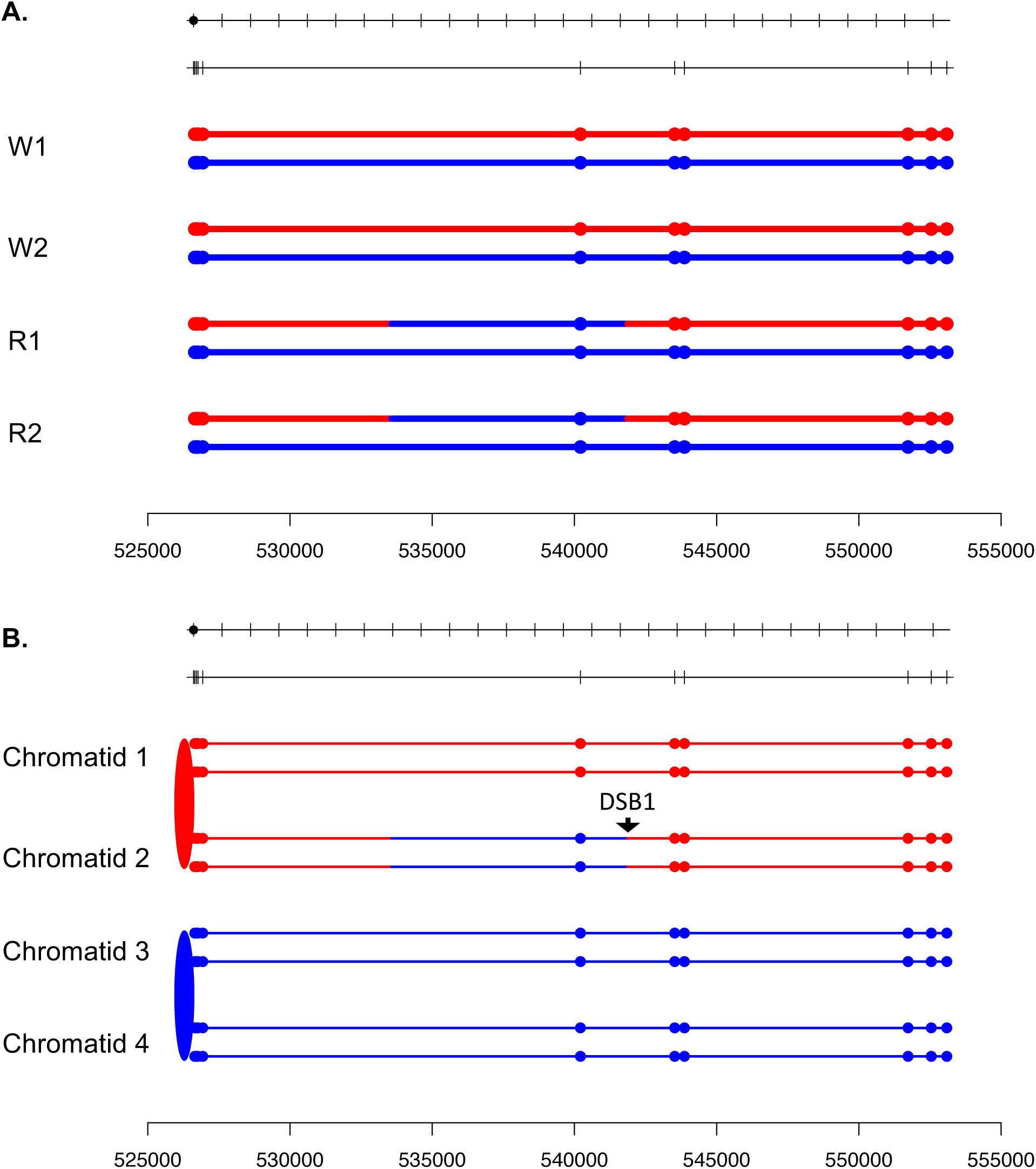

**Fig. S58.**
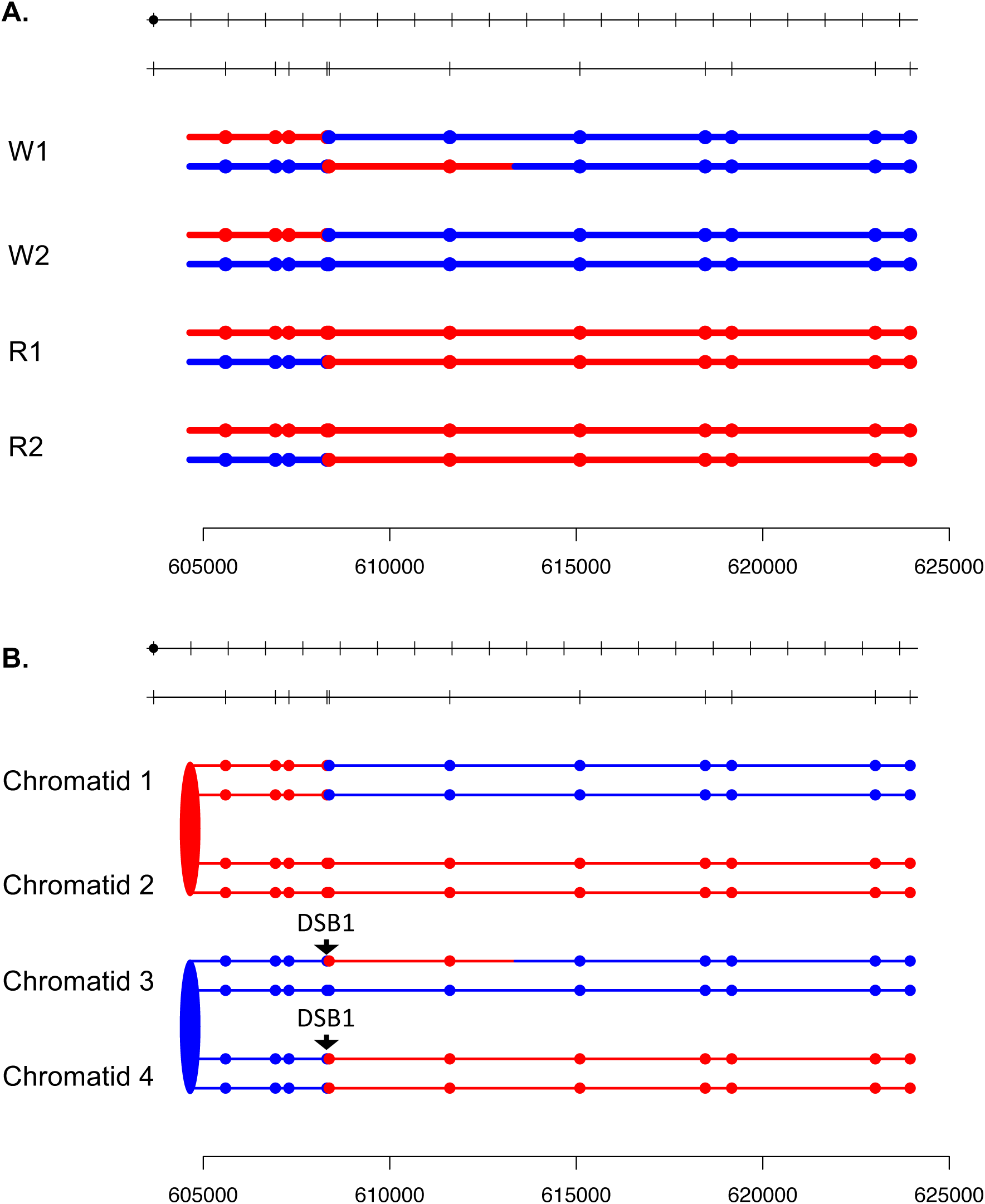

**Fig. S59.**
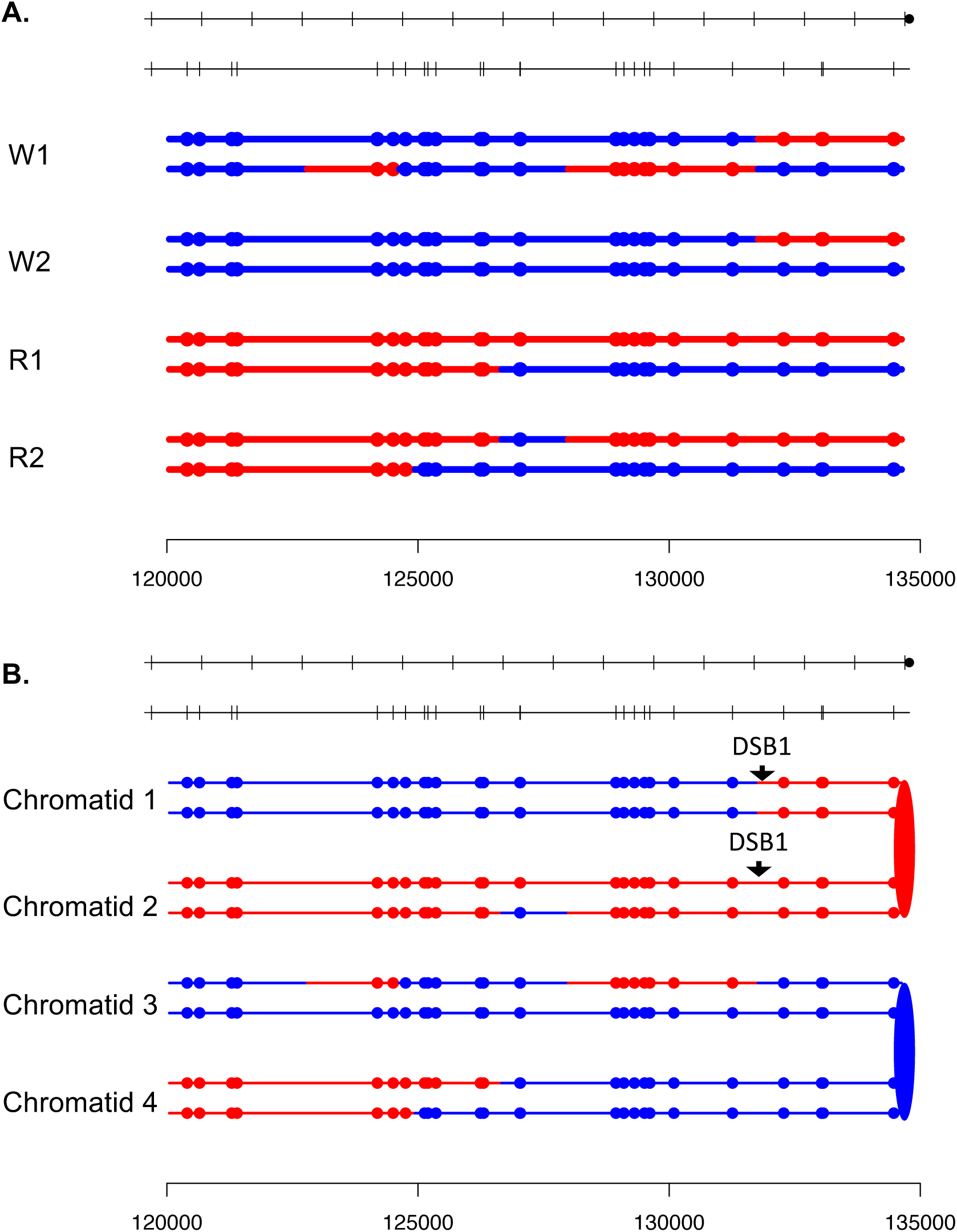

**Fig. S60.**
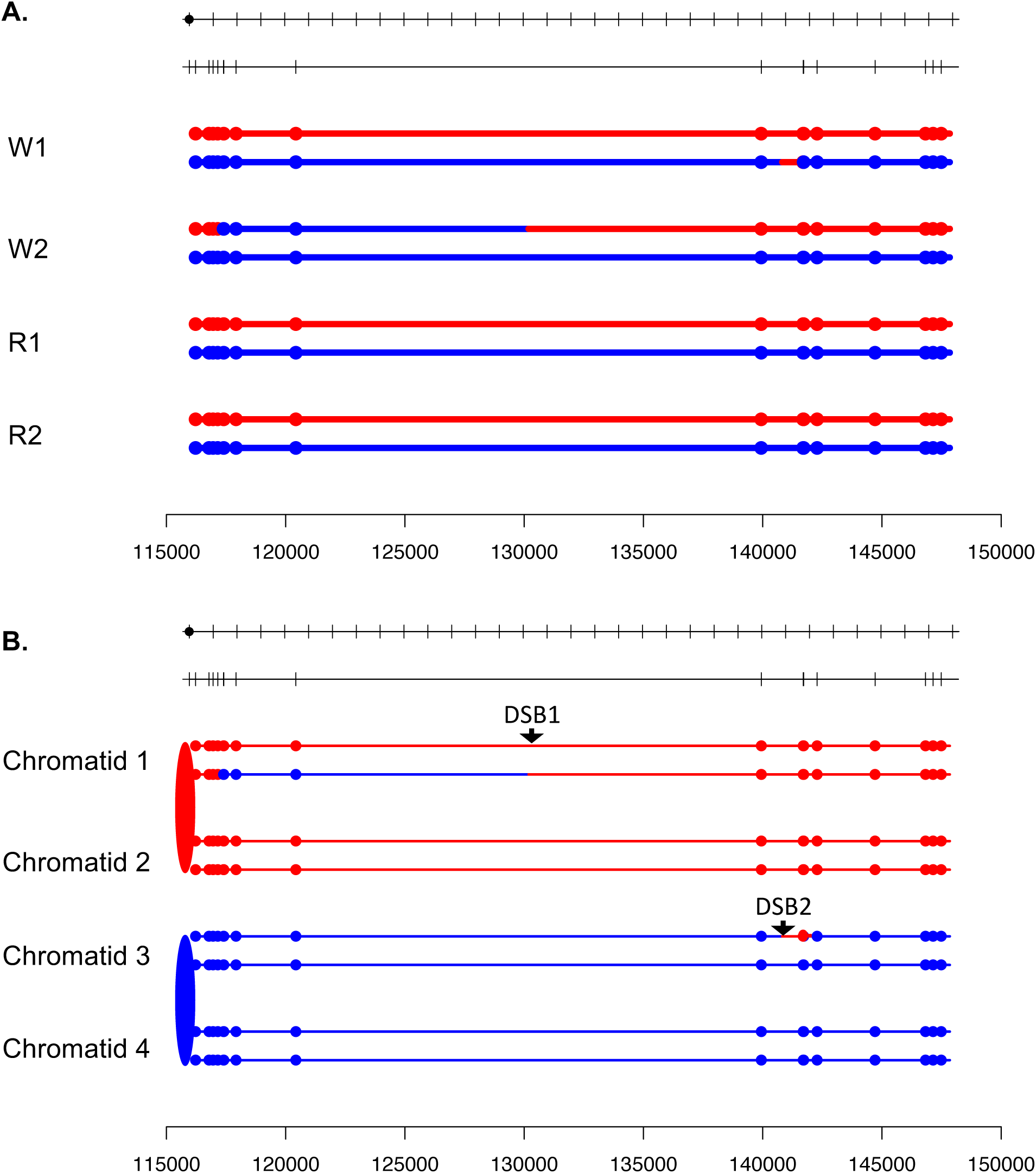

**Fig. S61.**
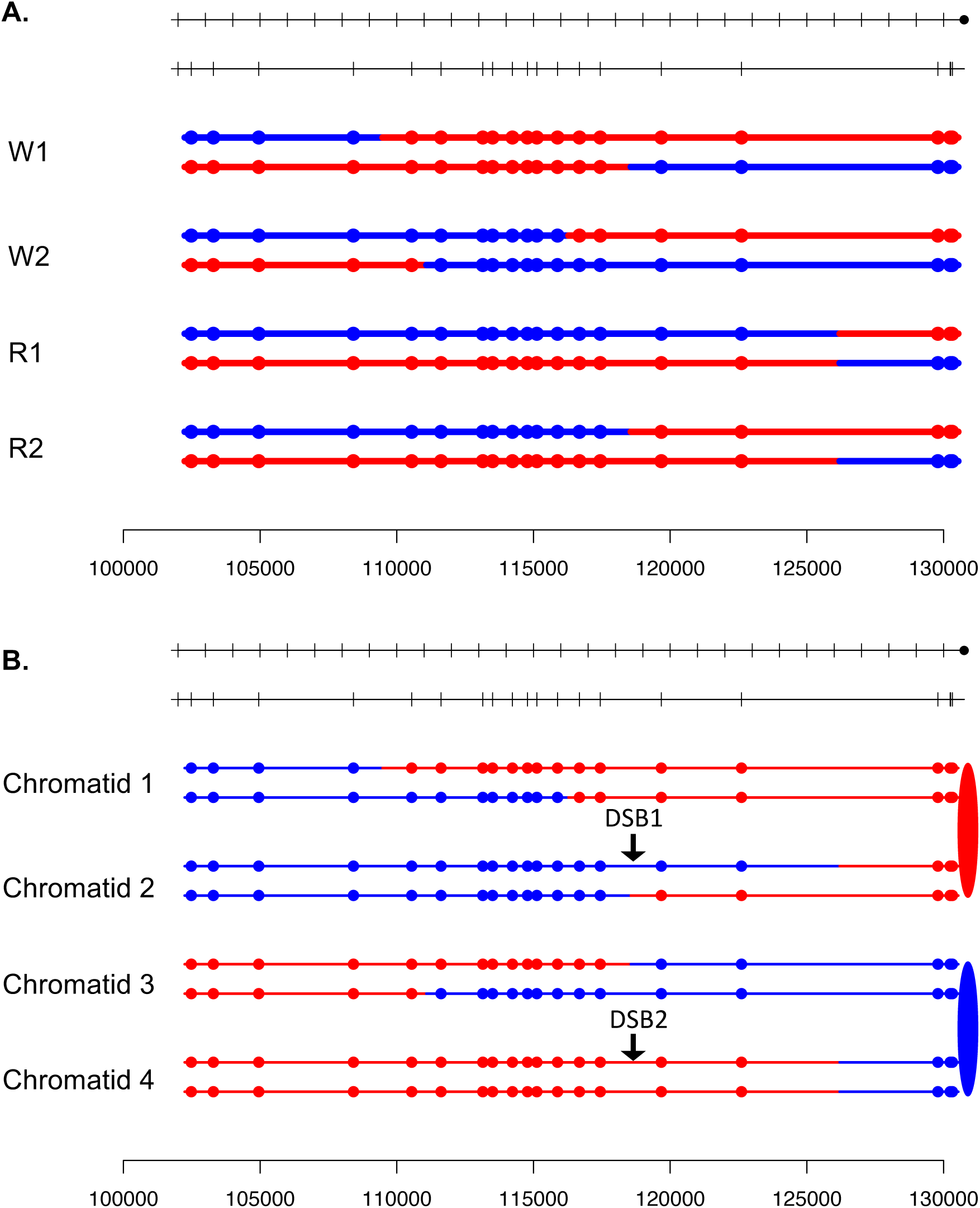

**Fig. S62.**
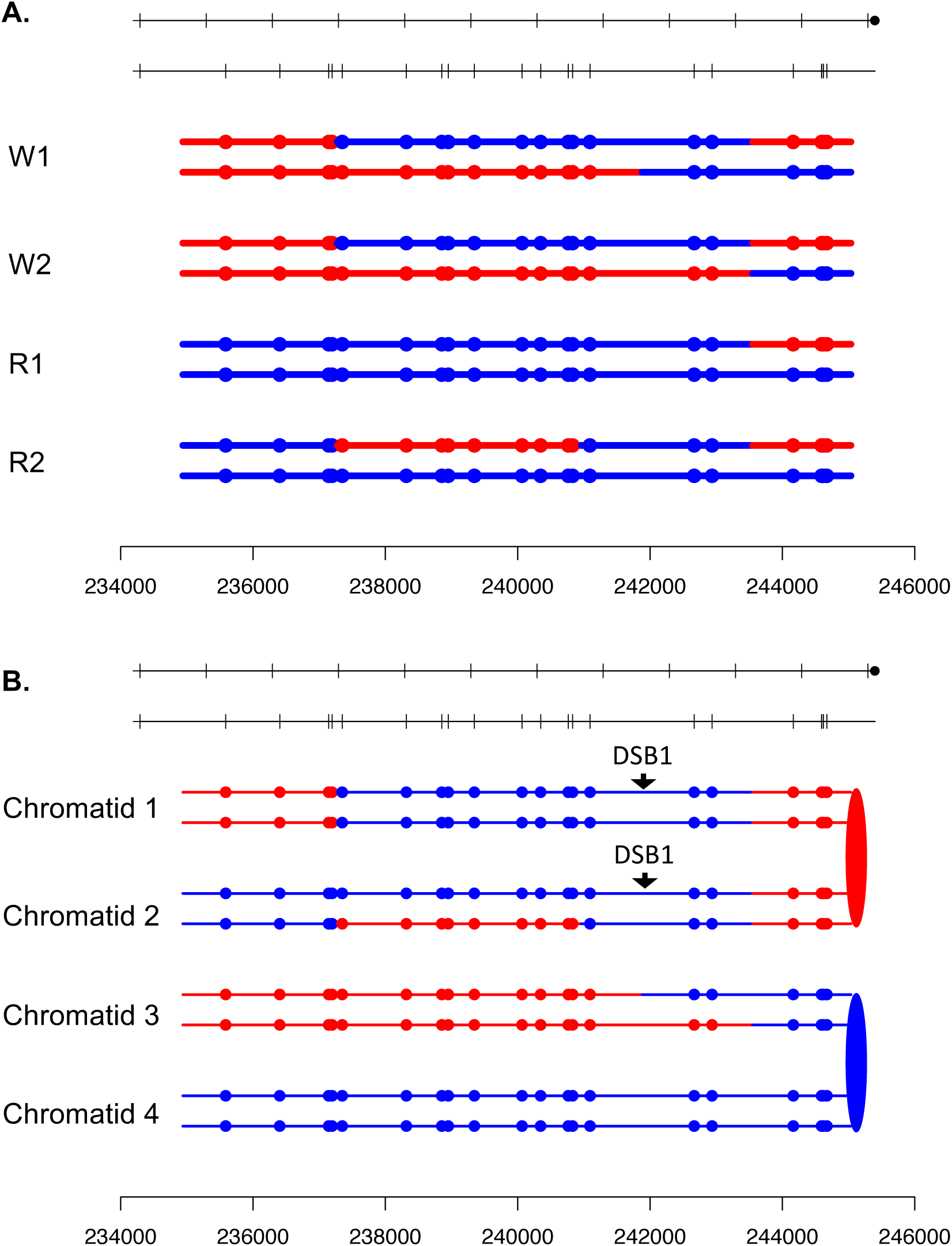

**Fig. S63.**
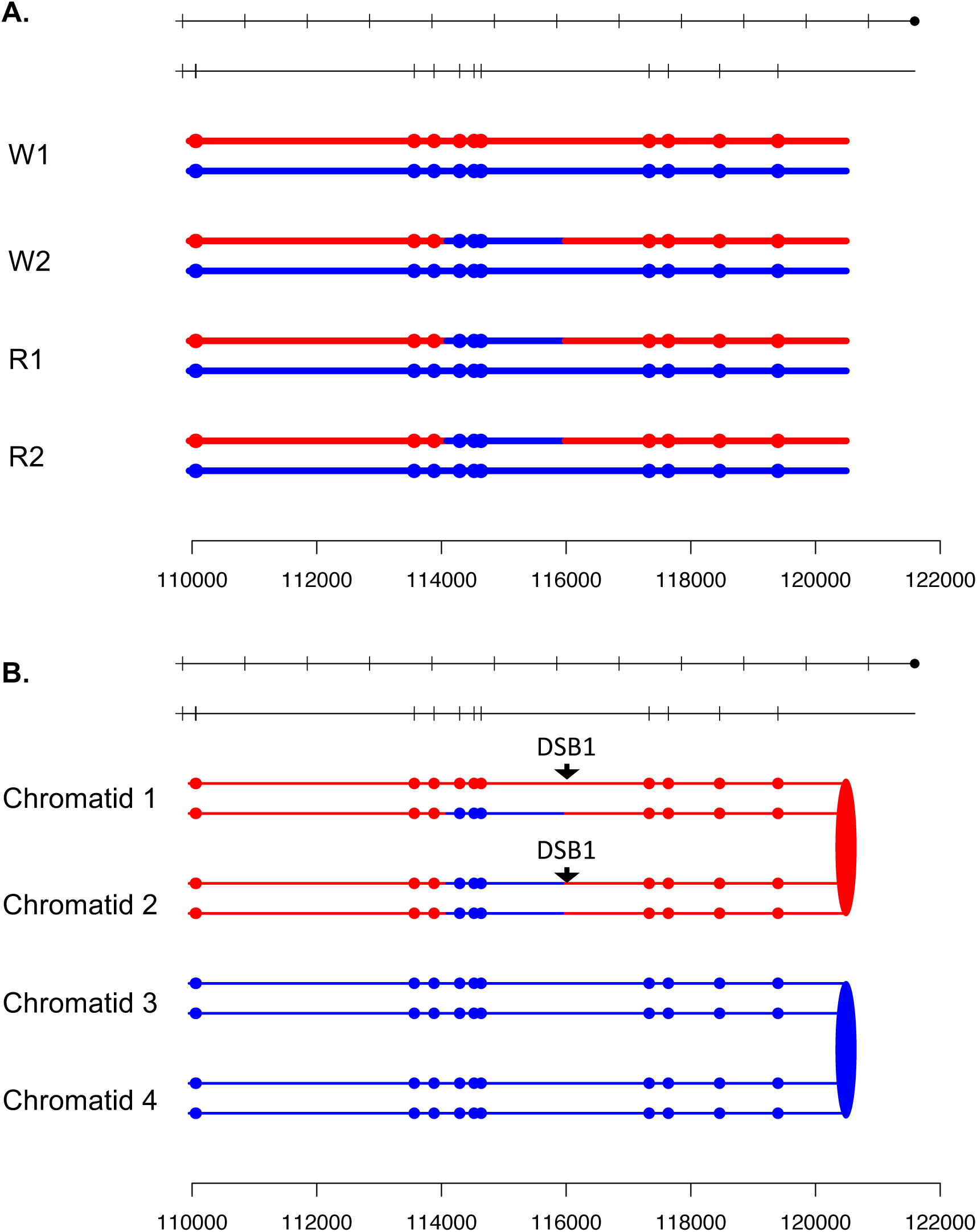

**Fig. S64.**
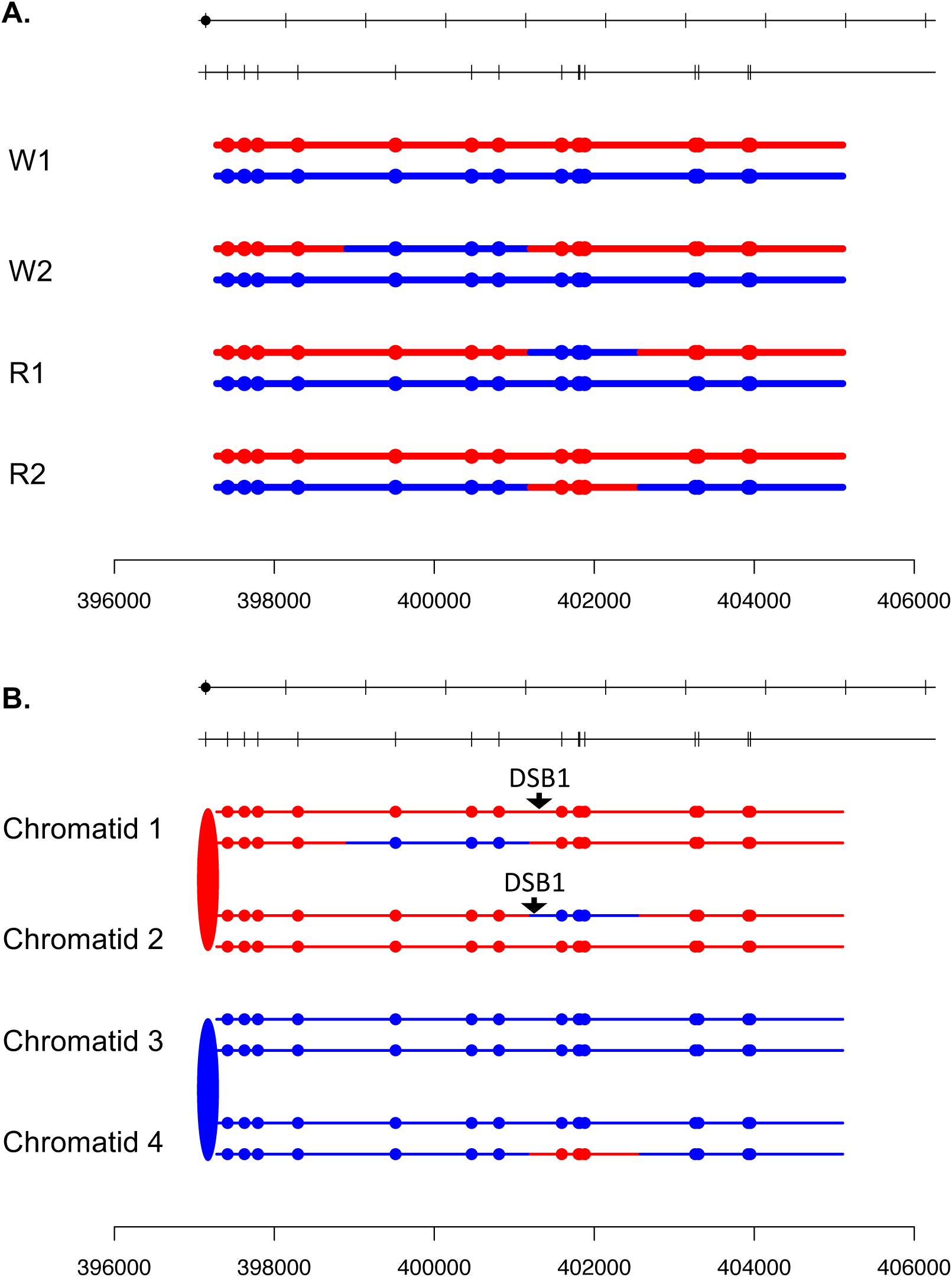

**Fig. S65.**
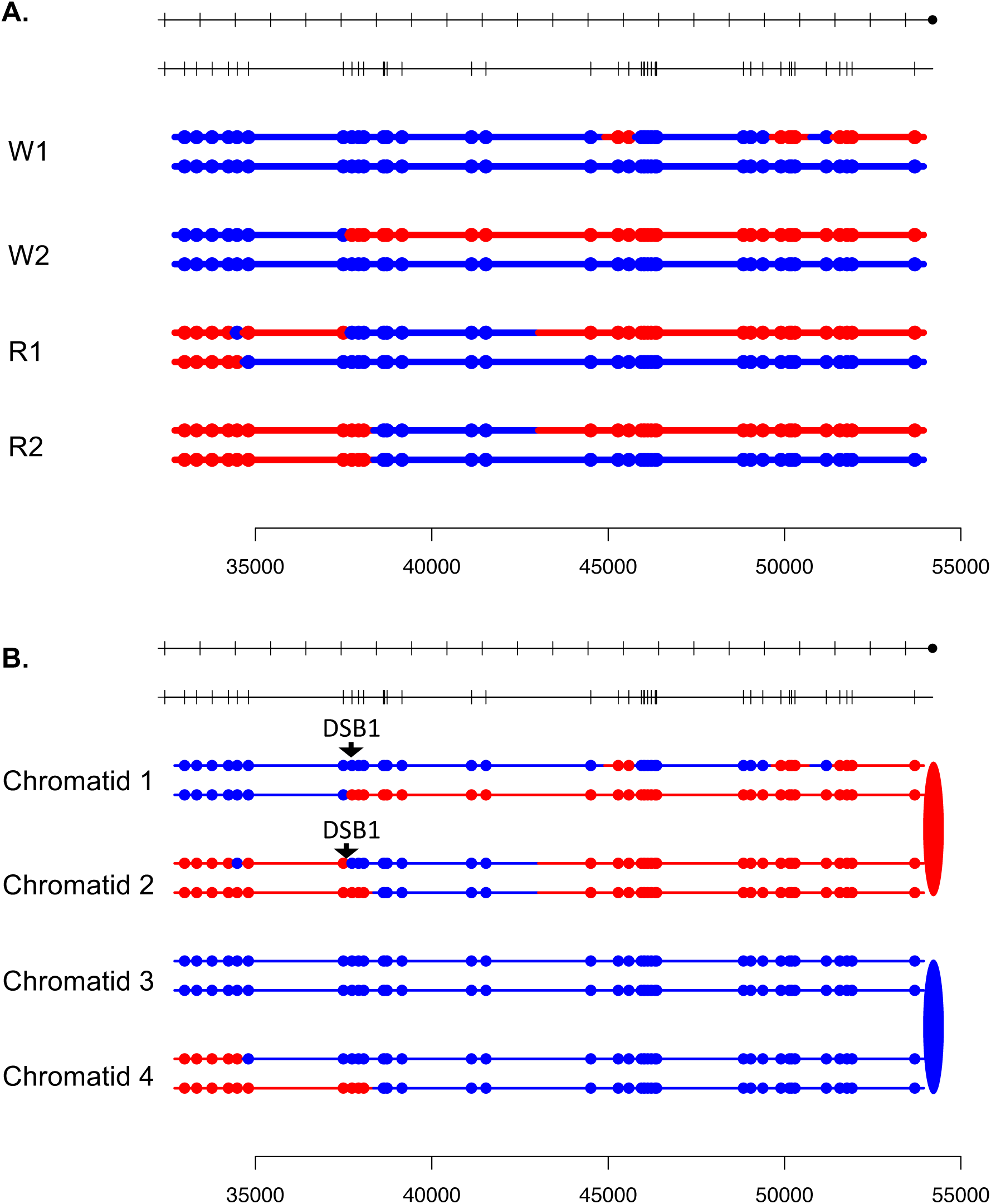

**Fig. S66.**
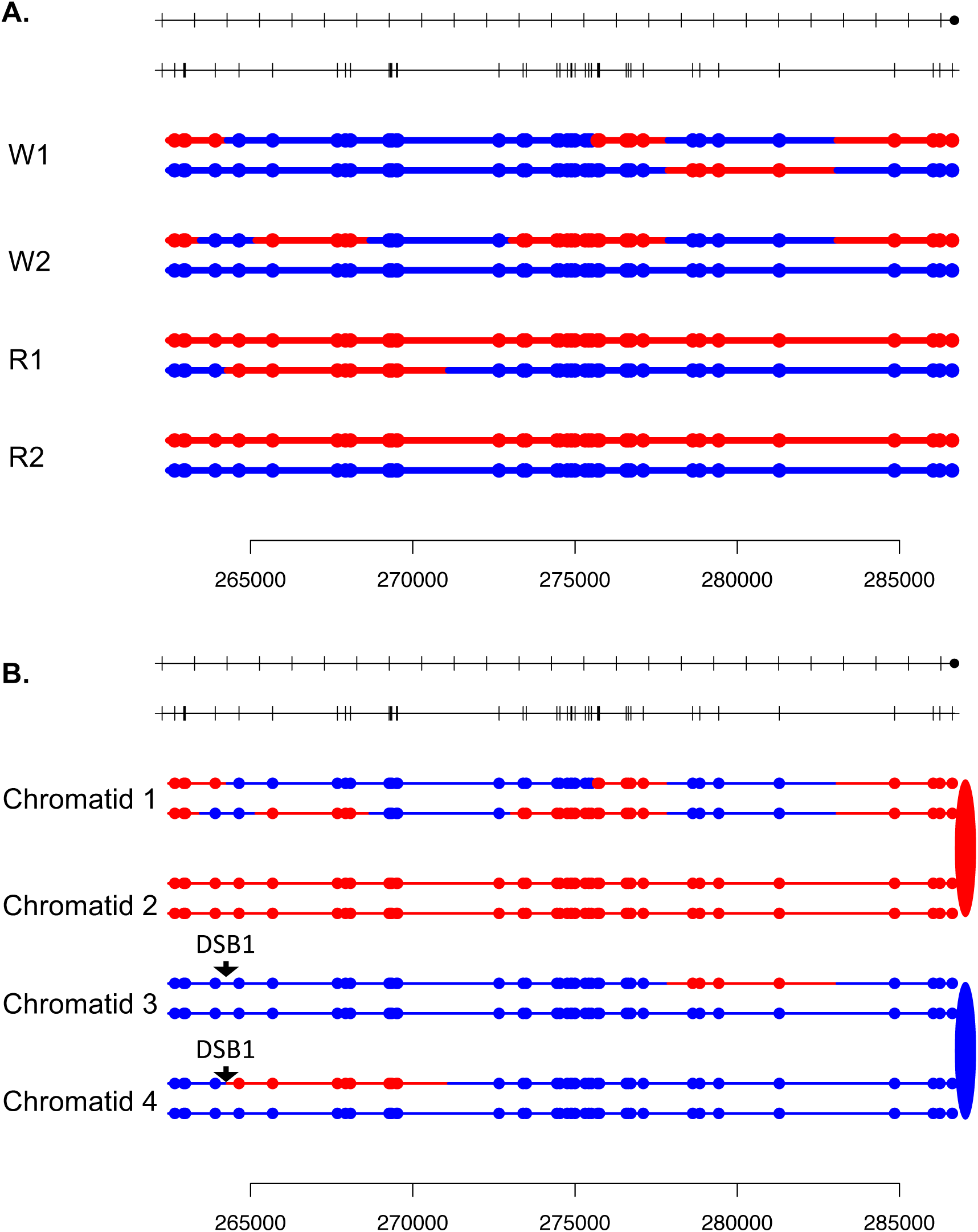

**Fig. S67.**
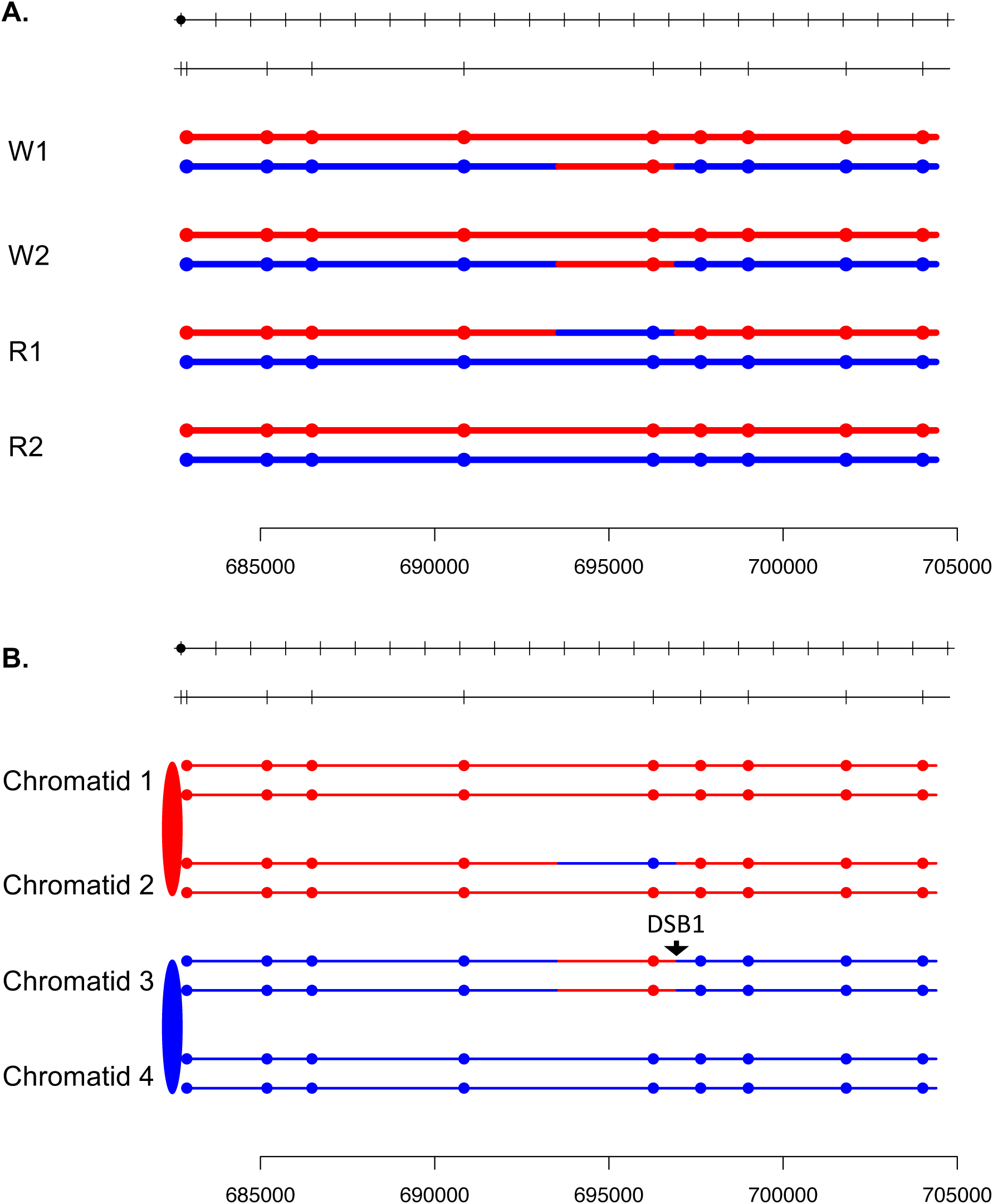

**Fig. S68.**
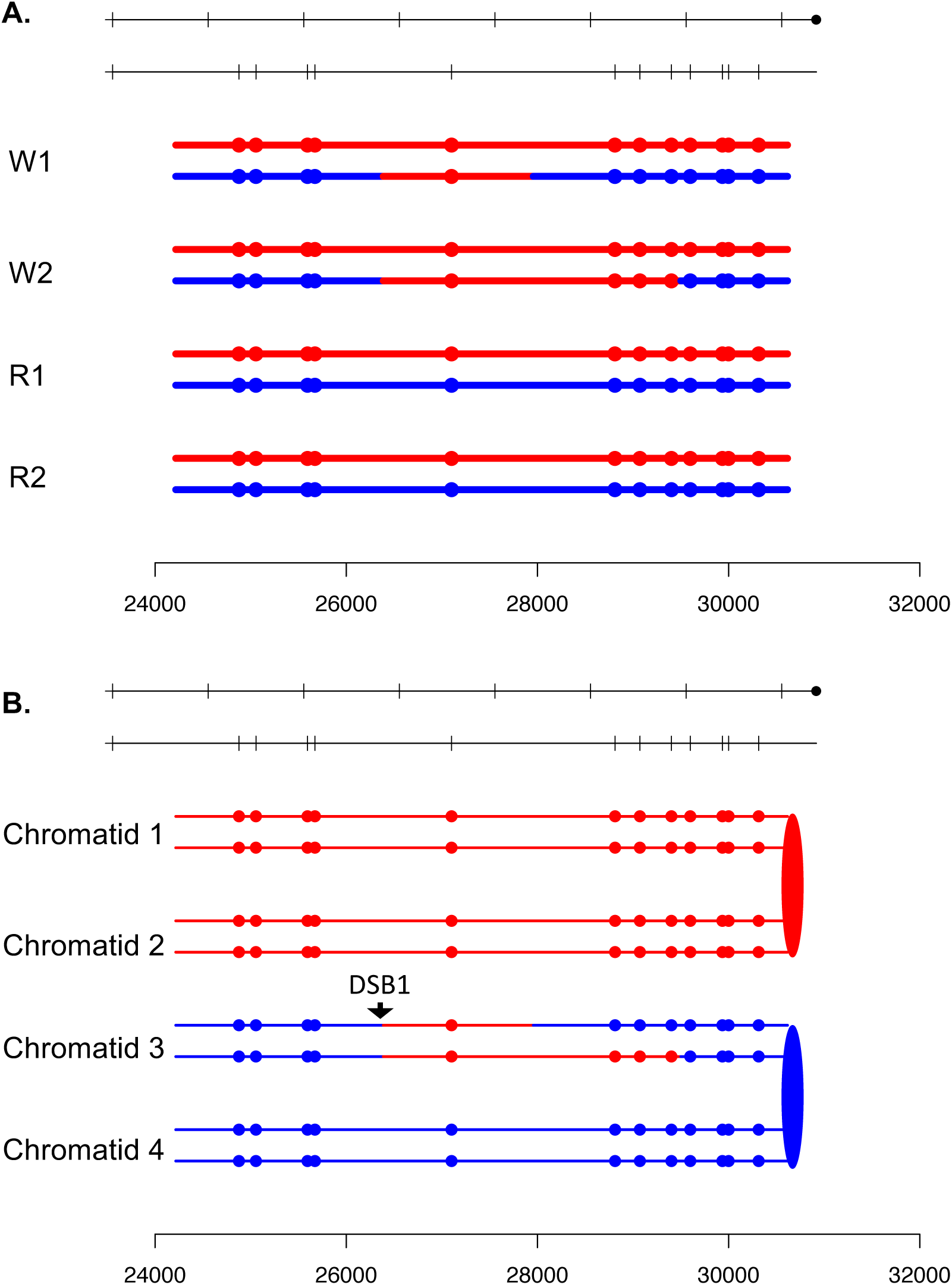

**Fig. S69.**
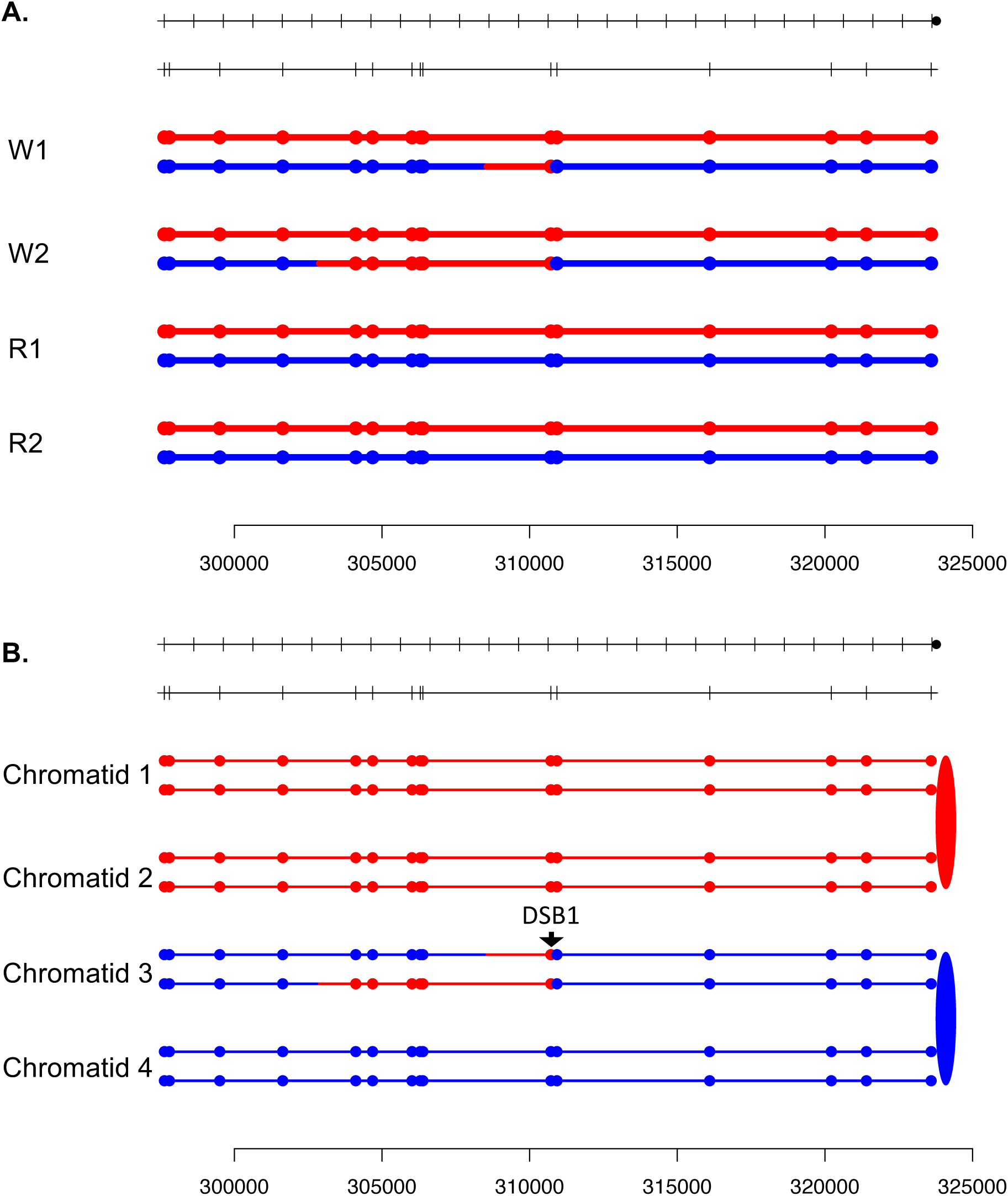

**Fig. S70.**
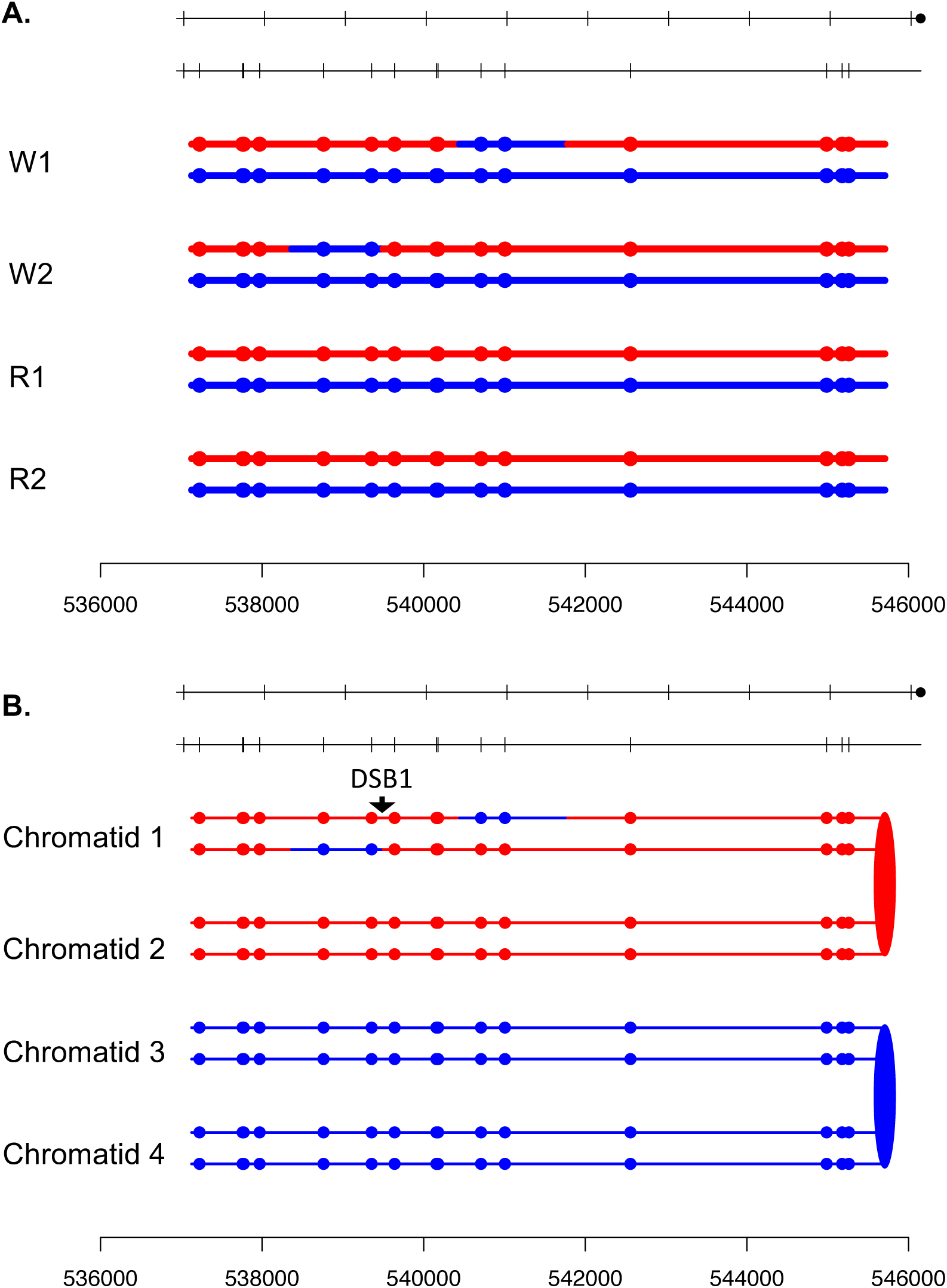

**Fig. S71.**
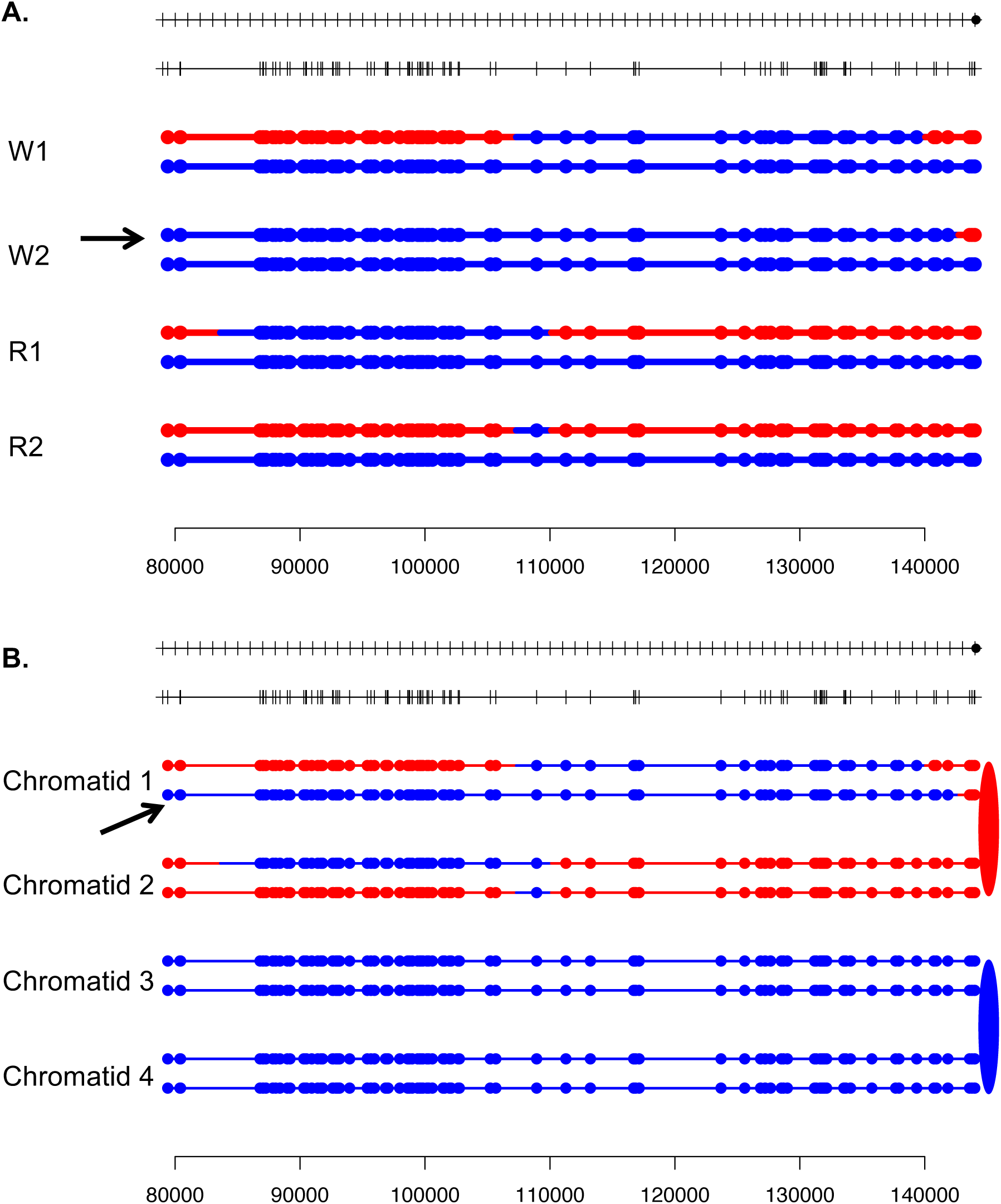

**Fig. S72.**
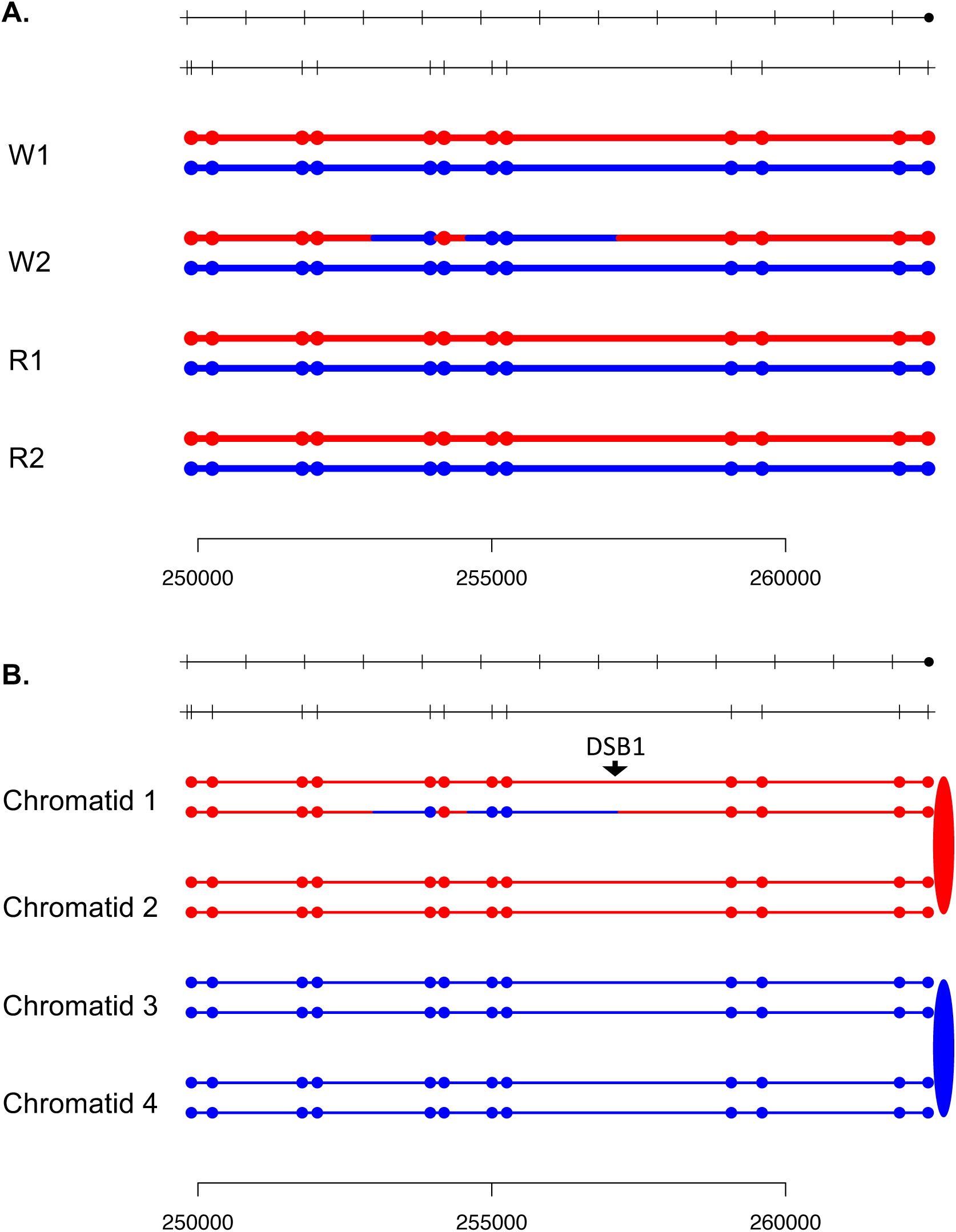

**Fig. S73.**
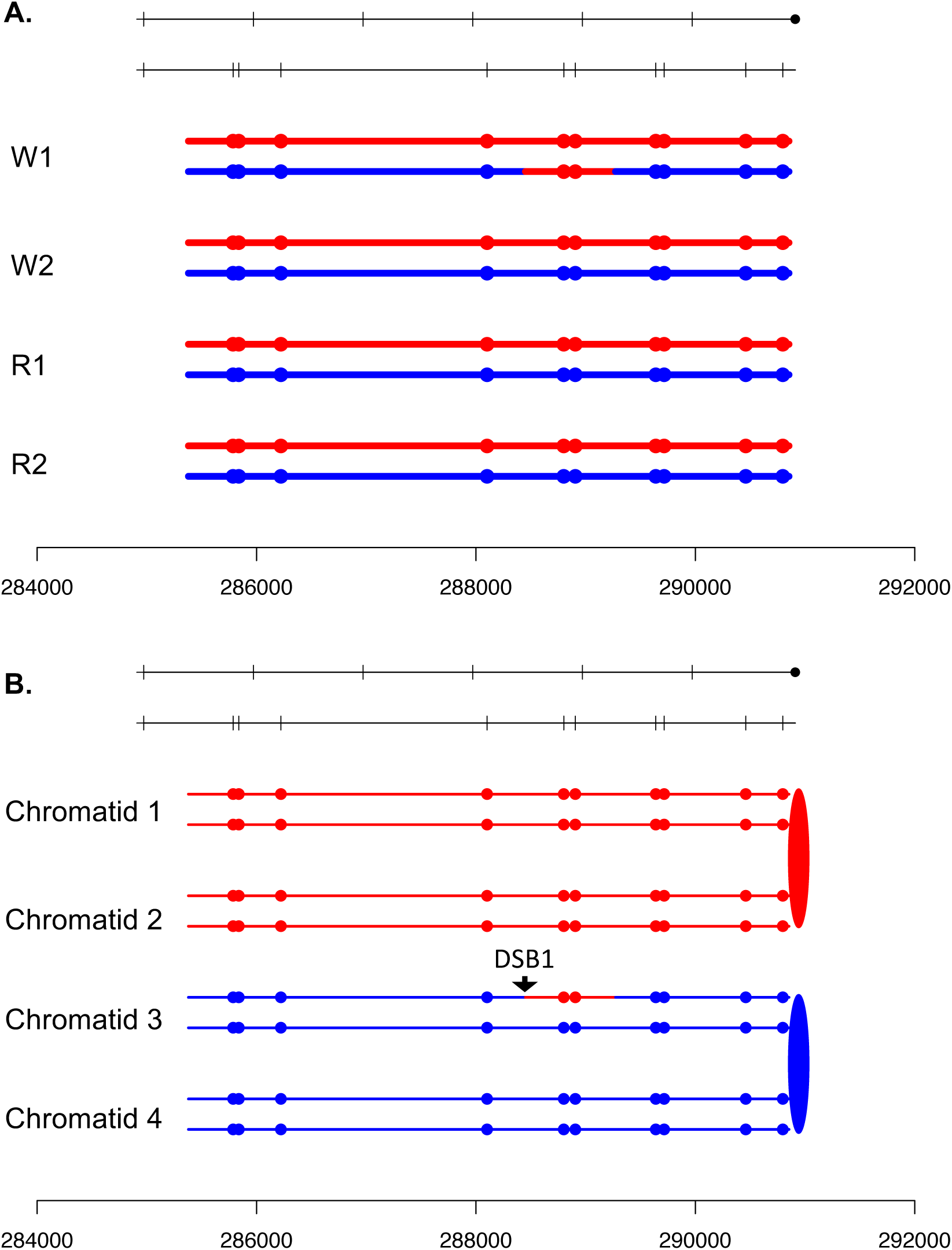

**Fig. S74.**
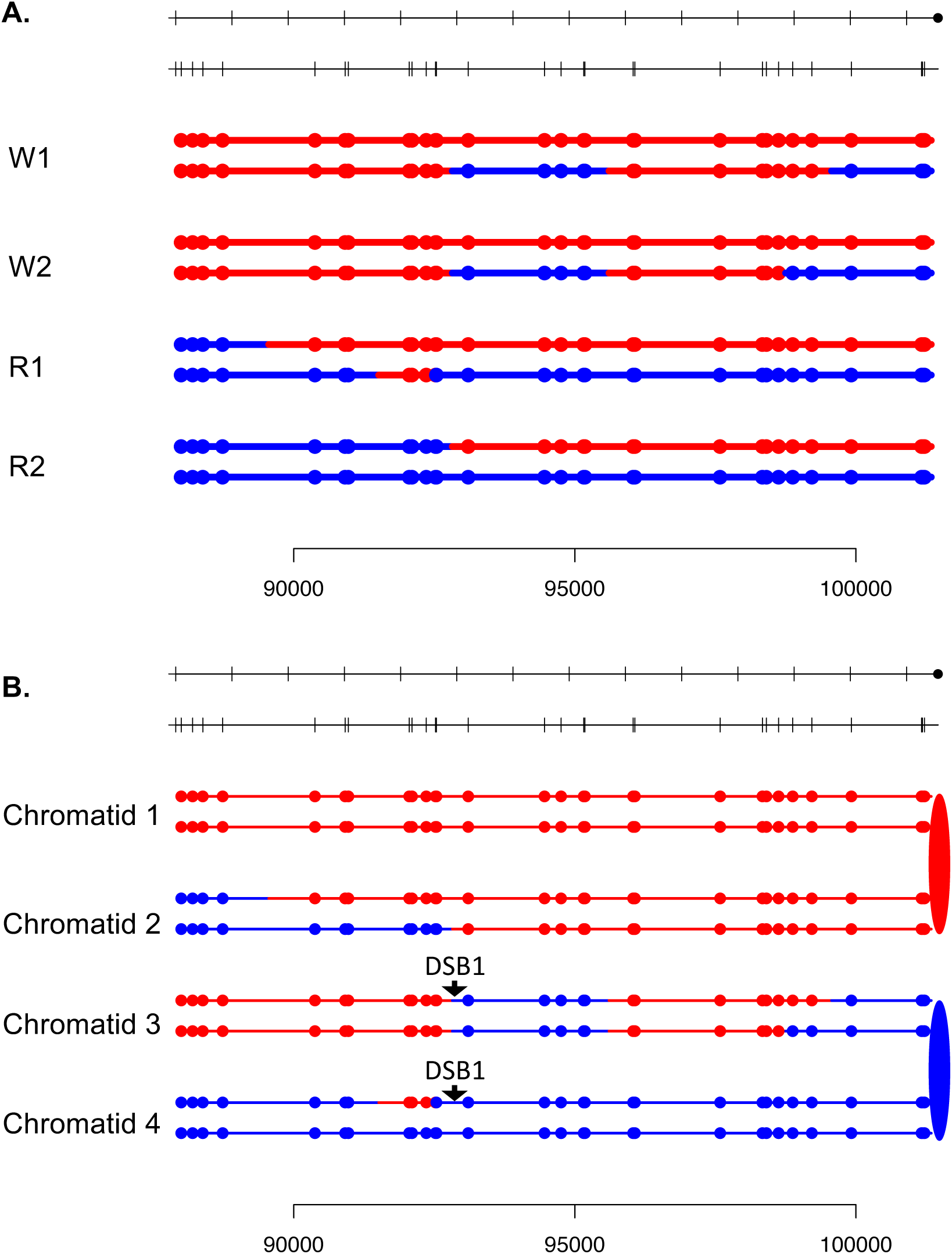

**Fig. S75.**
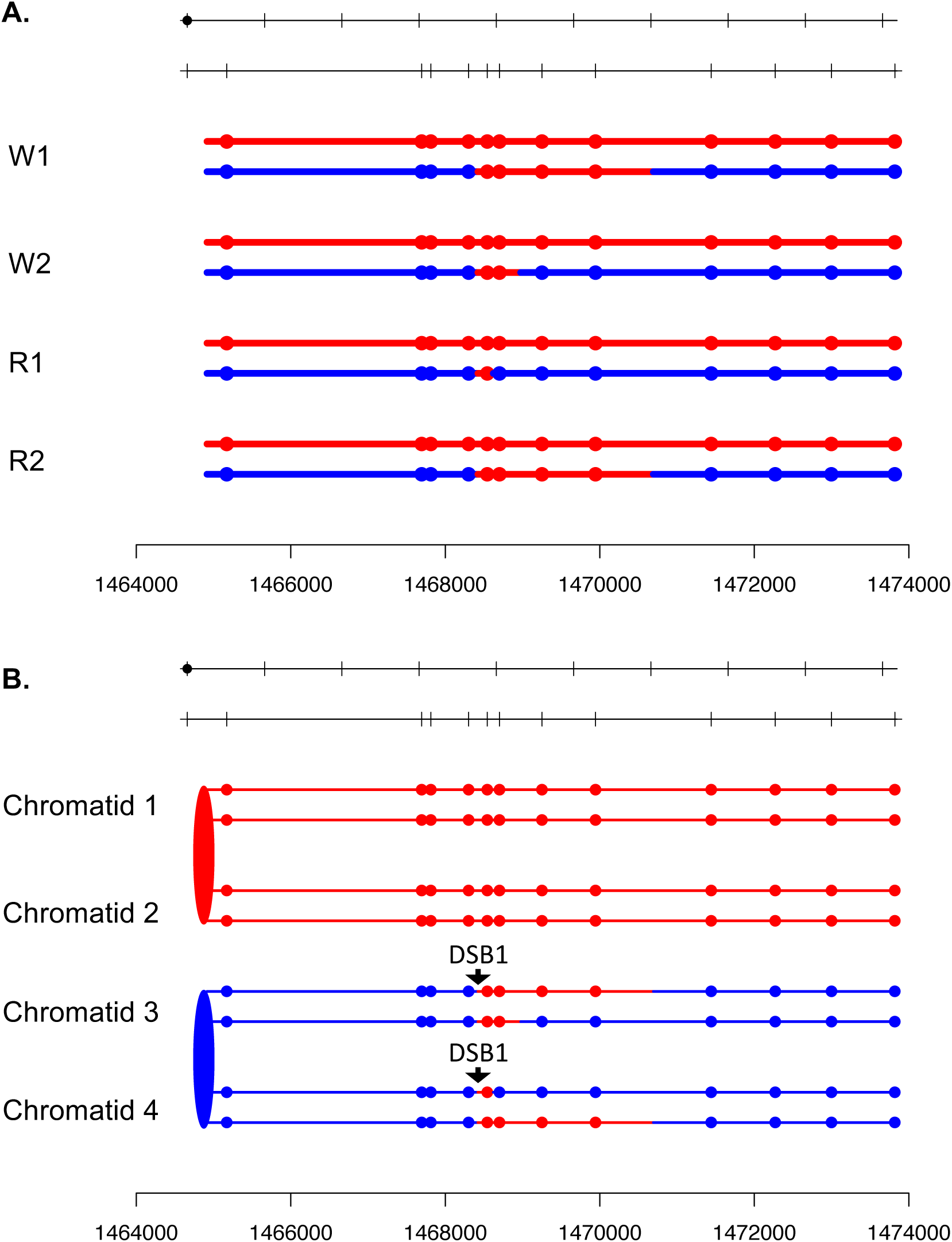

**Fig. S76.**
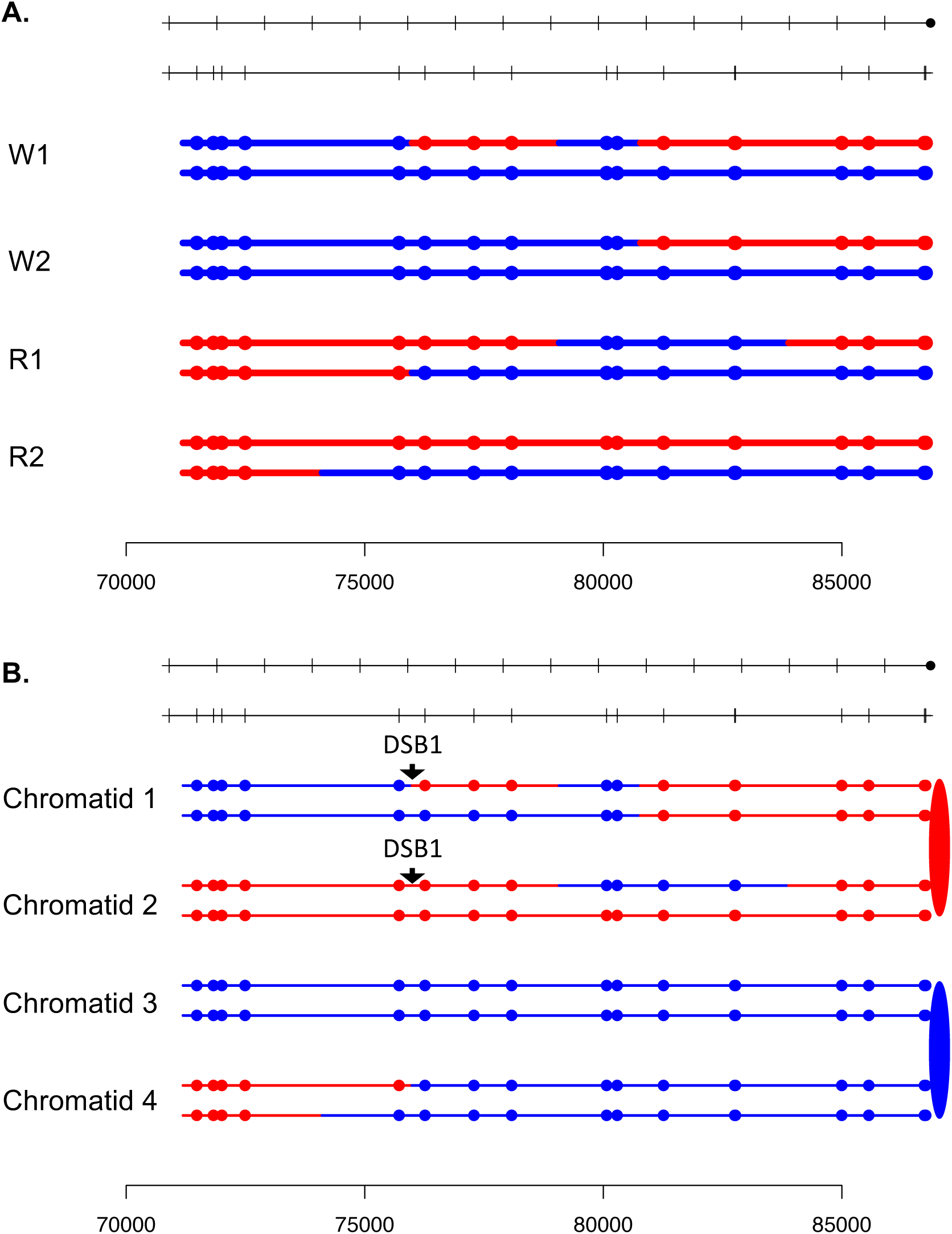

**Fig. S77.**
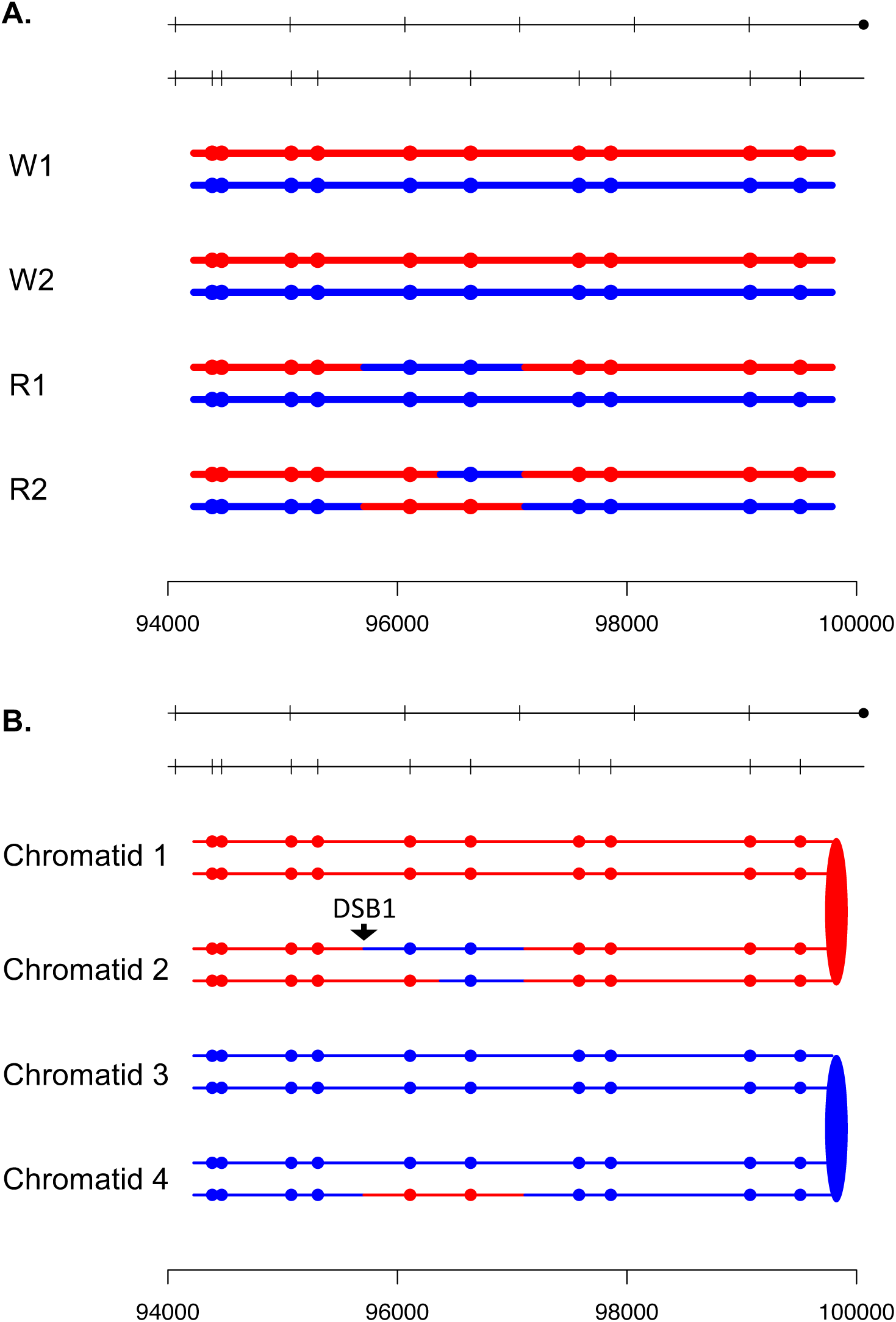

**Fig. S78.**
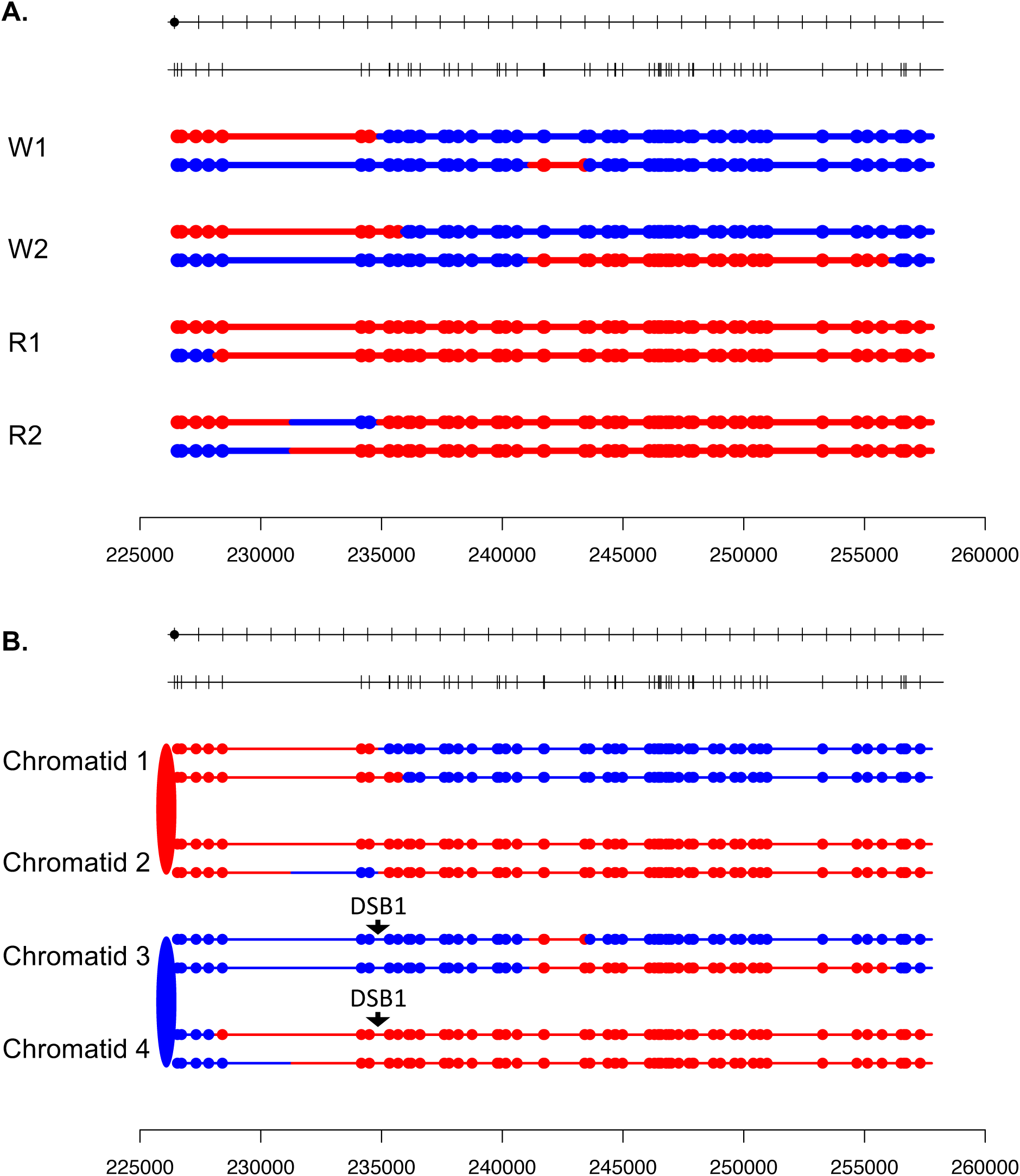

**Fig. S79.**
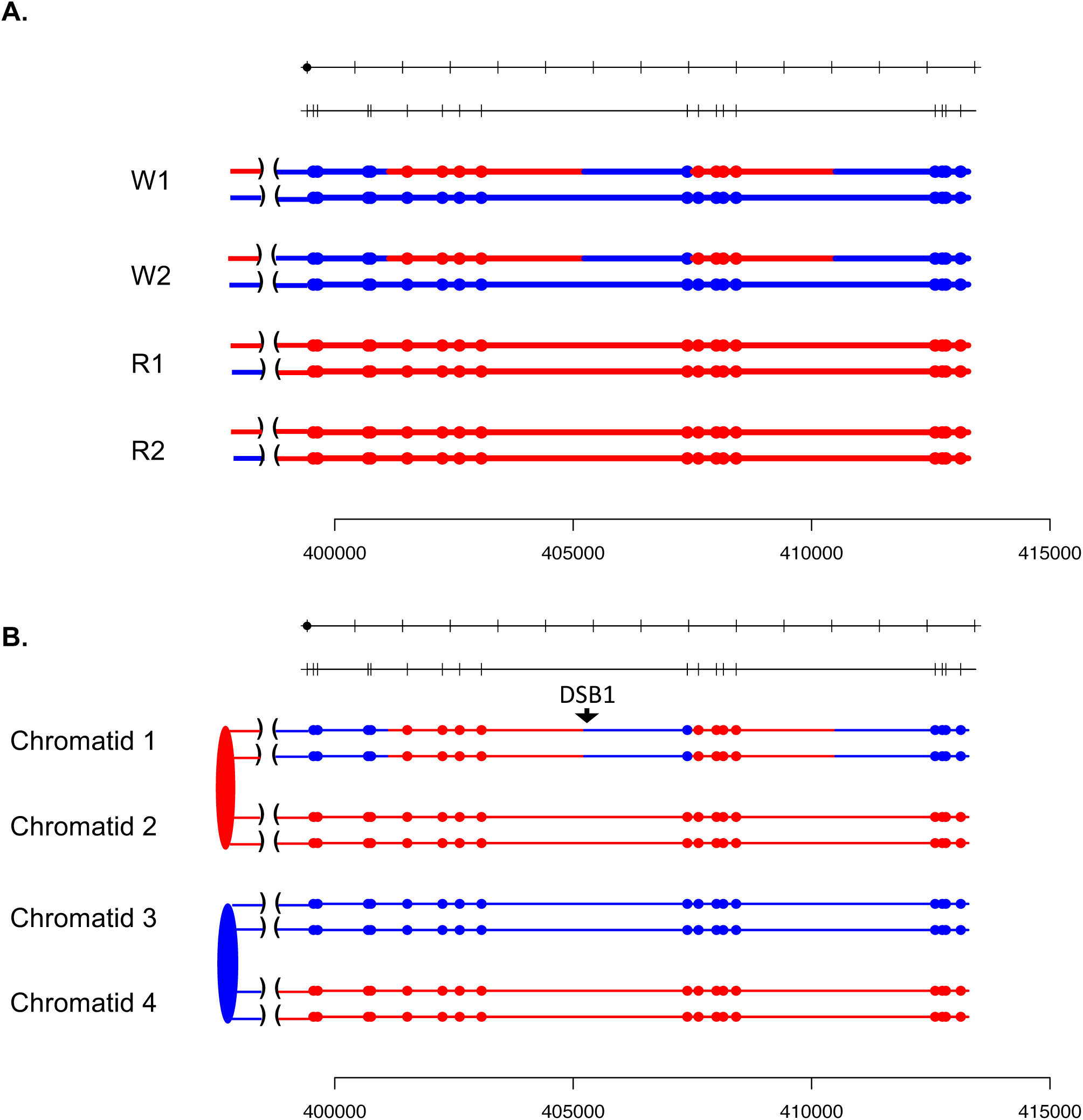

**Fig. S80.**
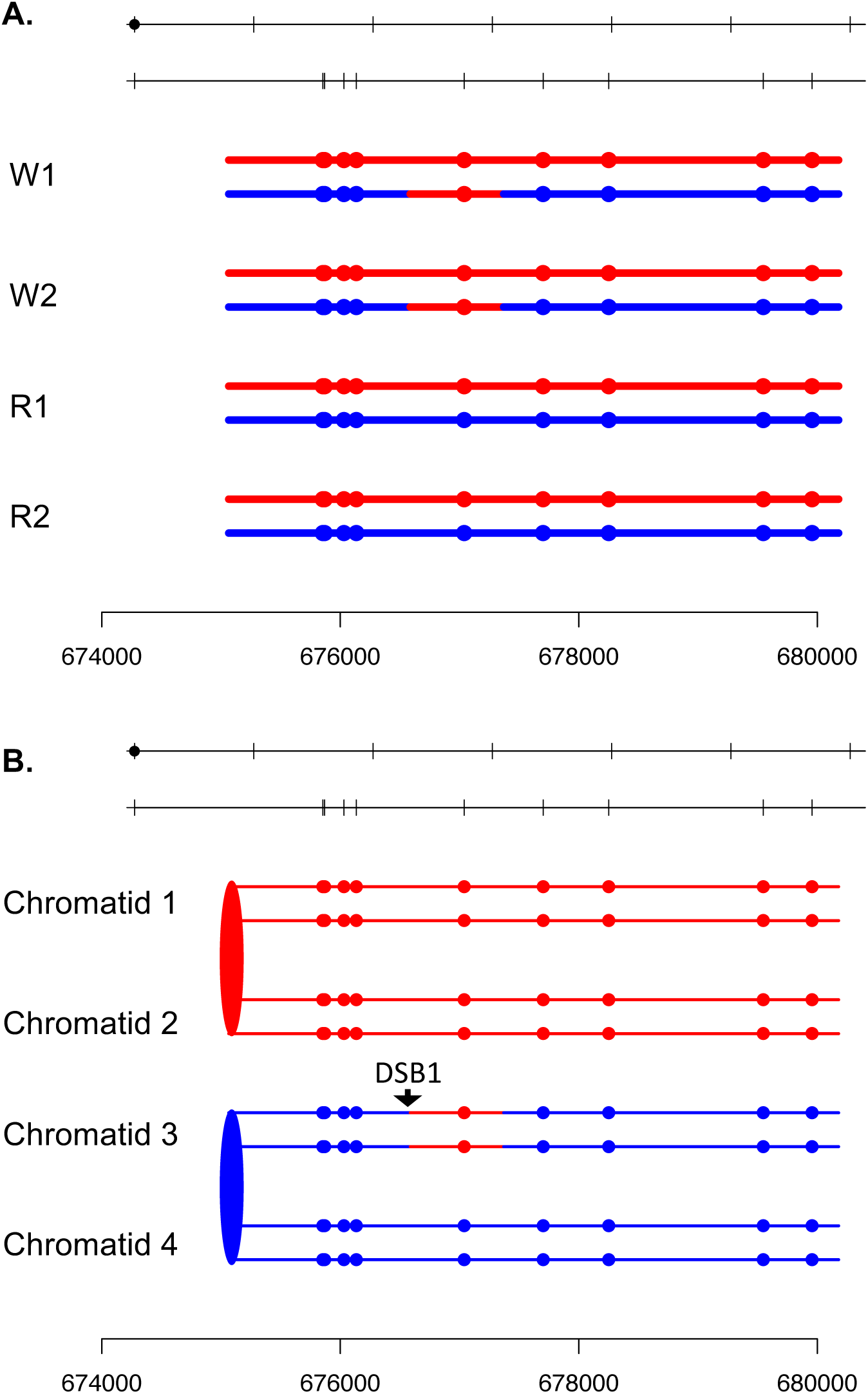

**Fig. S81.**
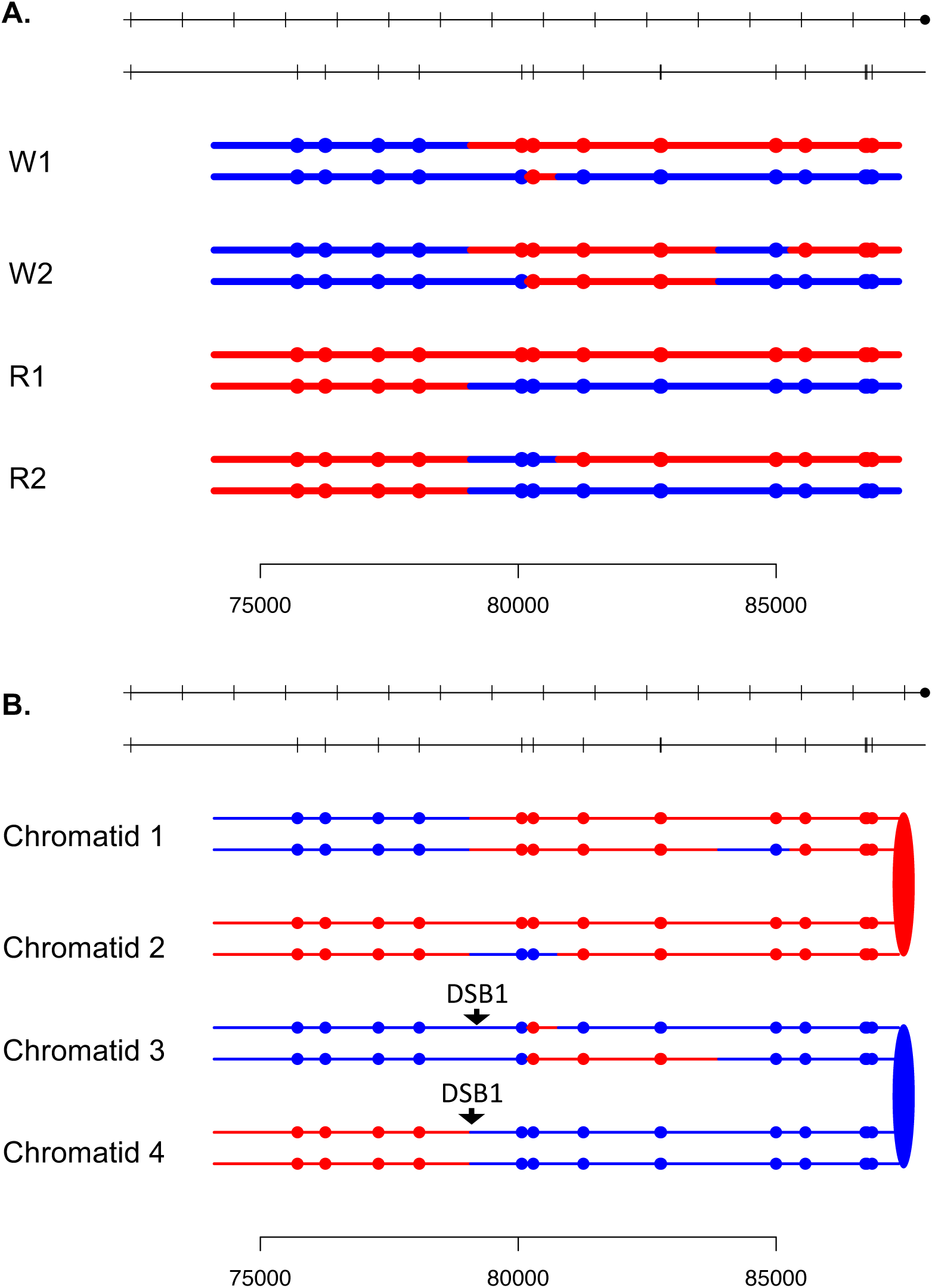

**Fig. S82.**
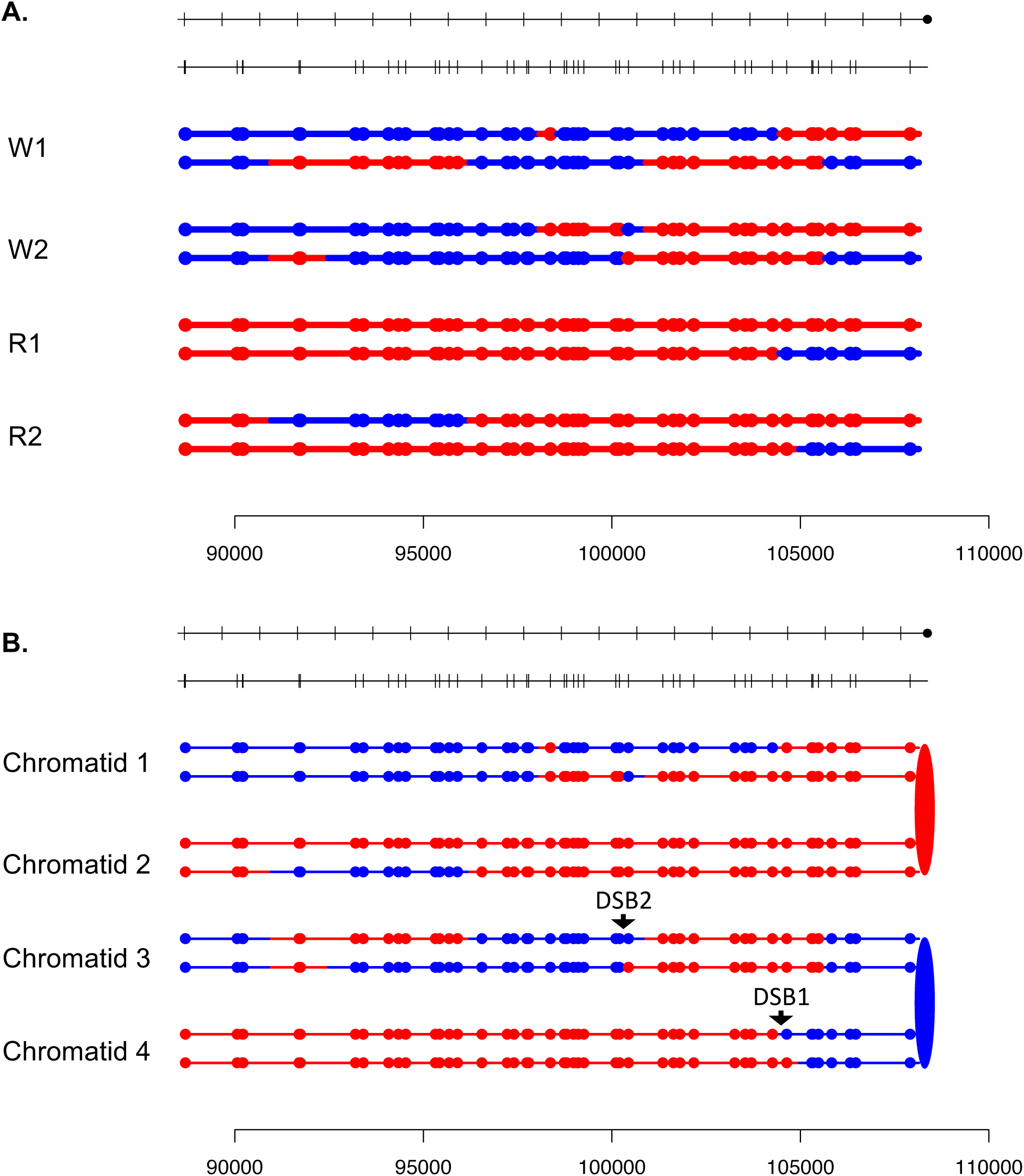

**Fig. S83.**
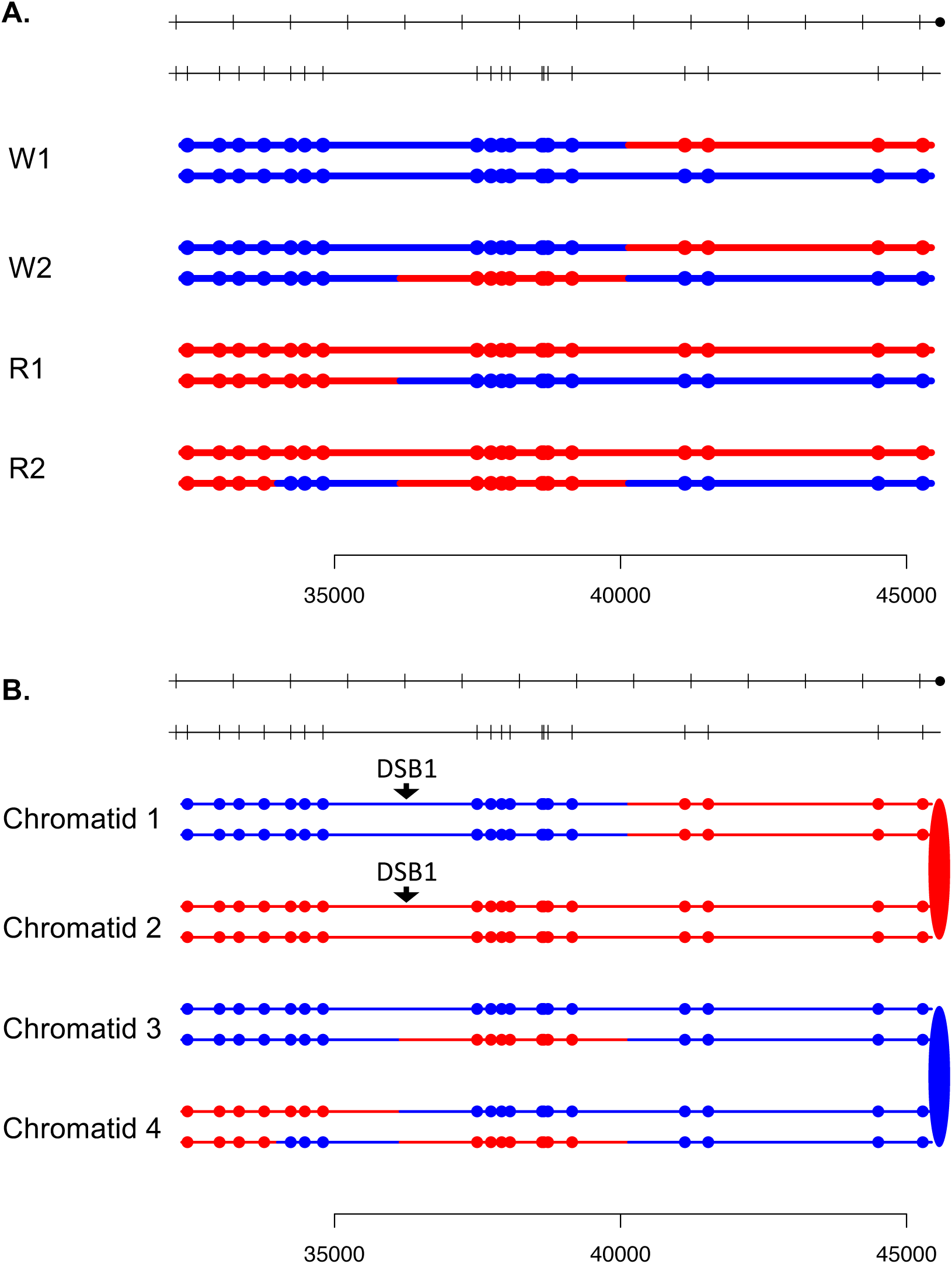

**Fig. S84.**
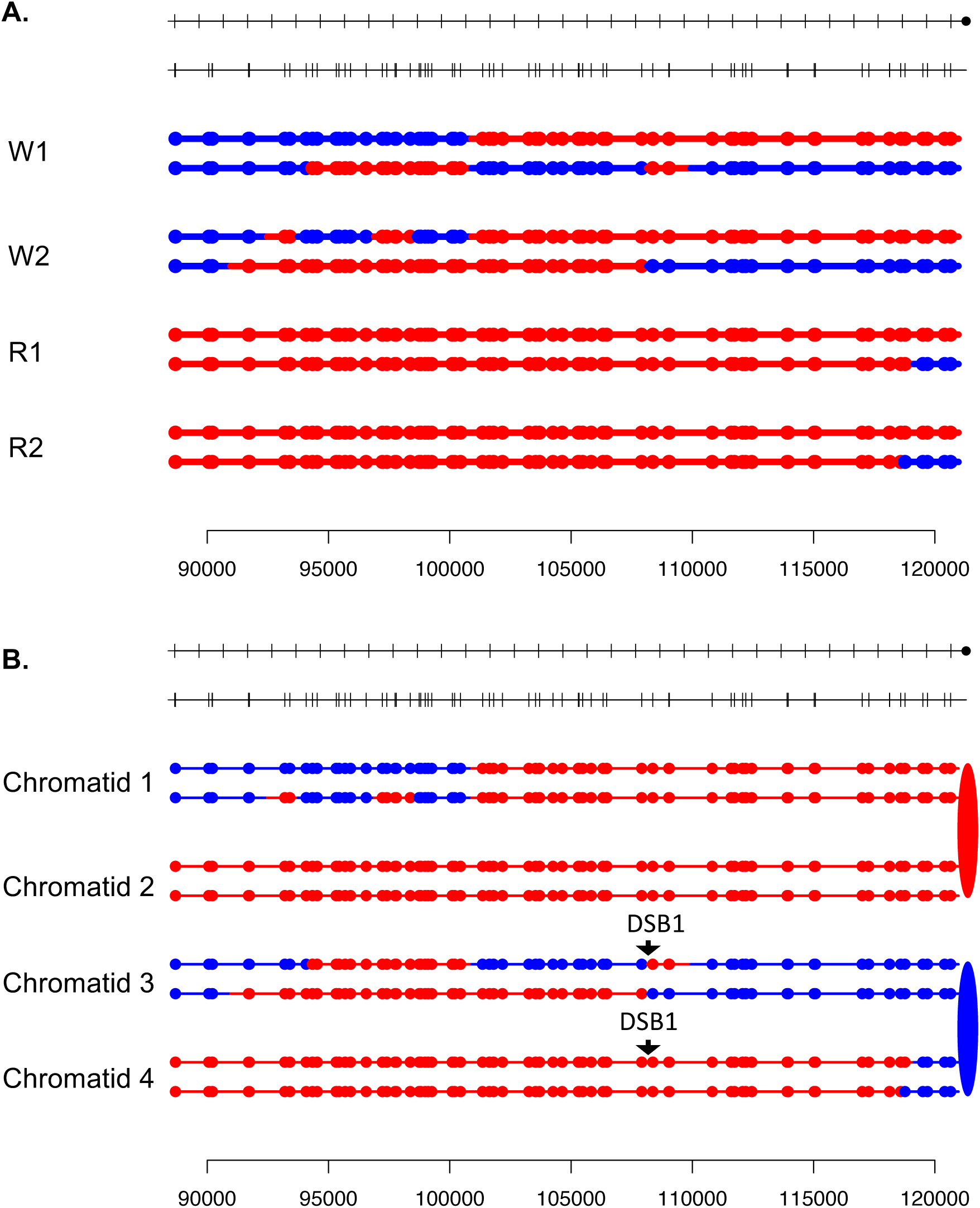

**Fig. S85.**
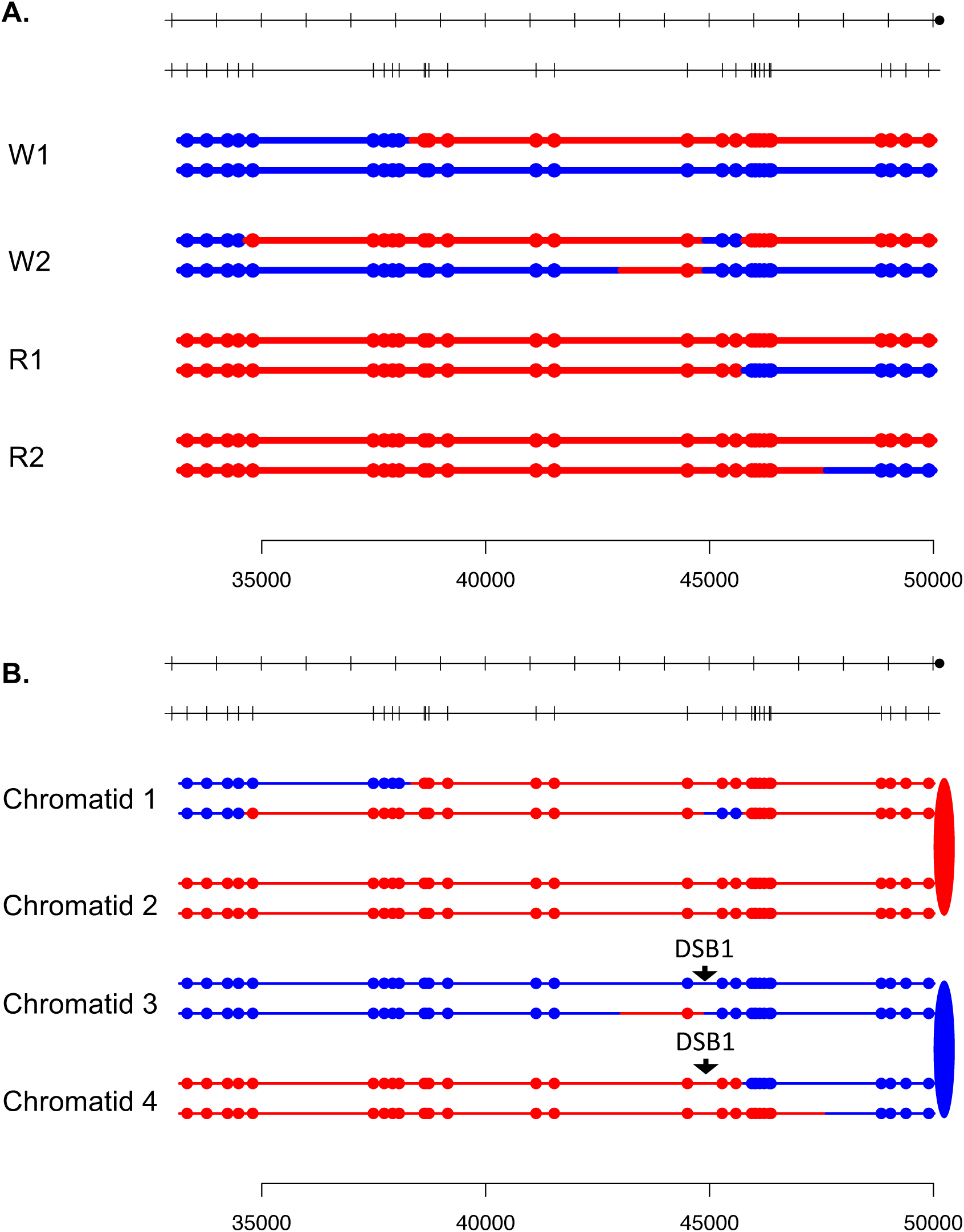

**Fig. S86.**
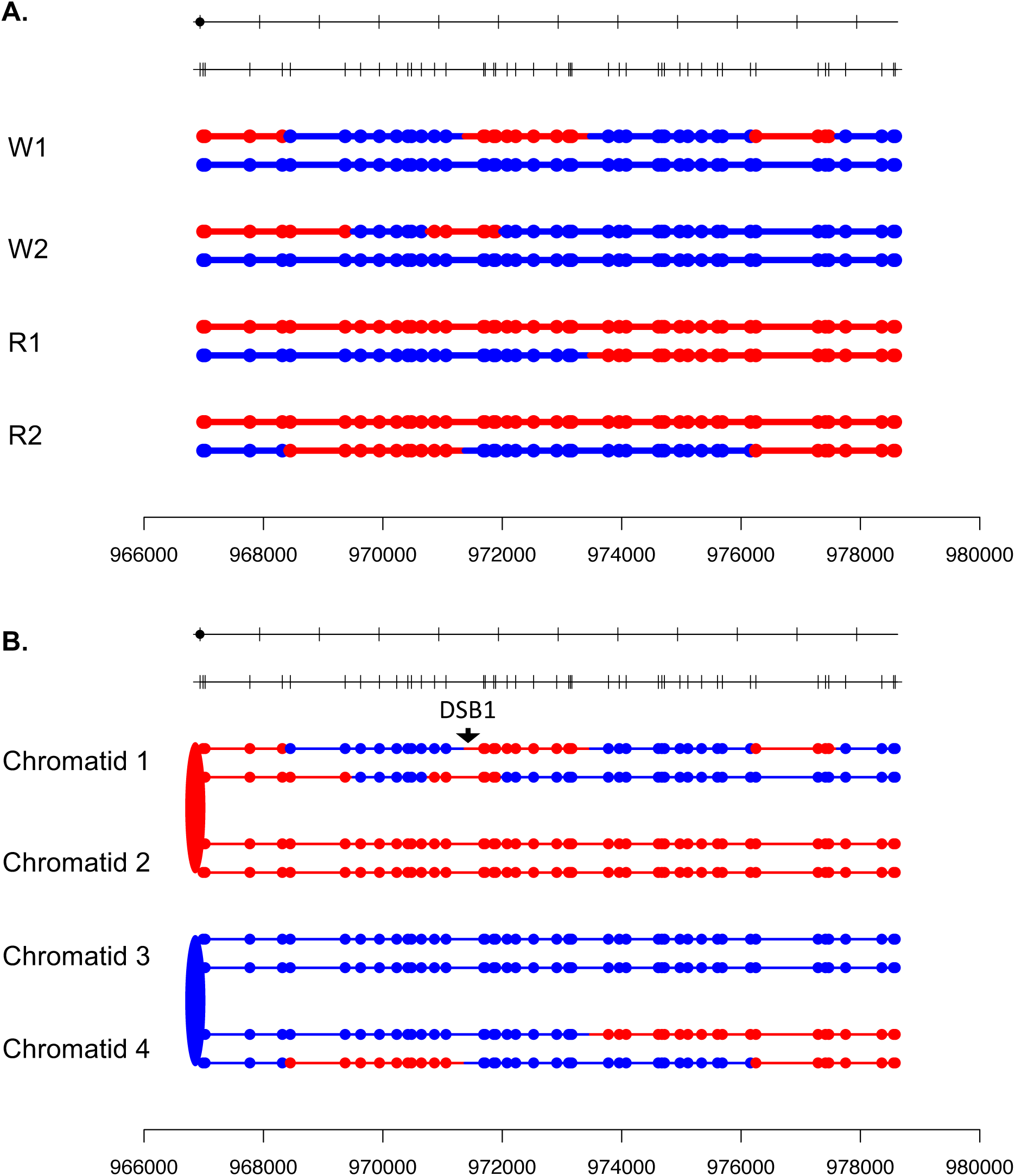

**Fig. S87.**
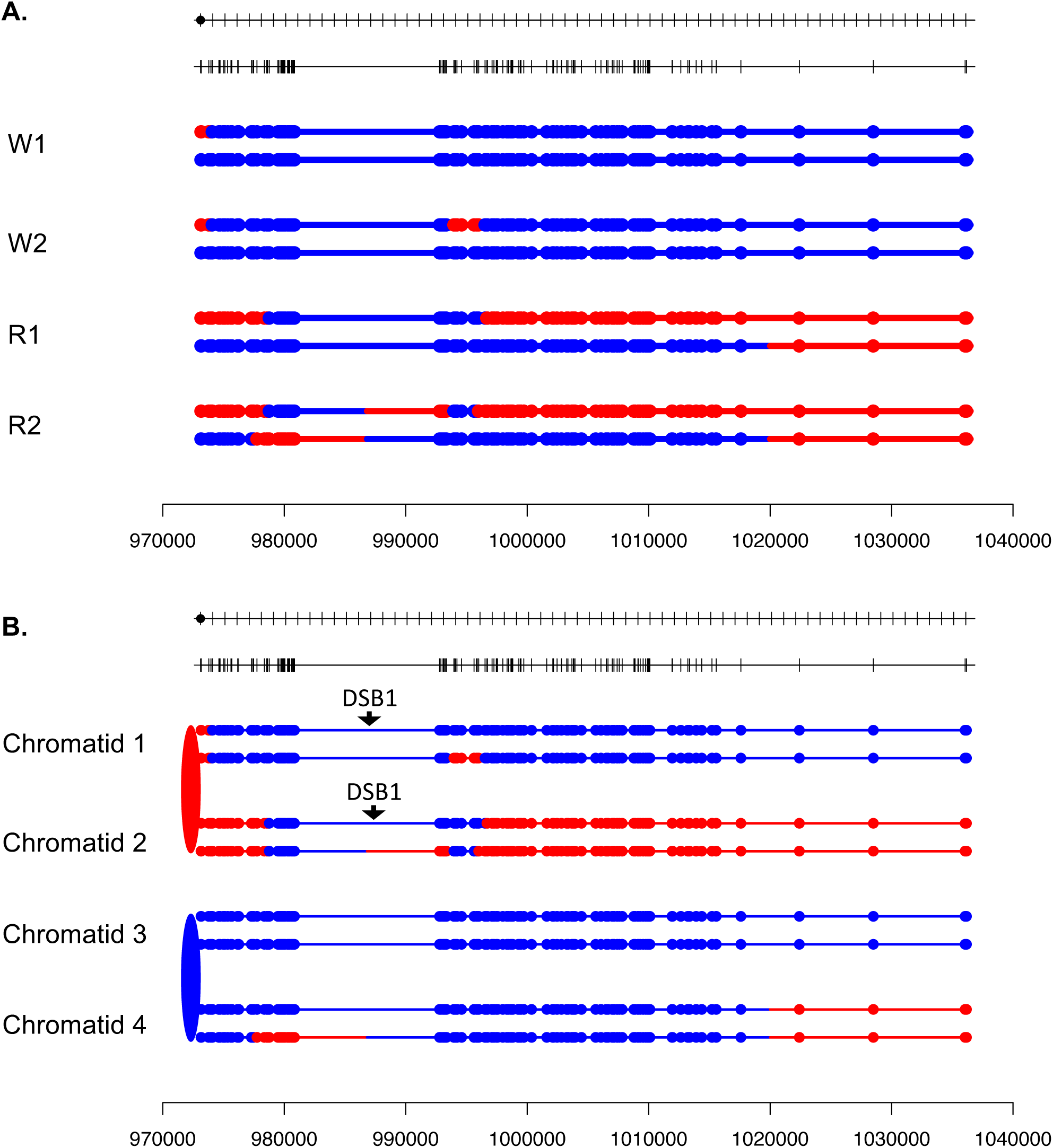

**Fig. S88.**
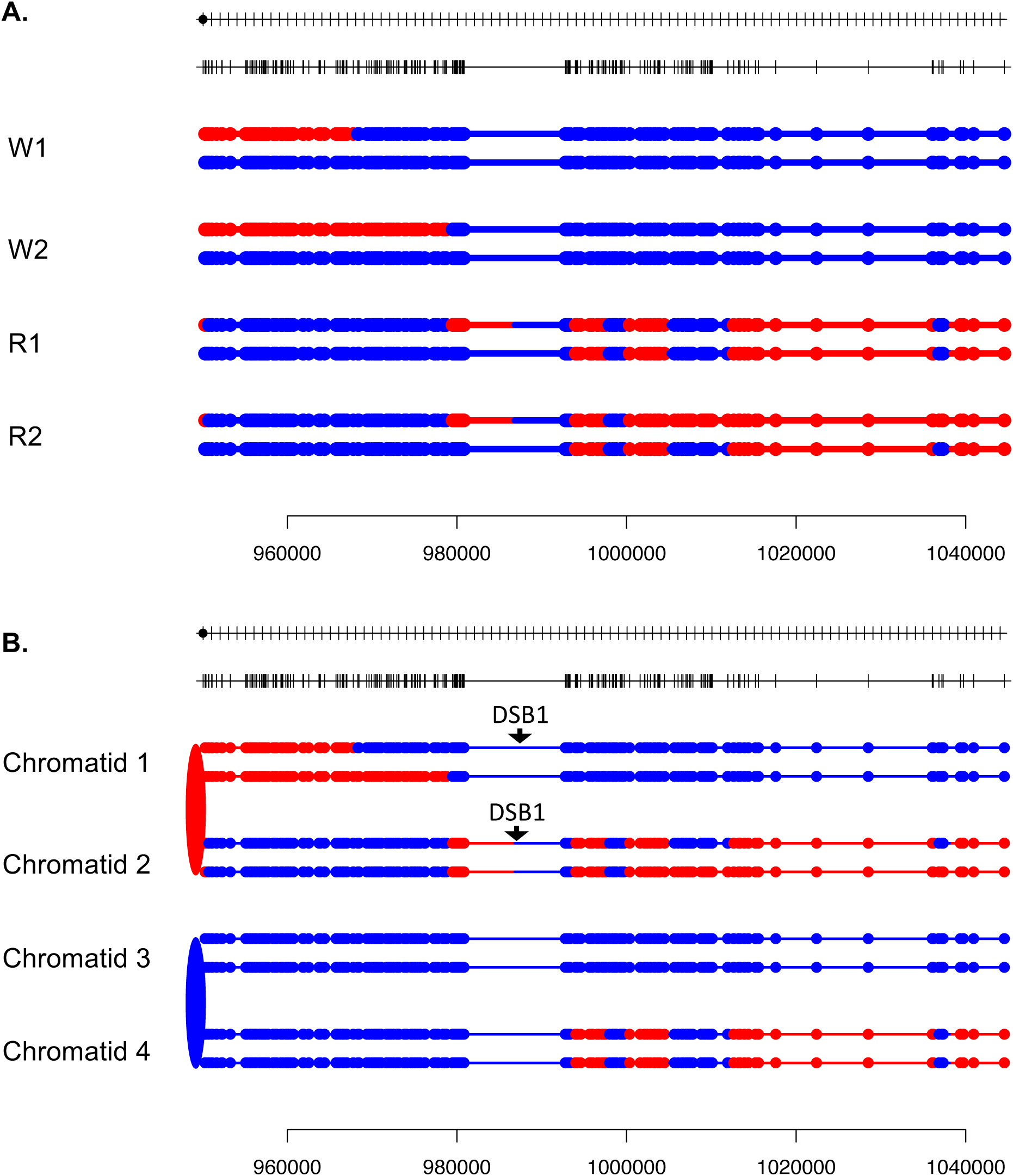

**Fig. S89.**
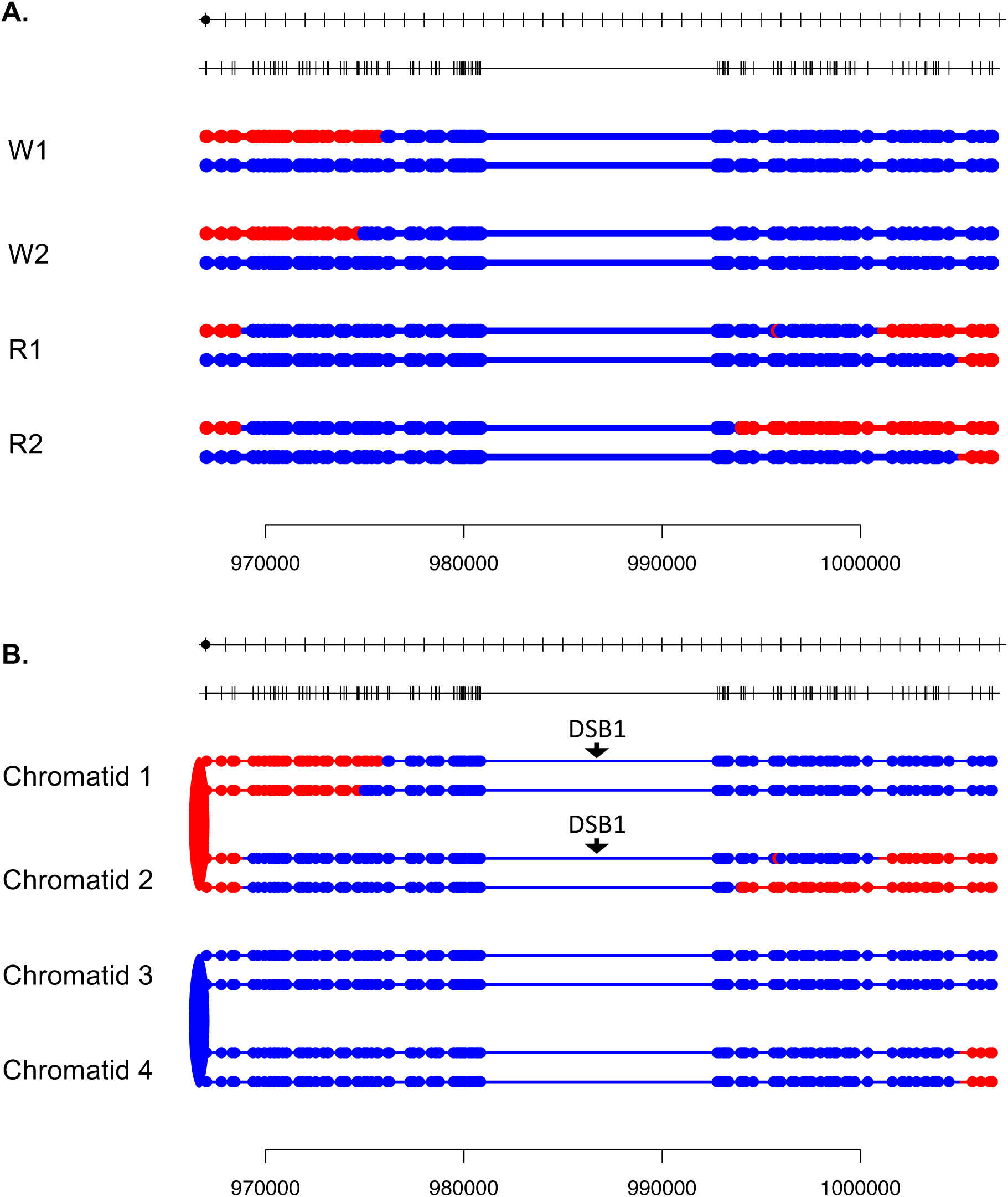

**Fig. S90.**
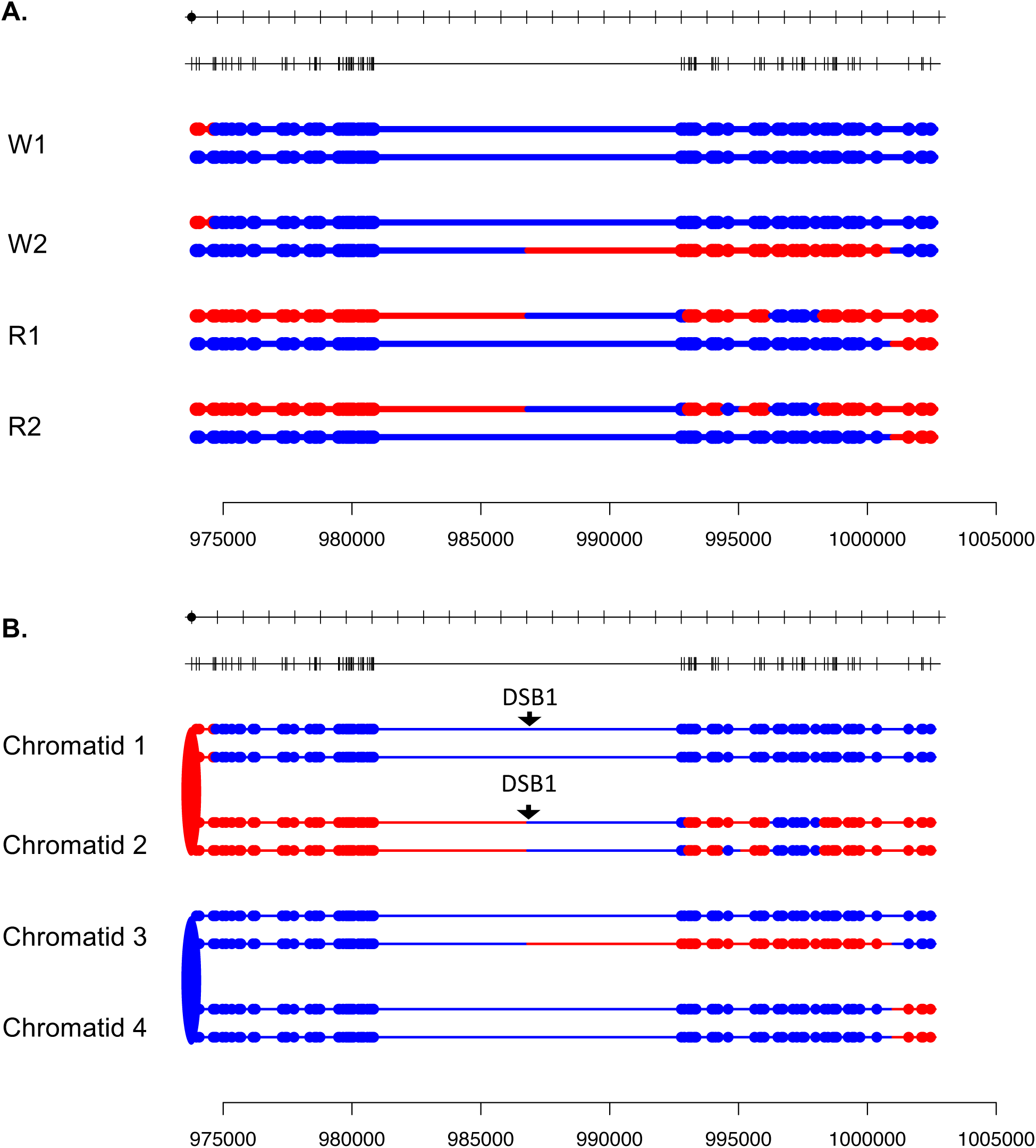

**Fig. S91.**
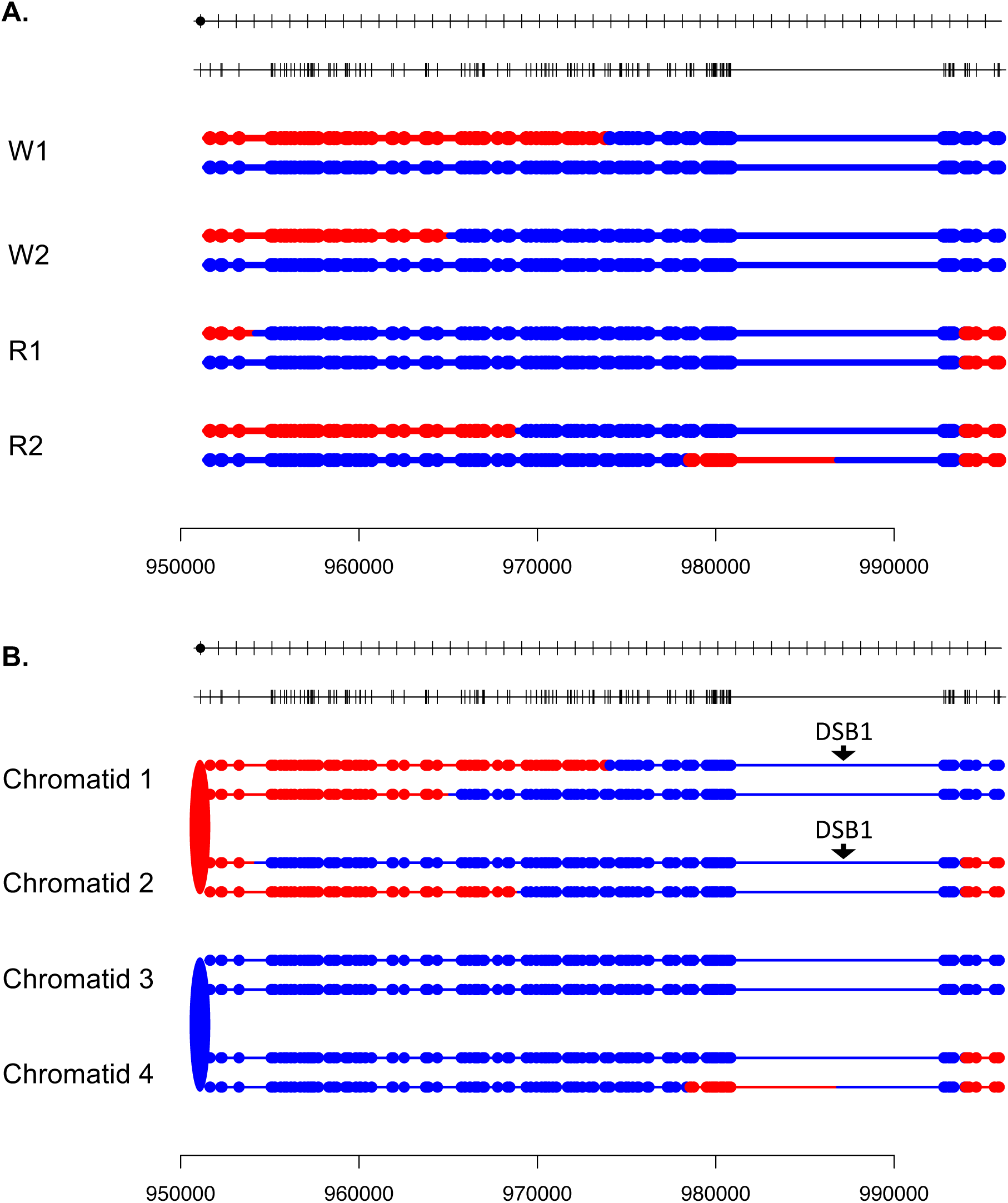

**Fig. S92-135.**
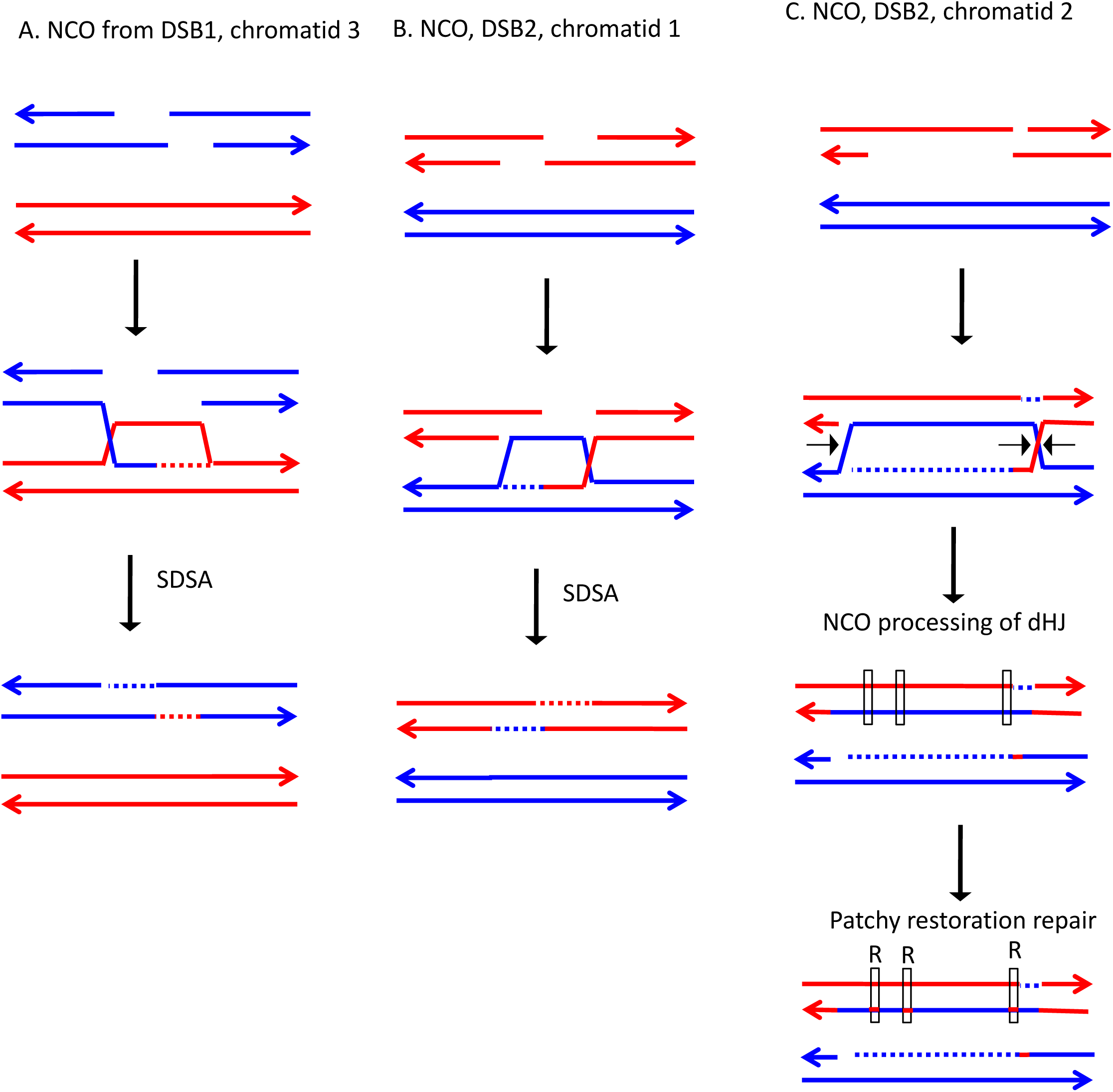
These figures show possible pathways of recombination to account for the pattern of the SNPs observed in the granddaughter cells. For events that involve the repair of two DSBs, we show each repair event separately. We show only one pathway for the repair of each DSB, although many repair events could be explained by alternative pathways. Fig. 92. Repair pathways for events shown in Fig. S3. Fig. 93. Repair pathways for events shown in Fig. S7. Fig. 94. Repair pathways for events shown in Fig. S8. Fig. 95. Repair pathways for events shown in Fig. S13. Fig. 96. Repair pathways for events shown in Fig. S16. Fig. 97. Repair pathways for events shown in Fig. S17. Fig. 98. Repair pathways for events shown in Fig. S24. Fig. 99. Repair pathways for events shown in Fig. S26. Fig. 100. Repair pathways for events shown in Fig. S28. Fig. 101. Repair pathways for events shown in Fig. S34. Fig. 102. Repair pathways for events shown in Fig. S38. Fig. 103. Repair pathways for events shown in Fig. S41. Fig. 104. Repair pathways for events shown in Fig. S43. Fig. 105. Repair pathways for events shown in Fig. S44. Fig. 106. Repair pathways for events shown in Fig. S45. Fig. 107. Repair pathways for events shown in Fig. S48. Fig. 108. Repair pathways for events shown in Fig. S53. Fig. 109. Repair pathways for events shown in Fig. S56. Fig. 110. Repair pathways for events shown in Fig. S59. Fig. 111. Repair pathways for events shown in Fig. S60. Fig. 112. Repair pathways for events shown in Fig. S61. Fig. 113. Repair pathways for events shown in Fig. S62. Fig. 114. Repair pathways for events shown in Fig. S64. Fig. 115. Repair pathways for events shown in Fig. S65. Fig. 116. Repair pathways for events shown in Fig. S66. Fig. 117. Repair pathways for events shown in Fig. S67. Fig. 118. Repair pathways for events shown in Fig. S70. Fig. 119. Repair pathways for events shown in Fig. S72. Fig. 120. Repair pathways for events shown in Fig. S74. Fig. 121. Repair pathways for events shown in Fig. S76. Fig. 122. Repair pathways for events shown in Fig. S77. Fig. 123. Repair pathways for events shown in Fig. S78. Fig. 124. Repair pathways for events shown in Fig. S79. Fig. 125. Repair pathways for events shown in Fig. S81. Fig. 126. Repair pathways for events shown in Fig. S82. Fig. 127. Repair pathways for events shown in Fig. S83. Fig. 128. Repair pathways for events shown in Fig. S84. Fig. 129. Repair pathways for events shown in Fig. S85. Fig. 130. Repair pathways for events shown in Fig. S86. Fig. 131. Repair pathways for events shown in Fig. S87. Fig. 132. Repair pathways for events shown in Fig. S88. Fig. 133. Repair pathways for events shown in Fig. S89. Fig. 134. Repair pathways for events shown in Fig. S90. Fig. 135. Repair pathways for events shown in Fig. S91.

**Fig. S93.**
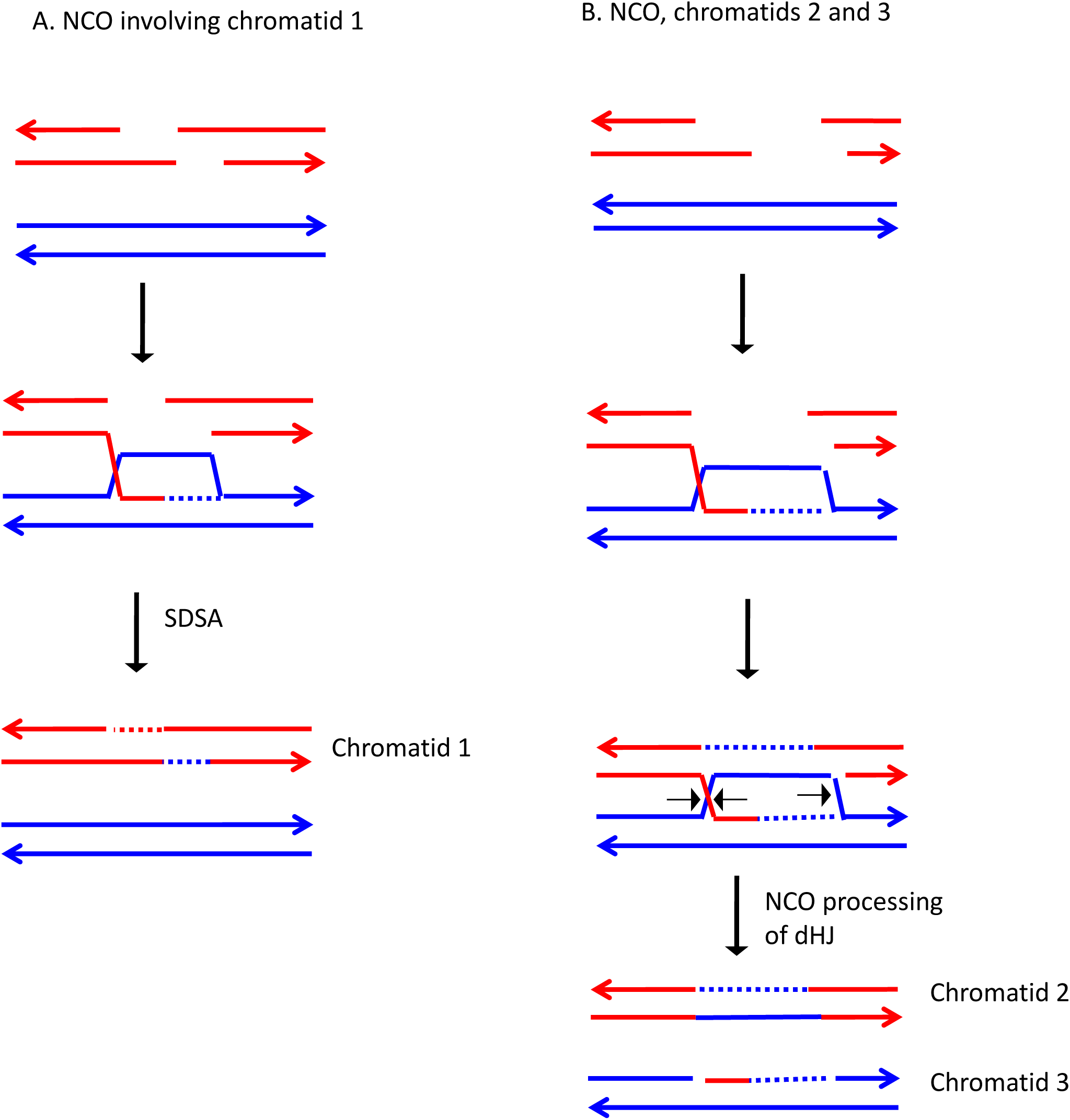

**Fig. S94.**
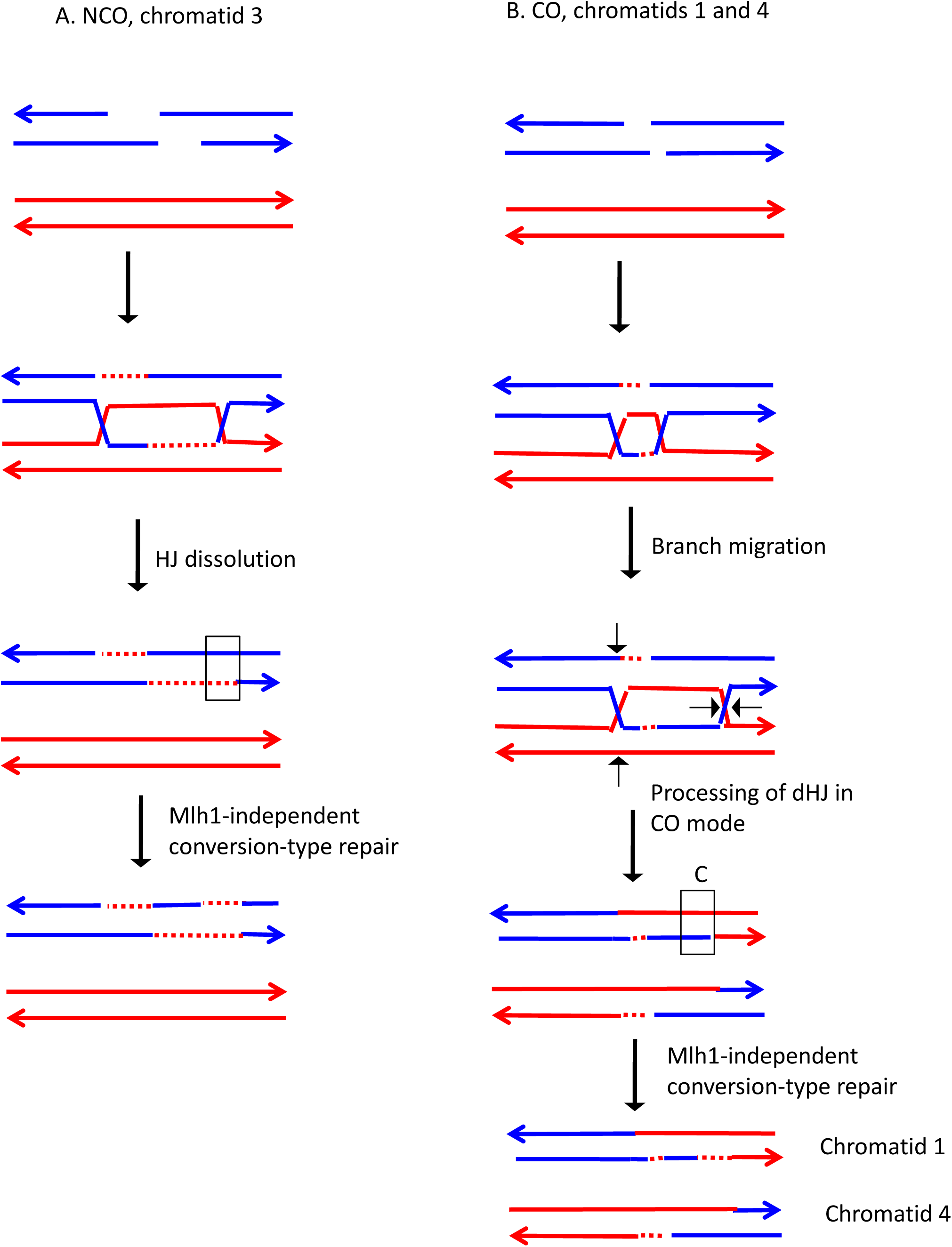

**Fig. S95.**
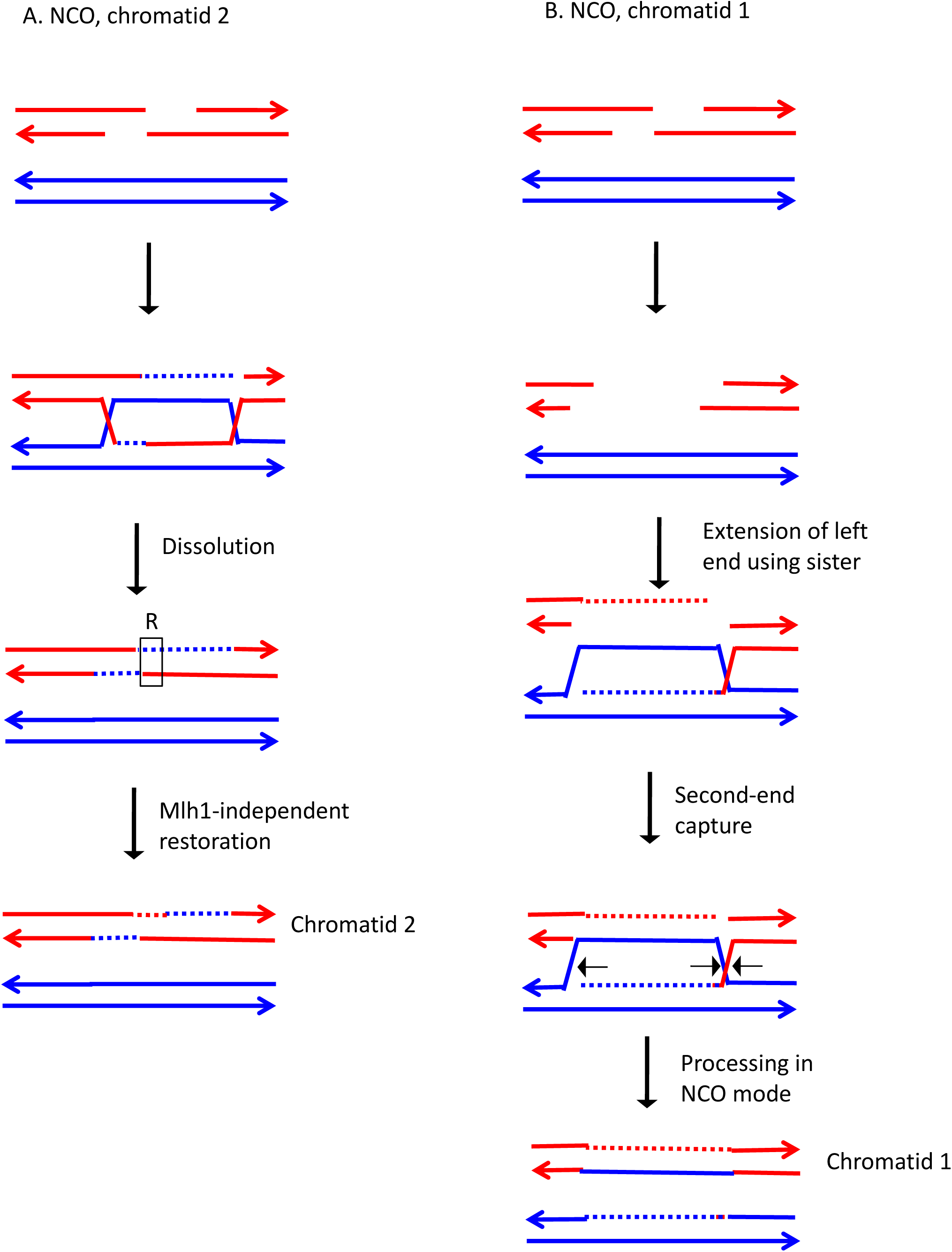

**Fig. S96.**
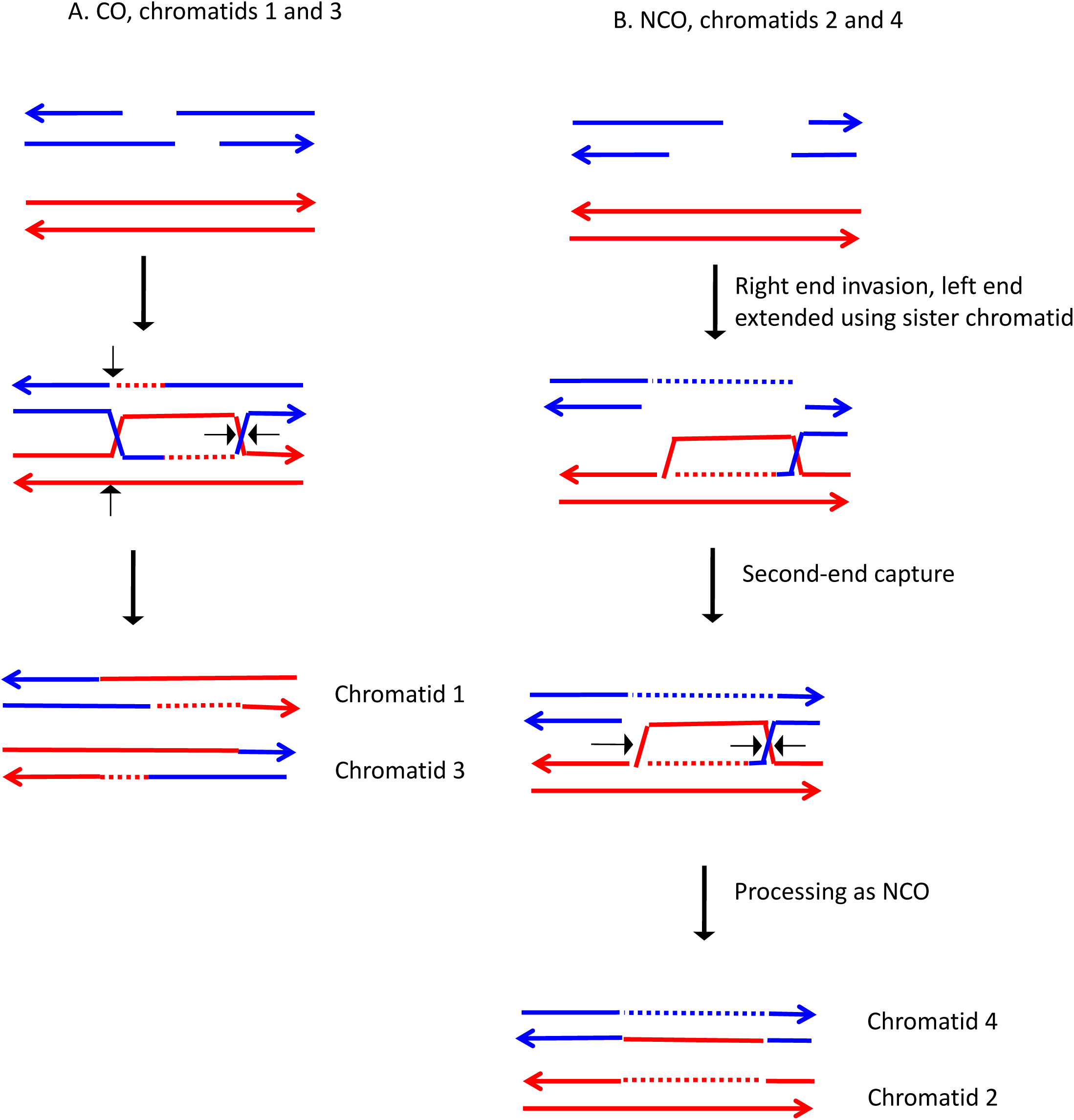

**Fig. S97.**
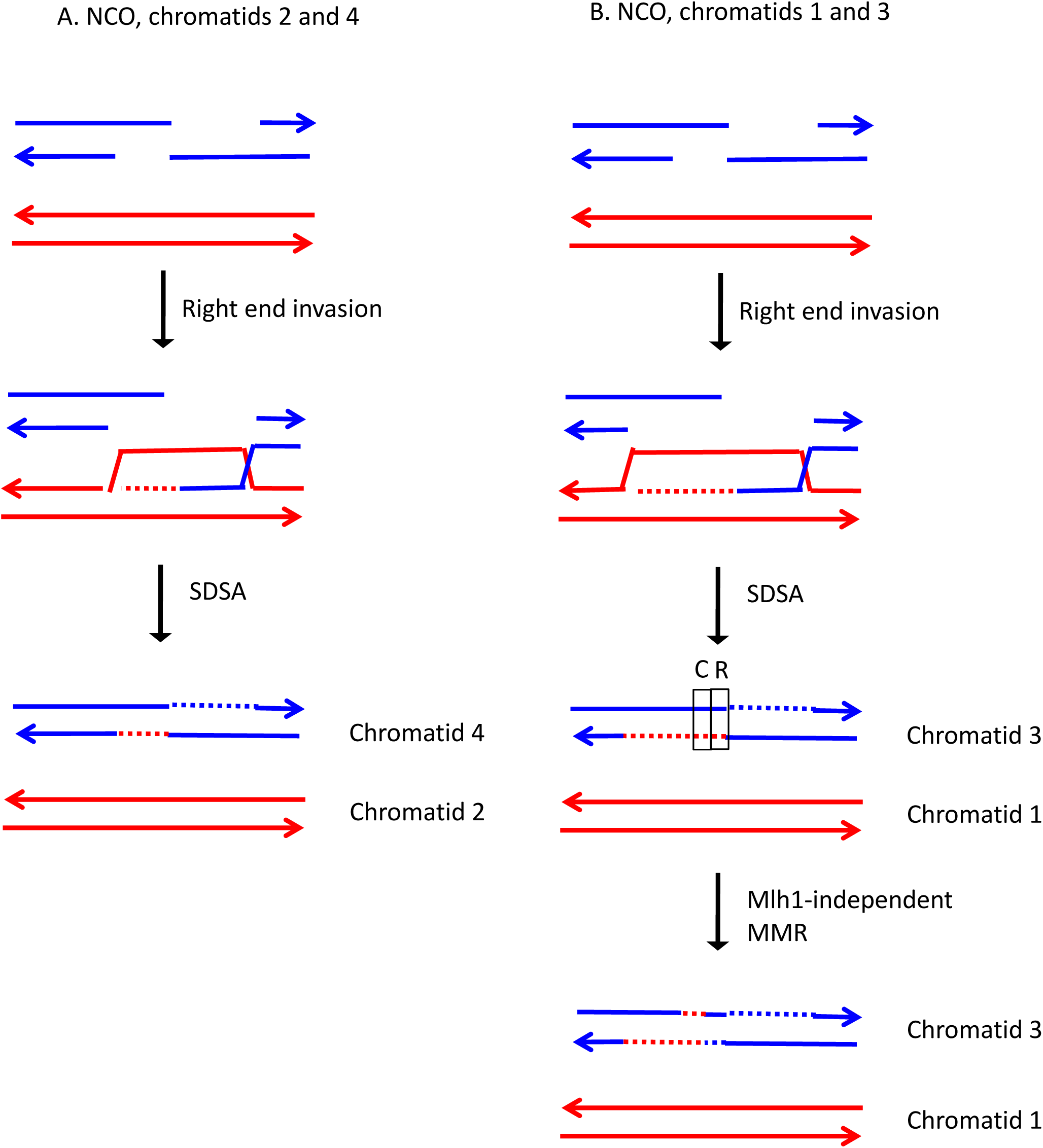

**Fig. S98.**
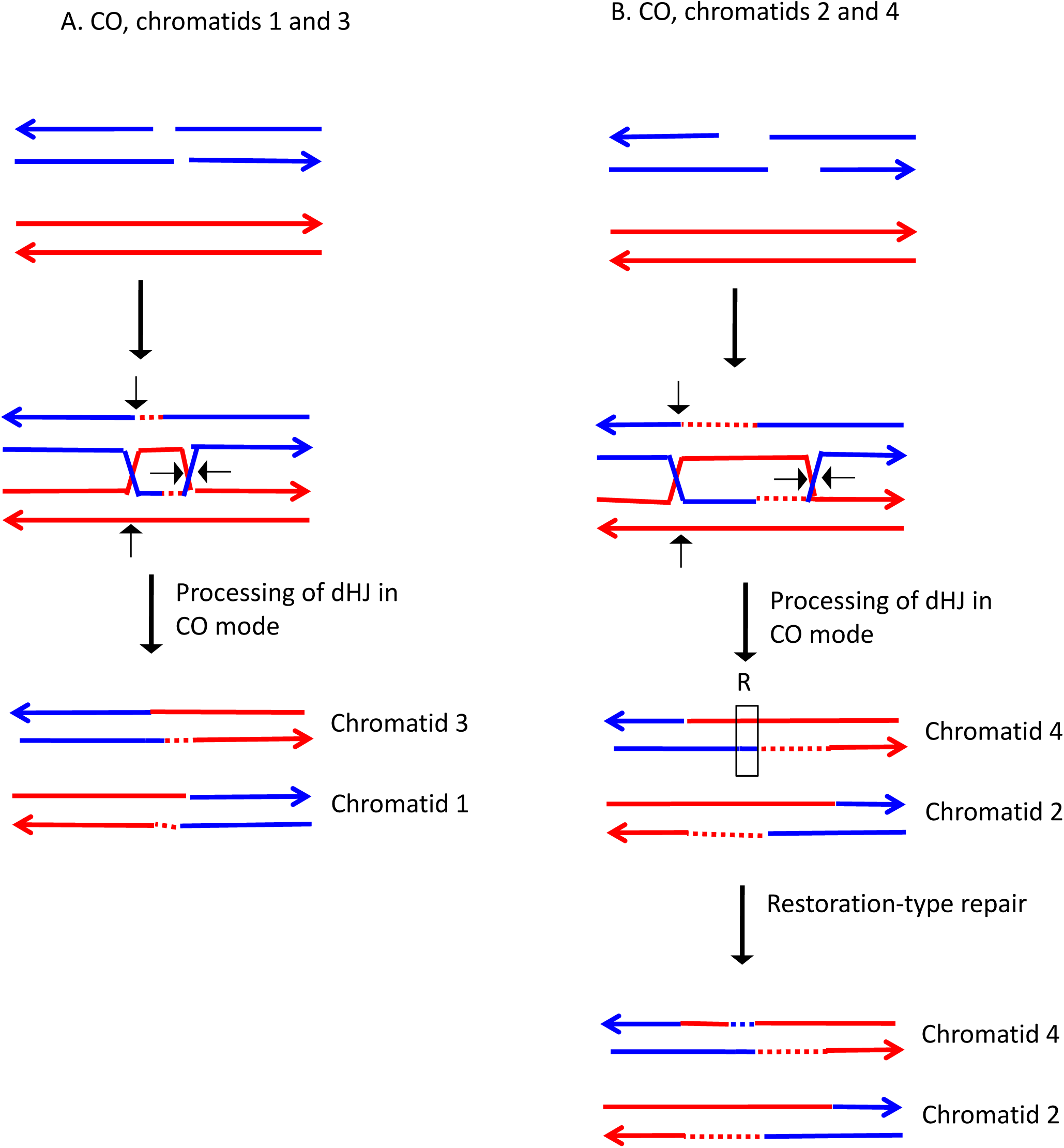

**Fig. S99.**
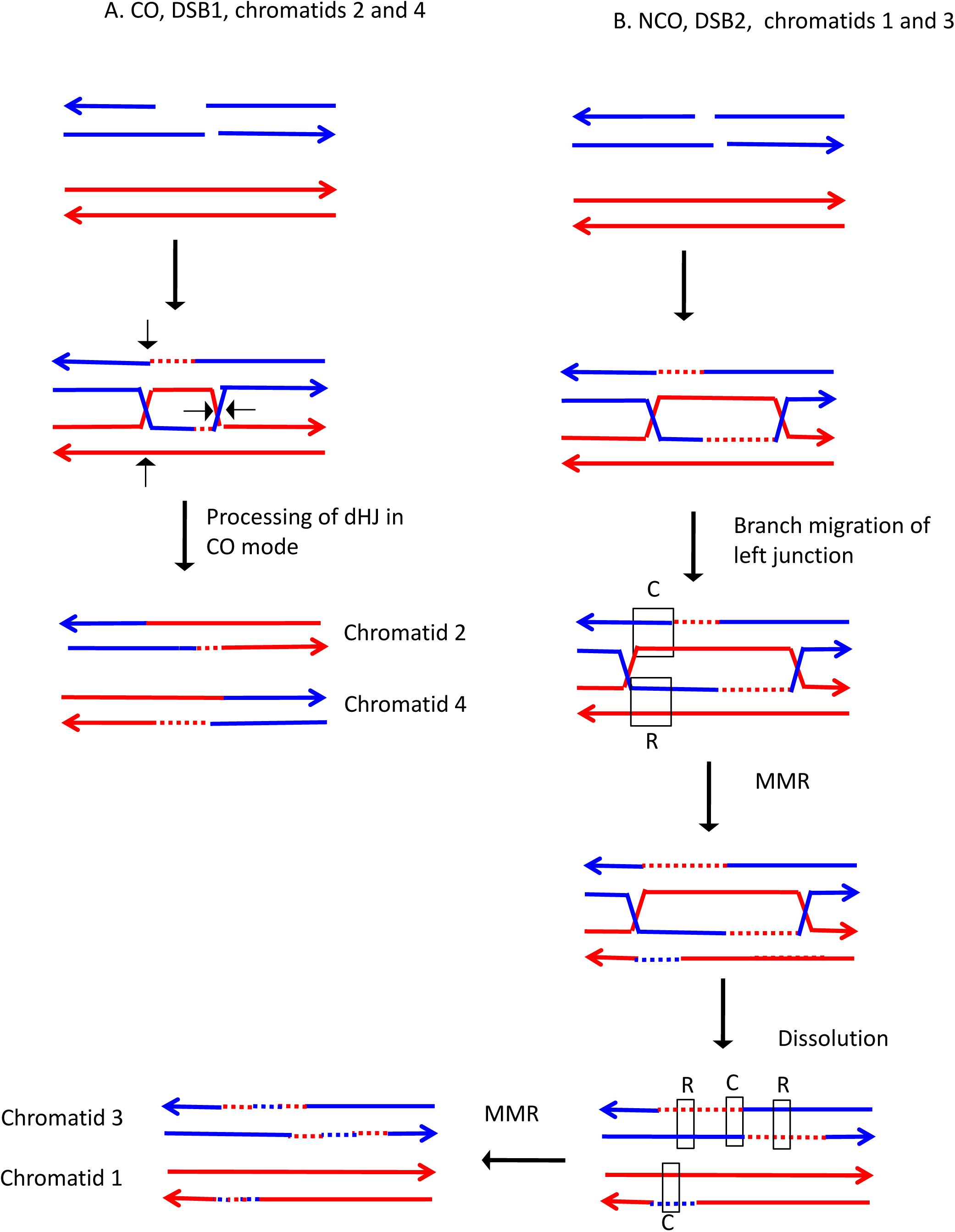

**Fig. S100.**
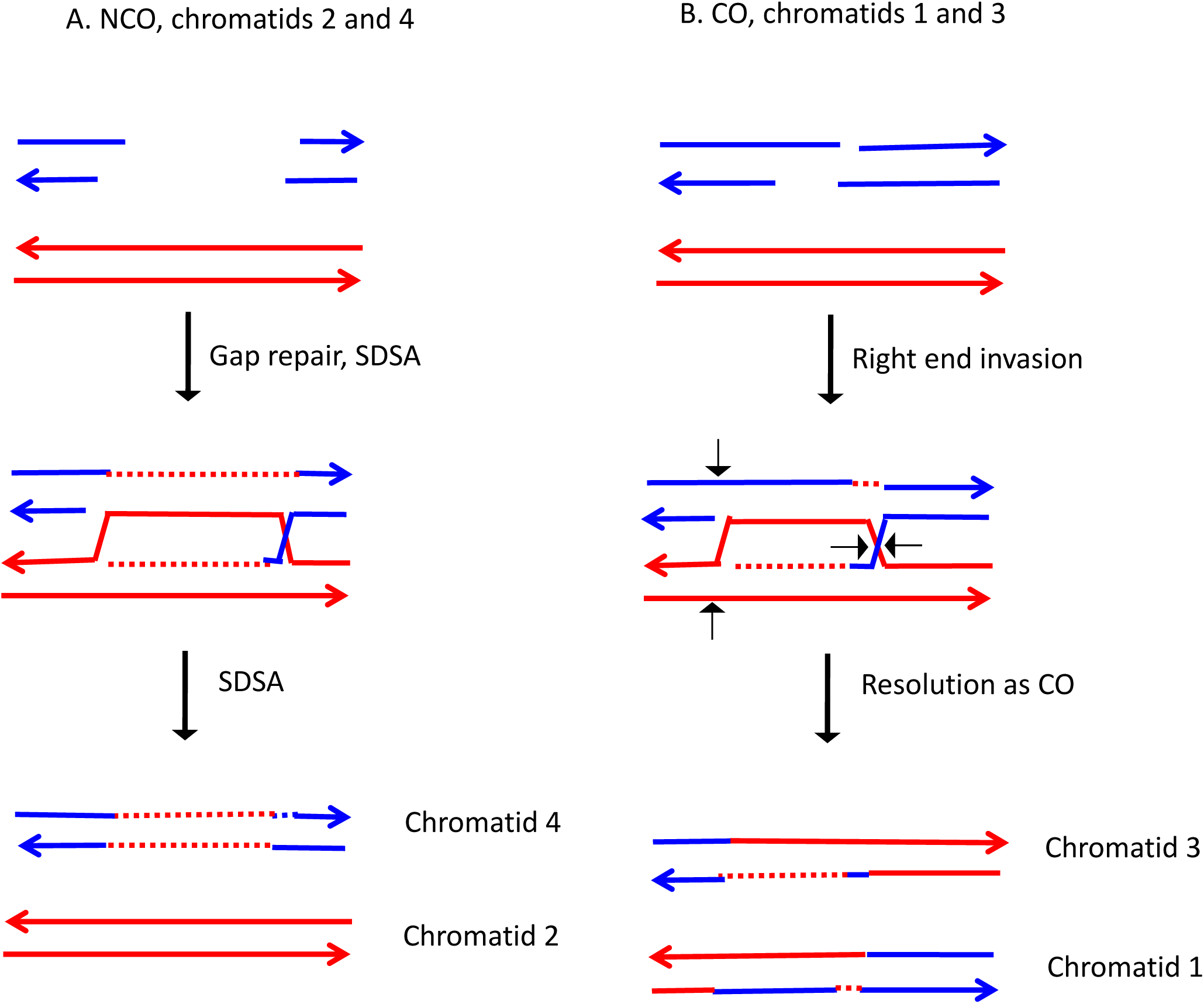

**Fig. S101.**
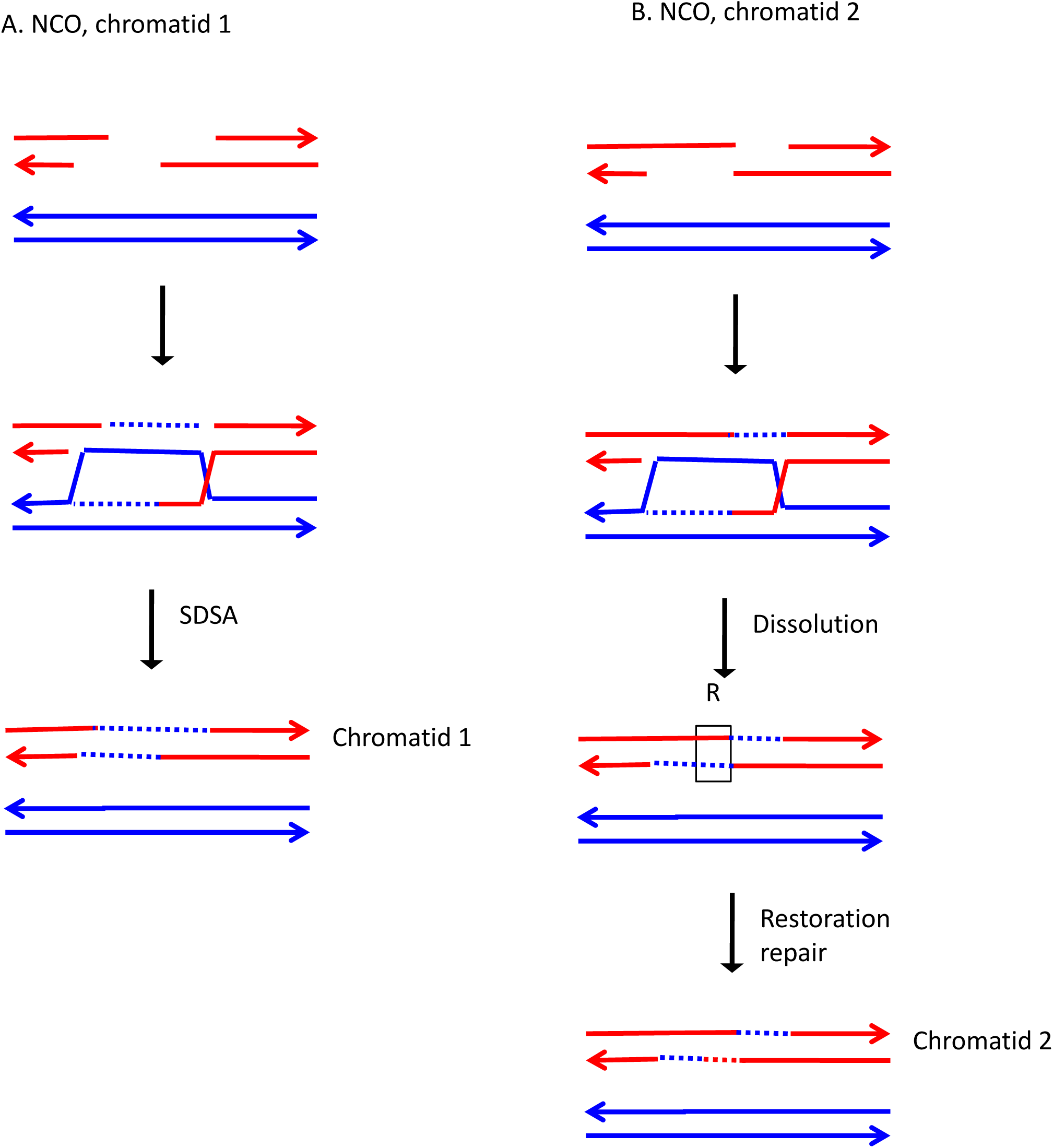

**Fig. S102.**
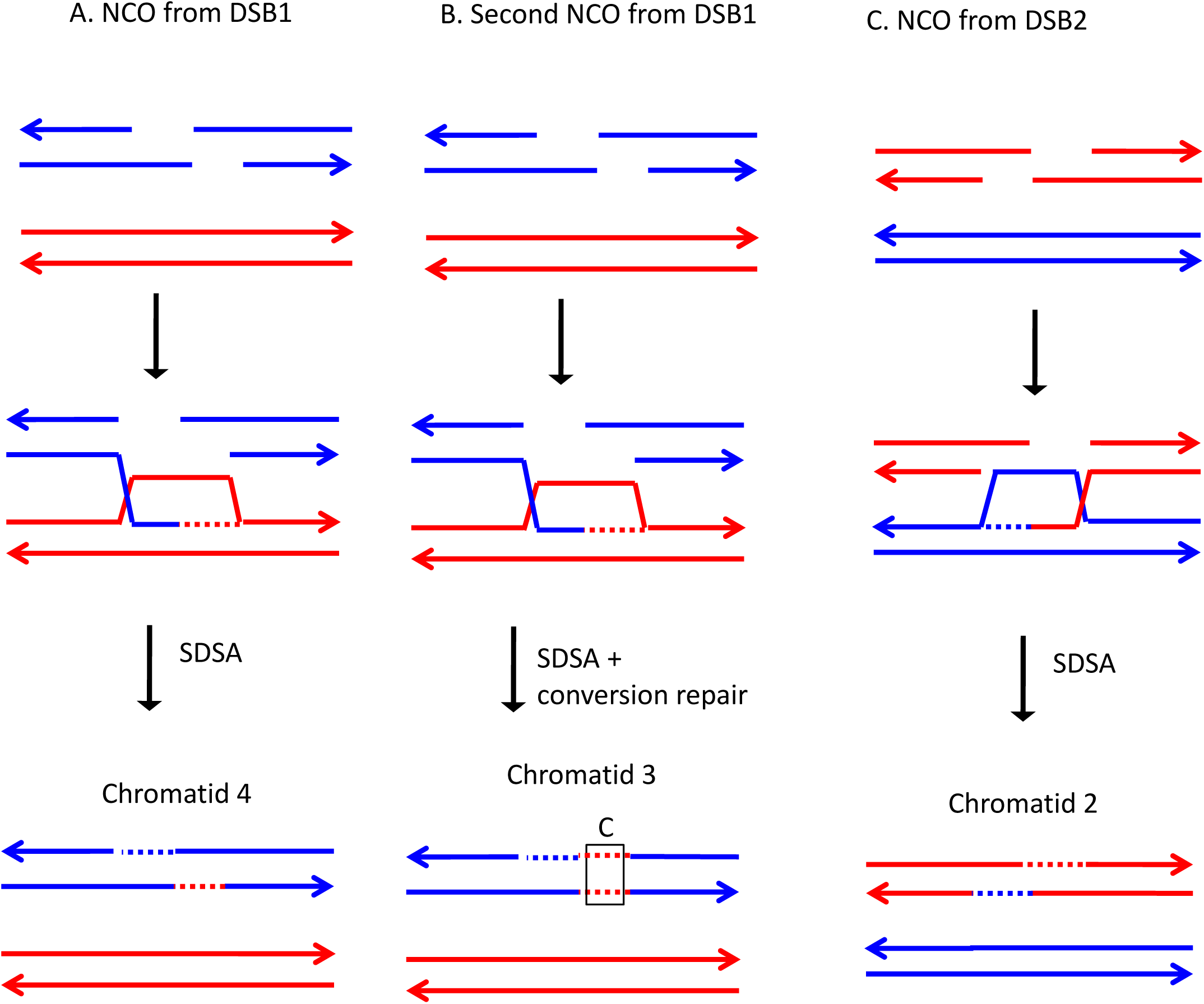

**Fig. S103.**
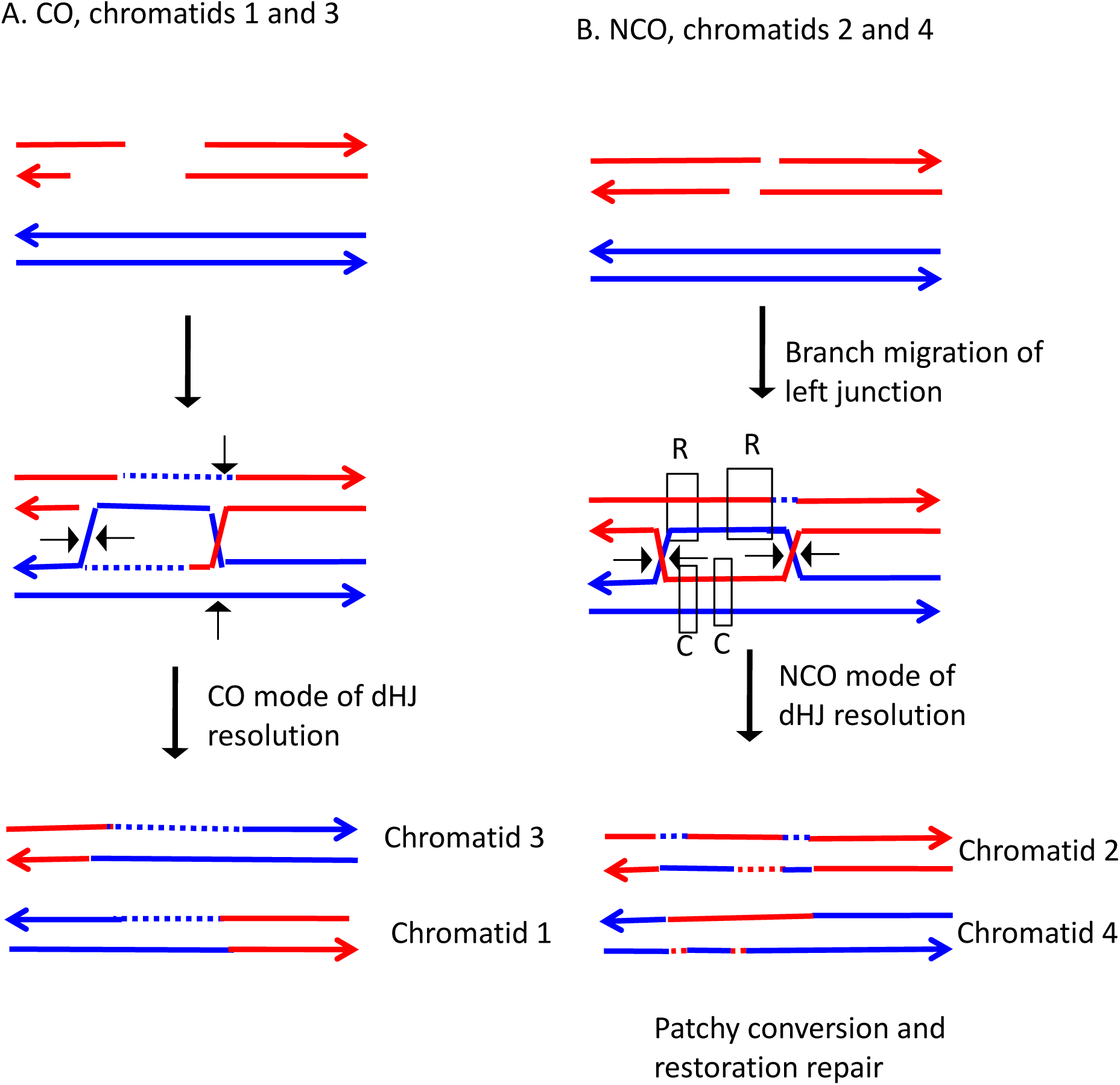

**Fig. S104.**
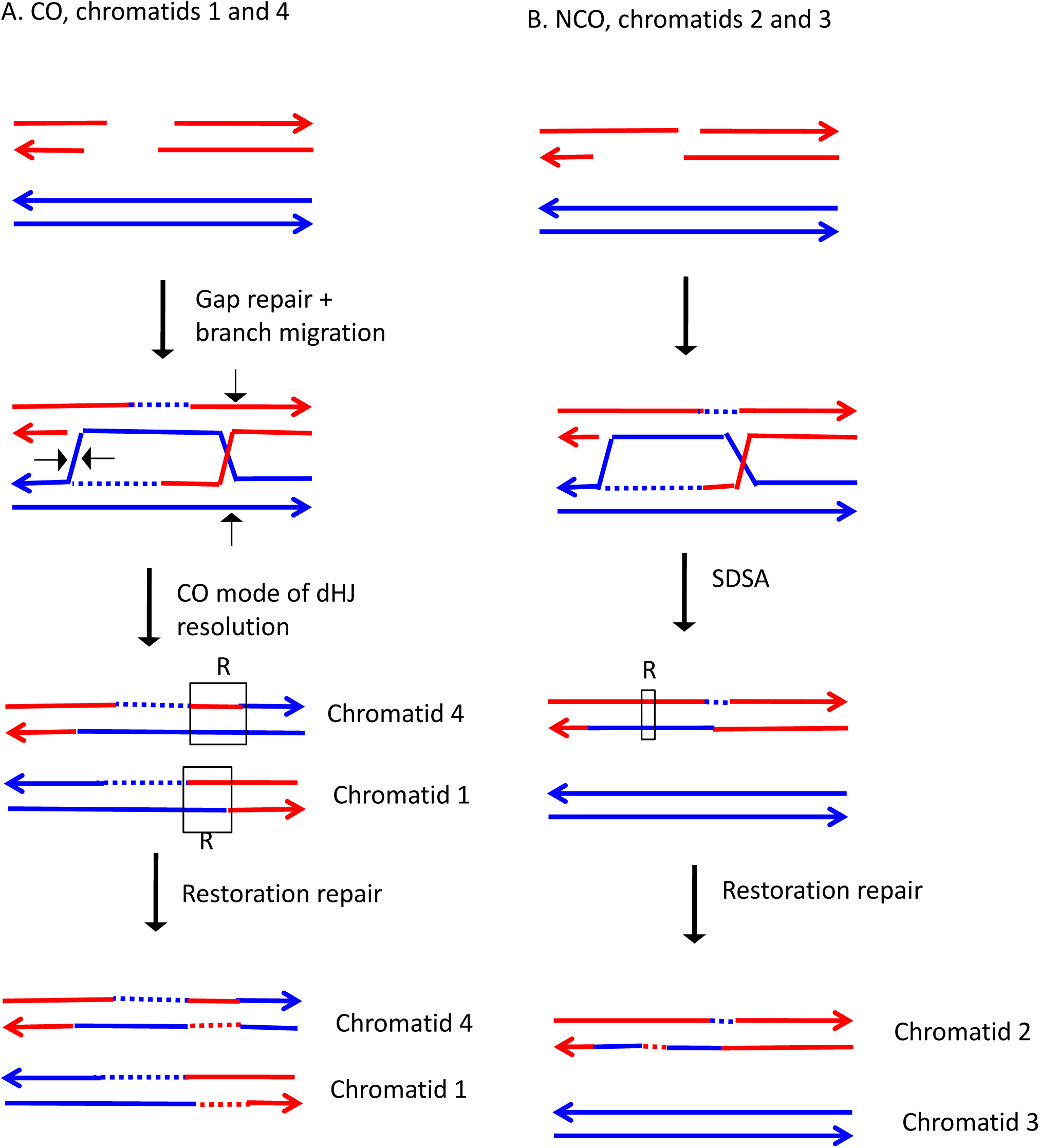

**Fig. S105.**
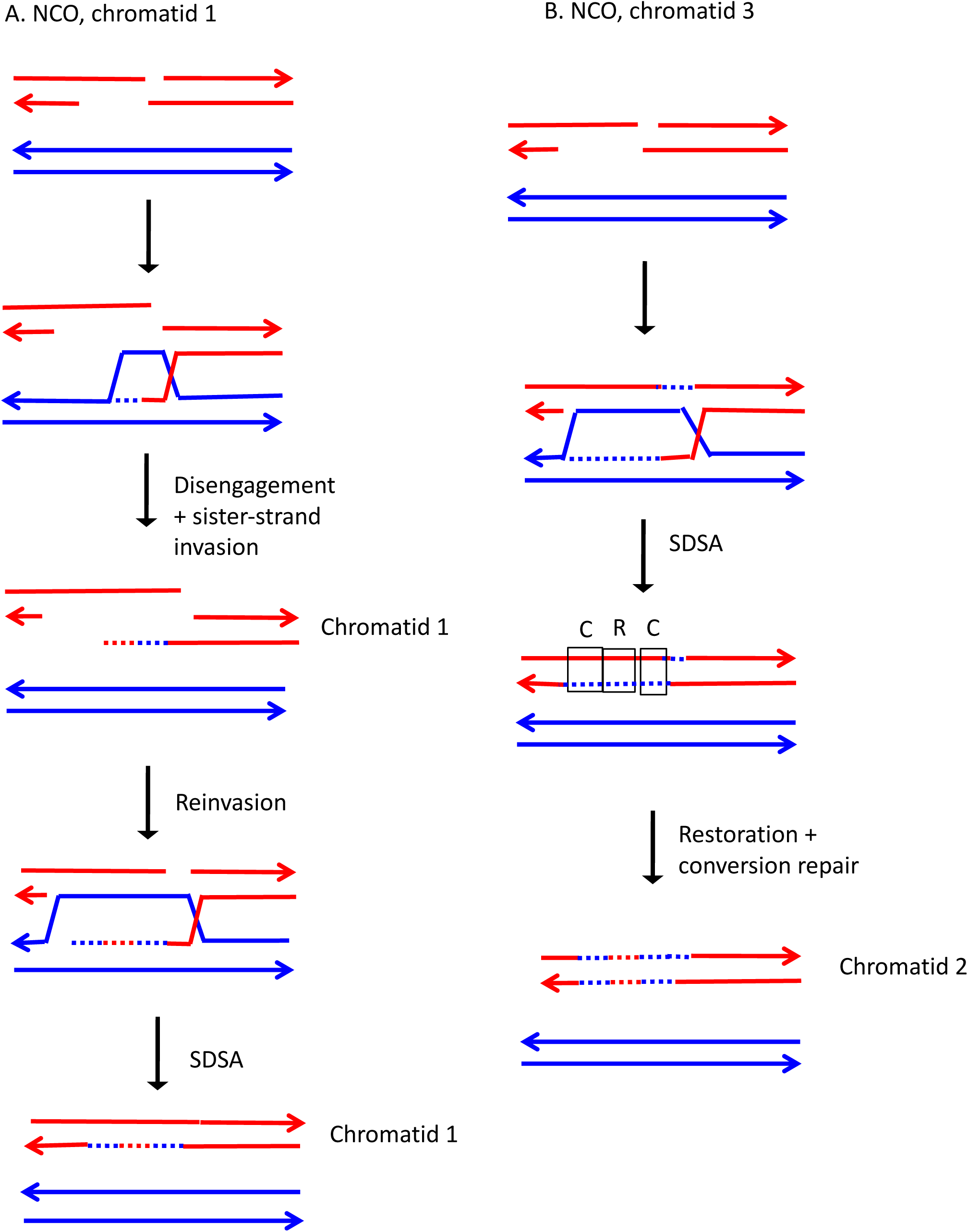

**Fig. S106.**
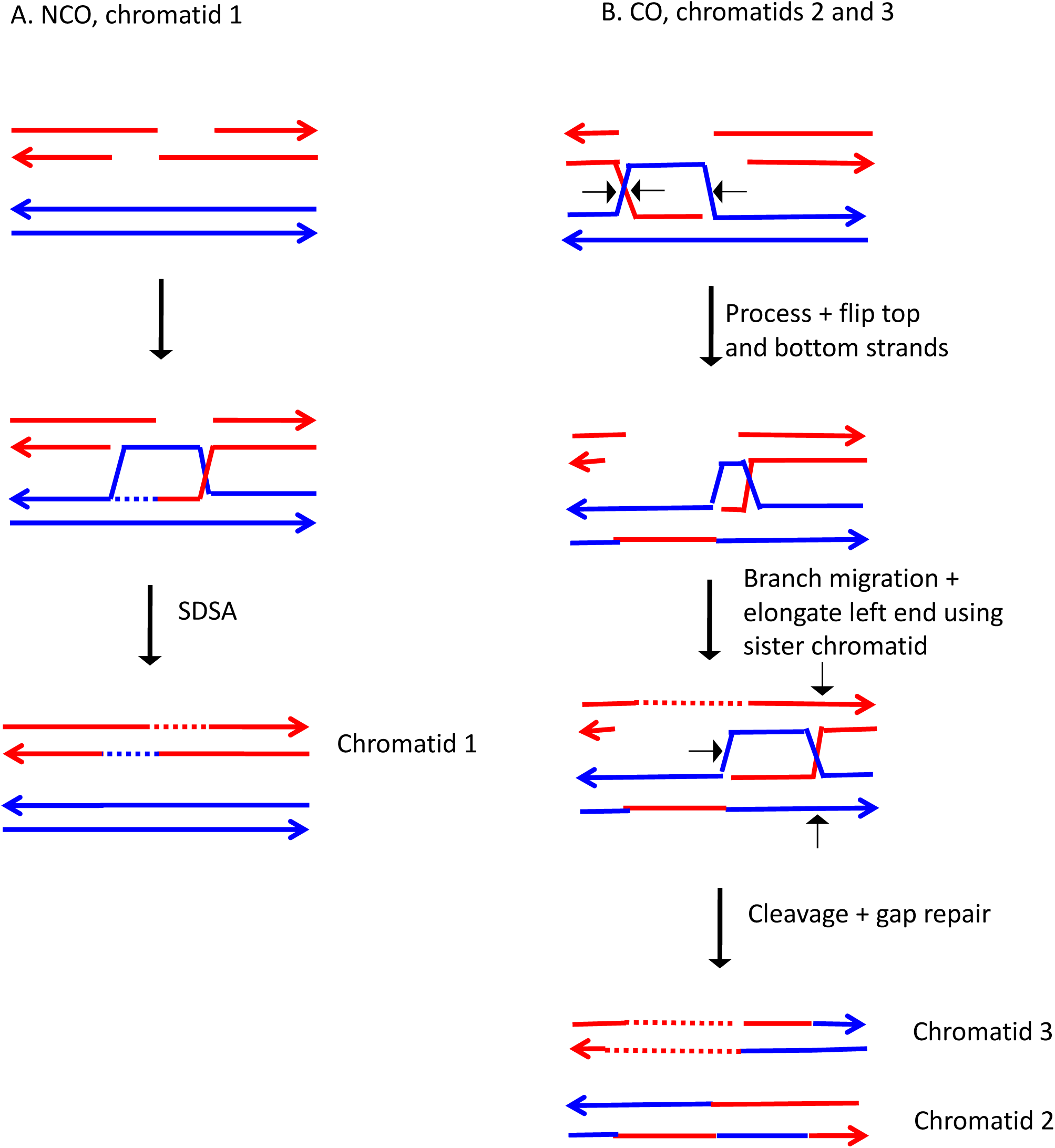

**Fig. S107.**
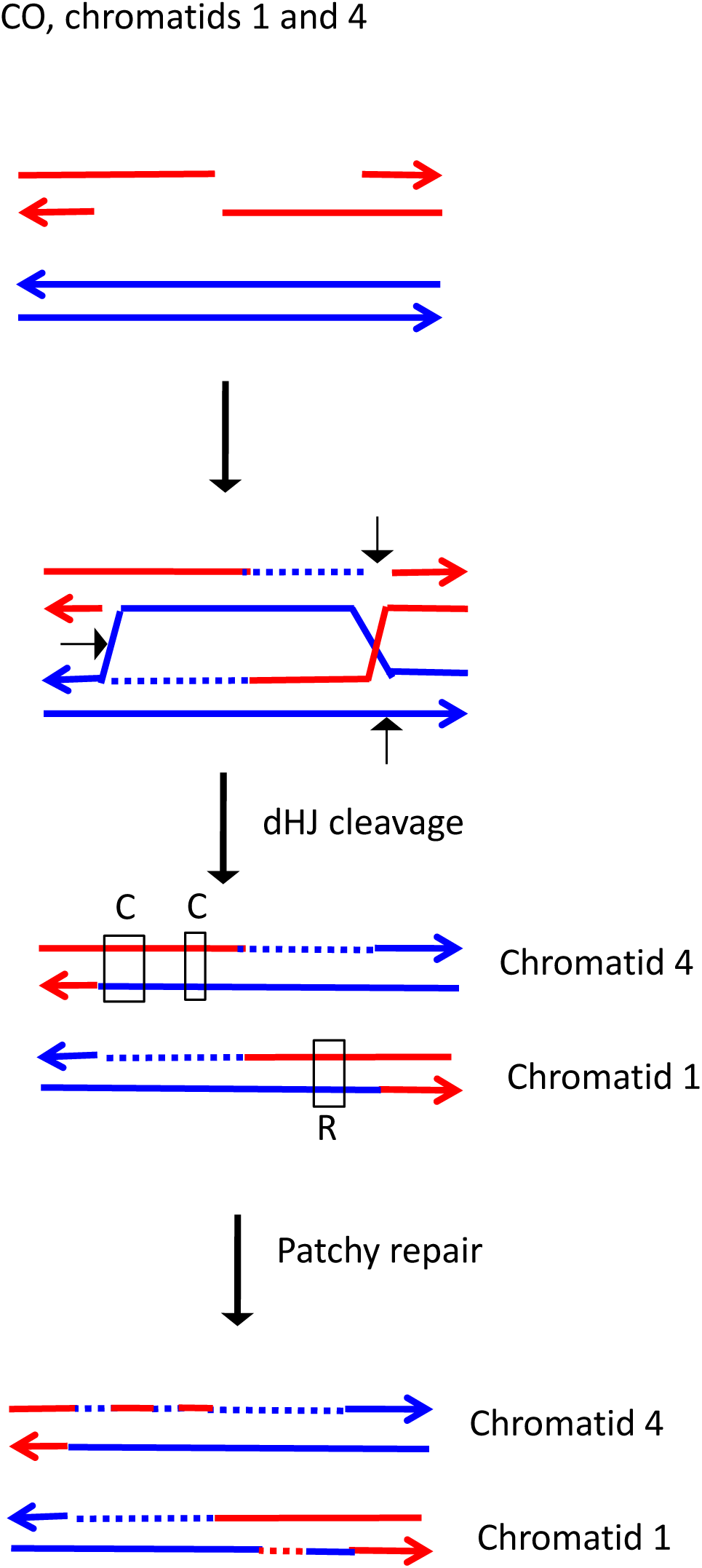

**Fig. S108.**
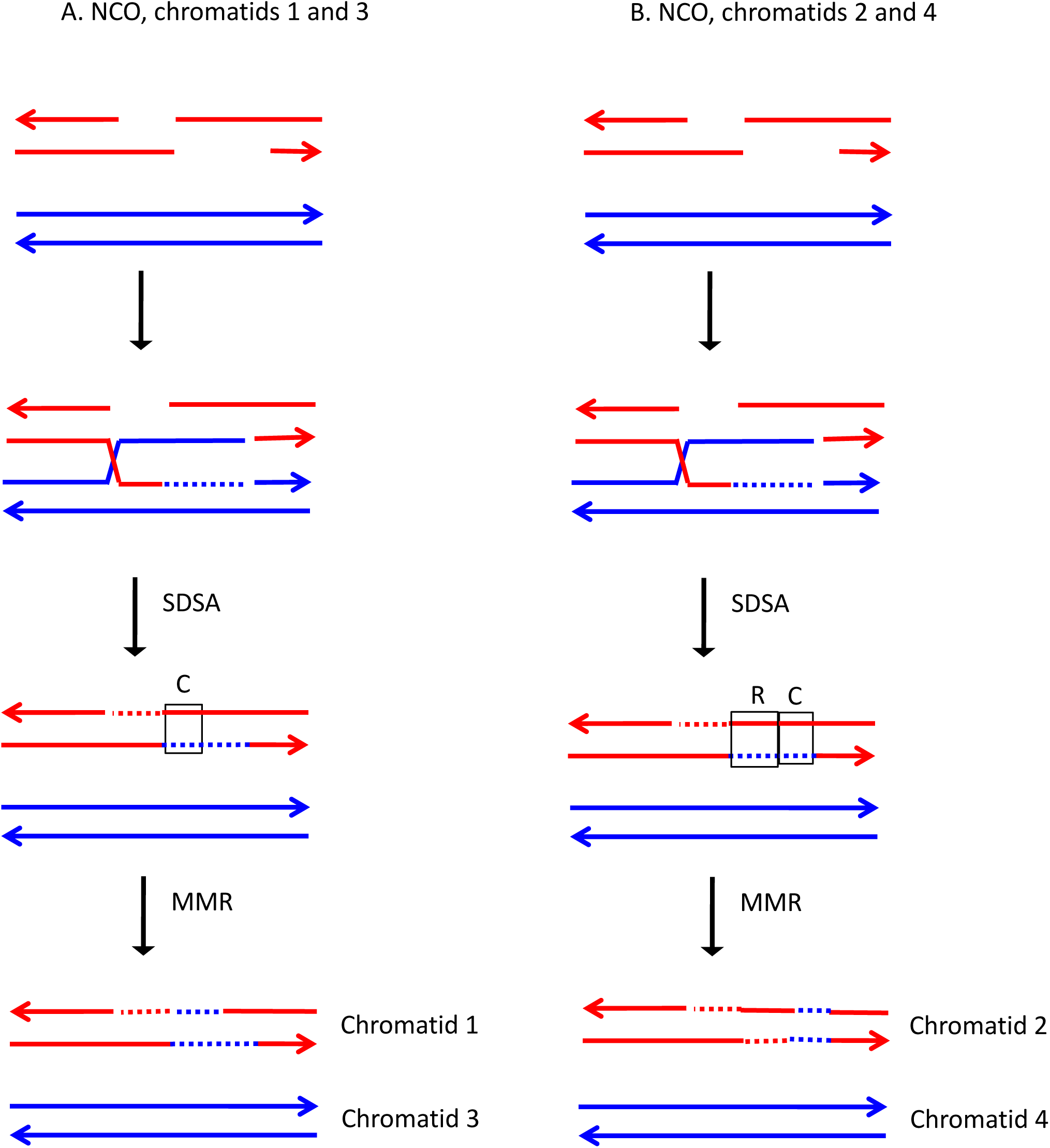

**Fig. S109.**
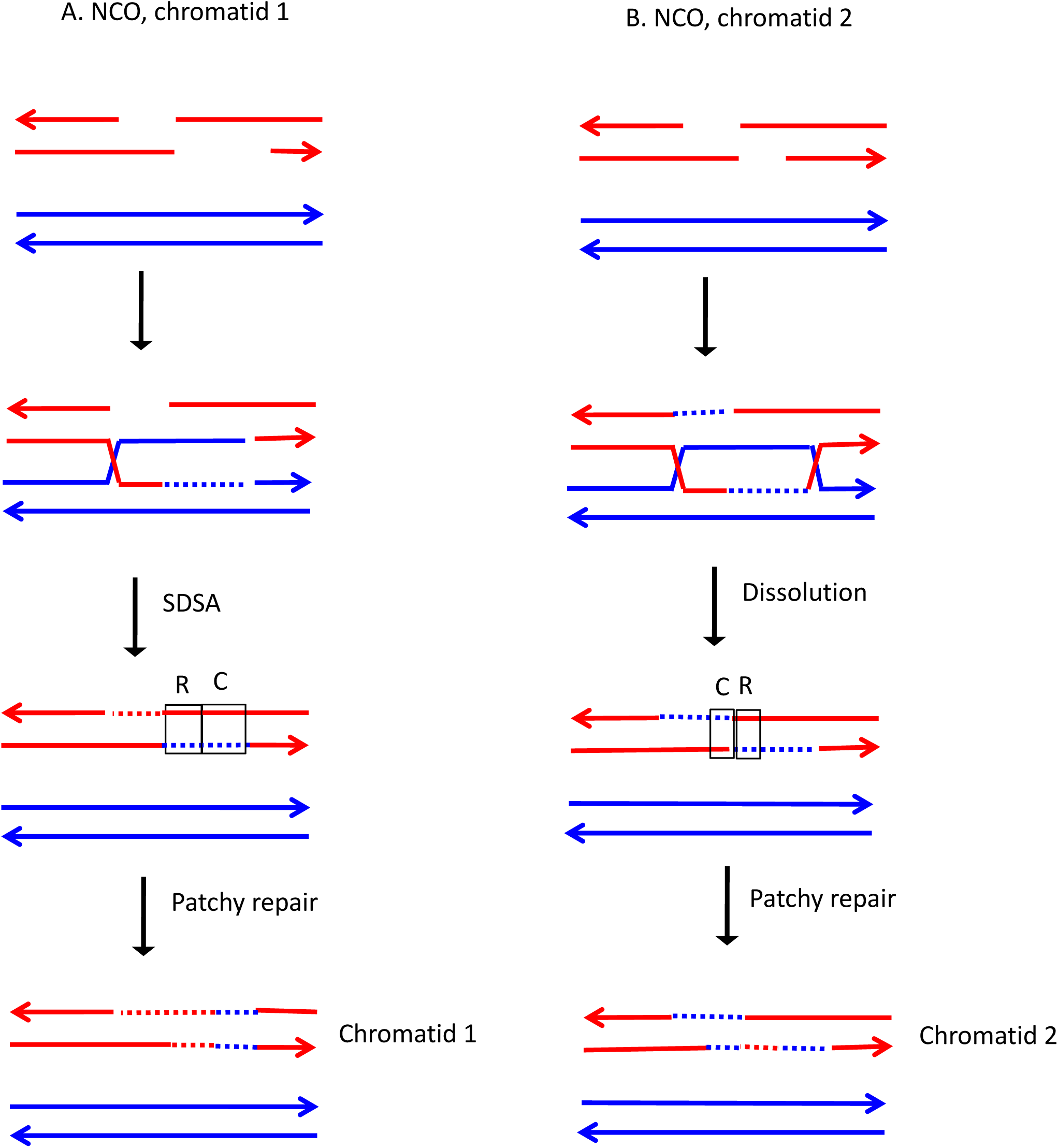

**Fig. S110.**
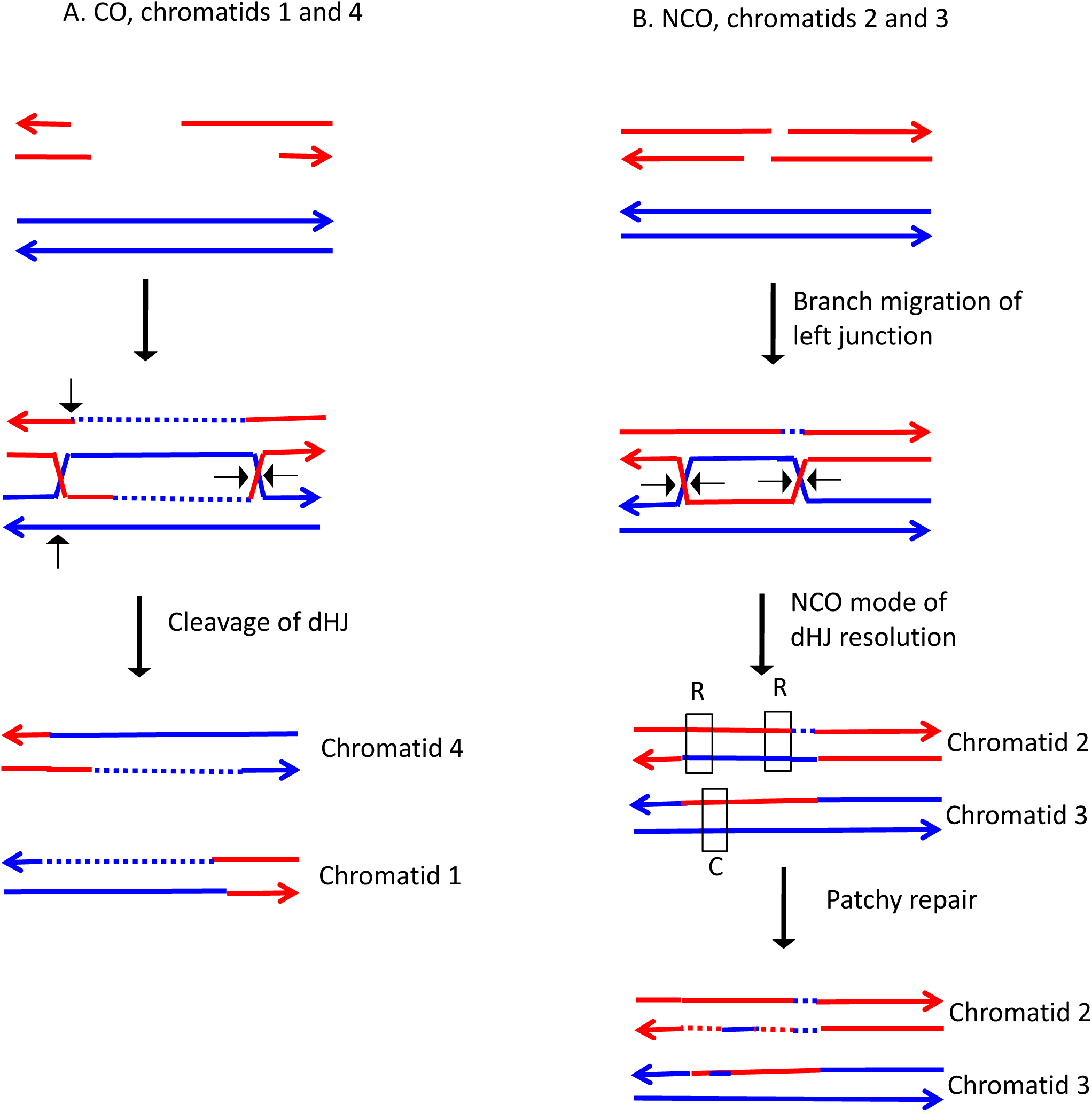

**Fig. S111.**
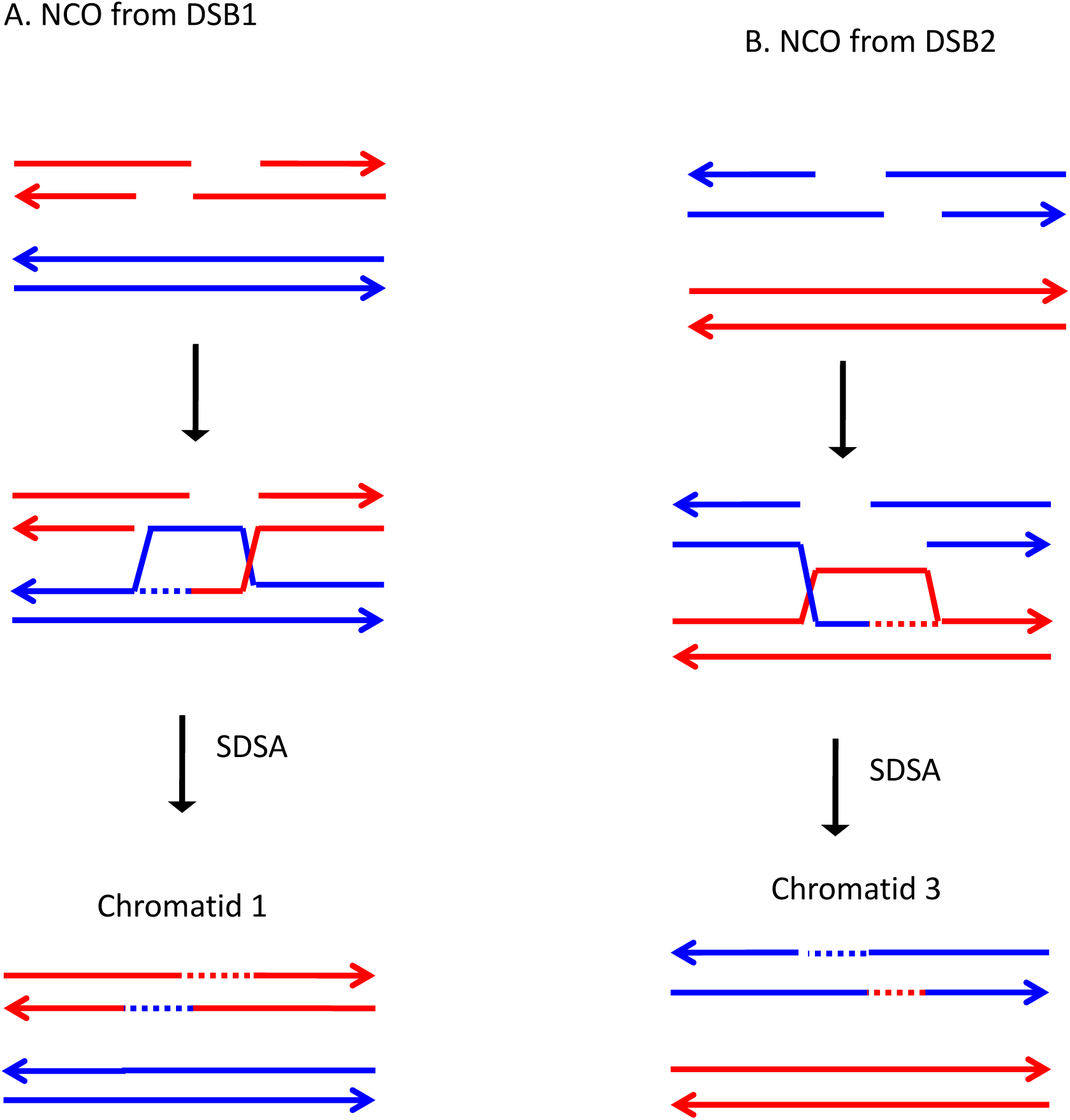

**Fig. S112.**
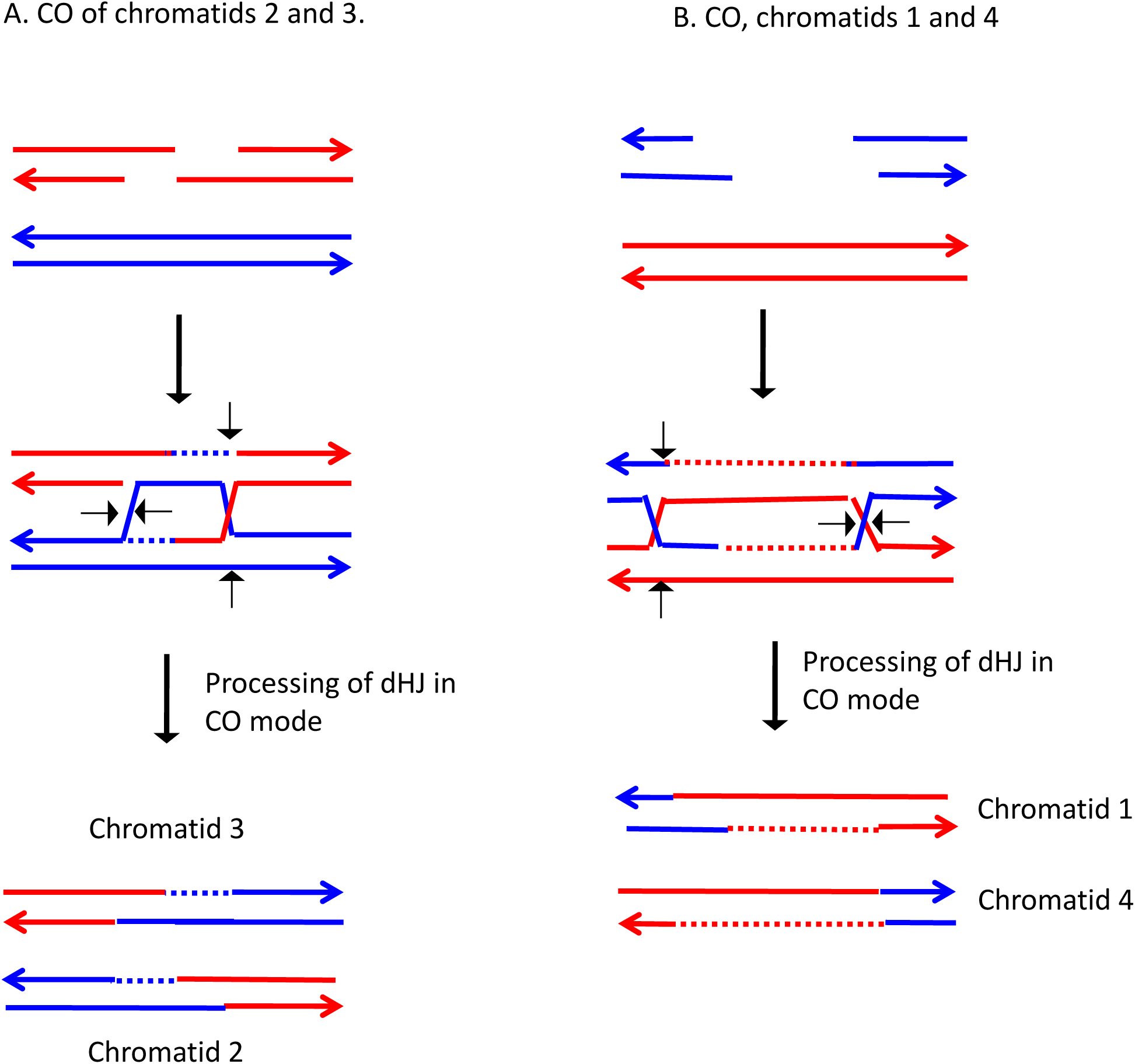

**Fig. S113.**
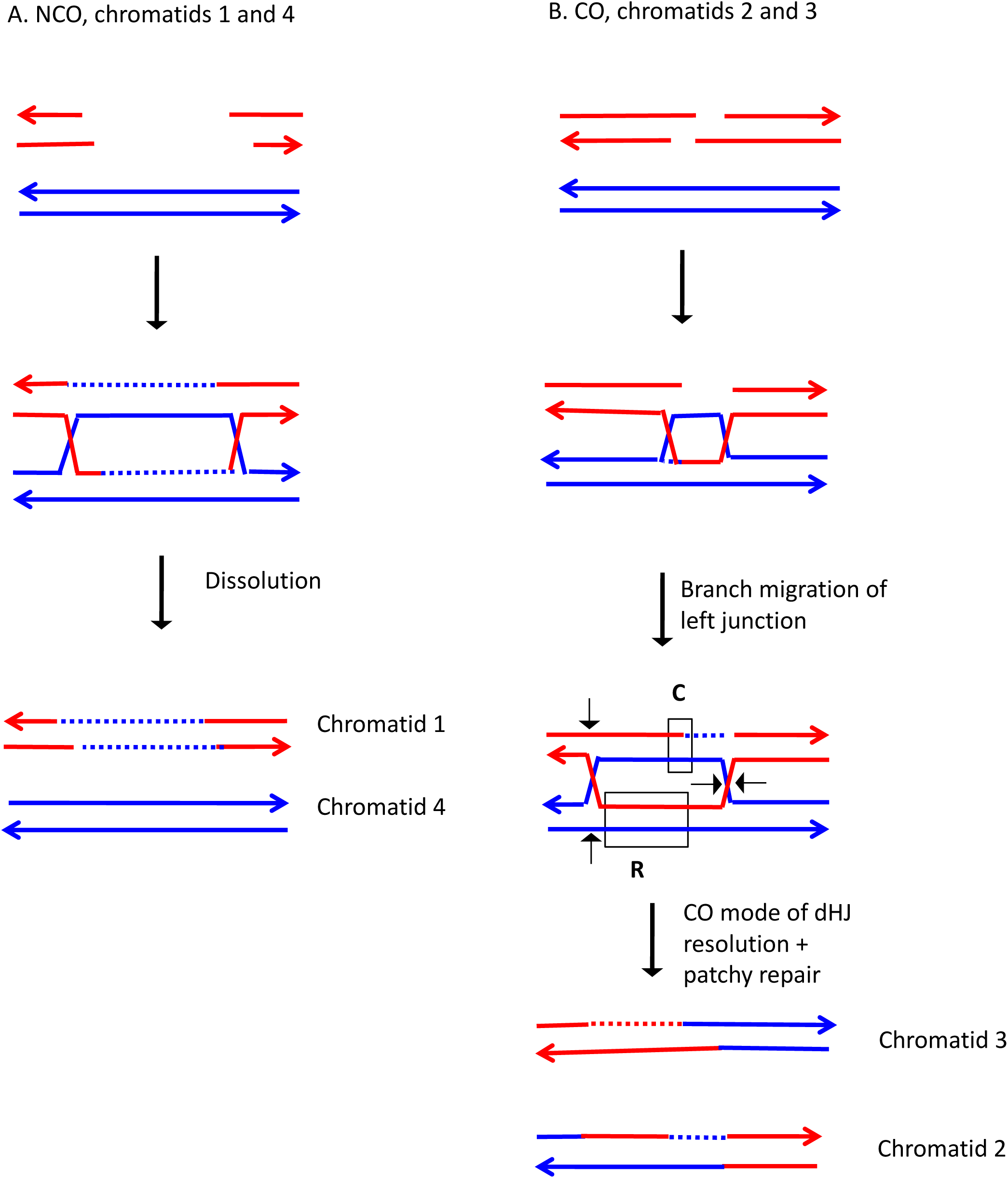

**Fig. S114.**
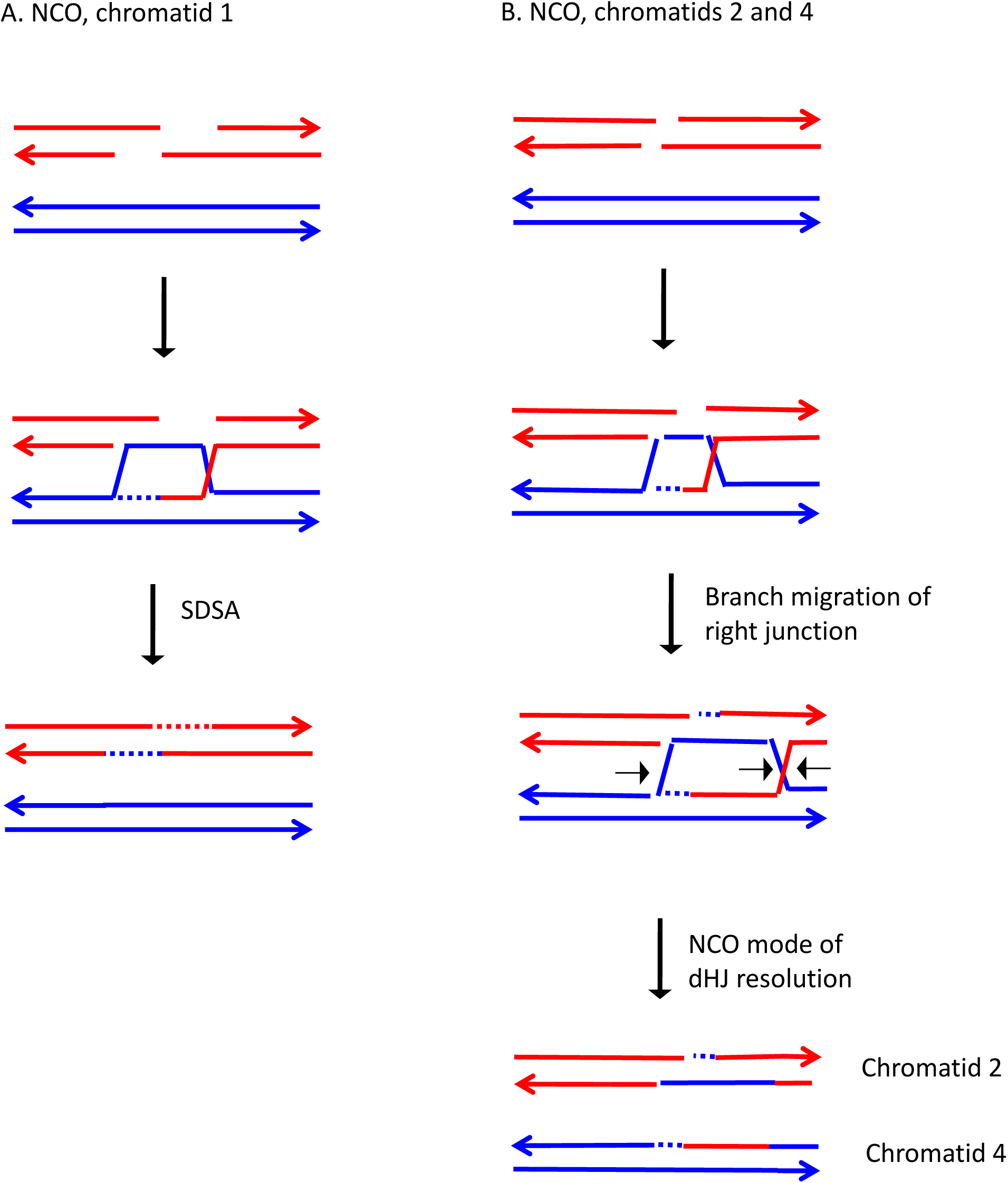

**Fig. S115.**
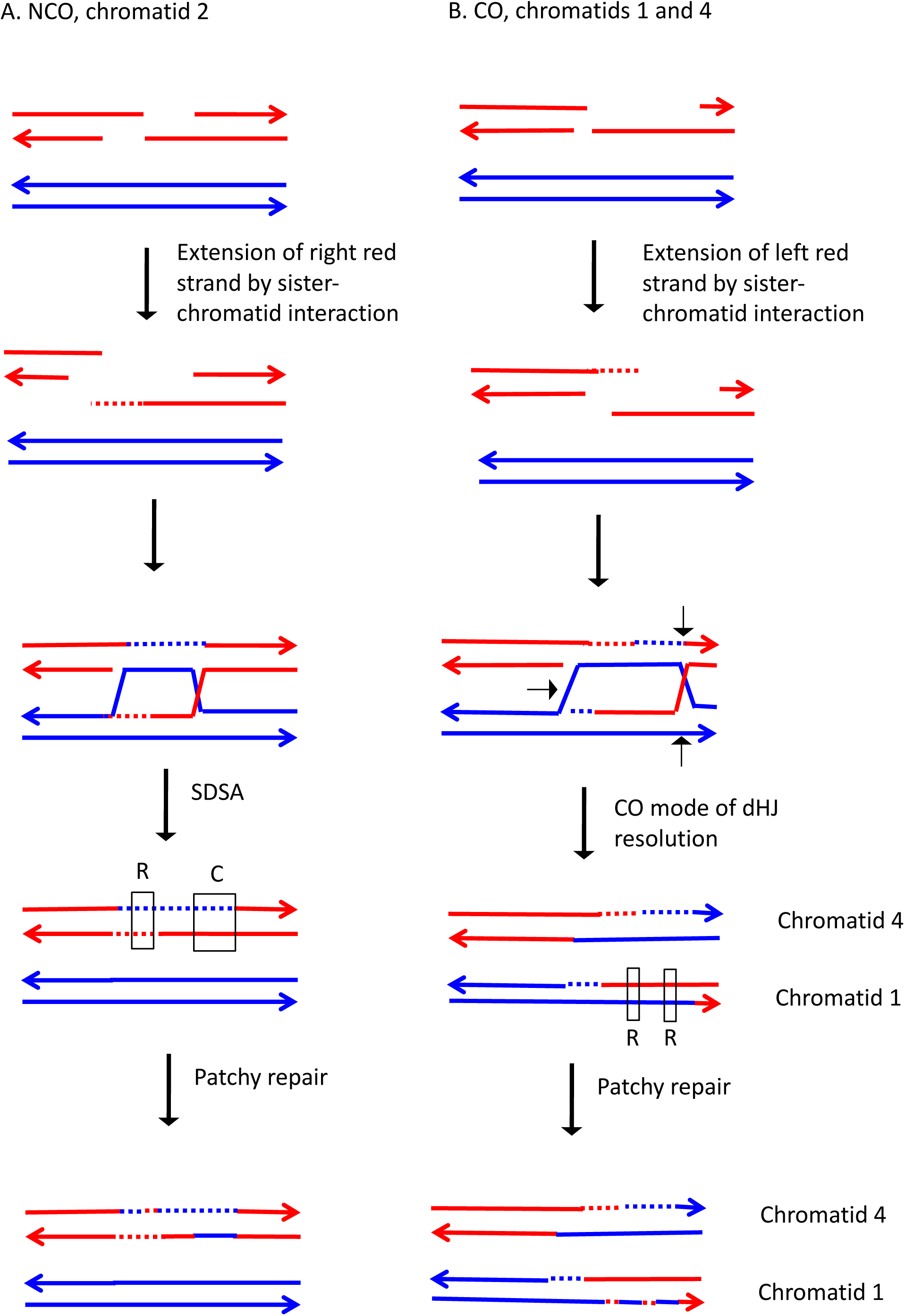

**Fig. S116.**
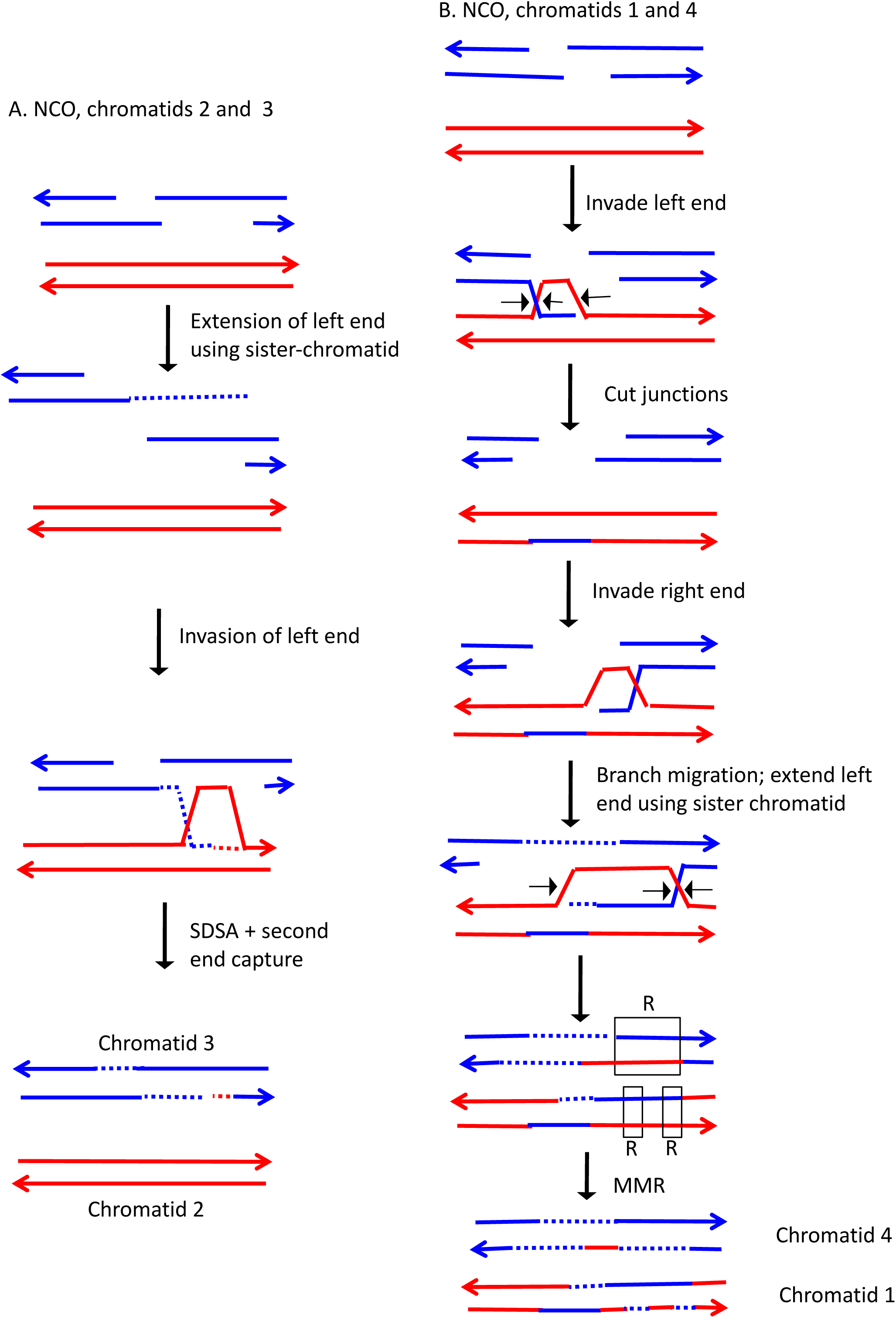

**Fig. S117.**
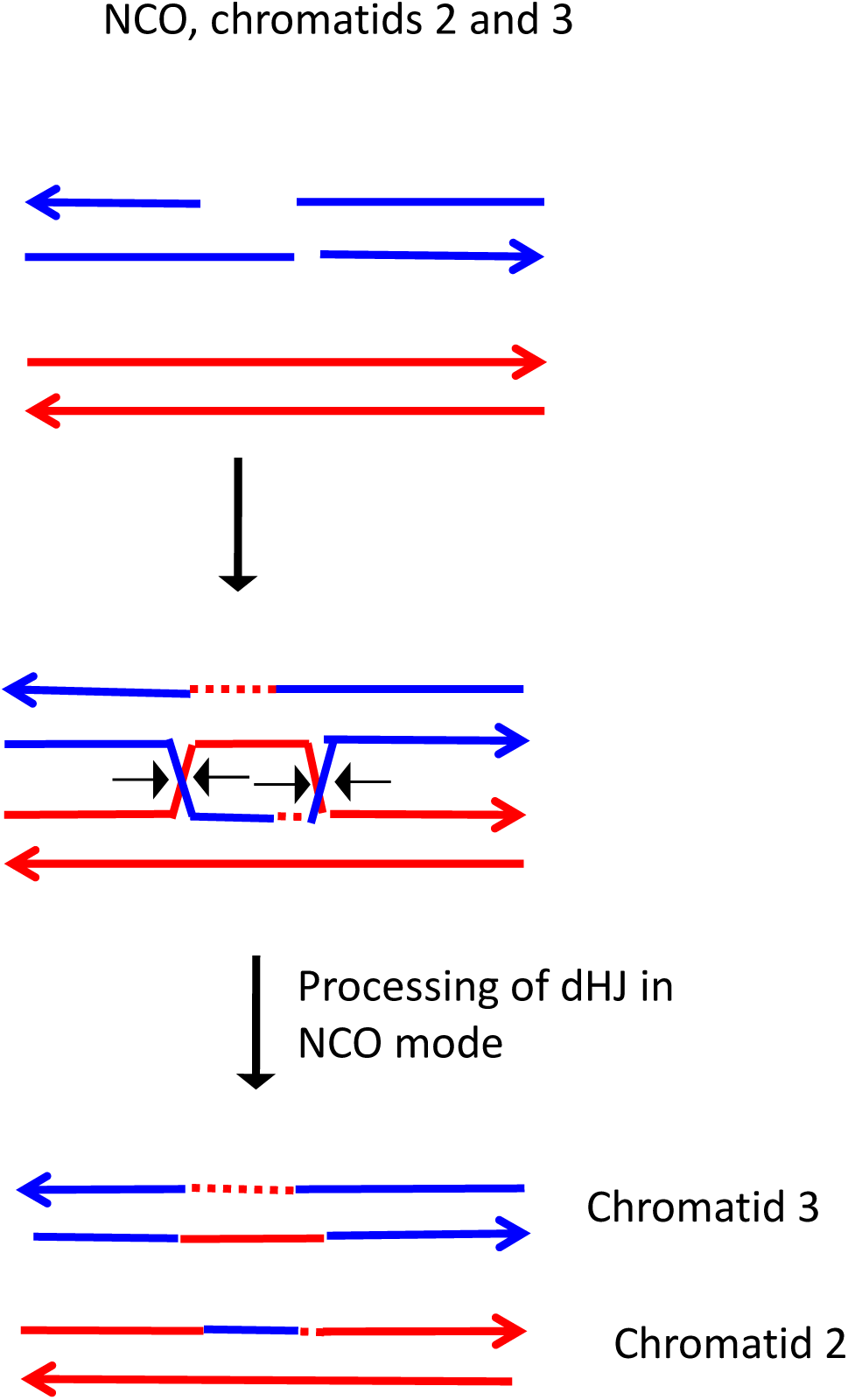

**Fig. S118.**
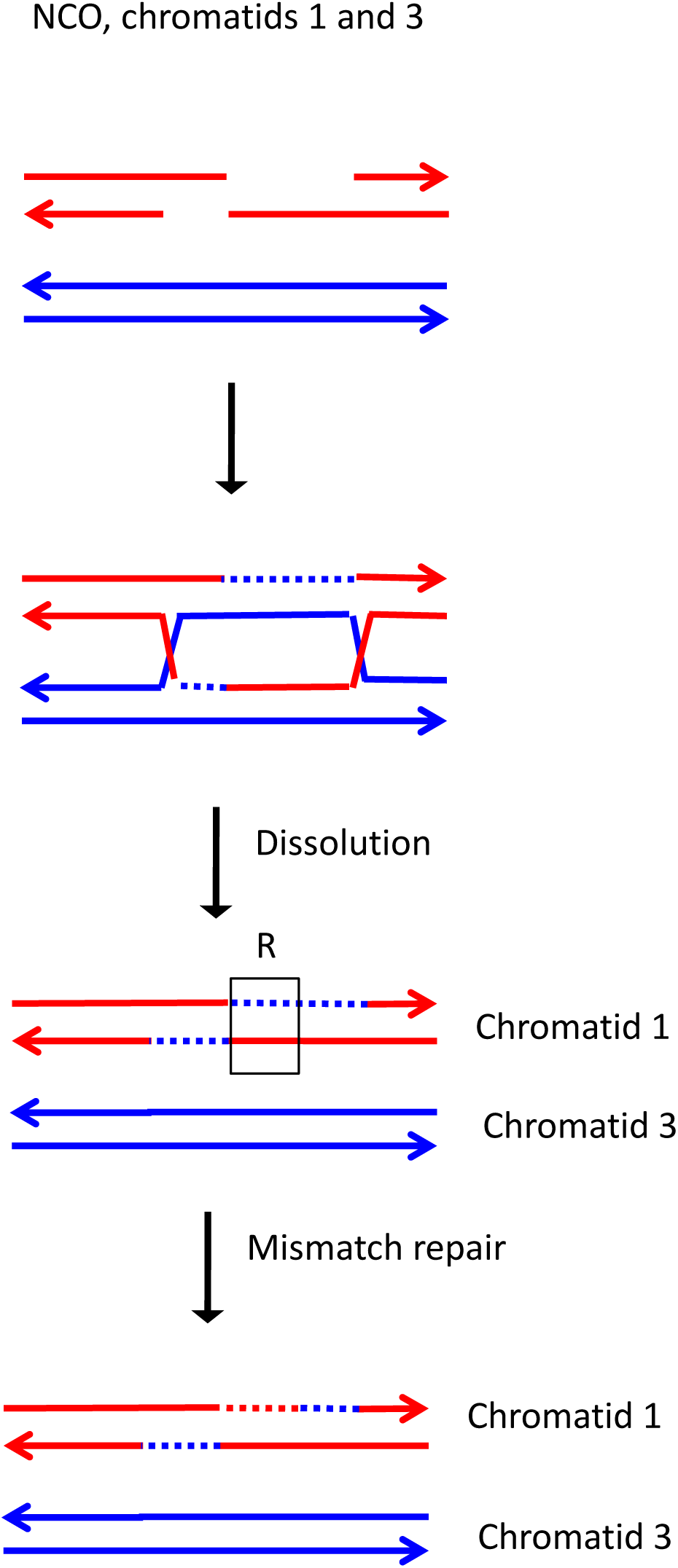

**Fig. S119.**
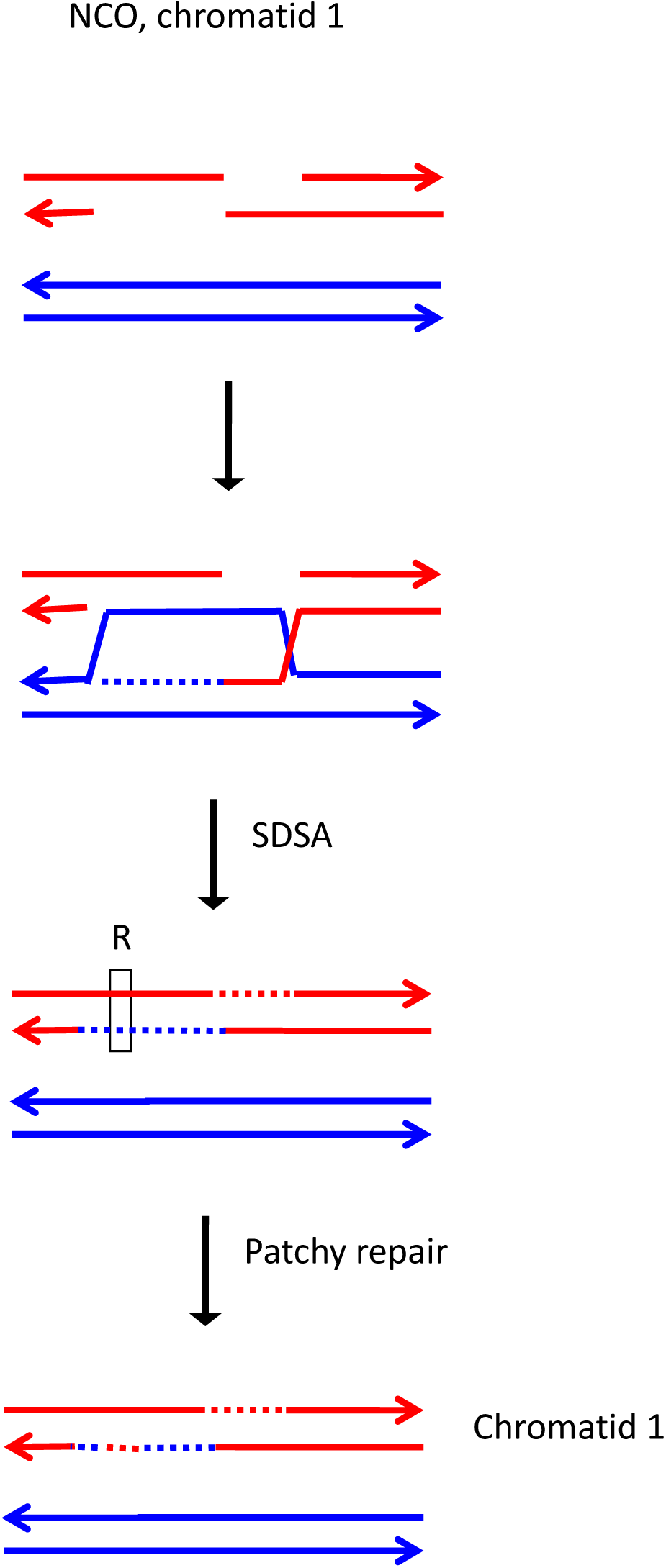

**Fig. S120.**
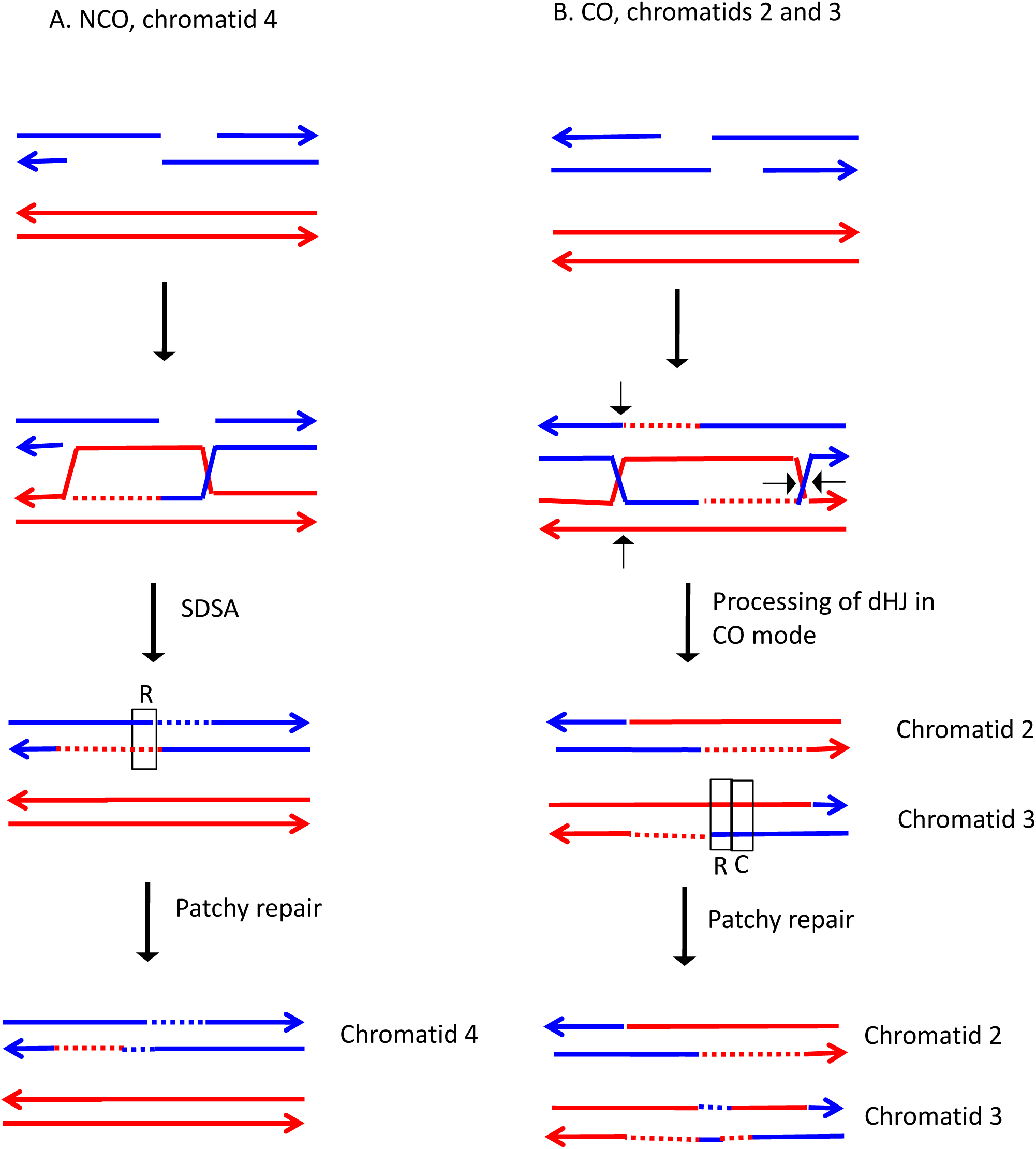

**Fig. S121.**
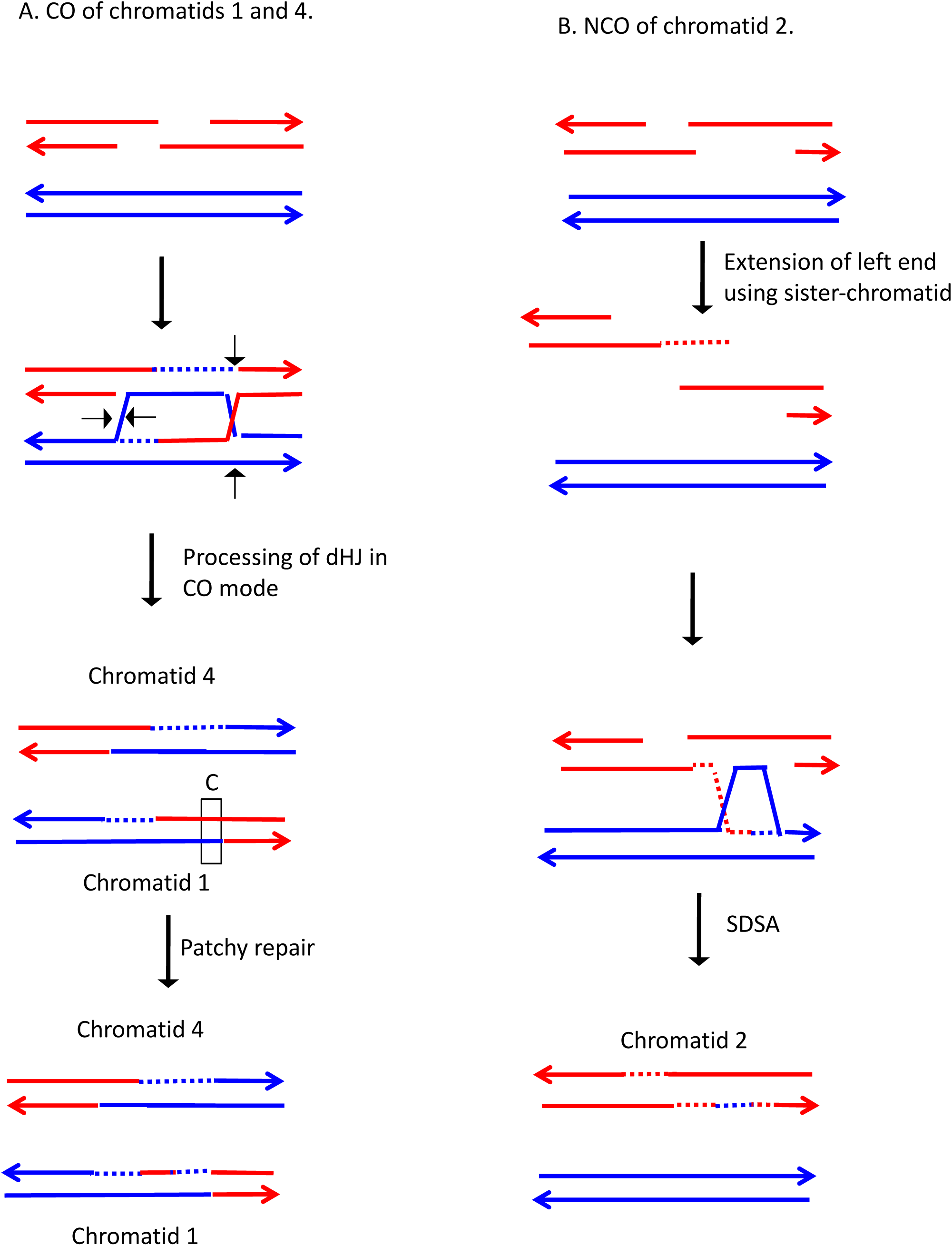

**Fig. S122.**
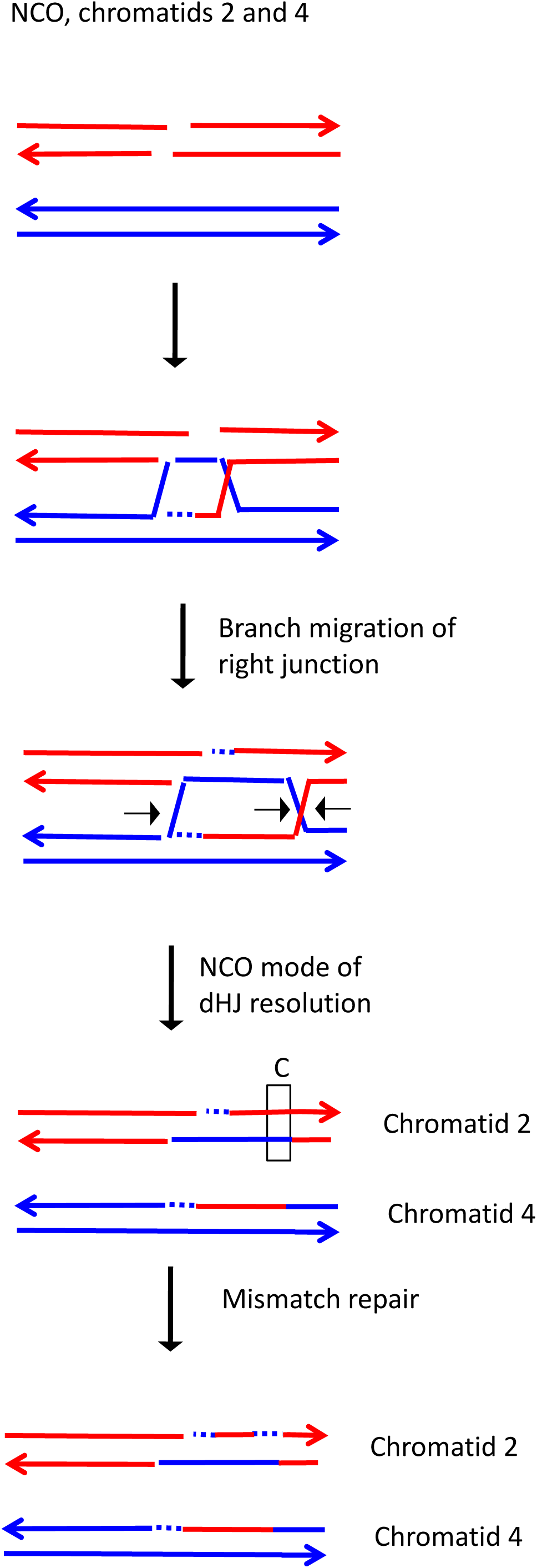

**Fig. S123.**
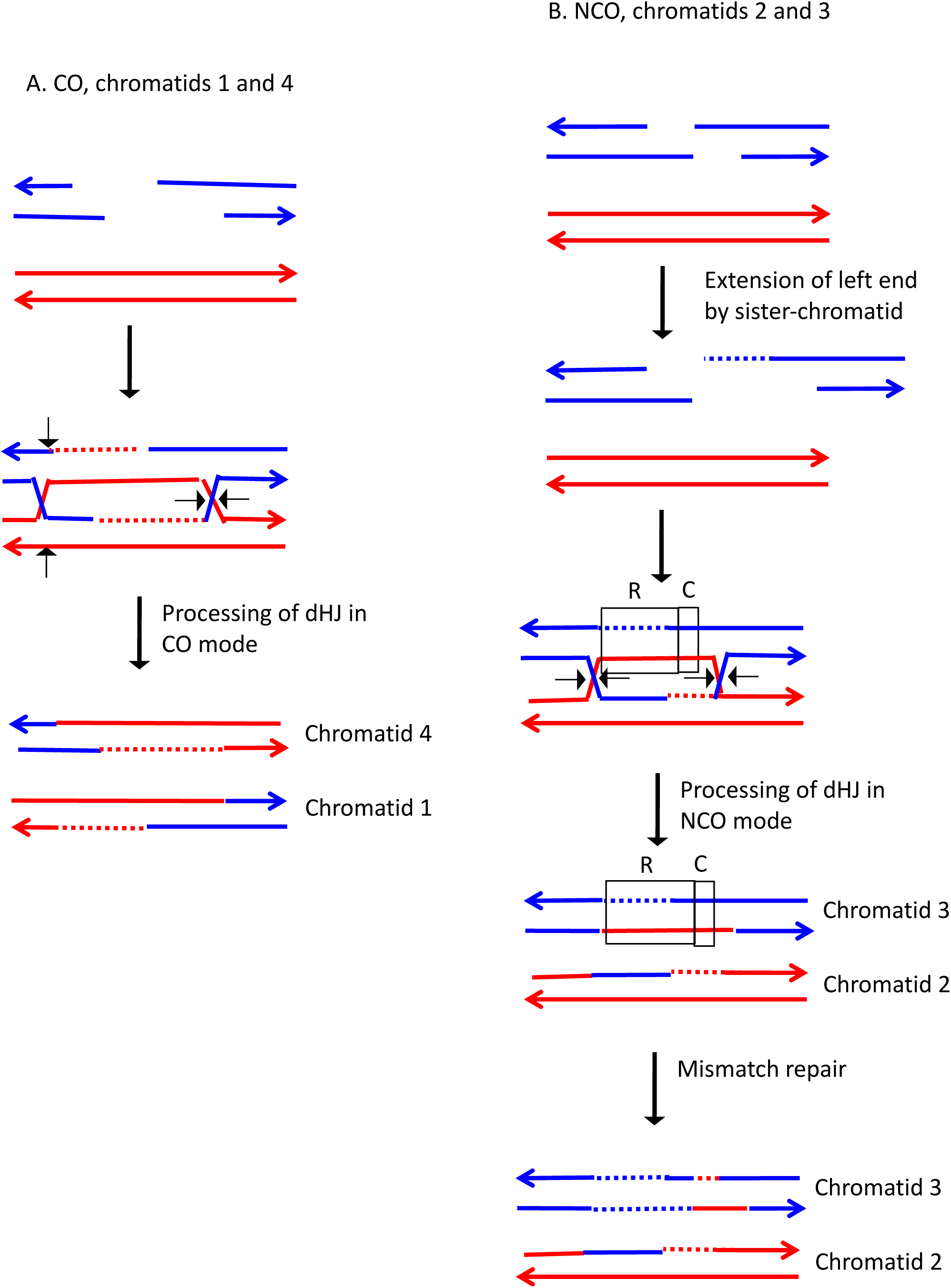

**Fig. S124.**
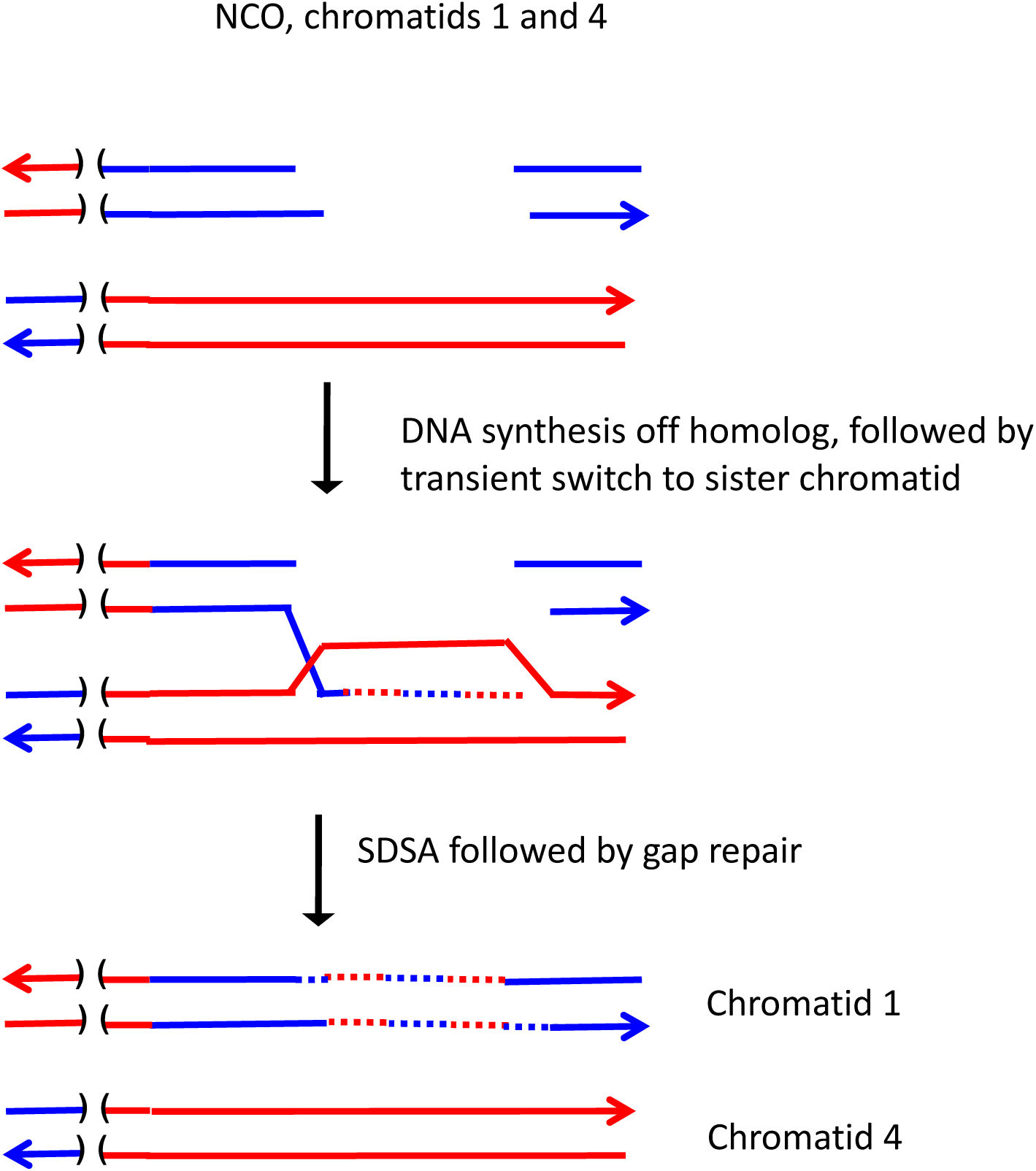

**Fig. S125.**
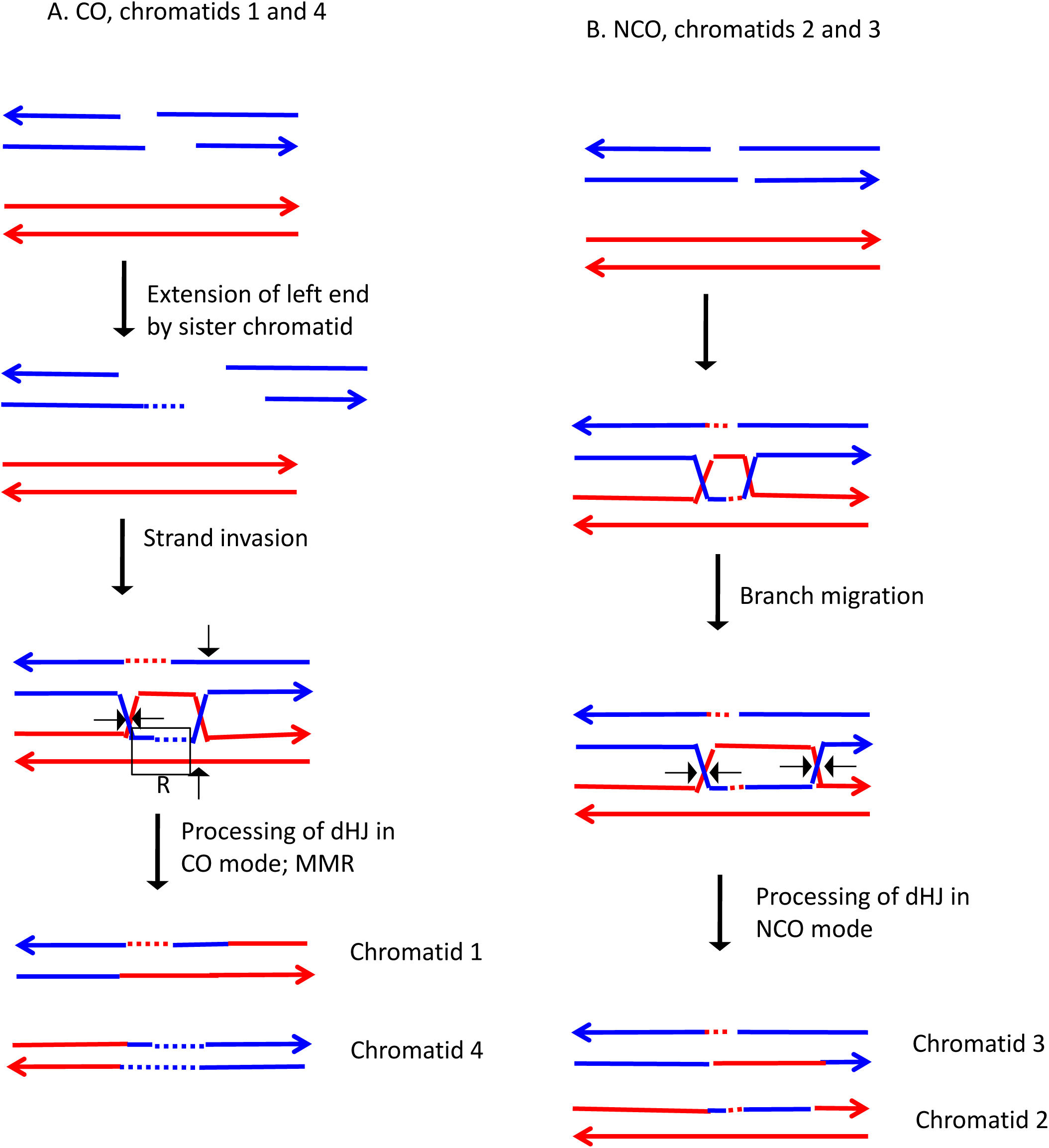

**Fig. S126.**
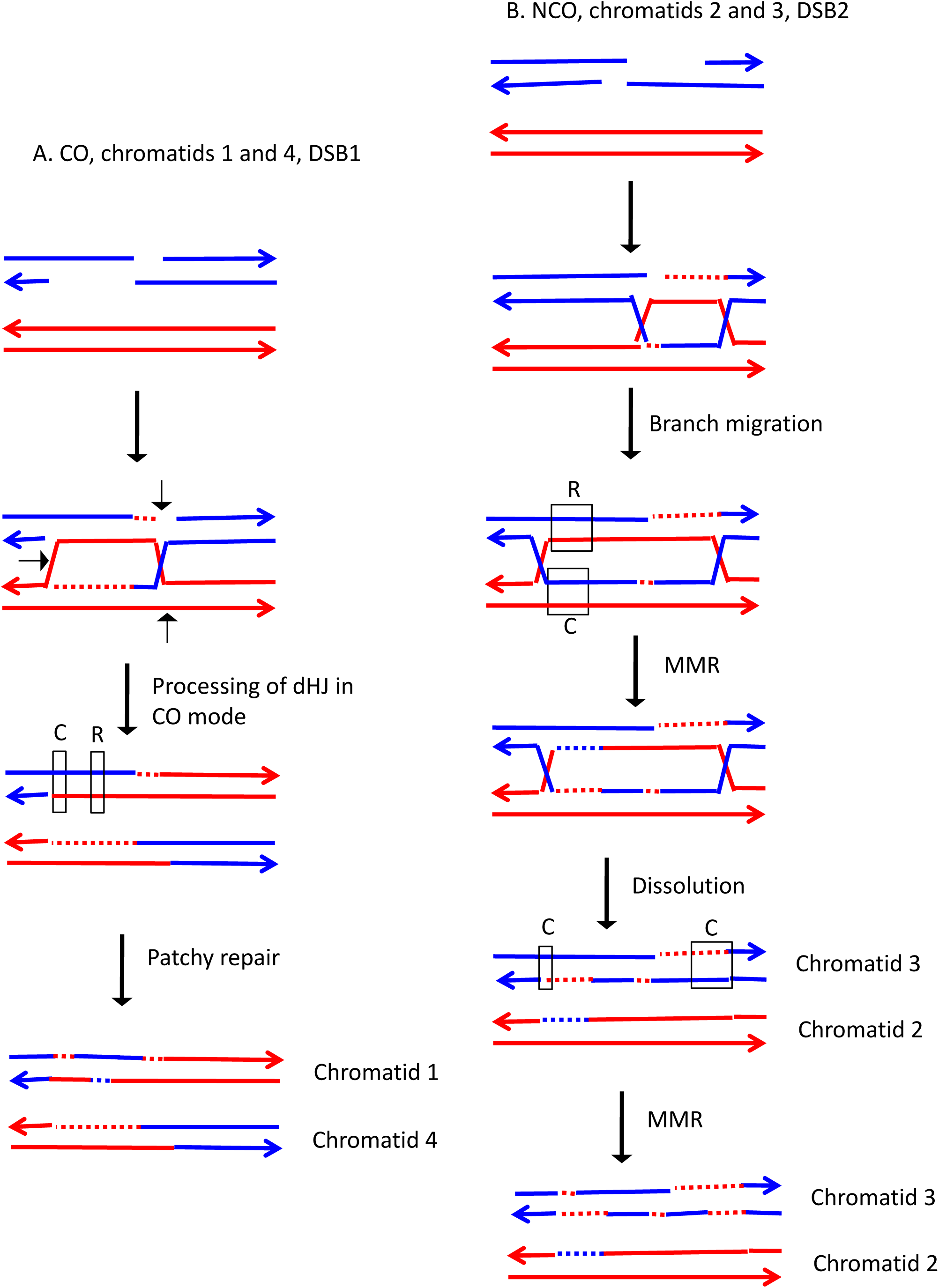

**Fig. S127.**
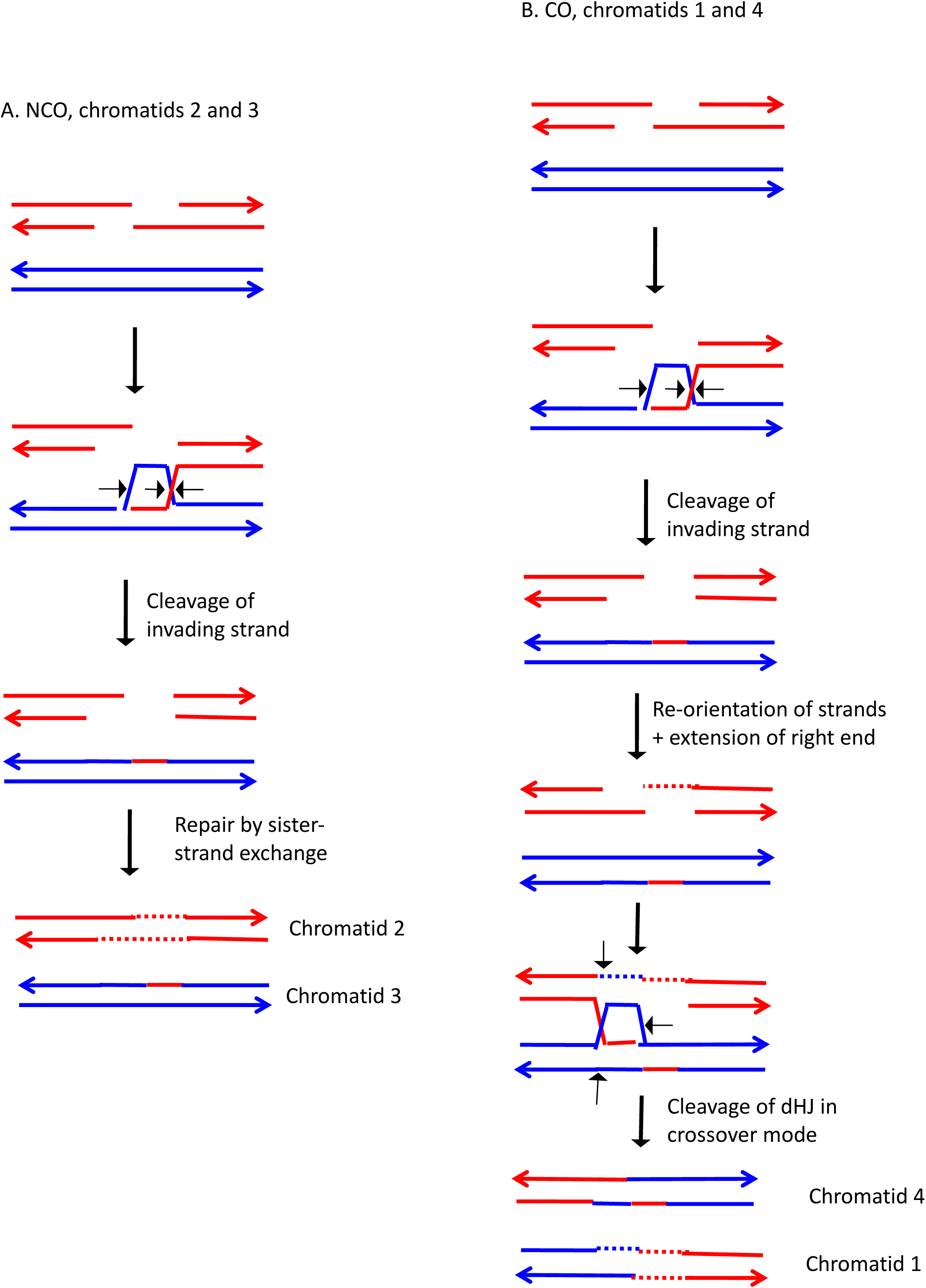

**Fig. S128.**
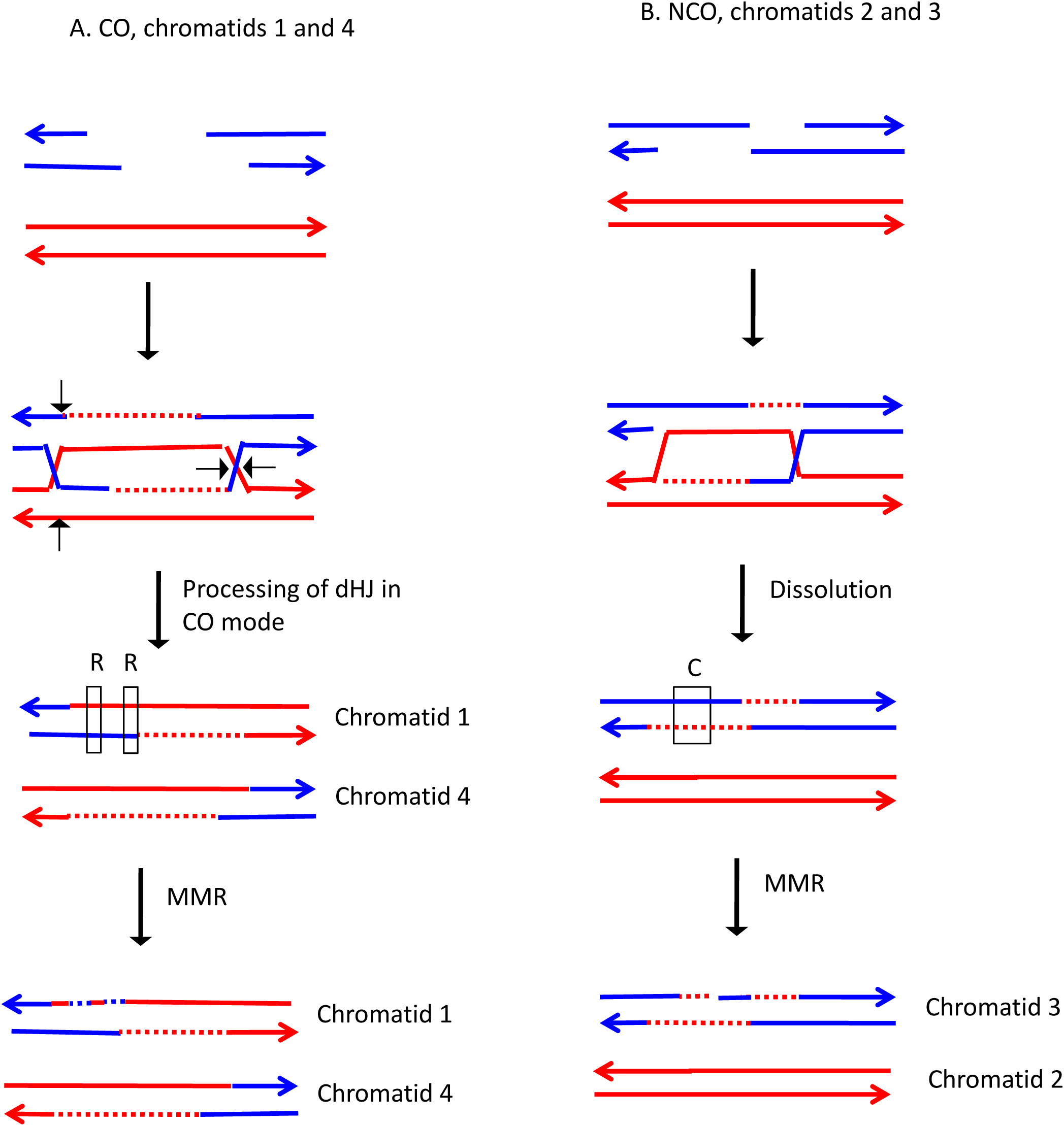

**Fig. S129.**
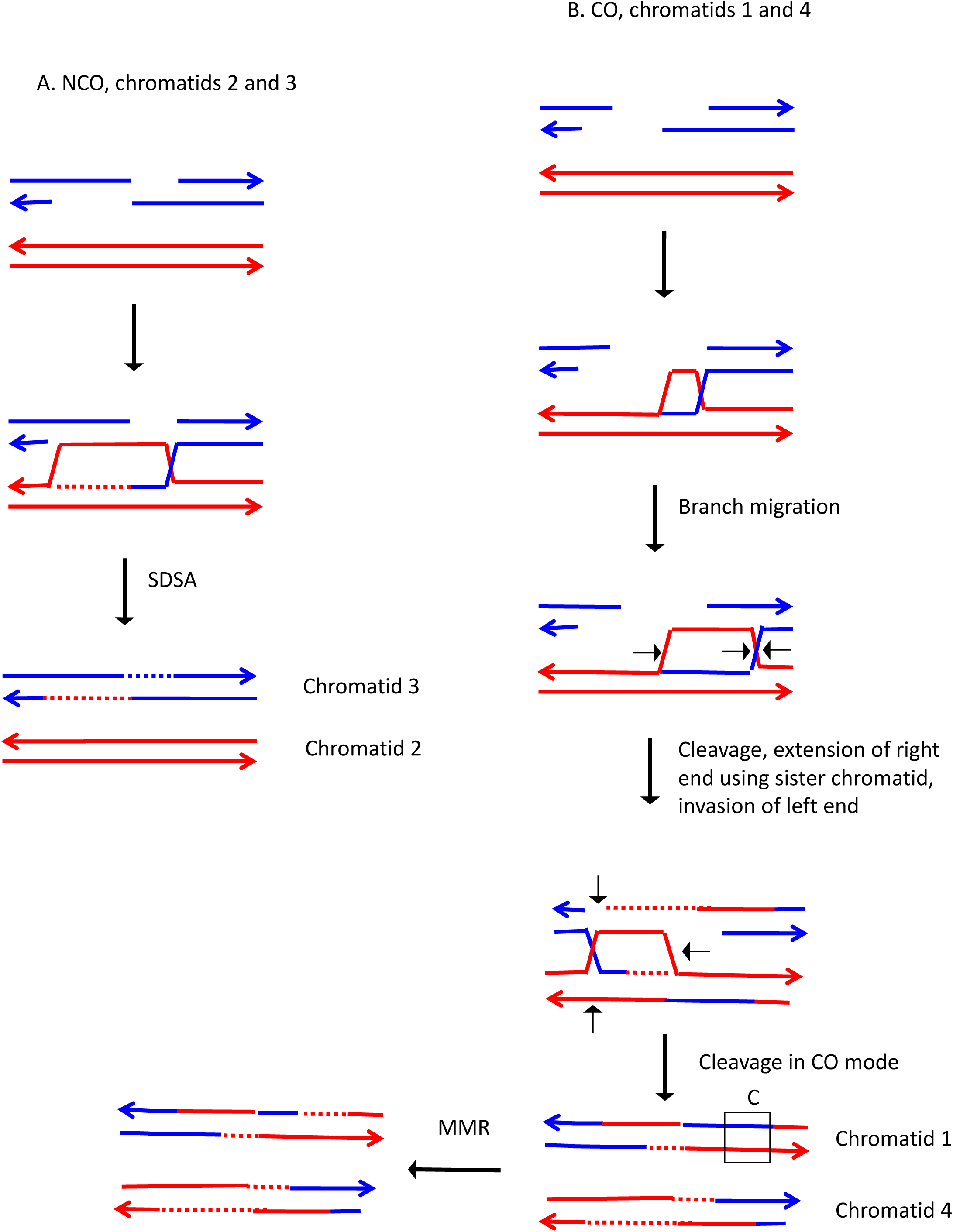

**Fig. S130.**
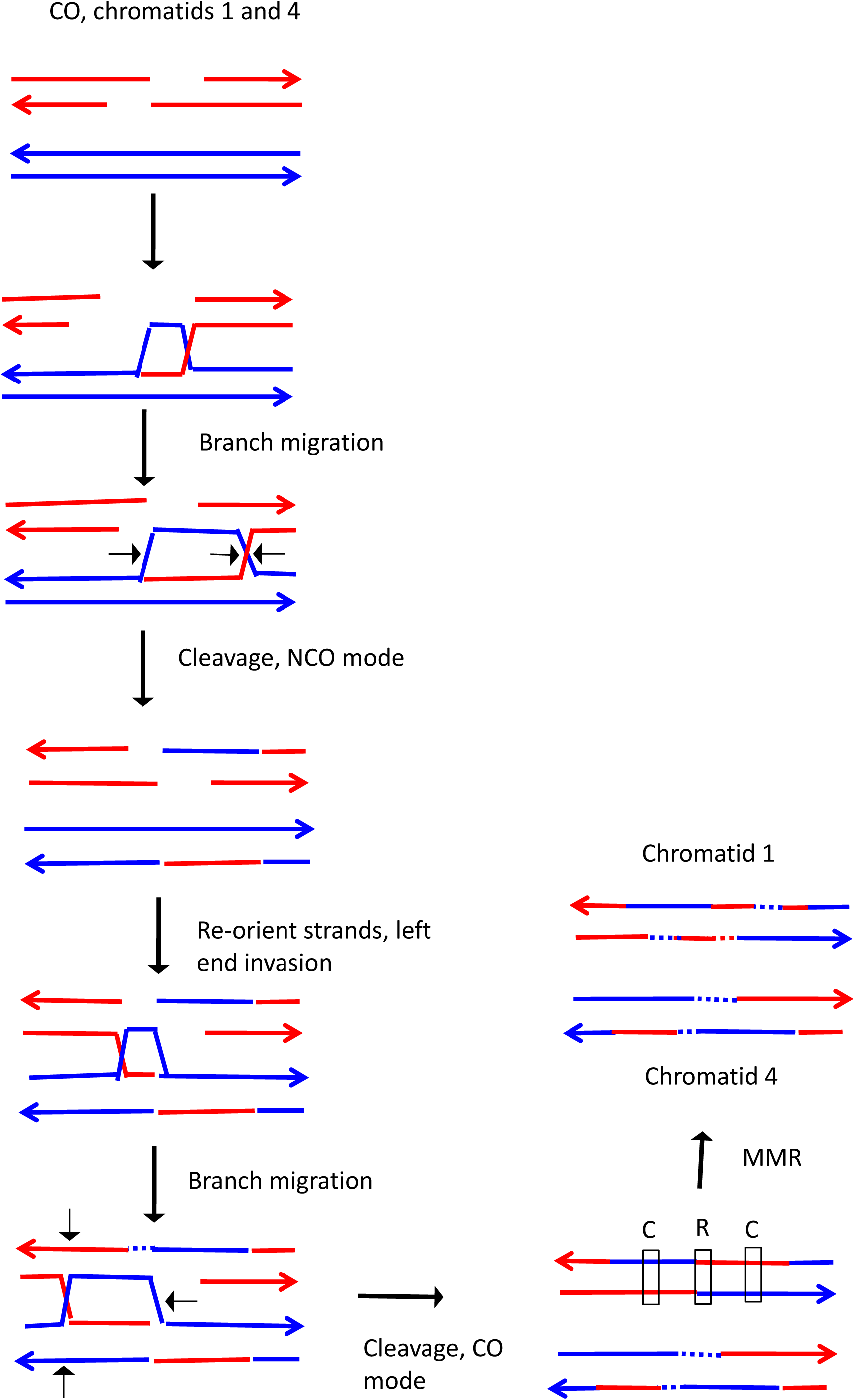

**Fig. S131.**
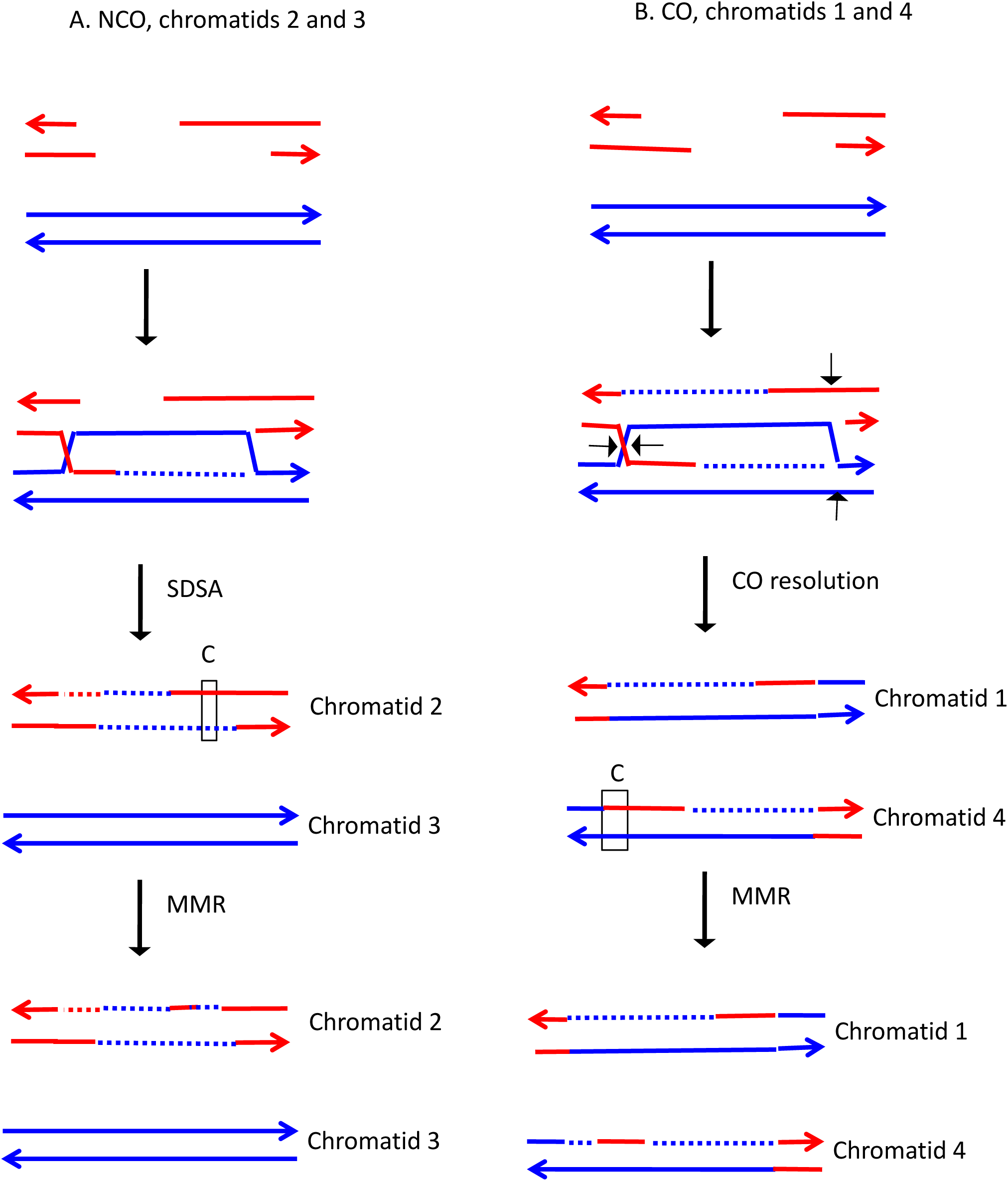

**Fig. S132.**
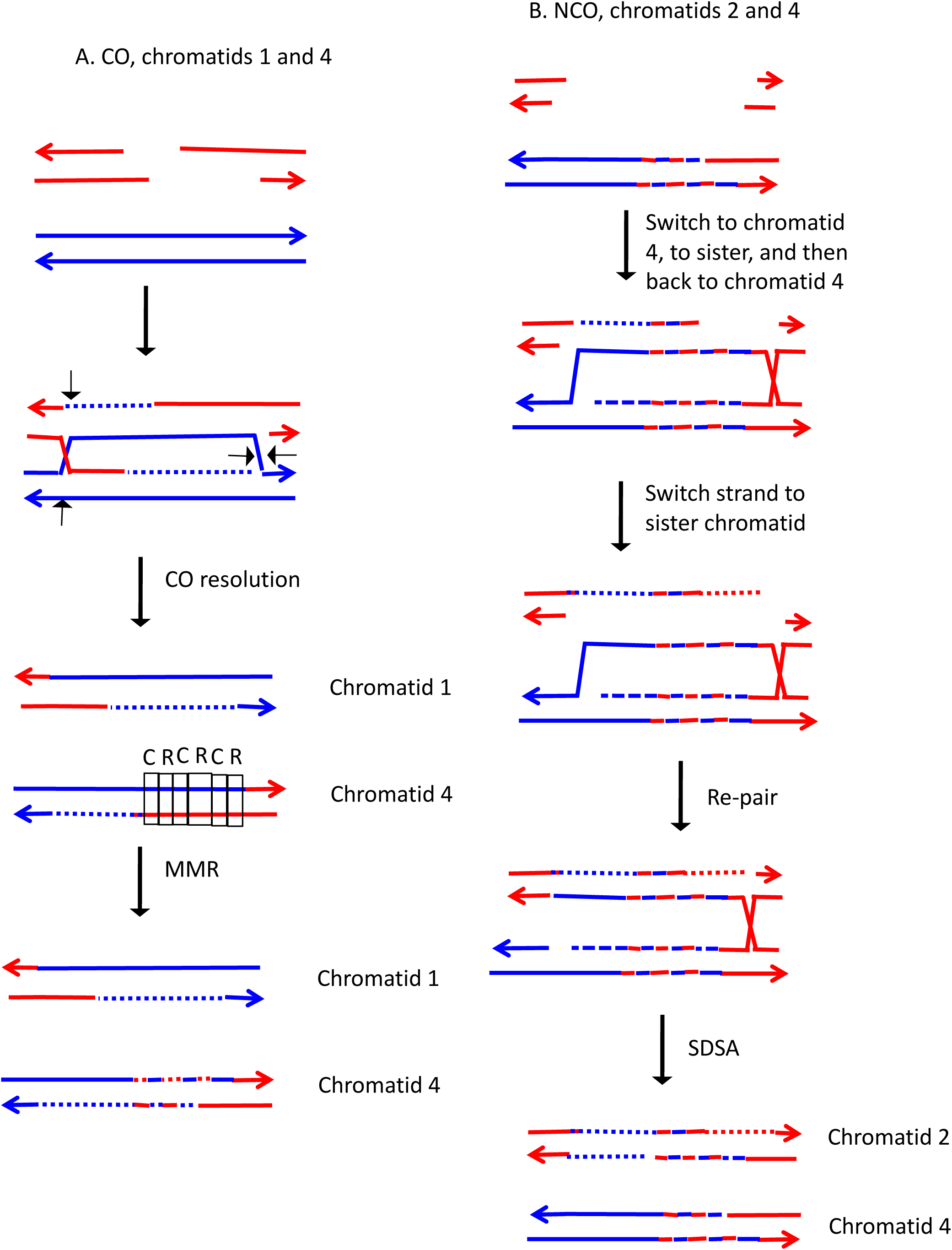

**Fig. S133.**
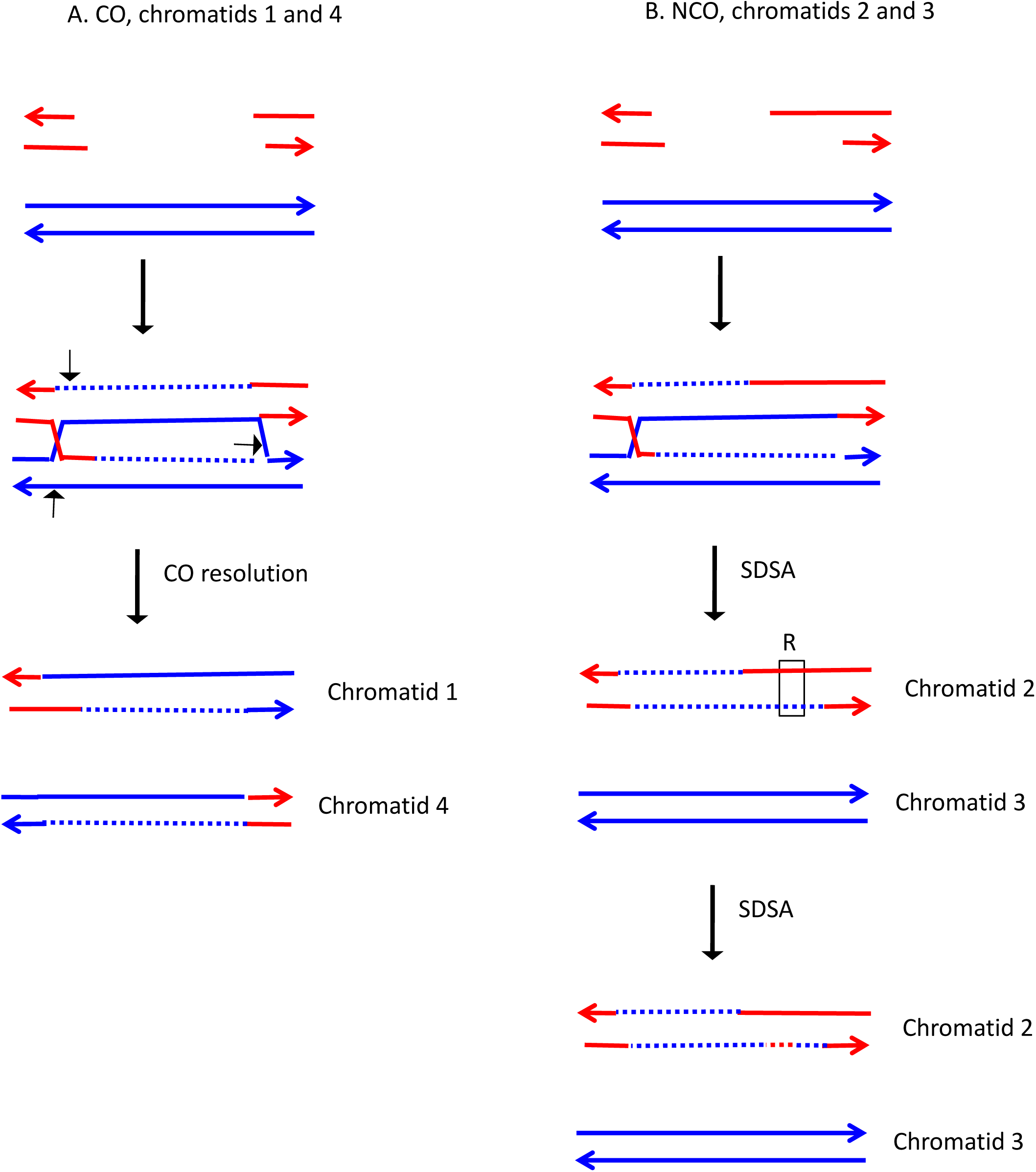

**Fig. S134.**
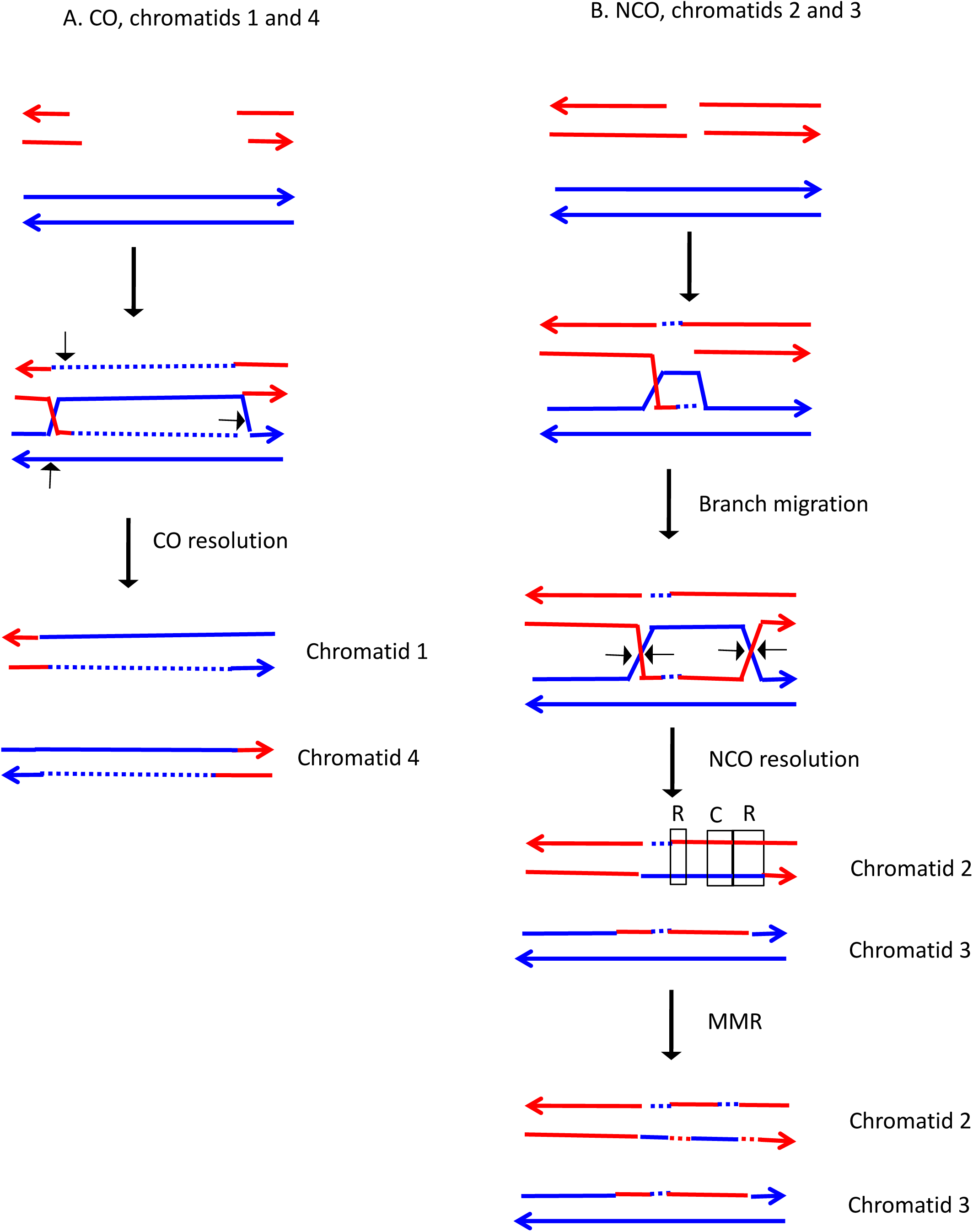

**Fig. S135.**
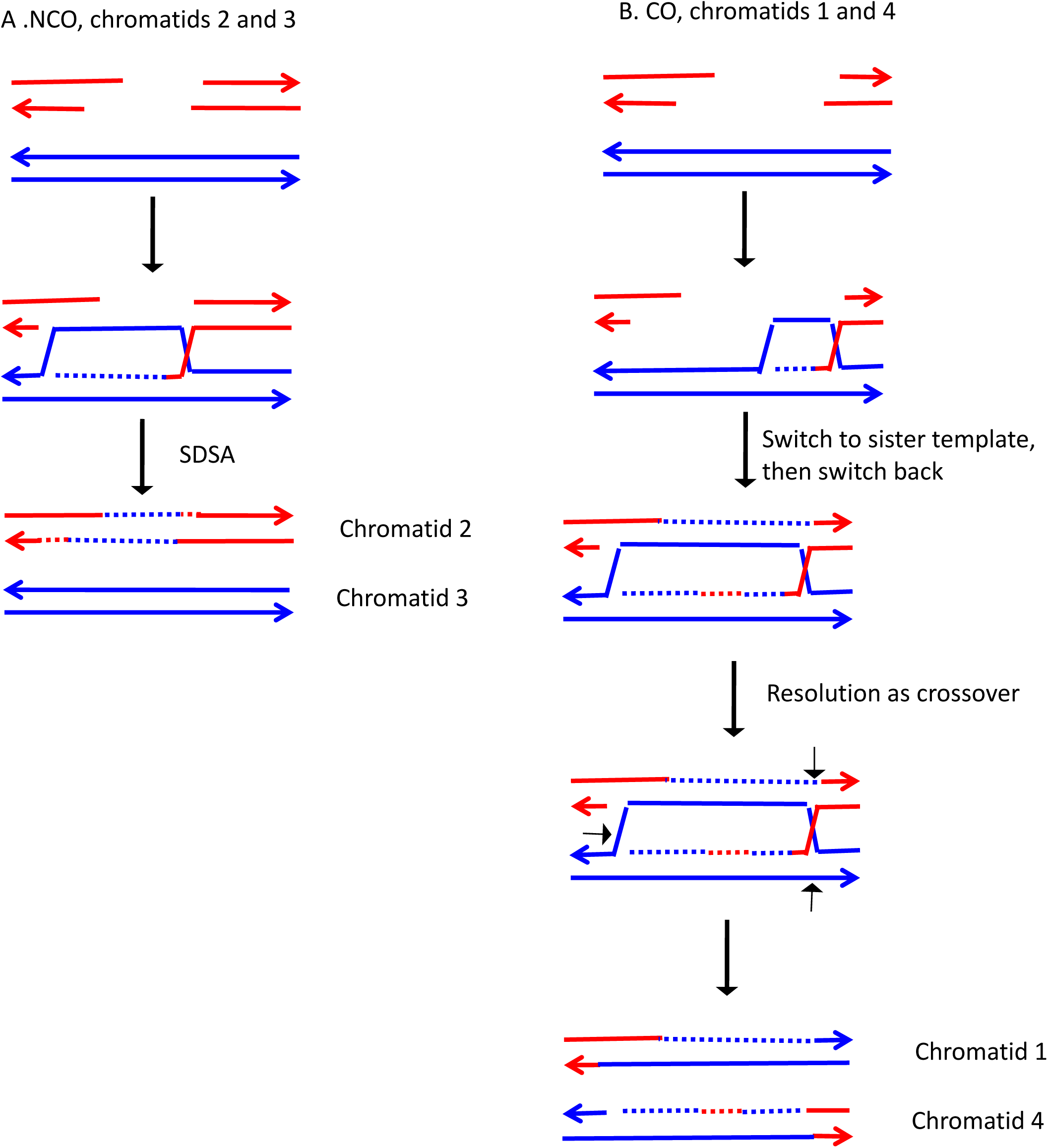

